# Epiphytic and endophytic microbiome of the seagrass *Zostera marina*: Do they contribute to pathogen reduction in seawater?

**DOI:** 10.1101/2023.08.21.554113

**Authors:** Deniz Tasdemir, Silvia Scarpato, Caroline Utermann-Thüsing, Timo Jensen, Martina Blümel, Arlette Wenzel-Storjohann, Claudia Welsch, Vivien Anne Echelmeyer

## Abstract

Seagrass ecosystems provide crucial ecosystem services for coastal environments and were shown to reduce the abundance of pathogens linked to infections in humans and marine organisms. Among several potential drivers, seagrass phenolics released into seawater have been suggested to play role in pathogen suppression, but the potential involvement of the seagrass microbiome in such effect has not been studied. Here we hypothesized that the microbiome of the eelgrass *Zostera marina*, especially the leaf epiphytes that are at direct interface between the seagrass host and surrounding seawater, inhibit such pathogenic microorganisms, hence, contribute to their suppression. Using a culture-dependent approach, we isolated 88 bacteria and fungi associated with the surfaces and inner tissues of the eelgrass leaves (healthy and decaying) and the roots, plus 19 strains from surrounding seawater and sediment. We first assessed the broad-spectrum antibiotic activity of microbial extracts against a large panel of common aquatic, human (fecal) and plant pathogens, and finally mined the metabolome of 88 most active extracts. The healthy leaf epibiotic bacteria, particularly *Streptomyces* sp. strain 131, displayed broad-spectrum and potent antibiotic activity superior to some control drugs. Gram-negative bacteria abundant on healthy leaf surfaces, and few endosphere-associated bacteria and fungi also showed remarkable antimicrobial activity. UPLC-MS/MS-based massive untargeted metabolomics analyses showed the rich specialized metabolite repertoire of strains with low annotation rates, indicating the presence of many undescribed antimicrobials in the extracts. This study contributes to our current understanding on microbial and chemical ecology of seagrasses, implying potential involvement of the seagrass microbiome, especially the leaf epiphytes, in reduction of pathogen load in seawater. Such antibiotic activity is not only beneficial for the health of ocean, human and aquaculture sector, especially in the context of climate change that is expected to exacerbate all infectious diseases, but may also assist seagrass conservation and management strategies.

## 1. Introduction

Microorganisms are highly abundant in seawater. Estimated number of bacteria in the seawater ranges from 10^4^ to 10^6^ cells per milliliter (Sunagawa et al., 2015) with a varying composition determined by biotic and abiotic processes. Seawater associated microbial communities can be beneficial, saprophytic, commensal, or pathogenic. Waterborne pathogens have diverse origins, i) some belong to the natural microbiota of aquatic environments, e.g., ubiquitous marine *Vibrio* spp., ii) some shed by humans (e.g., fecal enterococci, *Escherichia coli*), or iii) introduced naturally e.g., by wind, freshwater inflow, rainwater runoff or riverbank soils (Ortega et al., 2009). However, anthropogenic effects still count the main source of pathogens in the coastal zones. Urbanization, tourism, litter, wastewater pollution, leakage from sewage systems and agricultural or urban runoff constantly introduce pollutants, allochthonous organic matter, dust particles and various microbes pathogenic to humans, plants or animals into coastal waters (Orel et al., 2022). This has multi-faceted impacts such as negatively affecting the quality of water and ecosystem health, thus leading to eutrophication (Boesch et al., 2001), decreasing dissolved oxygen rates (Breitburg et al., 2018), or triggering infections or mass mortality of ecologically and economically important marine species (Orel et al., 2022). It is a growing health concern that some marine pathogens (e.g., *Vibrio* spp.) are transmitted to humans through recreational activities (swimming, sea bathing) or consumption of raw or undercooked seafood (Shuval, 2003; Cavicchioli et al., 2019). Also, climate change related challenges, e.g., elevated water temperatures, are expected to increase the load of pathogens such as *Vibrio* spp. in coastal areas, leading to increased exposure to pathogens and global disease outbreak risks to affect both ocean and human health, as well as aquaculture industry (Cavicchioli et al., 2019; de los Santos et al., 2020). Another foreseen impact of the climate change is the higher incidence of other infectious human diseases and higher rate of antibiotic resistance of some human pathogens (MacFadden et al., 2018; Semenza and Paz, 2021).

Seagrasses are marine plants (angiosperms) that form extensive underwater meadows in temperate coastal ecosystems and represent a widespread foundation species. They provide vital habitats and a nursery ground for many species, including economically important fish species (Jeyabaskaran et al., 2018; Conte et al., 2021). Being key benthic ecosystem engineers, seagrasses stabilize soft sediments and store large quantities of CO_2_ as blue carbon, contributing to the mitigation of anthropogenic emissions (Duarte et al., 2010; de Los Santos et al., 2019). An additional and recently discovered ecosystem service of seagrass meadows is the suppression of pathogenic bacteria in the water column. In a pioneering study, Lamb et al. (2017) showed that mixed seagrass meadows effectively reduced the abundance of human fecal bacteria (enterococci) and a variety of marine pathogens (e.g., *Vibrio* spp.) that are harmful for fish, marine invertebrates and marine mammals in polluted Indonesian coastal waters. The authors investigated over 8,000 reef-building corals and reported two-fold lower levels of disease on reefs with adjacent seagrass meadows than on reefs without adjacent seagrass meadows (Lamb et al., 2017). Several subsequent studies carried out on seagrass meadows in other parts of the world also confirmed lower levels of pathogens in their vicinity, including the fecal pathogens *E. coli* and various enterococci (Palazón et al., 2018), *Salmonella* spp. (Deng et al., 2021) and *Vibrio* spp. (Reusch et al., 2021). The underlying reason(s) for this ‘sanitary’ effect is currently unclear but it has been partly linked to the high content of seagrass phenolics (e.g., flavonoids and other phenylpropanoids) that are well-known for their antimicrobial effects and involvement in seagrass chemical defense (Migliore et al., 2007; Guan et al., 2017; Papazian et al., 2019; Mannino and Micheli, 2020; Conte et al., 2021, Deng et al., 2021).

Seagrasses harbor complex assemblies of microorganisms on their surfaces and in their inner tissues, and live intimately with both beneficial and harmful microorganisms in their environment (Hurtado-McCormick et al., 2019; Conte et al., 2021). Although some interactions between seagrasses and their surrounding sediment microbiota have been demonstrated (Bourque et al., 2015; Sun et al., 2015; Brodersen et al., 2018; Martin et al., 2018a and 2018b; Conte et al., 2021), the relationships between seagrass microbiome and the surrounding water column have largely remained unexplored. Here we investigated whether seagrass microbiomes, particularly the leaf epibionts that constitute a direct interface between the seagrass and the surrounding seawater, inhibit aquatic and other pathogens ‘introduced’ into the sea by natural and anthropogenic effects, thereby potentially contribute to the ‘sanitary’ effects of seagrasses. A culture-based approach allowed isolation and identification of 88 epiphytic and endophytic bacteria and fungi associated with the healthy leaves, decaying leaves and the roots of the Baltic eelgrass *Zostera marina*, plus 19 microbes deriving from the surrounding seawater and sediment. All 107 isolates were cultivated in two different growth media. Ethyl acetate (EtOAc) extracts of 214 microbial cultures were assessed for their antimicrobial effect against a panel of 28 pathogens comprised of aquatic, fecal, human and plant pathogens. The most active 88 extracts were investigated by an UPLC-MS/MS-based untargeted metabolomics approach to identify their chemical constituents that may contribute to the observed antimicrobial effects. We show both fungal and bacterial isolates, particularly those originating from the healthy leaf surfaces (HLS) of *Z. marina*, to have significant antipathogenic effects and rich secondary metabolomes, suggesting their potential involvement in the eelgrass ecosystem services.

## 2. Materials and methods

### 2.1. Collection of Zostera marina

Plant material was collected at Falckenstein Beach, Kiel Fjord, Baltic Sea (N 54°23’37.5, E 10°11’23.3) in August 2017. The environmental parameters, i.e., seawater temperature (17 °C), pH (pH 7, determined by pH-Fix indicator strips, Carl Roth GmbH, Karlsruhe, Germany) and salinity (13.3 psu, determined by conductivity measurement using a fast protein liquid chromatography system, GE ÄKTA Purifier FPLC System, GE Healthcare, Braunschweig, Germany) were measured. Several eelgrass plants each were collected from several spots of the same seagrass meadow by snorkeling (approx. 1.5 m depth). Ambient seawater and surface sediment in immediate vicinity of the sampled seagrass shoots were collected as reference. All samples were transported in a cooler box to laboratory and processed on the same day.

### 2.2. Isolation of Z. marina associated microbes

To isolate a broad bacterial and fungal diversity, nine different agar media were used for isolation of microorganisms from *Z. marina,* ambient seawater and sediment samples. We used six standard media for cultivation of marine-derived microorganisms: Marine Broth Agar (MA; 3.74% Marine Broth 2216, Becton Dickinson, Sparks, MD, USA; 1.5% agar bacteriology grade, AppliChem, Darmstadt, Germany), Potato-Dextrose-Agar (PDA; Oppong-Danquah et al., 2018 with 2% glucose instead of 0.4%), modified Wickerham medium (WSP30; adjusted to pH 5.8, 3% Instant Ocean, Blacksburg, VA, USA; Silber et al., 2013), modified Wickerham medium prepared with Baltic seawater instead of Instant Ocean (WSP+BSW), Tryptic Soy Broth agar (TSB3+10; Utermann et al., 2020) and TSB agar prepared with Baltic seawater instead of sodium chloride (TSB+BSW). In addition, modified Melin-Norkrans-Agar was used for cultivation of fungi (https://www.uamh.ca/-/media/uamh/OrderCultures/Documents/Media.pdf). In order to mimic the natural conditions in the seagrass meadow, two “*Z. marina* media” for cultivation of bacteria (ZMB) and fungi (ZMF) were designed. For this aim, freeze-dried *Z. marina* leaves were ground into powder using a pulverisette (Pulverisette 14, Fritsch, Idar-Oberstein, Germany). 1% of the resulting seagrass powder and 1.5% agar were added to Baltic seawater (ZMB). For cultivation of fungi, a penicillin-streptomycin mixture (Gibco^TM^, Thermo Fisher Scientific, Dreieich, Germany) was added to the ZMF medium to prevent bacterial growth.

For inoculation, plant samples were divided into roots (R) and leaves (L). The leaves were further split into healthy green (encoded as HL) and decaying leaves (DL). For inoculation of microbial epibionts from the plant surfaces (encoded by “S” for each plant organ, i.e., DLS, HLS, RS), two different methods were applied, namely surface swabbing and imprinting. For swabbing, roots, healthy and decaying leaves surfaces were wiped with a sterile cotton swab, placed in sterile saline (1.8% NaCl) and vortexed for 5 min. For imprinting, leaves and roots were cut into 1 cm pieces, the surface was wiped with a circular movement on the agar surface and finally left on the agar plate. For isolating endo-microbiota (encoded by “I” (for inner) for each plant tissue, i.e., DLI, HLI, RI), surface disinfection was performed using 70% EtOH. Subsequently, the surface sterilized tissue pieces of approximately 1 cm length were ground with a sterile pestle in a reaction tube containing sterile saline. The homogenized material was vortexed for 5 min and 100 µl were plated on the different media in two dilutions (undiluted, 1:10).

Sediment samples were resuspended in sterile saline and mixed for 5 min before plating 100 µL suspension (undiluted and 1:10 dilutions) on agar plates. Undiluted seawater samples were plated using two different volumes (100 µL, 500 µL). Samples were incubated in the dark at 22°C. Morphological different colonies were picked after 7 and 21 days of incubation. Colonies were transferred to fresh agar plates until pure cultures were obtained. All strains were cryopreserved at −80°C using the Microbank^TM^ system (Pro-Lab Diagnostics, Richmond Hill, ON, Canada).

### 2.3. Molecular identification of isolates

DNA extraction of pure bacterial strains was done by an established freeze-and-thaw procedure, while DNA from fungal cultures was obtained by mechanical disruption (Utermann et al., 2018). If initial DNA extraction failed, the extraction process was repeated with the DNeasy Plant Mini Kit (Qiagen, Hilden, Germany; Utermann et al., 2020). Amplification of bacterial and fungal DNA was performed with universal primers targeting the 16S or 18S rRNA gene or the ITS1-5.8S-ITS2 region by using established standard protocols (Utermann et al., 2018; Utermann et al., 2020). Successfully amplified DNA fragments were sequenced (Sanger et al., 1977) at IKMB (Institute of Clinical Molecular Biology, Kiel University, Kiel, Germany) or LGC Genomics GmbH (Berlin, Germany) using primers 1387R or Eub27F for the bacterial 16S rRNA gene and ITS1 for the fungal ITS fragment. Sequences were trimmed by using ChromasPro V1.33 (Technelysium Pty Ltd, South Brisbane, Australia) and generated FASTA files were submitted to BLAST (Basic Local Alignment Search Tool; Altschul et al., 1990) at NCBI (National Center for Biotechnology Information). DNA sequences are available under the accession numbers OR400253-OR400328 (bacteria) and OR400216-OR400246 (fungi) at NCBI GenBank.

### 2.4. Cultivation and extraction of isolated microorganisms

Selection for cultivation and extraction was based on phylogeny to reflect a broad phylogenetic diversity, and on safety level (exclusion of BSL-2 organisms). In total, 76 bacterial strains and 31 fungal strains were selected for cultivation (bacteria, Supplementary Table S2; fungi, Supplementary Table S3). To provide the microorganisms differential nutritional sources, two agar media each were selected for cultivation of fungal and bacterial strains. Fungal strains were cultivated on potato dextrose agar (PDA; potato infusion powder: 0.4%, D-glucose monohydrate: 2%) and casamino acids glucose medium (M34, casein hydrolysate: 0.25%, D-glucose monohydrate: 4%, magnesium sulfate: 0.01%, potassium dihydrogen phosphate: 0.18%). Bacterial strains were grown on MA and Glucose Yeast Malt Agar (GYM4; D-glucose monohydrate: 2%, malt extract: 0.4%, yeast extract: 0.4%, calcium carbonate: 0.2%). All cultivation media were supplied with 3.5% Instant Ocean (Aquarium systems, Sarrebourg, France) prior to adjusting the pH. Isolate pre-cultures were grown on plates with 1.5% Agar Bacteriology Grade (AppliChem); the respective main-culture plates for extraction contained 1.2% Agar Noble (Becton Dickinson).

Per pre-culture, three plates were inoculated using one bead per plate from the cryopreservation. For main cultures, eleven plates per medium were inoculated with cell material from the respective pre-cultures. The ten best grown plates were chosen for extraction. Fungal pre-and main cultures were grown for 14 d, while bacteria were grown for 7 d at 22 °C in under dark conditions. Main cultures were extracted with EtOAc (Pestinorm grade ≥ 99,8%; VWR, Leuven, Belgium) and water as described by Utermann et al. (2021). Organic crude extracts were stored at −20 °C until further use.

### 2.5. Antimicrobial activity assays

Supplementary Table S1 displays the list of all pathogens used in this study. All bacterial and fungal test strains were purchased from Leibniz Institute DSMZ (Braunschweig, Germany), Institut Pasteur (Paris, France) or Westerdijk Fungal Biodiversity Institute (Utrecht, Netherlands). Assays were performed in 96-well microplates at the initial test concentration of 100 µg/ml. Extracts inhibiting >50% of the pathogen growth were subjected to IC_50_ determinations.

#### 2.5.1. Assays against pathogenic bacteria

The extracts were tested against 28 bacterial and fungal pathogens divided into four panels reflecting i) 16 aquatic pathogens, ii) 5 human fecal pathogens, iii) 1 human pathogen (MRSA), and iv) 6 phytopathogens. The aquatic panel includes the bacteria *Algicola bacteriolytica* CIP 105725, *Lactococcus garvieae* DSM 20684, *Leifsonia aquatica* DSM 20146, *Pseudoalteromonas elyakovii* CIP 105338, *Shewanella algae* DSM 9167, *Vibrio aestuarianus* DSM 19606, *Vibrio alginolyticus* DSM 2171, *Vibrio anguillarum* DSM 21597, *Vibrio cholerae* DSM 100200, *Vibrio coralliilyticus* DSM 19607, *Aliivibrio fischeri* DSM 507, *Vibrio harveyi* DSM19623, *Vibrio ichthyoenteri* DSM 14397, *Vibrio parahaemolyticus* DSM 11058, *Vibrio splendidus* DSM and *Vibrio vulnificus* DSM 10143. All strains were cultivated in MB medium (0.5% peptone, Becton Dickinson; 0.1% yeast extract, Merck, Darmstadt, Germany; 3% artificial sea salt Instant Ocean) except *L. garvieae*, *L. aquatica* and *V. vulnificus,* which were cultivated in M92 medium (3% tryptic soy broth, Becton Dickinson; 0.3% yeast extract). The human fecal panel covers the gram-positive bacteria *Enterococcus casseliflavus* DSM 7370, *Enterococcus faecalis* DSM 20478, *Enterococcus faecium* DSM 20477, *Enteroccous hirae* DSM 27815 and the gram-negative bacterium *Escherichia coli* DSM 1576. All *Enterococcus* species were cultivated in M92 medium. TSB12 medium (1.2% tryptic soy broth, 0.5% NaCl) was used for *E. coli*, as well for the human pathogen methicillin-resistent *Staphylococcus aureus* DSM 18827 (MRSA). Three phytopathogenic bacteria *Pseudomonas syringae* DSM 50252, *Xanthomonas campestris* DSM 2405, *Erwinia amylovora* DSM 50901 were cultivated in TSB12 medium, whereas *Ralstonia solanacearum* DSM 9544 *R* was grown in M186 medium containing 1% glucose, 0.5% peptone from soybeans, 0.3% yeast extract and 0.3% malt extract (Becton Dickinson). In order to evaluate the antimicrobial activity of the crude extracts, a stock solution (20 mg/ml) in DMSO was prepared and transferred in duplicates into a 96-well microplate. 200 µl of the respective test organism were added to each well after diluting an overnight culture to an optical density (600 nm) of 0.01 – 0.03. Microplates containing bacteria of the phytopathogenic and aquatic panels were incubated for 5 - 6 h at 28 °C and shaken at 200 rpm. The exception applied for *V. splendidus*, which was incubated at 22 °C. *Vibrio ichthyoenteri, A. bacteriolytica* and *R. solanacearum* were cultivated for a longer period (18 h). *Lactococcus garvieae* and *Enterococcus* spp. were incubated for 5 - 7 h at 37 °C without shaking while MRSA and *E. coli* were incubated with shaking at 200 rpm. Subsequently, 10 µl resazurin solution (0.3 mg/ml in phosphate buffer) was added to each well and the microplates were incubated again for 5-60 min before measuring fluorescence (560/590 nm) using a microplate reader (Tecan Infinite M200, Tecan, Männedorf, Switzerland). For *Enterococcus* sp. and *L. garvieae*, the pH indicator bromocresol purple was used as detection reagent to determine color/pH change due to acidification caused by growth of the respective test strains. The color change was detected by absorbance measurement (600 nm/reference 690 nm). The percentage of inhibition was calculated based on a negative control (no extract) and compared to a positive control. Chloramphenicol was used as positive control for most pathogens, except for *Enterococcus* sp. and *L. garvieae*, for which ampicillin was used, and for *R. solanacearum*, for which tetracycline was the standard drug. To determine the IC_50_ values (i.e., the concentration that inhibits the half of the growth of a test organism), a 1:2 dilution series of the extracts was prepared and tested as described above. A concentration depending graph was created with Excel and the IC_50_ value was calculated.

#### 2.5.2. Assays against pathogenic fungi and oomycetes

The phytopathogenic fungus *Magnaphorte grisea* DSM 62938 and the oomycete *Phytophthora infestans* CBS 120920 were cultivated on agar medium for 2 weeks until sporulation. *Phytophthora infestans* was inoculated on carrot medium (4% grated carrot, 1.5% agar) and *M. grisea* on GPY medium (0.1% glucose, 0.05% peptone, 0.01% yeast extract, 1.5% agar). A suspension of 1 - 5*10^4^ spores/ml was prepared for each pathogen in liquid media. For this aim, pea medium (150 g peas cooked, filtrated, filled up to 1 L with H_2_O and 5 g glucose added, pH 6.5) was used for *P. infestans* and M186 medium for *M. grisea.* Crude extracts were prepared in microplates as described above and a volume of 200 µl of the spore suspension was added to each microplate well. Microplates were incubated at 22 °C for 3 days (*M. grisea*) or 4 days (*P. infestans*), and the absorbance was measured at 600 nm. The percentage inhibition and IC_50_ determinations were carried out as described above.

### 2.6. UPLC-ESI-QToF-MS/MS Analyses

Ultra-high-performance liquid chromatography analysis of the selected crude extracts (concentration 1 mg/mL in MeOH) was performed on an Acquity UPLC I-Class System (Waters, Milford, MA, United States) coupled to a Xevo G2-XS QToF Mass Spectrometer (Waters, Milford, MA, United States) operated in fast data-dependent acquisition mode (LC–MS/MS in fast DDA mode) controlled by MassLynx version 4.2. ULC-MS grade solvents were purchased from Biosolve Chimie, Dieuze, France. The volume injected was set to 0.3 µL for bacterial crude extracts and 0.6 µL for fungal crude extracts. All samples were analyzed in triplicate. Chromatographic separation was achieved on a CORTECS UPLC C18 column (1.7 mm, 100 × 2.1 mm, Waters, Milford, MA, United States) at a temperature of 40 °C. The mobile phase was composed of a mixture of (A) water with 0.1% formic acid (v/v) and (B) acetonitrile with 0.1% formic acid and pumped at a rate of 0.4 mL/min. The elution gradient was set as follows: 0.0–8.5 min, gradient from 30% to 99% B; 8.5–14.5 min, isocratic 99% B; back to initial condition in 0.1 min, for 5 min. Mass spectrometry data in the range of *m/z* 50–1200 Da were acquired with electrospray ionization source (ESI) in positive ion detection mode with the following parameters: spray voltage of 3 kV, cone gas flow of 50 L/h, desolvation gas flow of 1000 L/h, source temperature of 150 ^◦^C, desolvation temperature of 500 °C with sampling cone and source offset at 40 and 60, respectively. The MS/MS experiments were carried out in tandem with ramp collision energy (CE): Low CE from 6 to 60 eV and a high CE of 9 to 80 eV. The solvent (MeOH) was run under the same conditions.

#### 2.6.1. UPLC–MS/MS data processing and molecular networking

UPLC–MS/MS raw data were converted to mzXML file format using MSconvert, from the ProteoWizard suite (Chambers et al., 2012) and then processed in batch mode with the software MZmine 3.2.8 (Schmid et al., 2023). The output files (.csv and .mgf) were exported to GNPS (Global Natural Products Social Molecular Networking) and networks were created with the feature-based molecular networking (FBMN) workflow (Wang et al., 2016; Nothias et al., 2020). Briefly, nodes in the FBMN represent consensus MS/MS spectra and are connected by edges to other nodes, which share similar MS/MS fragmentation patterns, hence produce similar MS/MS spectra. Networks were then generated with the following parameters: a precursor ion mass tolerance 0.05 Da, MS/MS fragment ion tolerance 0.05 Da, cosine score > 0.6, minimum number of matched peaks ≥ 6, maximum number of neighbour nodes = 10, maximum number of nodes in a single network = 100. The spectra in the networks were then searched against GNPS spectral libraries. A score above 0.6 and at least 6 matching peaks were required to keep matches between network spectra and library spectra. Generated networks were visualized using Cytoscape version 3.9.1 (Shannon et al., 2003) with the edges modulated by cosine score. The analysis of the networks was simplified showing only nodes representing ions in a *m/z* range of 180–1200 Da. Nodes originating from the solvent (MeOH) were removed from the MN. In the networks, node colors were mapped based on the source of the spectra files or on the culture conditions. In addition to the library search, annotation of compounds was performed by combining automated and manual dereplication tools. Further *in-silico* annotations were done by SIRIUS with CANOPUS and CSI:FingerID tools as well as MolDiscovery (v.1.0.0) and DEREPLICATOR+ tool, available on GNPS (Dührkop et al. 2015; Djoumbou Feunang et al., 2016; Dürkop et al., 2019; Cao et al., 2021; Dührkop et al., 2021; Kim et al., 2021). Additional compound annotations were performed by searching putative molecular formulae, predicted with MassLynx version 4.1, against natural product databases such as COCONUT (https://coconut.naturalproducts.net/ <u>accessed 11 August 2023</u>, Sorokina et al., 2021) LOTUS, (https://lotus.naturalproducts.net/ <u>accessed 11 August 2023,</u> Rutz et al., 2022) MarinLit (http://pubs.rsc.org/marinlit/ <u>accessed 11</u> <u>August 2023</u>), Dictionary of Natural Products (https://dnp.chemnetbase.com <u>accessed 11 August</u> <u>2023</u>), The Natural Products Atlas (https://www.npatlas.org/joomla/index.php/search/basic-search, accessed 11 August 2023 van Santen et al., 2019) and Scifinder (https://scifinder-n.cas.org <u>accessed 11 August 2023</u>).

## 3. Results

### 3.1. Isolation and identification of microorganisms from Z. marina and its surrounding environment

By using 9 different isolation media, over 300 microbial strains were retrieved from the surfaces and the inner tissues of healthy and decaying eelgrass leaves, eelgrass roots, plus the adjacent sediment and surrounding seawater references. The number of strains was reduced to 107 based on the exclusion of strains with identical sequence and biological safety level (i.e., removal of organisms belonging to BSL-2) according to German safety guidelines TRBA 460 and TRBA 466 (Technical rules for biological agents for fungi and for bacteria, respectively). The majority of strains were identified as bacteria (76 strains, 71%), while only 31 strains (29%) were assigned to fungal taxa (Fig. 1, Supplementary Tables S2 and S3). The MA medium afforded the largest portion of bacterial strains (28%), while PDA and the newly designed *Z. marina* media ZMB and ZMF represented the most prolific media for fungal growth yielding altogether 68% of fungal isolates (PDA: 26 %, ZMB: 19%, ZMF: 23%).

**Fig. 1.**
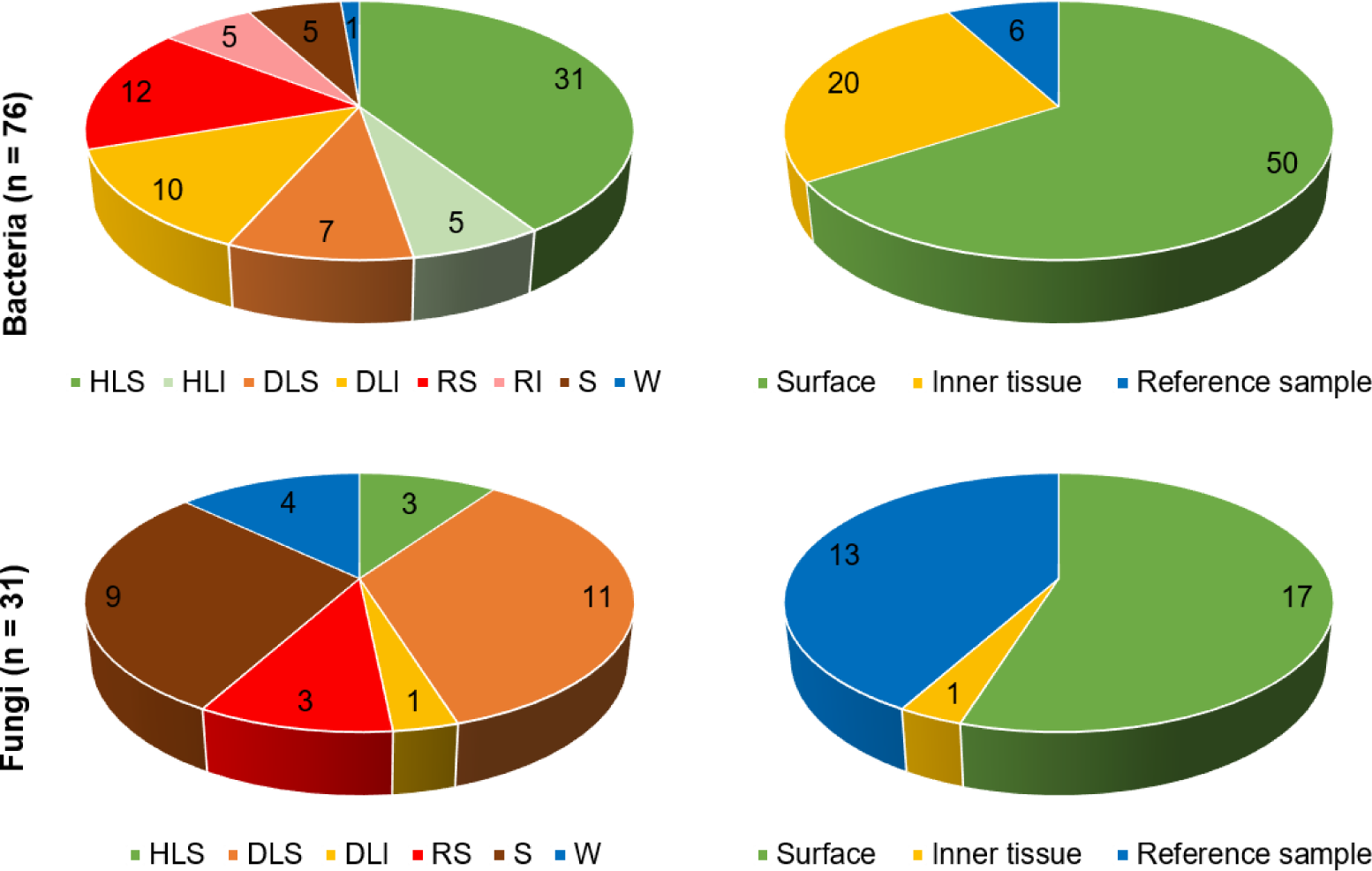
Isolation source of bacterial (top) and fungal (bottom) strains from eelgrass and reference samples. HLS: Healthy leaf surface, HLI: Healthy leaf inner tissue, DLS: Decaying leaf surface, DLI: Decaying leaf inner tissue, RS: Root surface, RI: Root inner tissue, W: Seawater reference, S: Sediment reference.

Bacterial strains derived from the surface of healthy *Z. marina* leaves (HLS, 31 strains) dominated the isolates (41%; Fig. 1). Additional 19 bacterial strains were obtained from decaying leaf surfaces (DLS, 7 strains) and root surfaces (RS, 12 strains), while 20 strains were retrieved from inner tissues. The striking difference in the number of isolates between the HLS (31 strains) and healthy leaf inner tissues (HLI, 5 strains) indicated the richness of the epiphytic bacterial community on eelgrass phyllosphere. Most fungal strains were obtained from decaying leaf surfaces (11, 35%), whereas inner tissues of decaying leaves (DLI) afforded only one single fungal strain. No fungi were obtained from the inner tissues of healthy eelgrass leaves and roots. Altogether 6 bacterial and 13 fungal strains were retrieved from the reference samples, with sediment representing the second-best source of fungi (9 isolates, 29%). Figure 1 shows the overall distribution of the isolates on surfaces, inner tissues and reference samples, indicating the abundant microbial diversity of the *Z. marina* epibiome (63 % of all strains). The eelgrass endosphere was much lesser colonized by culturable bacteria and fungi (20% of all strains). Few bacterial strains were obtained from reference samples (sediment, S: 5 strains, seawater, W: 1 strain), while fungi deriving from the seawater (4 strains) and sediment (9 strains) accounted for 42% of all isolated fungi in this study.

Sanger sequencing followed by comparison to the NCBI nucleotide database using the nucleotide BLAST algorithm (Altschul et al., 1990) allowed the identification of most microbial strains at species (44%) or genus level (54%; Supplementary Tables S2 and S3). Bacterial isolates belonged to four phyla, namely Proteobacteria, Actinomycetota, Bacteroidota and Firmicutes (Fig. 2A). Proteobacteria dominated most seagrass sources (HLS: 58%, DLS: 71%, DLI: 40%, RS: 50%, RI: 80%). Among the Proteobacteria, strains were assigned either to the class of Gammaproteobacteria (66% of all proteobacterial strains) or Alphaproteobacteria (34%) with healthy blades being the richest source for both classes. Notably, Proteobacteria were fully absent in the healthy leaf endosphere (HLI: 0%). All bacterial endophytes of the healthy eelgrass blades were assigned to Firmicutes, which showed either low abundance (HLS, RS, DLI) or were absent (DLS, RI) in other eelgrass samples. The phyla Actinomycetota and Bacteroidota showed comparably low abundances in *Z. marina* samples (Actinomycetota: ≤ 20%, Bacteroidota: ≤ 33%). Bacteroidota were isolated from all surfaces (HLS: 19%, DLS: 14%, RS: 33%), however endophytic Bacteroidota were only detected in decaying leaf inner tissues (20%). The largest number of Actinomycetes were obtained from HLS (5 strains) and except for HLI, all other eelgrass samples afforded 1 (DLS, RS, RI) or 2 (DLI) Actinobacteria isolates. Actinomycetota was the sole or main component of the microbiota of the reference samples (W: 100%, S: 80%), while the only remaining sediment-associated bacterium was assigned to Gammaprotobacteria. Hence, the cultivable eelgrass microbiota (both phyllosphere and endosphere) did not resemble the bacterial composition of the reference samples.

**Fig. 2.**
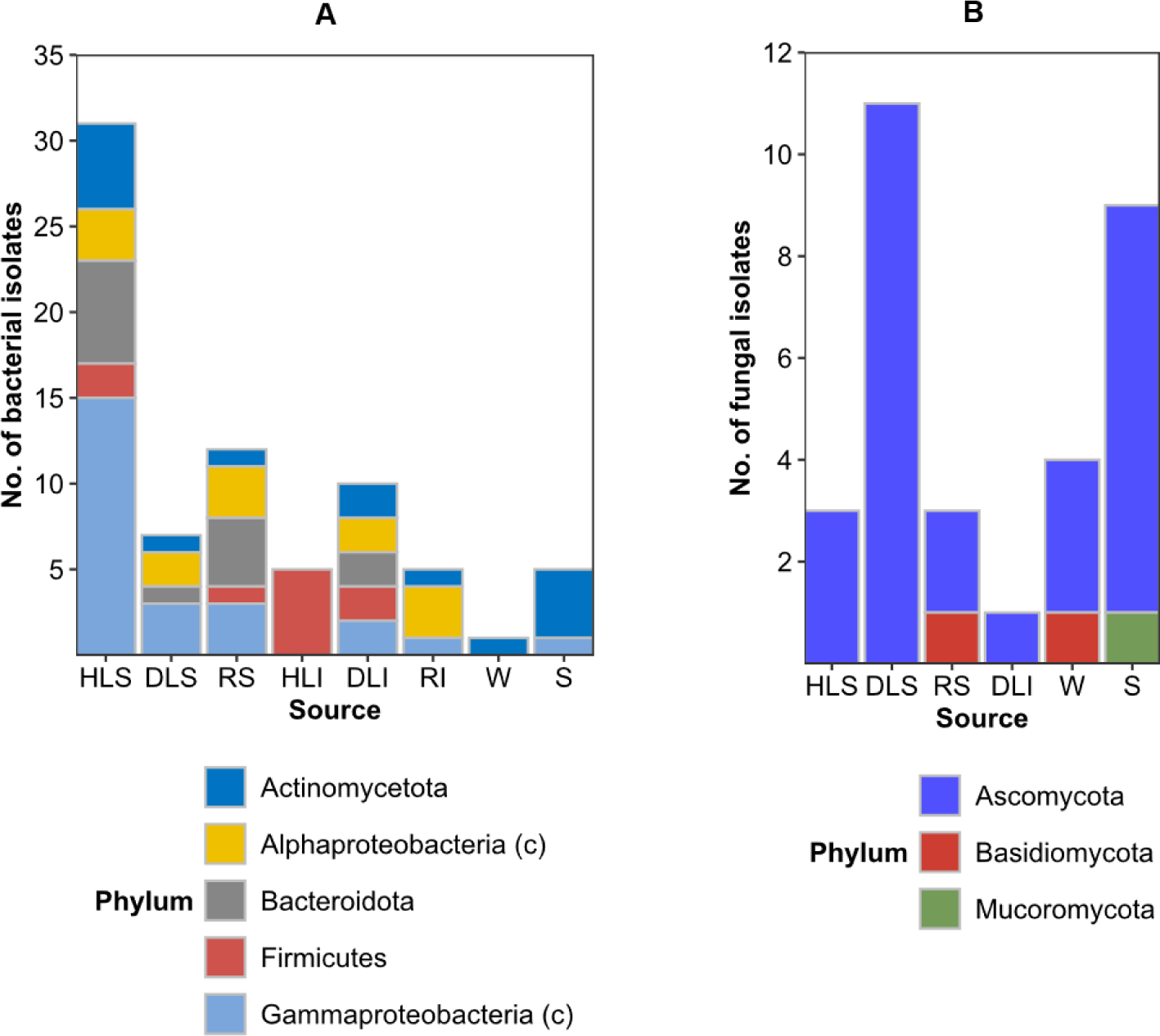
Diversity of bacterial (**A**) and fungal strains (**B**) at phylum level. Proteobacteria are shown at class level (c). HLS: Healthy leaf surface, HLI: Healthy leaf inner tissue, DLS: Decaying leaf surface, DLI: Decaying leaf inner tissue, RS: Root surface, RI: Root inner tissue, W: Seawater reference, S: Sediment.

The culturable fungal community was assigned to three different phyla. Ascomycota clearly dominated the culturable eelgrass mycobiome (67-100%, Fig. 2B), and was the sole component of healthy (HLS 3 strains) and decaying leaf surfaces (DLS 11 strains) and the endosphere of the decaying leaves (DLI 1 strain). The fungal epiphytes of roots were either assigned to Ascomycota (2 strains) or Basidiomycota (1 strain). Seawater-derived fungi showed a very similar distribution at phylum level (Ascomycota: 3 strains, Basidiomycota: 1 strain). The comparably rich mycobiome of the sediment reference was dominated by the phylum Ascomycota (89%). One strain was identified as member of the phylum Mucoromycota, which was the only representative of this phylum in this study. The culturable diversity of sediment-dwelling fungi was also different to those obtained from *Z. marina* samples.

Bacterial strains isolated from *Z. marina* and reference samples were affiliated to 49 unambiguously identifiable genera (Fig. 3A, Supplementary Table S2). Overall, there was only little overlap between different eelgrass tissues, since the majority of bacterial genera were exclusive to one sample source (82%) and no bacterial genus was common to all *Z. marina* samples. With 26 different genera, healthy leaf surfaces (HLS) were by far the most diverse source of bacteria, with 21 genera being exclusive to HLS (e.g., *Exiguobacterium*, *Glaciecola*, *Micrococcus*, *Stenotrophomonas*, *Vibrio*). Only five genera were shared with other surface samples: *Flavobacterium* and *Microbacterium* (roots), *Alteromonas* and *Streptomyces* (decaying leaves) and ubiquitous *Pseudoalteromonas* sp. from all surfaces. Root surfaces were the second richest source of bacteria, yielding 11 different genera, five of which being unique to RS (e.g., *Dokdonia*, *Octadecabacter*). Bacteria associated with DLS generally resembled to other plant tissues, *Sulfitobacter* was the only genus exclusive to decaying leaf phyllosphere. Notably, the root and leaf endosphere (both healthy and decaying) were much less diverse than the eelgrass surfaces. All five strains obtained from HLI were identified as members of the genus *Bacillus*, which was overall the most abundant bacterial genus in this study. Root endophytic bacteria were assigned to 5 genera exclusive to RI, e.g., *Hoeflea alexandrii* and *Rhodococcus* sp. Notably, DLI and DLS shared several bacterial genera common to the marine habitat (*Aquimarina*, *Pseudoalteromonas*, *Streptomyces*). In addition, the decaying leaf endosphere hosted seven genera (e.g., *Mycolicibacterium*, *Thalassospira*) that are absent from all other eelgrass tissues.

**Fig. 3.**
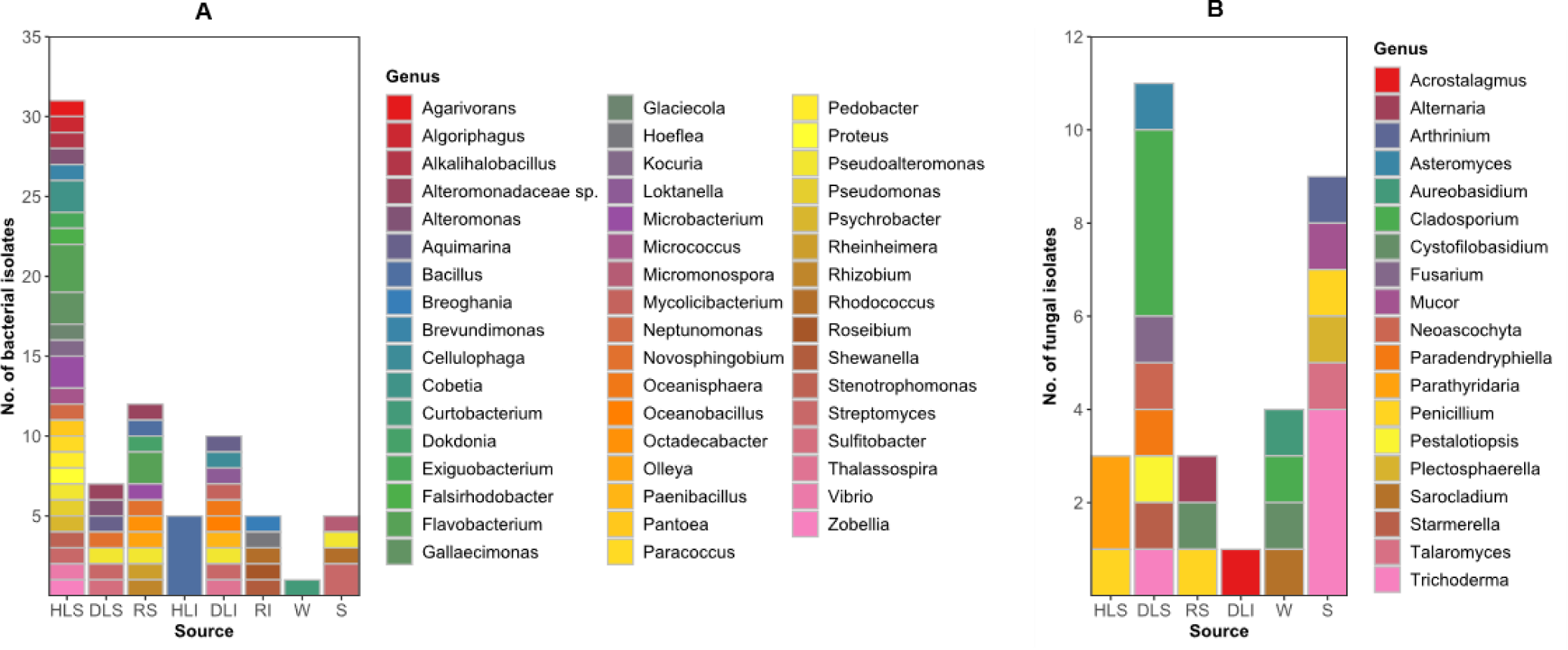
Diversity of all bacterial (**A**) and fungal (**B**) strains at genus level. HLS: Healthy leaf surface, HLI: Healthy leaf inner tissue, DLS: Decaying leaf surface, DLI: Decaying leaf inner tissue, RS: Root surface, RI: Root inner tissue, W: Seawater reference, S: Sediment reference.

Reference samples were among those with the lowest bacterial diversity: *Curtobacterium herbarum* strain 218 was the only bacterium obtained from ambient seawater (Fig. 3A, Supplementary Table S2). Sediment-associated bacteria were assigned to four different genera, of which all but one (*Micromonospora*) were detected also in eelgrass samples: *Pseudoalteromonas* (HLS, DLS, DLI, RS), *Rhodococcus* (RI) and *Streptomyces* (HLS, DLS, DLI). Nevertheless, reference samples showed much lower bacterial diversity and overall did not resemble the majority of the bacterial community of *Z. marina*.

The cultivable eelgrass mycobiome was considerably less diverse, yielding only 19 different genera (Fig. 3B, Supplementary Table S3). Decaying leaf surfaces were the richest source of fungal strains (8 genera) hosting six genera exclusively (*Asteromyces*, *Fusarium*, *Neoascochyta*, *Paradendryphiella*, *Pestalotiopsis*, *Starmerella*). The remaining decaying leaf epiphytes were assigned to the fungal genera *Cladosporium* and *Trichoderma.* Surfaces of healthy leaves and roots showed comparably lower fungal abundance (n = 3 each). The phylloplane of healthy blades hosted the fungal genera *Parathyridaria* and *Penicillium*. The root phylloplane mycobiome was composed of *Penicillium glabrum* (strain 403), *Cystofilobasidium bisporidii* (strain 417) and *Alternaria* sp. (strain 903a). Notably, culturable fungi were absent from the endosphere of healthy leaves and roots, and the only fungal endophyte isolated from decaying leaves belonged to the class Sordariomycetes (*Acrostalagmus luteoalbus* strain 720), which was absent from all other tissues. Accordingly, the surface and inner tissues of *Z. marina* accommodated distinct culturable fungal communities, with DLS yielding the most diverse fungal consortium.

Sediment-associated fungi (Fig 3B, Supplementary Table S3) were, with six different genera, the second most diverse sample source and were dominated by *Trichoderma* sp. (4 strains). This genus and *Penicillium* (1 strain) were also detected in the eelgrass phylloplane, while all other genera (*Arthrinium*, *Mucor*, *Plectosphaerella*, *Talaromyces*) were exclusive to the sediment reference. Seawater samples yielded only four fungal isolates, two of which were assigned to genera unique for the seawater reference (*Aureobasidium*, *Sarocladium*). Hence, despite some overlap, the vast majority of seagrass-associated taxa were not found in seawater or sediment reference samples indicating a seagrass-specific mycobiome.

### 3.2. Assessment of antimicrobial activity of the isolates

Each isolate was cultivated in two solid media: bacteria in MA and GYM, and fungi in M34 and PDA. In total, 214 crude microbial EtOAc extracts (152 bacterial, 62 fungal) were assessed for their *in vitro* antimicrobial activity against 28 bacterial, fungal or oomycete type pathogens from 4 panels of test organisms (Supplementary Tables S4-S11). The first test panel (aquatic) included common pathogens of the aquatic/marine environments affecting marine animals (fish, shellfish/bivalves, shrimps, corals, sea urchins) and seaweeds, but also causing illness in humans through ingestion (seafood consumption, e.g., *V. parahaemolyticus*, drinking of contaminated water, e.g., *V. cholerae*) or skin contact (e.g., *V. vulnificus*) (Baker-Austin et al., 2018). The second (fecal) pathogen panel included commensal intestinal bacteria that are mainly discarded to seawater by humans (*Enterococcus* spp., *E. coli*; Noble et al., 2003). Contact with seawater has also been associated with an enhanced risk of *Staphylococcus* infections, including MRSA, which is shed by human skin, feces or by wastewater contamination, and transmitted back to human by sea bathing (Levin-Edens et al., 2011; Plano et al., 2011). MRSA is often detected in surface- and wastewaters (Denissen et al., 2022), hence was included as the third ‘human’ pathogen panel. The pathogens of fecal and human panels cause gastrointestinal and several other critical infections in human (Ascioti et al., 2022). The last panel entailed economically important terrestrial plant pathogens, some belonging to genera reportedly affecting seagrasses, e.g., *Phytophora* (Govers et al., 2016; Sullivan et al., 2018), which are washed into the sea by large watersheds and riverine input. Due to the large number of samples and test pathogens, we visualized antimicrobial activity of bacterial and fungal isolates that had an IC_50_ value (up to 100 μg/ml) in heatmaps (Fig. 4). Supplementary Tables S4-S11 show the bioactivity data of all 214 extracts against all pathogens. For simplicity, we use, whenever appropriate, abbreviations showing the origin (e.g. HLS) and the culture medium (e.g., GYM) together to describe the extract of an isolate, e.g., HLS-GYM bacterium.

**Fig. 4.**
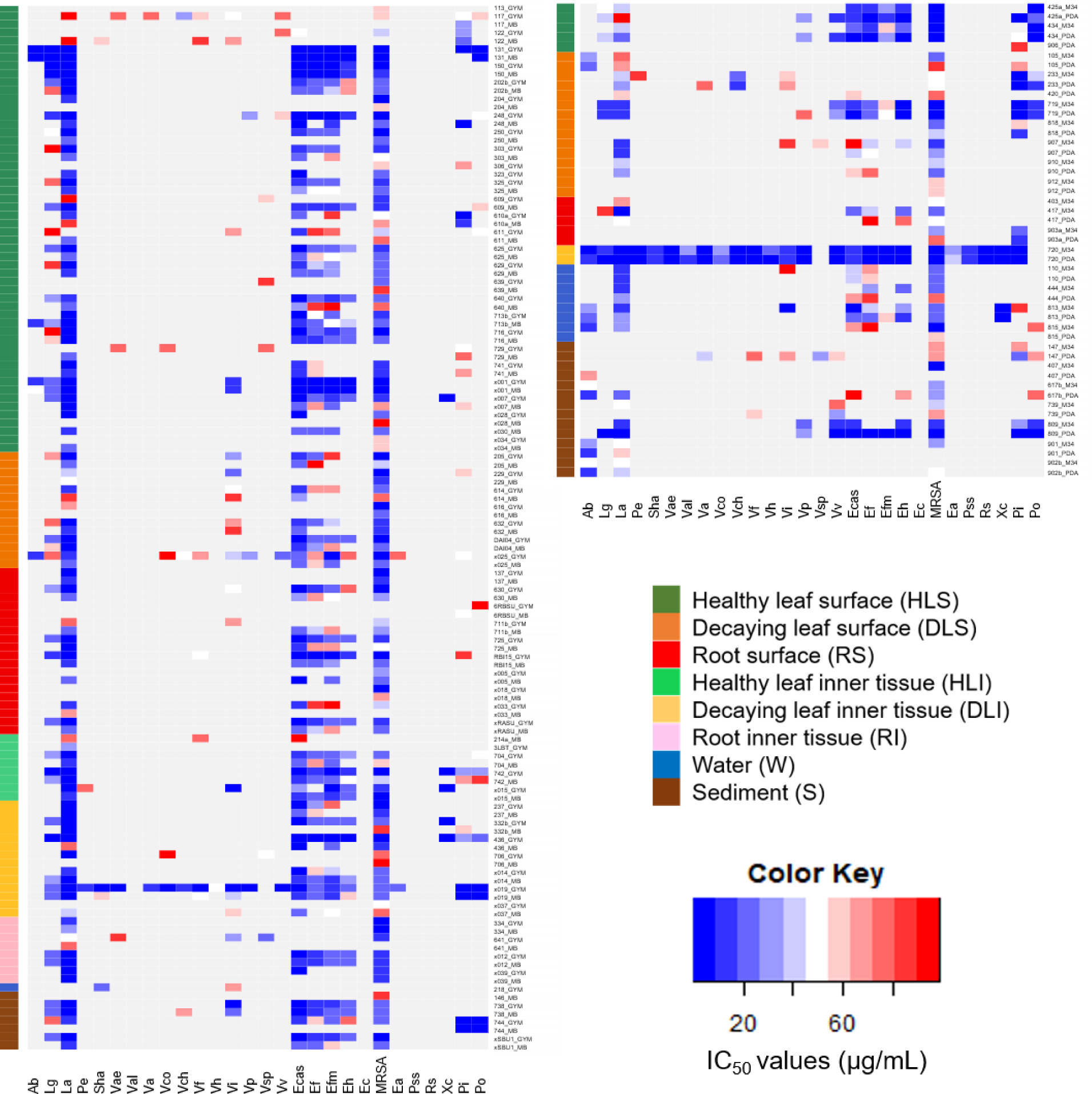
Antimicrobial activity of bacterial (left) and fungal (right) extracts with IC_50_ values ≤ 100 µg/ml against test pathogens. Abbreviations for test pathogens are shown in the Supplementary Table S1.

#### 3.2.1. Bioactivity of epiphytic bacterial extracts

The GYM extract of HLS-associated (HLS-GYM) *Streptomyces* sp. 131 was specifically active against three test organisms in the aquatic panel, *L. aquatica* (La), the seaweed pathogen *Algicola bacteriolytica* (Ab, both IC_50_s 1.8 μg/ml) and the fish pathogen *Lactococcus garvieae* (Lg, IC_50_ 6.9 μg/ml, Fig. 4, Supplementary Table S4). Gammaproteobacteria dominated the observed activities of the HLS-GYM bacterial extracts with varying spectrum and potency. For example, *Psychrobacter nivimaris* 741 and *Pseudomonas* sp. x028 inhibited only La (IC_50_s ∼ 7.0 μg/ml); *Stenotrophomonas* sp. 150 was equally active against La and Lg (IC_50_s 6.6 and 8.6 μg/ml), *Cobetia* sp. x001 also inhibited Ab and *V. ichthyoenteri* (Vi) with moderate IC_50_ values (16.7 and 14.0 μg/ml, resp.). *Vibrio* sp. 248 had good activity against La, Lg, *V. parahaemolyticus* (Vp) and *V. vulnificus* (Vv) with IC_50_s ranging from 6.5 to 56.1 μg/ml. Several Flavobacteria (phylum Bacteroidota), e.g., *Flavobacterium* sp. 625 inhibited La and Lg (IC_50_s 6.2 and 25.8 μg/ml, resp.).

Marine broth agar (MA) extract of HLS bacterium *Streptomyces* sp. 131 inhibited solely the same aquatic pathogens with higher potency (Fig. 4, Supplementary Table S6). Its IC_50_ values against La and Lg (0.6 and 1.1 μg/ml, respectively) were comparable to the positive controls, while its potency towards the gram-negative seaweed pathogen *Algicola bacteriolytica* (Ab) was approximately four times greater than the standard chloramphenicol (IC_50_s 0.8 and 2.9 μg/ml, resp.). Seven HLS originated Gammaproteobacteria exerted low IC_50_ values (<10 μg/ml) against La, plus generally lower activity against Lg. Two of them, *Proteus* sp. 713b and *Cobetia* sp. x001, inhibited La, Lg and Ab. The best anti-*Vibrio* effect was exhibited by the MA extracts of *Cobetia* sp. x001 against the fish pathogen Vi (IC_50_ 17.6 μg/ml). *Alkalihalobacillus hwajinpoensis* strain x007 (Bacillaceae) was exclusively active against La (IC_50_ 7.0 μg/ml).

The majority of HLS-GYM extracts had remarkable activity against multiple fecal pathogens, particularly against *Enterococcus casseliflavus* (Ecas), *E. faecalis* (Ef) and *E. faecium* (Efm, Fig. 4, Supplementary Table S5). Indeed only 11 HLS-GYM extracts lacked activity against enterococci. HLS-GYM extract of *Streptomyces* sp. 131 inhibited the growth of all fecal *Enterococcus* spp. (IC_50_s 1.6-6.4 μg/ml). Half of the bacterial HLS-MA extracts showed inhibitory activity against at least two fecal pathogens (Fig. 4, Supplementary Table S7). HLS-MA *Streptomyces* sp. 131 extract potently inhibited the growth of all fecal enterococci with equal (against *E. hirae*, Eh) or 8 and 2.5-times higher efficacy (against Ecas and Ef, resp.) than the positive control ampicillin. Both GYM and MA extracts of many Gammaproteobacteria, *Stenotrophomonas* sp. 150, *Vibrio* sp. 248 and *Cobetia* sp. x001, had anti-*Enterococcus* potential (Fig. 4, Supplementary Tables S5 and S7). The MA extract of the latter bacterium had equal potency with ampicillin (IC_50_ 2.4 μg/ml) against Ecas (Fig. 4, Supplementary Table S7). *Escherichia coli* (Ec) was not inhibited by any bacterial or fungal extract.

The human pathogen MRSA was susceptible to HLS-derived GYM extracts, many of which showed cross activity against enterococci (particularly Ecas) (Fig. 4, Supplementary Table S5). Except for three isolates, all HLS GYM extracts were active against MRSA. The anti-MRSA activity of the Gammaproteobacterium *Pantoea* sp. 716 was higher than of the reference compound chloramphenicol (IC_50_s 0.9 and 1.5 μg/ml, resp.), followed by *Streptomyces* sp. 131 and *Vibrio* sp. 248. The MA extracts of 24 HLS bacteria were also active against MRSA (Fig. 4, Supplementary Table S7). Again *Streptomyces* sp. 131 showed extraordinary anti-MRSA activity with almost 3-fold greater potency than chloramphenicol (IC_50_s 0.6 and 1.5 μg/ml, resp.). *Cobetia* sp. x001 was the second most active strain, while the remaining bacterial extracts had moderate to low inhibitory potentials.

Activity rates were generally lower against phytopathogens. Phytopathogenic oomycete *Phytophthora infestans* (Pi) and fungus *Magnaphorte grisea* (Mg) were those with highest susceptibility towards a smaller portion of GYM extracts with varying potencies (Fig. 4, Supplementary Table S5). Only 9 HLS-derived bacteria exhibited antiphytopathogenic activity. *Streptomyces* sp. 131 was the most toxic strain versus Mg (IC_50_ 1.2 μg/ml). *Phytophthora infestans* (Pi) was inhibited by the same bacterium and a *Flavobacterium* sp. 610a (IC_50_s 6.7 and 5.7 μg/ml). The HLS bacterium *Alkalihalobacillus hwajinpoensis* (strain x007) was the only isolate that inhibited the phytopathogenic bacterium *Xanthomonas campestris* (IC_50_ 6.2 μg/ml). MA extract of *Streptomyces* sp. 131 was selectively active against Mg with an IC_50_ value close to the standard antifungal drug nystatin (IC_50_s 0.5 and 0.3 μg/ml, resp.), while *Vibrio* sp. 248 strongly inhibited Pi (IC_50_ 0.9 μg/ml, Fig. 4, Supplementary Table S7).

All DLS-GYM extracts inhibited the growth of at least one aquatic pathogen (Fig. 4, Supplementary Table S4). The most sensitive aquatic pathogen was La, which was efficiently inhibited by multiple Gammaproteobacteria, e.g., Alteromonadaceae sp. 205, *Alteromonas stellipolaris* 632, *Pseudoalteromonas* sp. DAI04 (IC_50_s 6.2 to 9.1μg/ml). Five bacteria inhibited Lg and Vi with moderate potency. The GYM extract of *Streptomyces griseorubens* x025 had the broadest activity against six *Vibrio* spp. (IC_50_s 18.8-97.3 μg/ml), besides Ab and La with IC_50_s 9.4 and 11.0 μg/ml, respectively. MA extracts of the most DLS epibionts were found to exert moderate to low activity against aquatic pathogens, La, Lg and *V. harveyi* (Vh), with *Pseudoalteromonas* sp. DAI04 being the most active strain (La, IC_50_ 6.2 μg/ml, Fig. 4, Supplementary Table S6)

The majority of the DLS-GYM extracts inhibited three to four fecal pathogens with *Streptomyces griseorubens* x025 and *Pseudoalteromonas* sp. DAI04 being the most active ones (IC_50_s 7.0 vs Efm and 7.1 μg/ml vs Ecas, Fig. 4, Supplementary Table S5). All DLS GYM extracts had anti-MRSA activity, and the lowest IC_50_ values (6.6 and 6.8 μg/ml, resp.) were obtained with *Streptomyces griseorubens* x025 and *Sulfitobacter pontiacus* 616 extracts. Only two DLS-GYM bacteria exhibited low antiphytopathogenic activity towards *Erwinia amylovora* (Ea) and *Phytophthora infestans* (Pi). Many MA-DLS extracts had modest activity against fecal pathogens (5 extracts) and MRSA (all 7 extracts) while none was anti-phytopathogenic even at the highest test concentration (100 μg/ml, Fig. 4, Supplementary Table S7).

Half of the GYM extracts of root surface (RS) bacteria exhibited activity versus the aquatic panel (Fig. 4, Supplementary Table S4). Besides few Gammaproteobacteria (e.g., *Pseudoalteromonas* sp. RBI15), the Alphaproteobacterium *Rhizobium* sp. 137 and *Bacillus* sp. 725 exhibited good activity against La, while several strains moderately inhibited the growth of Lg, *Aliivibrio fischeri* (Vf) and *V. ichthyoenteri* (Vi). A similar trend was observed against fecal pathogens; of the five bioactive RS-GYM extracts, four belonged gram-negative bacteria (IC_50_s 4.4-14.5 μg/ml), *Pseudoalteromonas* sp. RBI15 being the most active against Ecas (Fig. 4, Supplementary Table S5). *Bacillus* sp. 725 was the only gram-positive bacterium inhibiting all four enterococci. The majority of GYM-RS bacteria inhibited MRSA, and four Proteobacteria, e.g., *Pseudoalteromonas* sp. RBI15, *Rhizobium* sp. 137, Alteromonadaceae sp. 630 and *Octadecabacter* sp. x018, were the most active ones (IC_50_s 6.7-8.8 μg/ml). Negligible activity was exerted by two RS-GYM extracts against phytopathogen panel.

Root-surface bacteria (RS) cultivated in MA medium had moderate and very narrow-spectrum activity towards the aquatic pathogen La, with *Bacillus* sp. 725 displaying the best activity with an IC_50_ value of 8.1 μg/ml (Fig. 4, Supplementary Table S6). Six RS-MA extracts moderately inhibited multiple enterococci and the most interesting was the Flavobacterium *Dokdonia* sp. x005 (IC_50_s 4.7 μg/ml against Ecas; Fig. 4, Supplementary Table S7). Eight RS-MA extracts showed low anti-MRSA potency and the Actinobacterium *Microbacterium* sp. 6RBSU was the only bacterium with low activity against the phytopathogen *Phytophthora infestans* (Pi).

#### 3.2.2. Bioactivity of endophytic bacterial extracts

All five healthy leaf endophytic (HLI) bacteria belonged to *Bacillus* spp. (phylum Firmicutes). Four HLI-GYM extracts showed low μg/ml level inhibition against aquatic pathogen La, while two inhibited Lg. *Bacillus* sp. strain x015 had good activity against the fish pathogen Vi (IC_50_ 2.1 μg/ml; Fig. 4, Supplementary Table S4). Similar bioactivity profile was obtained when HLI bacteria were grown in MA medium (Fig. 4, Supplementary Table S6). Four HLI-MA bacteria inhibited the aquatic pathogen La with *Bacillus* sp. 742 being the most potent (IC_50_ 5.9 μg/ml).

Three GYM-HLI isolates inhibited three to four *Enterococcus* spp. and MRSA, with *Bacillus* sp. 742 being most potent against Ecas and MRSA (IC_50_s 2.7 and 7.5 μg/ml, resp.; Fig. 4, Supplementary Table S5). The same strain and *Bacillus* sp. x015 had notable anti-phytopathogenic effect towards *Xanthomonas campestris* (IC_50_ values 6.2 and 2.1 μg/ml, resp.). Except for *Bacillus pumilis* strains 3LBT, all HLI-MA extracts were moderately active against fecal pathogens (Fig. 4, Supplementary Table S7). Three HLI-MA bacteria exerted anti-MRSA activity, and the potency of *Bacillus* sp. x015 was comparable to that of chloramphenicol (IC_50_s 2.2 and 1.5 μg/ml, resp.). No significant anti-phytopathogenic activity was observed.

Approximately the half of the DLI-GYM bacteria inhibited aquatic pathogens La and Lg, some showing low activity against *V. coralliilyticus* (Vc) and *V. splendidus* (Vs; Fig. 4, Supplementary Table S4)*. Streptomyces* sp. x019 stood out for its broadest spectrum activity against many aquatic pathogens, plus strong activity towards six *Vibrio* spp. including *Vibrio fischeri* (Vf), *V. ichthyoenteri* (Vi), *V. parahaemolyticus* (Vp) and *V. vulnificus* (Vv) with IC_50_ values ranging from 1.5 to 2.2 μg/ml. It was the only bacterial DLI-GYM extract that inhibited *Shewanella algae* (Sha, IC_50_ 6.6 μg/ml). A similar bioactivity profile was observed with the DLI-MA extracts, many inhibiting La. *Streptomyces* sp. x019 exerted the highest potential against La (IC_50_ 3.3 μg/ml) and to some extent against Lg, and Gram-negative pathogens Sha, Vf and Vi (Fig. 4, Supplementary Table S6). Gram-negative bacteria (*Thalassospira* sp. and *Aquimarina* sp.) and the Gram-positive *Oceanobacillus* sp. 332b were also active towards La (IC_50_s 7.6-9.0 μg/ml).

Half of the both DLI-GYM and DLI-MA extracts were active against multiple enteric pathogens, particularly against Ecas and MRSA, with Gram-positive bacterium *Paenibacillus terrae* 436 being the most active extract (IC_50_s 3.8 (GYM) and 5.3 mg/ml (MA; Fig. 4, Supplementary Tables S5 and S7). This strain and *Thalassospira lucentensis* strain 237 had the best anti-MRSA activity regardless of the media. Several gram-positive isolates showed prominent activity against phytopathogens. Both media extracts of DLI bacterium *Streptomyces* sp. x019 showed comparable or better activity against Mg (IC_50_s 0.6 (GYM), 0.2 μg/ml (MA) than the positive control nystatin (IC_50_ 0.3 μg/ml).

Overall, a similar activity profile was seen with both GYM and MA extracts of five root-endophytic (RI) bacteria (Fig. 4, Supplementary Tables S4-S7). *Rhodococcus* sp. x029 was devoid of any activity regardless of the medium used. All remaining extracts RI displayed activity versus aquatic panel particularly against La, with Alphaprotobacterium *Hoeflea alexandrii* x039 and *Breoghania corrubedonensis* 334, as well as Gammaprotobacterium *Shewanella* sp. x012 being the most active. Both MA and GYM extracts of *Shewanella* sp. x012 had equally moderate activity against Lg. The DLI-GYM extracts had somewhat broader activity profile, for example Alphaprotobacterium *Roseibium marinum* 641 moderately inhibited *V. splendidus* (Vs), *V. vulnificus* (Vv) and *V. aestuarianus* (Vae).

As for fecal pathogens, *Shewanella* sp. x012 inhibited all four *Enterococcus* sp. regardless of the medium used, while the GYM-derived *Hoeflea alexandrii* x039 extract specifically targeted Ecas (IC_50_ 4.4 μg/ml; Fig. 4, Supplementary Tables S5 and S7). The majority of RI bacteria were active against MRSA. None of the phytopathogens was targeted by any medium extract of the RI associated bacteria.

Two of the sediment-derived bacteria (S) *Micromonospora* sp. 146 and *Rhodococcus* sp. 733 were generally inactive against all test pathogens, while the remaining three extracts had higher and broader activity when cultivated in GYM medium (Fig. 4, Supplementary Tables S4-S7). *Streptomyces* sp. 738 exerted its best activity against La, Vi and Ecas (IC_50_ s 4.6-7.4 μg/ml), while *Streptomyces scopiformis* 744 was most active against phytopathogens Pi and Po (IC_50_s from 1.3 to 2.8 μg/ml) in both media. The GYM extract of *Pseudoalteromonas ulvae* xSBU1 had generally good activity against Gram-positive pathogens, such as La, Ecas, MRSA but was devoid of any anti-phytopathogenic potential.

#### 3.2.3. Bioactivity of epiphytic fungal extracts

Of the three fungi derived from healthy eelgrass phylloplane (HLS), both media extracts of *Parathyridaria* spp. showed a mild inhibition towards two aquatic pathogens, La or Lg (Fig. 4, Supplementary Tables S8 and S10). Their PDA extracts showed an additional and moderate anti-*Vibrio* activity against Vp and Vv (IC_50_s 16.0 to 32.0 μg/ml). Both media extracts of *Parathyridaria* sp. were active against all fecal panel but the efficacy was 2-fold higher in the PDA extracts (Fig. 4, Supplementary Tables S9 and S11). The same trend applied towards MRSA, which was efficiently inhibited by both strains (IC_50_s 2.1 to 6.5 μg/ml). M34 media extracts of *Parathyridaria* spp. had a specific, notable activity against phytopathogenic fungus Mg (IC_50_s 2.9 and 4.5 μg/ml). Their PDA medium extracts displayed selectivity, *Parathyridaria* sp. 425a being active against Pi and *Parathyridaria* sp. 434 against Mg.

With 10 strains, DLS-associated fungi represented the largest cohort with moderate to low activity (Fig. 4, Supplementary Tables S8-S11). Within the aquatic panel, La was the most sensitive microorganism, but also Lg, Ab and several *Vibrio* spp., *V. cholerae* (Vch), Va, Vi, Vsp and Vv were inhibited modestly. *Cladosporium halotolerans* 233 was noted for its almost equal activity against Vch in both media. *Cladosporium* sp. 907 and *Fusarium* sp. 719 had the lowest IC_50_ values, regardless of the medium, against aquatic pathogens. The activities against fecal pathogens were more potent and broader when PDA medium was used. *Fusarium* sp. 719 showed the best potential towards Ecas and Efm (IC_50_s 3.7 and 5.4 μg/ml, resp.). Most DLS-derived fungi exhibited anti-MRSA activity in both media, with *Fusarium* sp. 719 exhibiting the best activity with IC_50_s 5.8 μg/ml (M34) and 3.9 μg/ml (PDA). As for the phytopathogens, Pi and Mg were effectively inhibited by *Cladosporium halotolerans* 233 and *Fusarium* sp. 719 with PDA extracts being 2-3 times more efficacious (IC_50_s 0.7 and 2.6 μg/ml, resp.)

Rhizoplane (RS) derived fungi were represented by 3 strains, but only the M34 extracts of *Cystofilobasidium bisporidii* 417 displayed some activity against aquatic pathogen La (IC_50_ on M34 medium 8.6 μg/ml), activities against other panel pathogens were low (Fig. 4, Supplementary Tables S8-S11). Only *Alternaria* sp. showed mild activity against the phytopathogen Pi (IC_50_s ∼ 20 μg/ml in both media).

#### 3.2.4. Bioactivity of endophytic fungal extracts

The broadest and the highest activity was exerted by the only representative of decaying leaf-endosphere (DLI) fungus, *Acrostalagmus luteoalbus* 720 (Fig. 4, Supplementary Tables S8-S11). Regardless of the culture medium, it inhibited all aquatic pathogens, except for *V. splendidus*. The IC_50_ values obtained for both media extracts were comparable, and the lowest IC_50_ values by M34 and PDA extracts, respectively, were observed against La (2.1 and 2.2. μg/ml), Pe (both 3.8 μg/ml), Vch (5.0 and 2.6 μg/ml), Vf (4.8 and 3.5 μg/ml), Vp (4.6 and 4.2 μg/ml) and Vv (3.5 and 3.7 μg/ml; Fig. 4, Supplementary Tables S8 and S10). *Acrostalagmus* sp. inhibited all 4 fecal pathogens, with a slightly higher potency observed with its M34 extracts (Fig. 4, Supplementary Tables S9 and S11. The activity against Ef (IC_50_s 0.8 and 1.0 μg/ml) was comparable to that of the reference drug ampicillin (IC_50_ 0.5 μg/ml). With IC_50_ values of 0.2 μg/ml (M34) and 0.5 μg/ml (PDA) against MRSA, it was 7- and 3-times more potent than chloramphenicol (IC_50_ 1.5 μg/ml). *Acrostalagmus* sp. inhibited almost all test phytopathogens particularly the fungus Pi with IC_50_s values of 0.5 (M34 medium) and 0.7 μg/ml (PDA medium), with milder efficacy against bacterial plant pathogens Xc and Rs.

Of the four seawater-associated (W) fungi, *Aureobasidium pullulans* 813 gave generally the best activity profile (Fig. 4, Supplementary Tables S8-S11). When cultured in M34 medium, *A. pullulans* killed reasonably the aquatic pathogens Vi and La (IC_50_s 5.2 and 12.4 μg/ml, resp.), with a modest efficacy against Ab (IC_50_ 30.8 μg/ml). Most of the W-derived fungi inhibited multiple *Enterococcus* sp., and towards Ecas, the M34 extract of *A. pullulans* was as active as ampicillin (IC_50_s 2.8 and 2.4 μg/ml, resp.). Independent of the growth medium, all four fungi were moderately active against MRSA with M34 extract of *Sarocladium* sp. 815 exhibiting the lowest IC_50_ value (8.3 μg/ml). *Aureobasidium pullulans* had moderate and equipotent against phytopathogenic bacterium Xc in both media (IC_50_ 8.3 and 7.1 μg/ml).

Only few reference sediment-derived (S) fungi showed activity towards test pathogens (Fig. 4, Supplementary Tables S8-S11). *Penicillium olsonii* 809 was noted for its broad activity against many pathogens, such as aquatic pathogens La, Lg and Vv (and lower potency against Vp) especially when cultured in PDA medium (IC_50_s 2.4 to 8.5 μg/ml). Its PDA extract was the only sample that inhibited all 4 fecal pathogens with comparable activity with some of the reference compounds). Against MRSA, the same extract had superior activity to the control drug chloramphenicol (IC_50_s 1.0 and 1.5 μg/ml, resp.), while other sediment-associated fungi e.g., *Trichoderma viride* 407 also revealed some anti-MRSA activity (M34: IC_50_ 7.0 µg/mL). Against the phytopathogen Mg, the PDA extract of *P. olsonii* 809 exerted an IC_50_ value lower than the standard nystatin (IC_50_s 0.2 and 0.3 μg/ml, resp.).

### 3.3. Untargeted metabolomics analyses

Next, we wanted to shed light into chemical composition of the bioactive *Zostera*-associated microorganisms and identify secondary metabolites that may underlie their antimicrobial activities. We set the bioactivity threshold (IC_50_ value) to 10 μg/ml-against any pathogen and originating from either culture medium-, and mined the metabolome of 88 extracts (71 bacterial and 17 fungal, Supplementary Tables S12-S13) by an untargeted metabolomics strategy using UPLC-ESI-QToF-MS/MS-based Feature-Based Molecular Networking (FBMN, Nothias et al., 2020) workflow. Compound annotation included the library search on GNPS web platform (Wang et al., 2016), *in-silico* analyses (i.e., SIRIUS, MolDiscovery, DEREPLICATOR+) and manual dereplication tools (Dührkop et al. 2015; Djoumbou Feunang et al., 2016; Dürkop et al., 2019; Cao et al., 2021; Dührkop et al., 2021; Kim et al., 2021).

For metabolome analyses, microbial extracts were divided into groups of epiphytes and endophytes, and further into bacterial and fungal cultures. Seawater and sediment references were analysed together with epiphytes due to their close contact with them. The metabolome of both media extracts obtained are presented together, however we mapped the nodes on the same FBMN to reflect chemical variations and relative abundance of metabolites expressed in different media (Supplementary Figs. S1-S3). Annotated molecular families were highlighted in frames. Supplementary Tables S14-S17 and Supplementary Fig. S5 show the relevant MS information, retention times, reported bioactivity, biological source and the chemical structures of the putatively annotated compounds. Clusters annotated as primary metabolites (e.g., fatty acids, phospholipids, amino acids) are colored grey to put focus on molecular families (MFs) of specialized metabolites that are most likely to contribute to the observed antimicrobial activities.

#### 3.3.1. Metabolome of epiphytic bacteria

Only one Gram-positive HLS associated bacterium (*Streptomyces* sp. strain 131) returned hits in the metabolome analyses and the majority of the annotated MFs were exclusive to this strain. The first small MF belonged to streptophenazines where we could annotate streptophenazines B, C, E, G, H, I (Fig. 5, cluster A) and two oxo-streptophenazines (A and G, Fig. 5, cluster B). Streptophenazines are phenazine alkaloids first detected in a Baltic Sea sponge-associated *Streptomyces* sp. and possess moderate to weak activity versus Gram-positive bacteria (Mitova et al., 2008; Kunz et al., 2014; Liang, et al., 2017). Oxo-streptophenazines, also sourced from a marine-derived *Streptomyces* sp. (Bauman et al., 2019), have marginal or no bioactivity. Notably, both streptophenazine clusters were most abundant in the MA medium extract of *Streptomyces* sp. strain 131 (Supplementary Fig. S1).

**Fig. 5.**
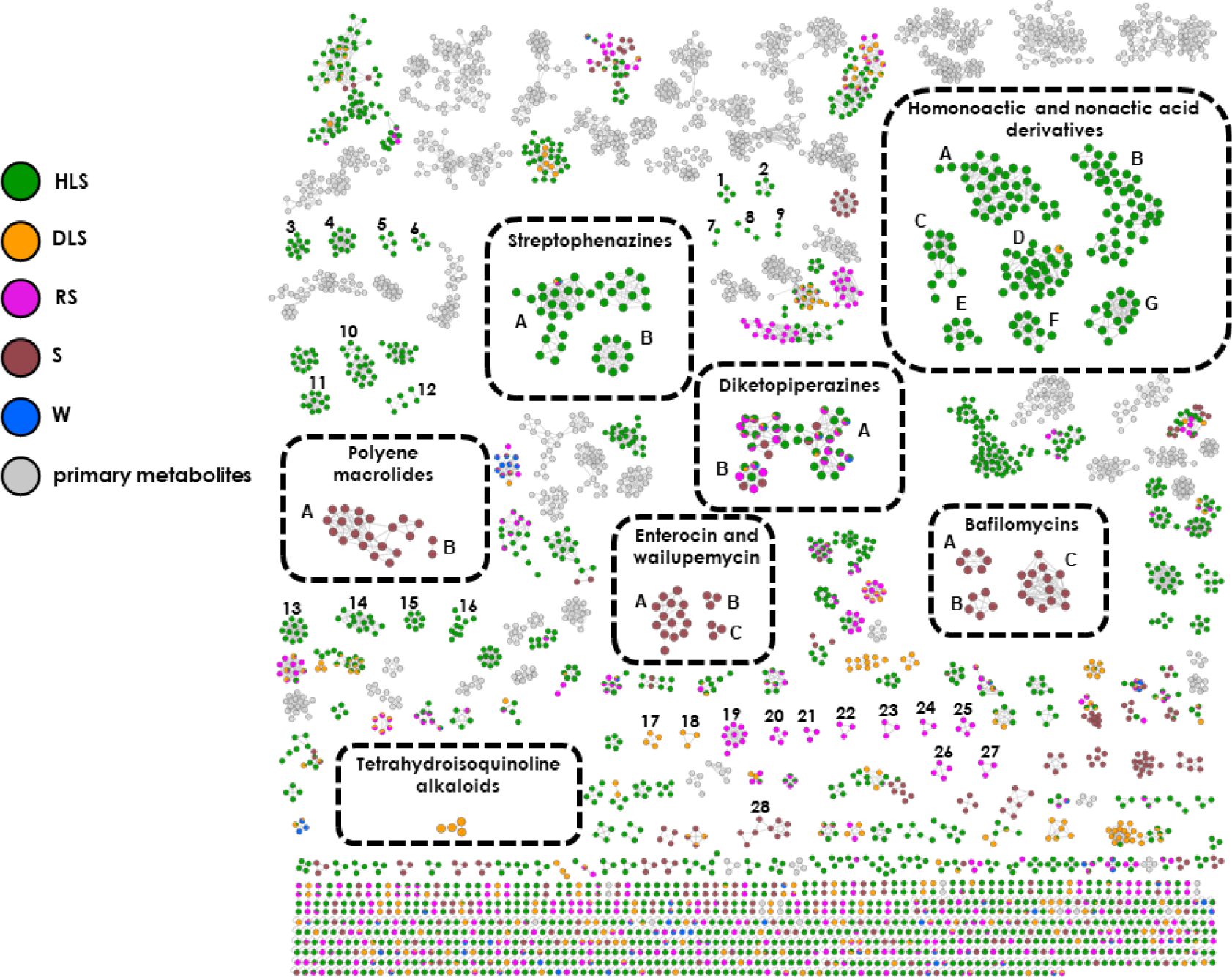
Feature-based molecular network analysis of the two media extracts of the most active bacteria isolated from the surface of *Zostera marina* healthy leaves (HLS), decaying leaves (DLS), roots (RS), as well as the sediment (S) and seawater (W) references. Clusters of primary metabolites are shown in grey color. Annotated molecular families (MF) are framed.

The second and the largest MF represented with nine clusters identified in *Streptomyces* sp. 131 was assigned to protonated (clusters A and E) and sodiated (clusters B-G) adducts of homononactic and nonactic acid type polyketides (Fig. 5). Notably, clusters A and D included linear ones (seco-dinactin, homononactyl homononactate, bonactin, nonactyl nonactoate, ethyl homononactyl nonactate), while cluster B, C, E-G included the cyclic derivatives (monactin, nonactin, dinactin, trinactin, tetranactin, macrotetrolide C (Fig. 5 and Supplementary Table S14). Homononactic and nonactic acid are the building units of the macrocyclic tetralactones (also called as macrotetrolide antibiotics or nactins), such as monactin, nonactin, dinactin, trinactin, tetranactin. Nactins are ionophoric compounds first reported from a soil-derived *Streptomyces* sp. showing a wide range of biological activities (Zizka, 1998). They potently inhibit Gram-positive bacteria, including *S. aureus*, *E. faecalis* (Kusche et al., 2009; Shishlyannikova et al., 2017) and the sporogenesis and the motility of the zoospores of peronosporomycetes, known pathogens of plants, fishes and vertebrates (Islam et al., 2016). Tetranactin (cluster C) and dinactin (cluster F) are potent inhibitors of fungal phytopathogens (Liu et al., 2019; Zhang et al., 2020). Bonactin, the linear homononactic acid derivative is of particular interest. First isolated from a marine sediment-derived *Streptomyces* sp. (Schumacher et al., 2003), bonactin exhibits strong antibiotic activity towards both Gram-positive and -negative bacteria, fungi (Schumacher et al., 2003; Huang et al., 2015) and inhibits the mycelial development the phytopathogen *Magnaporthe oryzae* pathotype *triticum* (Rabby et al., 2022). These metabolites can be linked the observed bioactivity of *Streptomyces* sp. 131 against Enterococci, MRSA and phytopathogens (Supplementary Table S12). Although the homononactic/nonactic acid derivatives were expressed in both culture media (MA and GYM), the majority of the nodes were exclusive or more abundant in the MA medium extract (Supplementary Fig. S1). This possibly underlies the stronger activity observed with the MA extract of this isolate (Supplementary Fig. S1, Supplementary Table S12). Many additional MFs exclusive to HLS-derived *Streptomyces* sp. 131 were found but none of them returned hits in databases (Fig. 5 and Supplementary Fig. S1, clusters 1-2, exclusive to GYM extract, clusters 3-6, shared by MA and GYM media extracts).

The only other HLS-derived bioactive Gram-positive bacterium, *Alkalihalobacillus hwajinpoensis* strain x007, accounted for three unique molecular family, produced only in MA medium (Fig. 5 and Supplementary Fig. S1, clusters 7-9) but no compounds could be annotated. Eighteen HLS-associated Gram-negative bacterial extracts selected for metabolome analyses produced mainly fatty acids, phospholipids and aminolipids, the common and abundant components of Gram-negative bacteria (Chianese et al., 2018). A number of MFs (detected in both culture media extracts) were produced by *Vibrio metschnikovii* strain 248 (Fig. 5 and Supplementary Fig. S1 clusters 10-15) or *Cobetia* sp. x001 (Fig. 5 and Supplementary Fig. S1, cluster 16), but no specific secondary metabolite could be annotated.

Two clusters mainly shared by HLS-, DLS-, RS- and reference isolates were annotated as diketopiperazines. From the large cluster A, we were able to annotate cyclo(L-Val-L-Phe), cyclo(L-Pro-L-Leu), cyclo(L-Pro-L-Val), and cyclo(L-Trp-L-Pro) from the cluster B (Fig. 5). Diketopiperazines are small cyclic-dipeptides that are produced by fungi, bacteria and other organisms. They display a wide range of biological activities, including antibacterial, antifungal and anticancer (Borthwick, 2012; Bojarsja et al., 2021). Cyclo(L-Pro-L-Val) and cyclo(L-Trp-L-Pro) isolated from various bacterial and fungal species showing antibiotic activity against various bacteria including *S. aureus* (Qi et al., 2009; Borthwick, 2012; Alshaibani et al., 2017). Cyclo(L-Trp-L-Pro) is an antifungal agent against human pathogenic fungi (Ben Ameur Mehdi et al., 2009; Borthwick, 2012; Nishant Kumar et al., 2014). Cyclo(L-Val-L-Phe), first reported from a sponge-derived *Pseudoalteromonas* sp., is a quorum sensing regulator (Guo et al., 2011). The cyclic-dipeptide cyclo(L-Pro-L-Leu), is of particular interest. Isolated from many bacterial and fungal species, including *Streptomyces* spp., it potently inhibits twelve vancomycin-resistant enterococci (VRE) strains (including *E. faecium* and *E. faecalis*), and several fungal phytopathogens including *M. oryzae* (Rhee, 2002; Rhee, 2003). Diketopiperazine clusters were expressed in both culture media (MA and GYM, Supplementary Fig. S1).

Bacteria retrieved from the surface of decaying eelgrass leaves (DLS) presented several small MFs, some of which were shared with the HLS-isolates. The majority of the MFs remained unannotated, except for one node each in streptophenazine and homononactic/nonactic acid clusters, and 14 nodes in diketopiperazine clusters (Fig. 5). One cluster that was unique to GYM extract of DLS isolate *Streptomyces griseorubens* strain x025 was attributed to tetrahydroisoquinoline alkaloids by manual dereplication (Fig. 5 and Supplementary Fig. S1). Two nodes were assigned as the protonated adduct of naphthyridinomycin *m/z* 418.1969, originally isolated from a soil-derived *Streptomyces lusitanus* (Kluepfel et al., 1975) and bioxalomycin β2 *m/z* 400.1866, sourced from a marine *Streptomyces* sp. (Bernan et al., 1994). Both compounds are potent and broad-spectrum antibiotics against Gram-positive and Gram-negative pathogens, including MRSA and *E. faecium* (Bernan et al., 1994; Zaccardi et al., 1994; Scott and Williams, 2002). The putative detection of these compounds could relate the observed moderate bioactivities towards these test pathogens (Supplementary Table S12). A few clusters were detected in DLS-derived Gram-negative bacterial extracts (Fig. 5, clusters 17-18), but none could be linked to a known compound reported in GNPS or other databases.

Overall, a few MFs that were exclusive to bacterial RS-isolates were observed in the FBMN (Fig. 5). Many clusters were shared with the bacterial isolates of HLS, DLS and sediment reference. The majority of the MFs remained unannotated, except for one node each in streptophenazines and 25 nodes in diketopiperazine clusters (Fig. 5). One cluster was specific to the single Gram-positive bacterium, *Bacillus* sp. strain 725 (Fig. 5, cluster 19). The MFs belonging to Gram-negative RS-derived bacteria were grouped in small clusters (Fig. 5, 20-27) or appeared as singletons. None of the MFs could be annotated.

Moving to reference isolates, several clusters were expressed solely in sediment-derived (S) bacterial extracts (Fig. 5). Few MFs were shared with the HLS-, DLS-, RS- and W-isolates. Most of them remained unannotated, except for one node in streptophenazine cluster and 17 nodes in diketopiperazine clusters. Two clusters, expressed in both media extracts of *Streptomyces scopiformis* 744 (Fig. 5, Supplementary Fig. S1) were attributed to polyene macrolides, in which we annotated the sodium adducts of dermostatin B *m/z* 757.4503 and PN00053 *m/z* 773.4456 (Fig. 5, clusters B and A, respectively). Dermostatins that were first isolated from a soil-derived *Streptomyces virdigreseus* are broad-spectrum fungicides, especially towards human pathogens involved in dermatomycosis (Thirumalachar and Menon, 1962). PN00053 originates from a soil-associated *Streptomyces* sp. and displays a broad-spectrum antifungal (but no antibacterial) activity (Vartak et al., 2014). No detailed activity against the pathogens tested in this work has been reported.

The other three clusters uniquely produced by *S. scopiformis* 744, were putatively annotated as bafilomycins, including the sodium adducts of bafilomycins A1 (*m/z* 645.3982), A2 (*m/z* 659.4135; Fig. 5, cluster C) and D (*m/z* 627.3869, Fig. 5, cluster B) plus 17,18-dehydro-19,21-di-O-methyl-24-demethylbafilomycin A1 (*m/z* 641.4033, Fig. 5, cluster A). Bafilomycins are a family of macrolide antibiotics, first reported from a soil-derived *Streptomyces griseus* (Werner et al., 1984), inhibiting Gram-positive bacteria, yeast, human and plant fungal pathogens (Werner et al., 1984; Zhang et al., 2011). Notable is the broad-spectrum activity against fungal phytopathogens, including *M. oryzae* reported for bafilomycin A1 and D (Zhang et al., 2011; Xu et al., 2013), which is in line with the activity observed here for both media extracts of *S. scopiformis* 744 against phytopathogens Pi and Mg (Supplementary Table S12). Clusters A and B are unique to MA medium, while cluster C is expressed in both media (Supplementary Fig. S1).

Three α-pyrone containing polyketide clusters (i.e., enterocin and wailupemycins) were exclusively detected in the second sediment-associated *Streptomyces* sp. strain 738 (Fig. 5). From cluster A, we annotated enterocin (*m/z* 445.1130) and 8-deoxyenterocin (*m/z* 429.1186). Enterocin, first reported from the soil-derived *Streptomyces* sp. (Miyairi et al., 1976), is a bacteriostatic agent against Gram-positive and Gram-negative bacteria, including *E. coli* (Miyairi et al., 1976). The other two small clusters belonged to wailupemycins, sourced from various marine-derived *Streptomyces* sp. (Xiang et al., 2002). We annotated wailupemycins F (cluster B) and G (cluster C, Fig. 5) that are also bacteriostatic agents like enterocin (Sitachitta et al., 1996).

One small cluster (cluster 28, Fig. 5) unique to sediment-derived Gram-negative bacterium *Pseudoalteromonas ulvae* XSBU1 was observed, but it returned no hits in the metabolome analyses.

#### 3.3.2. Metabolome of endophytic bacteria

The extracts of HLI- and DLI-derived bacteria dominated the metabolome of the endophytic bacteria (Fig. 6). Two HLI-derived Gram-positive strains (*Bacillus* sp. x015 and *Bacillus* sp. 742) returned hits in database searches.

**Fig. 6.**
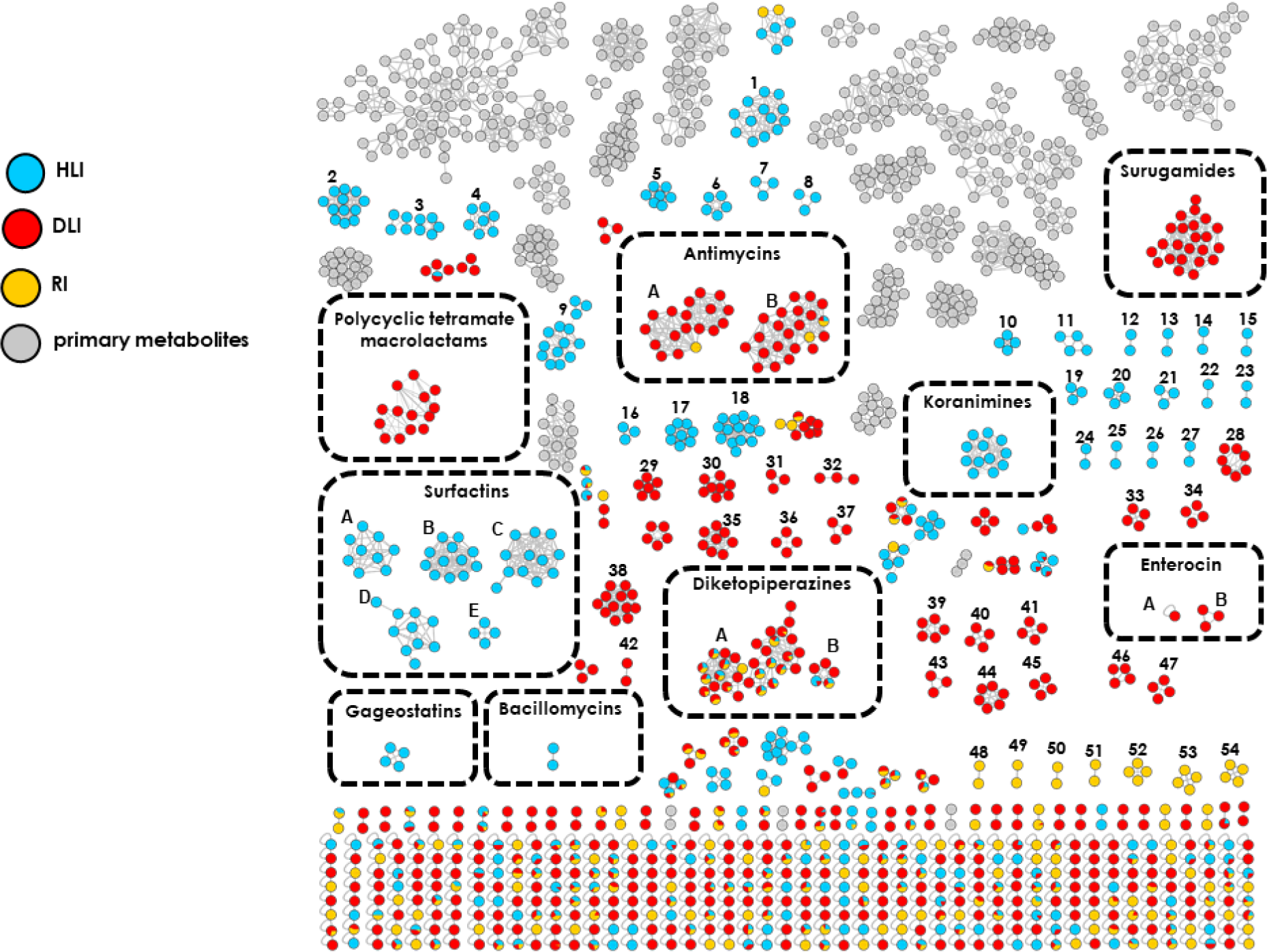
Feature-based molecular network showing ions detected in the most active bacterial isolates from the inner tissues of *Zostera marina* healthy leaves (HLI), decaying leaves (DLI), roots (RI) obtained from both culture media. Clusters grouping primary metabolites are coloured in grey. Annotated molecular families (MF) are framed.

Putatively identified compounds for *Bacillus* sp. x015 included lipopeptides, including both cyclic (bacillomycin and surfactin clusters) and linear (gageostatin cluster) ones (Fig. 6). C15-bacillomycin D, first isolated from the culture of terrestrial-derived *B. subtilis* (Peypoux et al., 1984) was putatively annotated in the two-node cluster detected only in the MA medium (Supplementary Fig. S2). Bacillomycins exhibit strong activity against *Staphylococcus* spp., including MRSA (Nam et al., 2021), that may underlie the antibacterial properties of the MA extract of *Bacillus* sp. x015 (Supplementary Table S12). Gageostatins A and B were mainly identified in the MA extract of the same strain (Supplementary Fig. S2). Originally discovered from the fermentation broth of a marine *B. subtilis* strain, gageostatins show moderate antimicrobial properties against pathogenic bacteria, including *S. aureus*, and phytopathogenic fungi (Tareq et al., 2014).

The large MF of surfactins solely produced by *Bacillus* sp. x015 in both culture media was clustered in six networks comprising different adducts, i.e., [M+NH_4_]^+^ (cluster A), [M+H]^+^ (clusters B, D, E), and [M+Na]^+^ (clusters C). This large class of cyclic peptidolipids with surfactant properties were first identified in a terrestrial *B. subtilis* (Arima et al., 1968) and shown to inhibit Gram-positive (e.g., *Staphylococcus, Enterococcus*), Gram-negative (*Vibrio, Escherichia*, and *Salmonella*; Oka et al., 1993; Kim et al., 2009) and phytopathogenic (e.g., *Xanthomonas campestris*) bacteria (Bartal et al., 2023), in line with the results of the present study (Supplementary Table S12). Many additional MFs exclusive to HLI-derived *Bacillus* sp. x015 were found, including the clusters 1-7 (common to both MA and GYM extracts), cluster 8 (exclusive to GYM extract) and clusters 9-15 (exclusive to MA extract), however none of them returned hits in databases (Fig. 6, Supplementary Fig. S2).

The cyclic imine heptapeptides koranimine and koranimine B that clustered in a single network were exclusive to the other HLI-derived *Bacillus* sp. strain 742, being particularly abundant in its MA extract (Fig. 6 and Supplementary Fig. S2). Koranimine (*m/z* 804.5026), originally isolated from a soil-derived *Bacillus* sp., is known for potent nematocidal activity (Evans et al., 2011; Montecillo and Bae, 2022). Few other clusters unique to HLI-derived *Bacillus sp*. strain 742, were shown in the network (Fig. 6 and Supplementary Fig. S2), but none of them could be annotated. This includes the clusters 16-17 (shared by MA and GYM extracts), clusters 18-22 (exclusive to MA extract) and clusters 23-26 (exclusive to GYM extract).

The GYM extract of another HLI-isolate, *Bacillus* sp. 704 showed only one exclusive cluster (Fig. 6, cluster 27), which could not be annotated.

Two clusters, shared by all endophytic bacteria, but present more abundantly in DLI-derived isolates belonged to diketopiperazines (Fig. 6). We annotated cyclo(L-Val-L-Phe), cyclo(L-Pro-L-Leu), cyclo(L-Pro-L-Val) in cluster A and cyclo(L-Trp-L-Pro) in cluster B. The majority of the nodes were exclusive or more abundant in the MA medium extract (Supplementary Fig. S2).

Of the DLI-isolates analysed, *Streptomyces* sp. strain x019 showed the most interesting chemistry. Annotated MFs included a large surugamide cluster that was detected in both media extracts (Fig. 6, Supplementary Fig. S2). Surugamides are cyclic peptides first reported from a deep-sea sediment-associated *Streptomyces* sp. (Takada et al., 2013) and exert anticancer activity (Almeida et al., 2019). Two other large clusters mostly identified in the same isolate were annotated as antimycin and its deformylated derivatives (Fig. 6, clusters A and B, resp.). Antimycin antibiotics isolated from soil- and marine sediment-derived *Streptomyces* sp. are potent fungicides, especially against phytopathogens, e.g., *Magnaporthe oryzae* (Belakhov et al., 2018; Paul et al., 2022). Both clusters were dominant in the MA extracts, possibly justifying the strong activity of the MA extracts against phytopathogens, particularly Mg (Fig. 6, Supplementary Fig. S2 and Supplementary Table S12). An additional cluster unique to *Streptomyces* sp. x019 belonged to polycyclic tetramate macrolactams, including ikarugamycin, ikarugamycin epoxide, maltophilin and dihydromaltophilin (Fig. 6). Ikarugamycin (*m/z* 479.2906) and ikarugamycin epoxide (*m/z* 495.2857), first reported from soil-derived *Streptomyces* sp. (Jomon et al.,1972; Bertasso et al., 2003) show moderate activities against Gram-positive bacteria, e.g., *S. aureus* (Bertasso et al., 2003). Maltophilin (*m/z* 511.2804) and dihydromaltophilin (*m/z* 513.2968), originate from rhizosphere- and soil-derived bacteria are broad spectrum fungicides, also inhibiting the phytopathogenic oomycete *Plasmopara viticola* (Jakobi et al., 1996; Graupner et al., 1997). Although this cluster was detected in both media, the majority of the nodes were solely or more abundantly expressed in the GYM extracts (Supplementary Fig. S2). Finally, two nodes in the three-node cluster were annotated as enterocin and 8-deoxyenterocin (Fig. 6, cluster B), while the sodium adduct of the latter was annotated as a singleton (Fig. 6, A). Enterocin was unique to GYM medium, while 8-deoxyenterocin to MA medium (Supplementary Fig. S2). Ten additional clusters (28-37, Fig. 6) exclusive to *Streptomyces sp.* strain x019 were also observed. As they did match to any known compounds, they may be new compounds with potential contribution to the observed activities herein.

The remaining DLI-derived Gram-positive bacteria (*Paenibacillus terrae* strain 436 and *Oceanobacillus* sp. strain 332b), produced clusters 38-41 (exclusive to *P. terrae* 436), and clusters 42-43 (exclusive *Oceanobacillus* sp. 332b), but no annotation was possible. Gram-negative bacterial DLI-isolates displayed many nodes clustered in lipid families and small unannotated clusters (44-47).

All selected bioactive bacterial root endophytes (from RI) were Gram-negative, showing only few specific clusters (48-54) that could not be matched to any known compound. Several MFs were shared with the HLI- and DLI-isolates. The majority of the MFs remained unannotated, except for three nodes in antimycin cluster and 18 nodes in diketopiperazine clusters (Fig. 6).

#### 3.3.3. Metabolome of epiphytic fungi

Fig. 7 displays the global MN of all selected epiphytic fungi, while Supplementary Fig. S3 indicates the media effect (including Venn diagrams of the nodes fungal extracts) on their metabolome.

**Fig. 7.**
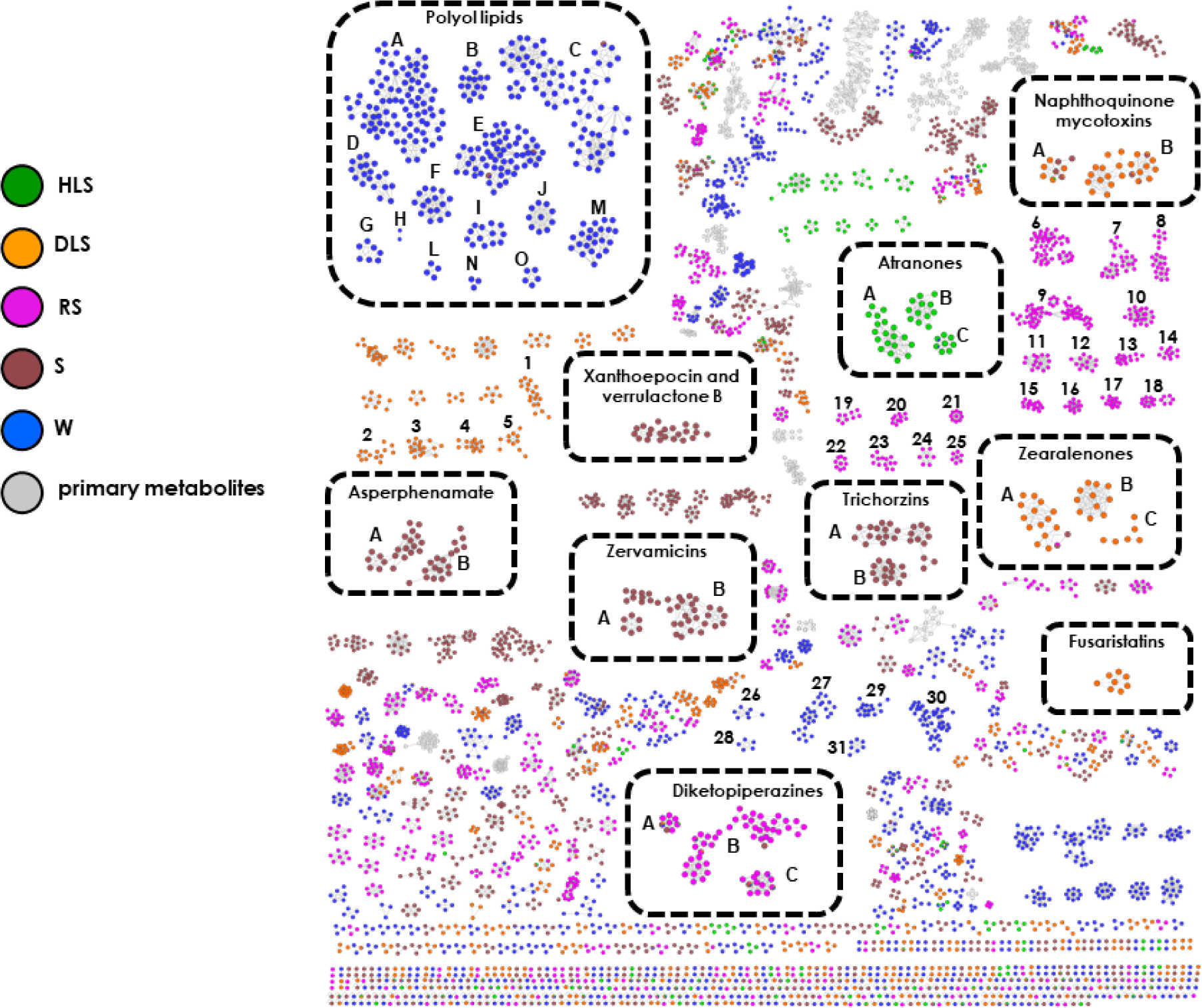
FBMN showing ions detected in the most active fungi isolated from the surface of eelgrass healthy leaves (HLS), decaying leaves (DLS), roots (RS), and the sediment (S) and seawater (W) references obtained from both culture media. Annotated molecular families (MF) are framed.

Of 12 clusters originating from the HLS-derived fungi, three could be dereplicated as sodiated (cluster A) and protonated (clusters C, B) adducts of atranone type diterpenoids (Fig. 7, Supplementary Fig. S3). Notably, atranone clusters were exclusive to HLS-associated *Parathyridaria* sp. strains 425a and 436, being present mostly in their PDA extracts (Supplementary Fig. S3). The atranones B and O derive from terrestrial and marine-derived *Stachybotrys* spp., respectively (Hinkley et al., 1999; Li et al., 2017). The recently reported atranones (but not those annotated herein) show antibacterial activity against MRSA and *E. faecalis* (Yang et al., 2019). The remaining clusters belonging to HLS-derived fungi could not be annotated.

Many MFs were produced by DLS-derived fungi (Fig. 7). However only few could be retrieved from databases, almost all of which deriving from *Fusarium* sp. strain 719. Naphthoquinone type terrestrial mycotoxins were represented in two clusters, i.e., dimeric rubrofusarin *m/z* 543.1289 (cluster A) and aurofusarin *m/z* 571.0877, fuscofusarin, *m/z* 557.1086, xanthomegnin, *m/z* 575.1187, morakotin D, *m/z* 557.0719, dimeric 9-hydroxyrubrofusarin-fuscofusarin, *m/z* 573.1031, dimeric rubrofusarin-9-hydroxyrubrofusarin, *m/z* 559.118 and 3,4,3’,4’-bisdehydroxanthomegnin *m/z* 571.0880 (cluster B, Fig. 7, Supplementary Fig. S3). Some nodes were shared with the sediment-associated fungus *Penicillium olsonii* 809, as quinones are common constituents of filamentous fungal genera e.g., *Aspergillus*, *Fusarium* and *Penicillium*. (Christiansen et al., 2021). Naphtoquinones exert a wide range of bioactivities including selective inhibition of Gram-positive bacteria, e.g., *S. aureus* (Medentsev and Akimenko, 1998; Wang et al., 2019). The majority of nodes in cluster B were expressed in both culture media M34 and PDA (Supplementary Fig. S3).

Another small, single network belonging to *Fusarium* sp. 719 was putatively assigned to fusaristatin A and B type cyclic lipopeptides (Fig. 7). First reported from a plant associated *Fusarium* sp., both compounds show cytotoxicity (Shiono et al., 2007; Evidente, 2022). Fusaristatin A inhibits the plant pathogen *Glomerella acutata* (Li et al., 2016), and may contribute to strong anti-phytopathogenic activities of *Fusarium* sp. strain 719 reported herein. As displayed in Supplementary Fig. S3, most of the nodes were exclusive to M34 medium.

Three additional MFs mainly detected in the M34 extract of *Fusarium* sp. strain 719 were annotated as zearalenone type polyketides, including zearalenol, zearalenone and 5β-hydroxyzearalenone (clusters A, B, C, resp.; Fig. 7). The first two compounds originate from a terrestrial *Fusarium* sp., while 5β-hydroxyzearalenone originates from a seagrass-derived *Fusarium* sp. (Urry et al., 1966; Stipanovic and Schroeder, 1975; Saetang et al., 2016). Zearalenone is a mycotoxin inhibiting the growth of competing fungi during the saprophytic growth phase (Utermark and Karlovsky, 2007). In addition, zearalenone shows a weak activity against MRSA (Seatang et al., 2016).

Five clusters (1-5, Fig. 7) were exclusively produced by DLS-derived *Cladosporium halotolerans* 233. However, no annotation was possible, implying that they may represent new, undescribed MFs.

The only RS-isolate selected for metabolome analysis, *Cystofilobasidium bisporidii* 417, was chemically prolific, with many MFs being unique for it, but none could be annotated (Fig. 7, clusters 6-25).

Two clusters, shared by all surface associated fungi, but more abundant in the only RS-derived isolate, belonged to diketopiperazines. The following metabolites were annotated: cyclo(L-Pro-L-Val) in cluster A; cyclo(L-Val-L-Phe) and cyclo(L-Tyr-L-Pro) in cluster B; and cyclo(L-Pro-L-Leu) in cluster C (Fig. 7). Cyclo(L-Tyr-L-Pro), isolated from several microorganisms, shows antimicrobial properties against Gram-positive (incl. *S. aureus*) and Gram-negative bacteria, and against fungal human and plant pathogens (Kumar et al., 2013; Wattana-Amorn et al., 2016). The majority of the nodes were exclusive or abundant in the M34 medium extract (Supplementary Fig. S3).

A number of molecular clusters expressed in the sediment-associated (S) reference fungal extracts were observed. Some of them were shared with HLS-, DLS-, RS- and W-isolates (Fig. 7). Two annotated clusters were exclusive to M34 extract of S-isolate *Trichoderma viride* strain 407. Trichorzin and zervamicins type peptaibols (linear peptides) were represented with two clusters each (Fig. 7), we were able to annotate trichorzin PA-VI and PA-VIII (cluster A), and trichorzin HA-2 (cluster B; Fig. 7). Originally isolated from a soil-derived *T. harzianum,* trichorzins PA series exhibit potent activity against Gram-positive bacteria including *S. aureus* (Duval et al., 1997), while trichorzin HA series inhibit phytopathogen *Sclerotium cepivorum* (Goulard et al., 1995). The other MF was represented by zervamicin IB (cluster A), plus zervamicin IIB and II-4 (cluster B). Zervamicins were originally reported from marine fungus *Emericillopsis microspora* and are mainly active against Gram-positive bacteria, including *S. aureus* (Argoudelis et al., 1974; Rinehart et al., 1981).

The small linear amino acid ester asperphenamate and its phenylalanine derivative were annotated in two clusters (as [M+H]^+^ and [M+Na]^+^ adducts) unique to both media extracts of the other sediment-derived fungus, *Penicillium olsonii* strain 809 (Fig. 7). Asperphenamate, first discovered from the terrestrial *Aspergillus flavipes*, is produced by wide range of *Aspergillus* and *Penicillium* spp., its tyrosine derivative was sourced from a marine-derived *Penicillium bialowiezense* (Clark et al., 1977; Kildgaard et al., 2014). Asperphenamates are best known as anticancer agents (Frisvad et al., 2013) and are inactive against *S. aureus* and *E. coli* (Zheng et al., 2013).

The dimeric polyketide family (xanthoepocin and verrulactone B) was unique to *P. olsonii* 809 (Fig. 7). Xanthoepocin, a yellow pigment common in many Penicillii (Igarashi et al., 2000) is an antibiotic active against pathogenic yeasts and multi-drug resistant Gram-positive bacteria, including MRSA and the VRE *E. faecium* (Vrabl et al., 2022). Also, verrulactone B, which was discovered from the soil fungus *P. verruculosum* is a potent MRSA inhibitor (Kim et al., 2012). This MF was predominantly expressed in the PDA medium (Supplementary Fig. S3) that may contribute to the potent activity of the PDA extract of *P. olsonii* 809 against aquatic, fecal and phytopathogenic panels (Supplementary Table S13).

Seawater (W) associated fungi produced a large number of MFs (Fig. 7). The largest and annotatable cluster, mainly identified in *Aureobasidium pullulans* 813, belonged to polyol lipids (glycolipids) such as liamocins (clusters A-E, I, L), exophilins (clusters F, G, M) and halymecins (H, J, N and O), through extensive *in*-*silico* and manual dereplication, most abundantly present in M34 extracts (Fig. 7, Supplementary Fig. S3). The liamocins were repeatedly isolated from *A. pullulans* strains (Manitchotpisit et al., 2011; Leathers et al., 2015) and exert selective activity against *Streptococcus* spp. (Price et al., 2013; Bischoff et al., 2015). In contrast, exophilins, originating from marine sponge-derived black yeast *Exophiala pisciphila,* inhibit Enterococci and *S. aureus* (Bischoff et al., 2015). Among the annotated halymecins, halymecin A, first isolated from algicolous fungus *Fusarium* sp., inhibits Gram-positive and Gram-negative bacteria (i.e., *Enterococcus faecium* and *Klebsiella pneumoniae*; Chen et al., 1996; Le Dang et al., 2014). Halymecin F, first reported from the biocontrol fungus *Simplicillium lamellicola* is a strong inhibitor of phytopathogenic bacteria (Le Dang et al., 2014). These results are in line with the moderate to good bioactivity observed for *A. pullulans* 813 extracts against environmental (e.g., *Vibrio ichthyoenteri*), fecal (*Enterococcus casseliflavus*) and phytopathogenic (*Xanthomonas campestris*) bacteria (Supplementary Table S13). Notably, few nodes in liamocin clusters A, C and F were shared with the extracts of another W-derived fungus, *Sarocladium* sp. 815. Six clusters (Fig. 7, clusters 26-31) were exclusive for this strain, but none could be annotated.

#### 3.3.4. Metabolome of endophytic fungi

The FBMN of the only bioactive DLI-associated fungus *Acrostalagmus luteoalbus* strain 720 (Fig. 8) revealed the presence of a wide variety of the epipolythiodioxopiperazine toxins, i.e., 11’-deoxyverticillin A, verticillin B and C in cluster A, chetracin C and C in cluster B and chetocin in cluster C (Fig. 8). Epipolythiodioxopiperazine clusters (A-C) were common to both culture media extracts (M34, PDA; Fig. 8). Epipolythiodioxopiperazines such as chaetocins, chaetracins and verticillins are mycotoxins all showing remarkable antitumor activities (Li et al., 2012; Huber, 2022). Chaetocin A, identified from *Chaetobium* spp., was also reported to inhibit *S. aureus,* penicillin-resistant *S. aureus* and *E. faecalis* (Hauser et al., 1970; Dwibedi et al., 2023).

**Fig. 8.**
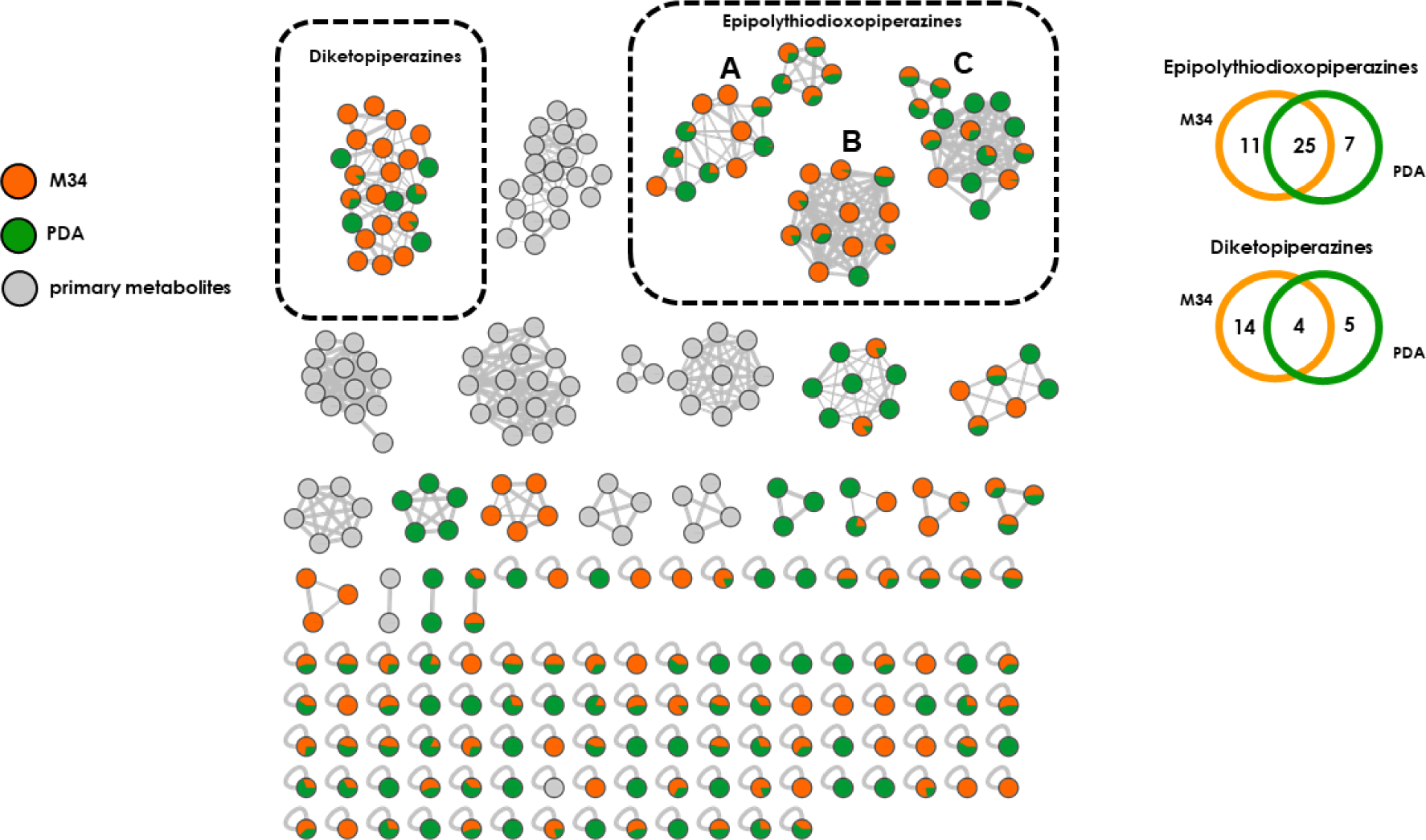
FBMN showing ions detected in M34 and PDA media extracts of *Acrostalagmus luteoalbus* sourced from DLI. Clusters containing primary metabolites are coloured in grey. Annotated molecular families (MF) are framed. The Venn diagram shows the number of exclusive and shared nodes of two culture media used for the annotated molecular families.

The second large MF was dereplicated as diketopiperazines, also expressed more abundantly in the M34 medium (Fig. 8). Within this class, we annotated cyclo(L-Pro-L-Val), cyclo(L-Pro-L-Leu), cyclo(L-Tyr-L-Pro) and cyclo(L-Phe-L-Pro). Cyclo(L-Phe-L-Pro), previously isolated from *Pseudomonas fluorescens* and *P. alcaligenes*, is a potent antibiotic inhibiting VRE strains (including *E. faecium, E. faecalis*) and *S. aureus.* It also exhibits antifungal activity against human and plant pathogens (Rhee et al., 2001; Ström et al., 2002).

A number of small clusters that were observed in the FBMN of *A. luteoalbus* remain unannotated. Considering the highly broad and often potent activity of *A. luteoalbus* extracts, the observed activities may rather be a non-specific toxicity towards the test pathogens, possibly originating from epipolythiodioxopiperazines and diketopiperazines.

## 4. Discussions

In this study, we characterized the cultivable epiphytic and endopytic microbiome of the healthy and decaying leaves and the roots of Baltic seagrass *Z. marina*, then assessed their inhibitory effects against a large panel of pathogens and finally mined their metabolomes for identifying putative compounds responsible for the detected anti-pathogenic activities. Our culture-based approach allowed the isolation of 70 bacteria and 18 fungi that were assigned to numerous microbial taxa. An early culture-dependent study on bacterial epiphytes of the leaves and the roots of Japanese *Z. marina* (Kurilenko et al., 2001) reported the same four bacterial phyla as in our work, with heterotrophic Gram-negative bacteria dominating surface microbiomes of both tissues and comparably higher bacterial diversity in (young) leaves (Kurilenko et al., 2001). Most *Z. marina* microbiome studies however used culture-independent methods, mainly amplicon sequencing of the 16S rRNA gene. Several studies comparing the eelgrass leaf and root epibiome detected specific associations with these two tissues and a high abundance of Alpha- and Gammaproteobacteria (Crump and Koch, 2008; Fahimipour et al., 2017; Crump et al., 2018), which is in line with our results. Such approaches naturally detected a much broader bacterial diversity with many taxa being absent from our strain collection, such as Beta- and Deltaproteobacteria, Planctomycetes (Alain and Querellou, 2009; Garza and Dutilh, 2015). The observed higher bacterial diversity of leaf surfaces compared to root surfaces was however not detected in metagenomics-based studies (Ettinger et al., 2017; Fahimipour et al., 2017; Crump et al., 2018), hence may be a result of a culture-based approach. Compared to phyllosphere, the eelgrass endosphere harbored reduced diversity but hosted specific taxa (e.g., *Mycolicibacterium* and *Paenibacillus* for DLI; *Breoghania* and *Hoeflea* for RI). The lowest diversity was observed within healthy leaves (5 *Bacillus* spp.) indicating that the inner eelgrasss tissues are well protected against potential pathogen invasions. This was further supported by comparably higher bacterial abundance in decaying leaf endosphere, indicating that leaf tissue degradation facilitates the entry of microorganisms into inner leaf tissues. Our results were in line with previous studies; the eelgrass microbiome was different than its surrounding seawater and sediment (Jensen et al., 2007; Sanders-Smith et al., 2020).

Similar findings were reported for *Z. marina*-associated fungi (e.g., Kirichuk and Pivki, 2015; Ettinger and Eisen, 2019; Ettinger and Eisen 2020; Ettinger et al., 2021), although fewer studies have been conducted so far. Most of the fungal taxa isolated in this study were previously isolated from *Z. marina*, including the common ones, e.g., *Cladosporium*, *Penicillium* and *Trichoderma*, as well as more rare genera, e.g., *Acrostalagmus* and *Paradendryphiella* (Kirichuk and Pivki, 2015; Ettinger and Eisen 2020). These culture-based studies also revealed high proportions of Ascomycetes in all tissues and differential fungal communities when comparing to ambient seawater and sediment (Kirichuk and Pivki, 2015; Ettinger and Eisen 2020). Dominance of Ascomycete fungi was also shown with culture-independent analyses and interestingly, high abundance of Basidiomycetes was detected in root tissues (Ettinger and Eisen, 2019). This is in line with our results, since the only Basidiomycete we isolated from *Z. marina* (*Cystofilobasidium bisporidii* strain 417) was retrieved from the root surface.

Overall, the majority of eelgrass associated microbes inhibited the growth of at least one pathogen. Gram-positive pathogens, such as the aquatic pathogen *Leifsonia aquatica*, fish pathogen *Lactococcus garvieae*, fecal *Enterococcus* spp. and human pathogen MRSA were the most susceptible test organisms towards eelgrass-associated bacteria and fungi. Gram-negative bacteria are in general difficult to inhibit, still, the seaweed pathogen *Algicola bacteriolytica,* fish pathogen *Shewanella algae* and multiple *Vibrio* spp. (e.g., *V. fischeri*, *V. ichthyoenteri*, *V. parahaemolyticus*) were hit, moderate to high levels, by some extracts. A number of extracts showed significant potency towards the phytopathogenic fungi *Phytophthora infestans* and *Magnaphorte grisea*. In general, HLS bacteria represented the largest and the most active group with broader antimicrobial activity (Fig. 4 and Supplementary Tables S4-S7). The potent antimicrobial bacterial isolate *Streptomyces* sp. 131 was an epiphyte of the healthy *Z. marina* leaves. Despite their lower potency, many gammaproteobacterial (Gram-negative) healthy leaf epiphytes, plus other surface associated bacteria (from DLS and RS) also proved active. The inner tissues also hosted diverse bacteria with antimicrobial properties. *Streptomyces* sp. strain x019 deriving from decaying leaf endosphere (DLI) showed a broad-spectrum activity particularly towards phytopathogenic and aquatic panel, including the majority of *Vibrio* spp. Root endopyhtic bacteria were a small cohort with moderate, narrow-spectrum activity towards few aquatic pathogens (*Vibrio* spp.), enterococci and MRSA. The most active fungi were those associated with DLS, with *Cladosporium halotolerans* 233 and *Fusarium* sp. 719 being potently active against plant pathogens. As mentioned before, the very broad antibiotic activity of the decaying leaf endophyte *Acrostalagmus luteoalbus* is likely to originate from a non-specific effect. Also, few sediment-derived bacteria (e.g., *Streptomyces* spp. 738 and 744) and fungi (e.g., *Penicillium olsonii*) displayed antimicrobial potential.

FBMN based untargeted metabolomics approach allowed i) analysing qualitative and semi-quantitative differences in chemical machinery of the active strains isolated from different plant parts, ii) comparison of different strains of the same genus and iii) assessment of the impact of the media on biosynthetic capacity of the isolates. Wide variety of complex secondary metabolite types, e.g., polyketides, alkaloids, terpenes, and various peptide classes were identified. Despite the employment of massive computational and *in silico* metabolomics and manual dereplication approaches using public and commercial databases, compound annotation rates were low, suggesting the presence of new chemistry in the extracts. Several, mainly Gram-positive bacterial isolates and common fungal taxa reputed with rich chemistry, e.g., *Streptomyces* or *Penicillium* spp. returned hits in database searches. Notably, we were generally able to dereplicate only few nodes in large networks, indicating the existence of many new, potentially bioactive members of the knowns. We also detected numerous clusters produced by both bacteria or fungi that could not be assigned to any class, hence are likely to be novel molecular families. Gram-negative bacteria were very rich in lipids, which may have masked their less abundant secondary metabolites.

The culture media used made significant qualitative and quantitative differences in the chemical inventory (and the antibiotic activity) of the isolates. GYM medium appeared to be favorable for overall bioactivity rate of bacteria, as the number of bioactive GYM-sourced extracts were 3-fold of the MA extracts, but the highest potency generally derived from the MA extracts. In fungi, growth media had a less prominent influence on the bioactivities, but the PDA extracts often exerted better activity. In some cases, MFs were expressed in both media, however generally one medium was favored for biosynthesis of unique MFs. The impact of medium is also noticed in the extracts of eelgrass-associated *Streptomyces* spp. with major differences in their antibiotic potency. The nutrient-poor medium MA medium often led to a higher potency and larger chemical space. *Streptomyces* strains 131 (HLS) and x019 (DLI) dominated the global metabolome of 5 bioactive *Streptomyces* strains (Supplementary Fig. S4) with striking differences in the chemistry and antibiotic potential of different strains. Epiphytic *Streptomyces* sp. 131 had generally high potency but narrow spectrum towards aquatic panel, but broader spectrum towards the remaining pathogens. DLS-originated *S. griseorubens* x025 had lower level but much broader activity towards all four test panels. DLI-associated *Streptomyces* sp. x019 had overall the broadest activity profile with highest potential towards multiple *Vibrio* spp. and the phytopathogen *Magnaphorte grisea.* Notably, several large MFs annotated in *Streptomyces* spp. are known for their antibiotic properties against human pathogens (e.g., *S. aureus*, MRSA) and/or pyhtopathogens, however their effect towards aquatic pathogens has remained almost completely unknown. Hence, it is possible that the described molecules and their new congeners herein are of ecological importance towards aquatic/waterborne pathogens.

Seagrasses influence their surrounding environments through multiple ecosystem services, ranging from nursery habitat provision, nutrient cycling, water purification by filtration to coastal protection and carbon sequestration (Webb et al., 2019; de los Santos et al., 2020). Another impact of seagrass meadows is the alteration of the microbial community structure, diversity and abundance in their vicinity (Webb et al., 2019; Mohapatra et al., 2022). High-throughput 16S rRNA gene amplicon sequencing studies on several seagrass genera revealed different compositions of seawater and sediment microbiome between seagrass-vegetated and - unvegetated areas (Ettinger et al., 2017; Webb et al., 2019; Alsaffar et al., 2020; Sun et al., 2020). It is known that seagrasses support certain microbial communities and this effect is prominent in sediments. Colonization of seagrasses has been shown to promote accumulation of specific bacteria, such as diazotrophic and sulfate-reducing sediment bacteria, hence altering the relative abundances of specific bacterial lineages involved in benthic carbon and sulfur cycling (Sun et al., 2015; Seymour et al., 2018). Seagrass vegetation affect the vertical organization of microbial communities in sediment (Sun et al., 2020).

Less is known on the effect of seagrass meadows on seawater microbiome, but differences in bacterial taxa between eelgrass-present and eelgrass-absent seawater samples has been shown (Webb et al., 2019). A growing body of evidence show that seagrasses significantly reduce the load of various pathogenic bacteria in their surrounding water, thereby changing microbial community structure therein. By using a combination of 16S rRNA gene sequencing and *Enterococcus* assays, Lamb et al. (2017) showed that tropical eelgrass meadows (mixture of six seagrass species in the Spermonde Archipelago, Indonesia) significantly reduced bacterial pathogen load in their surrounding waters. Based on the observed 50% pathogen reduction in these coastal areas, seagrass meadows were attributed to provide a sanitation servic” as another benefit for the health of both humans and marine organisms. A microbiological survey carried out in the coastal waters of 270 Spanish beaches found decreased concentrations of fecal bacteria (*E. coli* and enterococci) in the meadows of Mediterranean seagrass *Posidonia oceanica* (Palazón et al., 2018). A similar effect was detected in the South China Sea for *Thalassia hemprichii* and *Enhalus acoroides* with *Salmonella* and *Vibrio* spp. being significantly reduced in these seagrass meadows compared to non-vegetated sites (Deng et al., 2021). A recent study reported substantially lower levels of several *Vibrio* spp. in eelgrass meadows in the Baltic Sea coast by applying a plate counting approach using selective agar media targeting pathogenic *Vibrio* spp. (Reusch et al., 2021). Although studies on pathogen load reduction from seagrass colonized sediments are not available, detection of significantly higher levels of potentially harmful pathogens (Enterobacteriales and *Vibrio* spp.) in non-vegetated (bulk) sediment compared to seagrass-vegetated area (Mohapatra et al., 2022) suggests a similar pathogen suppression effect in sediments.

Seagrasses such as *Zostera marina* and their associated microbiota are known to produce of antimicrobial metabolites (Kannan et al., 2010; Marhaeni et al., 2010; Ravikumar et al., 2012; Papazian et al., 2019; Petersen et al., 2019). Based on our results on the reduction of above-mentioned pathogens, especially by the epibiome of healthy eelgrass leaves, we now hypothesize *Z. marina*-associated microorganisms to play role in removal of pathogens from coastal waters. The reduction of bacterial contamination in seawater by seagrasses has been attributed to various mechanisms, such as grazing (filter-feeding or direct consumption) of plankton and bacteria by seagrass epiphytic fauna (Petersen and Heck Jr., 2001; González-Ortiz et al., 2014; Reusch et al., 2021), trapping and enhanced sedimentation rate of organic particles associated with bacteria (Fonseca et al., 1982; Worcester 1995; Deng et al., 2021) and exudation of antibacterial phytochemicals by seagrasses (Conte et al., 2021; Deng et al., 2021). Being sessile benthic marine plants, seagrasses are in constant contact with waterborne microbes, including a broad array of potentially harmful pathogens, hence they produce defensive secondary metabolites (Engel et al., 2002; Seymour et al. 2018). Phenolic compounds such as flavonoids and other phenylpropanoids that form the basis of defensive mechanisms in plants are abundant in seagrasses, providing protection against pathogen attacks, inhibiting microbial settlement or growth, and controlling biofouling (Harrison, 1982; Engel et al., 2002; Guan et al., 2017; Laabir et al., 2013; Subhashini et al., 2013). A sulfated flavone glycoside (thalassiolin) has been shown to protect the healthy leaves of *Thalassia testudinum* from infestation by a thraustochytrid protist (Jensen et al., 1998). Zosteric acid (sulfated *p*-coumaric acid), a strong antiadhesive/antifouling seagrass metabolite, is well known for preventing the attachment microorganisms, algae and bivalves to solid surfaces or plant leaves (Vilas-Boas et al., 2017). On the basis of metabolomics, surface mass spectrometry-based chemical imaging coupled with bioassay-guided isolation and functional bioassays, we showed that surface deployed sulfated flavonoids and the depside rosmarinic acid control microfouling of bacteria and yeasts on the leaf surfaces of *Z. marina* (Guan et al., 2017; Papazian et al., 2019). The antimicrobial effect of total seagrass tissue extracts against wide ranging environmental, marine and human pathogens have also been shown (Puglisi et al., 2007; Kumar et al., 2008; Ross et al., 2008; Kannan et al., 2010; Trevathan-Tackett et al., 2015).

Seagrass surfaces and internal tissues harbor an abundant and diverse microbiota mediating several metabolic exchanges and biogeochemical transformations essential for resource provision, plant growth and health (Ugarelli et al., 2017; Hurtado et al., 2019; Tarquinio et al., 2019; Tarquinio et al., 2021). Seagrasses support heterotrophic epiphytic microbes with exudation of nutrients, and in turn receive several benefits e.g., nitrogen fixation, production of polymer-degrading enzymes, and release of chemicals protecting the host from pathogens and biofouling by other organisms (Cole and McGlathery, 2011; Seymour et al., 2018; Tarquinio et al., 2019). Due to their highly intimate spatial association with the seagrass host, endophytic microbes would also be expected to play a key role on seagrass health and physiology, but the current knowledge on their functions is scarce (Seymour et al., 2018). Endophytic bacteria promote the growth within reproductive tissues (Tarquinio et al., 2021), and based on our experience in terrestrial plants, endophytes are be expected to produce defense chemicals for the host (Trivedi et al., 2020). Endophytic bacteria such as Actinobacteria, which are common in the root tissue of seagrasses including *Z. marina* (Jensen et al., 2007), are talented producers of potent antibiotics (Dalisay et al., 2013). Notably, the ecological roles and interactions of seagrass microbiome with the surrounding seawater microbiome, including the pathogens therein remain severely understudied. Webb et al. (2019) have suggested the involvement of some microbial taxa (e.g., the genus *Tenacibaculum* identified in greater abundance in seawater samples from eelgrass-vegetated sites) in the sanitary effect of seagrasses. Interestingly, the surface of *Z. marina* blades has been found to contain high densities of (unidentified) bacteria that inhibit toxic dinoflagellates and algae that cause harmful algal blooms (HABs; Imai et al., 2009; Onishi et al., 2014; Inaba et al., 2017), an effect also shown for the extracts of *Z. marina* and *Z. noltii* (Laabir et al., 2013). Several early, narrow scope studies on the extracts of seagrass epi- and endophytic bacteria point out their antifouling and ecologically relevant activities e.g., towards fish pathogens (Marhaeni et al., 2010; Ravikumar et al., 2012). We recently reported the quorum quenching and antimicrobial activities of a small set of fungal strains isolated from the phyllosphere and the rhizosphere of *Z. marina* (Petersen et al., 2019). These encouraging studies, together with our results provide some evidence that seagrass epiphytes and endophytes, collectively the eelgrass symbionts, do not only protect their plant host, but are likely to be involved in the removal of pathogens in the surrounding seawater. The common antifouling activity of both endobionts and epibionts suggest the exudation of their chemical constituents onto leaf surface and then into seawater. Also, the seagrass host deploys its own chemical weapons onto surface (Papazian et al., 2019). Hence it is plausible that the ‘pathogen filtering/removal’ effect of seagrasses is a concerted effort of both seagrass host and its microbiome, i.e., seagrass holobiont.

Seagrass ecosystem service of filtering pathogens along the water reservoir implies decrease in exposure to pathogenic agents and global infection/disease outbreak risks, hence is a major benefit to health of ocean and human. This is particularly important within the context of climate change that may make grave impact on the fitness, virulence and dominance of certain pathogens in our coasts. A natural ‘sanitary’ service offers additional advantages, e.g., provision of sustainable fish and bivalve aquaculture that is cleaner and safer for coastal ecosystems and human consumers (de los Santos et al., 2020). Improvement of water quality by seagrasses may also assist maintenance of tourism services in coastal regions, and improve the health and life quality of low-income communities living in the coastal areas without sanitary systems (Lamb et al., 2017; Jamison et al., 2018). On the long term, it could make financial impact by decreasing the costs of global health care systems due to lower incidence of e.g., gastrointestinal and other infections (Ascioti et al., 2022). Alone in the United States, several virulent *Vibrio* spp. are cumulatively responsible for annual health costs over USD 250 M (Trevathan-Tackett et al., 2019).

Although seagrass meadows are declining worldwide, recent efforts for conservation and restoration of natural seagrass ecosystems are encouraging. Seagrass beds are widely used as indicators (and for monitoring) of the health status of coastal ecosystems (Sun et al., 2020; Soto, 2022). Recent investigations voice the need for a microbiome-driven perspective and strategies for conservation and protection of seagrass meadows (Trevathan-Tackett et al., 2019; Sun et al., 2020). Preserving healthy microbial assemblages through restoration efforts could create ecosystems that can control HABs and pathogen exposure globally. The current study paves the way for detailed studies to illuminate the multi-faceted roles of the seagrass microbiome in maintaining a healthy status of the seagrass meadows and coastal ecosystems. Further studies will not only improve our understanding in tripartite host–pathogen–microbiome interactions, but may also assist to predict the outcomes of such interactions across variable marine environments, especially under future climate change scenarios. Finally, eelgrass microbiome can deliver novel molecules for development of anti-infectives for environmental or human use.

## Credit authorship contribution statement

**DT:** Conceptualization, methodology, formal analysis, investigation, writing-original draft preparation, review & editing, supervision.

**SS:** Investigation, methodology, formal analysis, writing-original draft preparation.

**CUT**: Methodology, formal analysis, writing-original draft preparation.

**TJ**: Investigation, methodology. **MB**: Methodology, formal analysis. **CW**: Methodology, formal analysis **AWS**: Methodology, formal analysis.

**VAE**: Methodology, formal analysis.

## Data availability

Research data may be available upon request to corresponding author.

## Declaration of competing interest

The authors declare that they have no known competing financial interests or personal relationships that could have appeared to influence the work reported in this paper.

## Acknowledgments

We thank the Institute of Clinical Molecular Biology in Kiel for providing Sanger sequencing as supported in part by the DFG Clusters of Excellence “Precision Medicine in Chronic Inflammation” and “ROOTS”. We thank Dr. D. Langfeldt, Manuela Pendziwiat and Dr. B. Löscher for technical support.

## Notes

### Competing Interest Statement

The authors have declared no competing interest.

