## Supplementary Information for "Epiphytic and endophytic microbiome of the seagrass *Zostera marina*: Do they contribute to pathogen reduction in seawater?"

### List of Supplementary Tables

**Supplementary Table S1.** Test strains used for assessment of the antimicrobial activity of *Zostera marina* microbial extracts.

**Supplementary Table S2.** Taxonomical identification of bacteria isolated from *Zostera marina* and reference samples.

**Supplementary Table S3.** Taxonomical identification of fungi isolated from *Zostera marina* and reference samples.

**Supplementary Table S4.** Bioactivity of bacterial GYM medium extracts against aquatic panel.

**Supplementary Table S5.** Bioactivity of bacterial GYM medium extracts against fecal, human and plant pathogenic panel.

**Supplementary Table S6.** Bioactivity of bacterial MA medium extracts against aquatic panel.

**Supplementary Table S7.** Bioactivity of bacterial MA medium extracts against fecal, human and plant pathogenic panel.

**Supplementary Table S8.** Bioactivity of fungal M34 medium extracts against aquatic panel.

**Supplementary Table S9.** Bioactivity of fungal M34 medium extracts against fecal, human and plant pathogenic panel.

**Supplementary Table S10.** Bioactivity of fungal PDA medium extracts against aquatic panel.

**Supplementary Table S11.** Bioactivity of fungal PDA medium extracts against fecal, human and plant pathogenic panel.

**Supplementary Table S12.** The most active ($IC_{50} \leq 10$ μg/ml) bacterial extracts selected for metabolome analysis.

**Supplementary Table S13.** The most active ($IC_{50} \leq 10$ μg/ml) fungal extracts selected for metabolome analysis.

**Supplementary Table S14.** Putative annotation of metabolites produced by the extracts of the most active bacteria isolated from eelgrass surfaces.

**Supplementary Table S15.** Putative annotation of metabolites produced by the extracts of the most active bacteria isolated from the inner tissues of eelgrass.

**Supplementary Table S16.** Putative annotation of metabolites produced by the extracts of the most active fungi isolated from eelgrass surfaces.

**Supplementary Table S17.** Putative annotation of metabolites produced by two media extracts of *Acrostalagmus luteoalbus* 720 isolated from the inner tissues of the decaying eelgrass leaves (DLI).

**List of Supplementary Figures**

**Supplementary Fig. S1:** FBMN showing the relative abundance in MA and GYM media of ions detected in the most active bacterial isolates from the surface of *Zostera marina*, i.e., healthy leaves (HLS), decaying leaves (DLS), roots (RS), plus the sediment (S) and seawater (W) references.

**Supplementary Fig. S2:** FBMN showing the relative abundance in MA and GYM media of ions detected in the most active bacterial isolates from the inner tissues of *Zostera marina*, i.e., healthy leaves (HLI), decaying leaves (DLI), roots (RI).

**Supplementary Fig. S3:** FBMN showing the relative abundance of ions detected in M34 and PDA media extracts of the most active fungi isolated from all eelgras surfaces, i.e., healthy leaves (HLS), decaying leaves (DLS), roots (RS), plus and the sediment (S) and seawater (W) references.

**Supplementary Fig. S4:** Comparative metabolomics of five *Streptomyces* species cultured in two culture media (MA and GYM).

**Supplementary Fig. S5:** Chemical structures of annotated compounds.

**Supplementary Table S1.** Test strains used for assessment of the antimicrobial activity of *Zostera marina* microbial extracts. B: Bacterium, F: Fungus, O: Oomycete.

| Test organism | Type | Abbrev. | Panel | Host organism/occurence | Disease caused (in the host, and the human) |
|---|---|---|---|---|---|
| *Algicola bacteriolytica* | B | Ab | Aquatic | Seaweed | Red spot disease in seaweed *Laminaria japonica* |
| *Lactococcus garviae* | B | Lg | Aquatic | Fish | Lactococcosis in salt-/freshwater (fish), endocarditis & other infections (human) |
| *Leifsonia aquatica* | B | La | Aquatic | Aquatic environment | Bloodstream infections (septicemia, bacteremia), peritoneal dialysis peritonitis, meningitis (all human) |
| *Pseudoalteromonas elyakovii* | B | Pe | Aquatic | Seaweed | Spot disease in seaweeds, e.g., *Laminaria japonica* |
| *Shewanella algae* | B | Sha | Aquatic | Fish, shellfish, seawater | Ulcer disease (fish); bacteremia, ear, eye, skin & other infections, endocarditis (human) |
| *Vibrio aestuarianus* | B | Vae | Aquatic | Fish, shellfish, seawater, sediment | Pacific oyster infection |
| *Vibrio alginolyticus* | B | Val | Aquatic | Fish, shellfish | Vibriosis (i.e., pathogenic hemorrhagic sepsis, fish); eye, ear, wound infections, septicemia (human) |
| *Vibrio anguillarum* | B | Va | Aquatic | Marine/freshwater fish crustaceans, molluscs, aquatic environment | Vibriosis, fatal hemorrhagic septicemia (fish) |
| *Vibrio cholerae* | B | Vch | Aquatic | fish, shellfish, seawater | Colonizes the intestine and destroys cells (fish), Cholera, gastroenteritis (human) |
| *Vibrio coralliilyticus* | B | Vco | Aquatic | Coral, fish, shellfish | Coral bleaching, larval oyster mortality |
| *Vibrio (Aliivibrio) fischeri* | B | Vf | Aquatic | Squid, fish, shrimps, free living in seawater, sediment | Liver haemorrhagia in flatfish brill *Scophthalmus rhombus* |
| *Vibrio harveyi* | B | Vh | Aquatic | Marine invertebrates and vertebrates | Vasculitis, gastroenteritis, eye lesions, skin ulcers (fish), luminous vibriosis (prawns); acute septicemia, gastroenteritis, necrotizing soft-tissue infection, high lethality through ingestion of *V. harveyi* contaminated seafood (human) |
| *Vibrio ichthyoenteri* | B | Vi | Aquatic | Fish | Bacterial enteritis, opaque intestines, intestinal necrosis in flatfish |
| *Vibrio parahaemolyticus* | B | Vp | Aquatic | Fish, shellfish, crustaceans | Gastrointestinal ailments, tailrot disease (fish), gastroenteritis, septisemia, wound infections, death (human) |
| *Vibrio splendidus* | B | Vsp | Aquatic | Fish, shellfish, sea urchins | Vibriosis in fish, also in aquaculture |
| *Vibrio vulnificus* | B | Vv | Aquatic | Fish, shellfish, seawater | Vibriosis, skin lesions, ulcers, abdominal effusion, enteritis, tissue & organ lesions (fish); Gastroenteritis, wound infection, septicemia, death (human) |
| *Enterococcus casseliflavus* | B | Ecas | Fecal | Human | Commensal, opportunistic infections esp. in the immunocomprised, urinary tract infections (UTI) and abdominal infections, bacteremia |
| *Enterococcus faecalis* | B | Ef | Fecal | Human | Endocarditis, UTI, meningitis, sepsis, hospital/catheter infections |
| *Enterococcus faecium* | B | Efm | Fecal | Human | Bloodstream, UTI and wound infections, catheter infections |
| *Enteroccous hirae* | B | Eh | Fecal | Human | Endocarditis, sepsis, UTI/bile infections, common infection in many animals |
| *Escherichia coli* | B | Ec | Fecal | Human | Diarrhea, dysentery, cystitis, pneumonia, bacteremia, peritonitis and others |

| | | | | | |
|---|---|---|---|---|---|
| Methicillin-resistant *Staphylococcus aureus* | B | MRSA | Human | Human | Infections of skin, bloodstream, lungs, hospital, medical devices and other infections |
| *Erwinia amylovora* | B | Ea | Phytopathogen | Rosaceous plants | Fireblight disease in apples, pears |
| *Pseudomonas syringae* pv. *aptata* | B | Pss | Phytopathogen | Sugar beet, melon | Leaf spot disease (sugar beet) |
| *Ralstonia solanacearum* | B | Rs | Phytopathogen | Solanaceous & other plants | Bacterial wilt in potato, tomato, tobacco, eggplants; ginger, olive tree, rose |
| *Xanthomonas campestris* | B | Xc | Phytopathogen | Cruciferous plants | Black rot disease (crucifers) |
| *Phytophthora infestans* | O | Pi | Phytopathogen | Solanaceous plants | Late blight/potato blight disease in potato, tomato |
| *Magnaphorte grisea* | F | Po | Phytopathogen | Rice plants, palm grass | Rice blast disease |

**Supplementary Table S2:** Taxonomical identification of bacteria isolated from *Z. marina* and reference samples. Strains are given with their isolation medium and source as well as Genbank accession number (Acc. no.). Closest three related strains are given according to a full BLAST (Altschul et al., 1990) search (BLAST all) and a BLAST search limited to sequences from type material (BLAST type). Code for isolation source: HLS: Healthy leaf surface, HLI: Healthy leaf inner tissue, DLS: Decaying leaf surface, DLI: Decaying leaf inner tissue, RS: Root surface, RI: Root inner tissue, W: Seawater reference, S: Sediment reference.

| Isolate no. | Medium | Source | Closest related species (BLAST all) | Acc. no. next related (all) | Closest related species (BLAST type) | Acc. no. next related (type) | Lowest taxonomic classification (order) | Acc. no. |
|---|---|---|---|---|---|---|---|---|
| 111 | MMN | HLS | Pedobacter silvilitoris strain W-WS1<br>Unidentified bacterium isolate R-20818<br>Pedobacter sp. RSR28 | NR_134787.1<br>AJ786785.1<br>KP876467.1 | Pedobacter silvilitoris strain W-WS1<br>Pedobacter alpinus strain RSP19<br>Pedobacter lignilitoris strain W-WS13 | NR_134787.1<br>NR_144597.1<br>NR_137365.1 | *Pedobacter silvitoris* (Sphingobacteriales) | OR400255 |
| 113 | MMN | HLS | Brevundimonas sp. strain FW305-C-20-1<br>Brevundimonas sp. 'scallop'<br>Brevundimonas vesicularis strain JM91 | MT160402.1<br>CP039382.1<br>MN758851.1 | Brevundimonas vesicularis strain NBRC 12165<br>Brevundimonas vesicularis type strain DSM 7226T<br>Brevundimonas nasdae strain W1-2B | OL880545.1<br>LN681560.1<br>NR_028633.1 | *Brevundimonas* sp. (Caulobacterales) | OR400256 |
| 115 | MMN | RS | Uncultured bacterium clone nck189a07c1<br>Sphingomonadaceae bacterium DVSMS<br>Novosphingobium sp. Rr 2-17 | KF094233.1<br>KJ004493.1<br>EU984513.1 | Novosphingobium clariflavum strain CICC11035s<br>Novosphingobium clariflavum strain CICC<br>Novosphingobium soli strain CC-TPE-1 | NR_157981.1<br>MT760066.1<br>NR_116654.1 | *Novosphingobium* sp. (Sphingomonadales) | OR400257 |
| 117 | MMN | HLS | Kocuria rhizophila strain RW13<br>Kocuria rhizophila strain D2<br>Kocuria sp. strain BAB-2791 | MH715224.1<br>MH005095.1<br>MF188117.1 | Kocuria rhizophila strain DSM 11926<br>Kocuria arsenatis strain CM1E1<br>Kocuria rhizophila strain TA68 | KJ476722.1<br>NR_148610.1<br>NR_026452.1 | *Kocuria* sp. (Micrococcales) | OR400258 |
| 122 | MMN | HLS | Microbacterium saccharophilum strain CCBhMc-4<br>Microbacterium kitamiense strain SW-138<br>Microbacterium sp. SL014B-14A2 | MK239627.1<br>KY382822.1<br>KP711500.1 | Microbacterium laevaniformans strain DSM 20140<br>Microbacterium saccharophilum strain K-1<br>Microbacterium kitamiense strain kitami C2 | MN543870.1<br>NR_114342.1<br>NR_112042.2 | *Microbacterium* sp. (Micrococcales) | OR400259 |
| 131 | MMN | HLS | Streptomyces pratensis strain JK-A-21<br>Streptomyces sp. strain FW305-F13<br>Streptomyces sp. EO1-12 | OM909122.1<br>OM867411.1<br>LC685868.1 | Streptomyces anulatus strain NRRL B-2000<br>Streptomyces griseus subsp. griseus strain DSM 40236<br>Streptomyces pluricolorescens strain NBRC 12808 (T) | MT569979.1<br>MK734067.1<br>MK424312.1 | *Streptomyces* sp. (Streptomycetales) | OR400260 |
| 137 | MMN | RS | Rhizobium sp. strain ICMP 22292<br>Rhizobium sp. strain ICMP 22260<br>Agrobacterium sp. BE516 | MH392633.1<br>MH392684.1<br>JQ764998.1 | Rhizobium taibaishanense CCNWSX 0483<br>Agrobacterium vitis strain NBRC 15140<br>Agrobacterium vitis strain K309 | NR_117899.1<br>NR_113735.1<br>NR_036780.1 | *Rhizobium* sp. (Hyphomicrobiales) | OR400261 |
| 144 | MMN | DLI | Pseudoalteromonas sp. BSs20004<br>Pseudoalteromonas sp. MM1<br>Pseudoalteromonas spiralis strain LAMA 0631 | EU365506.1<br>LC767762.1<br>OQ442358.1 | Pseudoalteromonas issachenkonii strain KMM 3549<br>Pseudoalteromonas tetraodonis strain GFC<br>Pseudoalteromonas issachenkonii strain KCTC 12958 | CP011030.1<br>CP011041.1<br>CP013350.1 | *Pseudoalteromonas* sp. (Alteromonadales) | OR400262 |
| 146 | MMN | S | Micromonospora sp. strain Atrumurd12<br>Micromonospora sp. strain BSP4<br>Micromonospora sp. strain SL97 | MK894186.1<br>MT176511.1<br>MN822727.1 | Micromonospora azadirachtae AZ1-19<br>Micromonospora siamensis strain DSM 45097<br>Micromonospora chokoriensis strain DSM 45160 | LC224297.1<br>LT607751.1<br>LT607409.1 | *Micromonospora* sp. (Micromonosporales) | OR400263 |
| 150 | MMN | HLS | Uncultured bacterium clone A_0706<br>Uncultured bacterium clone A_0706_2<br>Stenotrophomonas maltophilia strain KNUC311 | GQ379598.1<br>GQ379588.1<br>EU239136.1 | Stenotrophomonas nematodicola culture CPCC:101271<br>Stenotrophomonas rhizophila strain e-p10<br>Stenotrophomonas rhizophila strain DSM14405 | MT126327.1<br>NR_121739.1<br>CP007597.1 | *Stenotrophomonas* sp. (Xanthomonadales) | OR400264 |

| Isolate no. | Medium | Source | Closest related species (BLAST all) | Acc. no. next related (all) | Closest related species (BLAST type) | Acc. no. next related (type) | Lowest taxonomic classification (order) | Acc. no. |
|---|---|---|---|---|---|---|---|---|
| 204 | WSP30 | HLS | Falsirhodobacter sp. alg1<br>Falsirhodobacter deserti strain WQM-13<br>Falsirhodobacter deserti strain WQM-11 | AB916500.1<br>MT974352.1<br>MT974329.1 | Falsirhodobacter halotolerans strain JA744<br>Tabrizicola oligotrophica strain KMS-5<br>Cypionkella psychrotolerans strain PAMC 27389 | NR_108884.1<br>MK215682.1<br>NR_148653.1 | *Falsirhodobacter* sp. (Rhodobacterales) | OR400266 |
| 205 | WSP30 | DLS | Salinimonas sediminis strain N102<br>Alteromonas indica strain IO390401<br>Alteromonas sp. M71_D56 | CP031769.1<br>NR_179950.1<br>FM992724.1 | Salinimonas sediminis strain N102<br>Alteromonas indica strain IO390401<br>Alteromonas indica strain IO390401 | CP031769.1<br>NR_179950.1<br>MG852173.1 | Alteromonadaceae sp. (Alteromonadales) | OR400267 |
| 218 | WSP30 | W | Curtobacterium sp. strain OVL10<br>Curtobacterium sp. strain W3<br>Curtobacterium herbarum isolate PDD-32b-46 | MN811065.1<br>MK100500.1<br>HQ256845.1 | Curtobacterium herbarum strain P 420/07<br>Curtobacterium herbarum strain DSM 14013<br>Curtobacterium allii strain 20TX0166 | NR_025461.1<br>NR_115036.1<br>OK275102.1 | *Curtobacterium herbarum* (Micrococcales) | OR400269 |
| 229 | WSP30 | DLS | Novosphingobium sp. strain AX3-1<br>Novosphingobium sp. MBES04<br>Novosphingobium sp. F189 | MZ312133.1<br>AB733576.1<br>KT900241.1 | Novosphingobium decolorationis strain 502str22<br>Novosphingobium decolorationis strain 502str22<br>Novosphingobium soli strain CC-TPE-1 | MN940439.1<br>CP054856.1<br>NR_116654.1 | *Novosphingobium decolorationis* (Sphingomonadales) | OR400270 |
| 237 | WSP30 | DLI | Alpha proteobacterium MBF<br>Thalassospira sp. R-52913<br>Thalassospira sp. R-52699 | AB378715.1<br>KT185151.1<br>KT185150.1 | Thalassospira lucentensis MCCC 1A00383 = DSM 14000 strain QMT2<br>Thalassospira lohafexi strain 139Z-12<br>Thalassospira alkalitolerans strain MBE#61 | NR_115011.1<br>NR_136875.1<br>NR_114386.1 | *Thalassospira lucentensis* (Rhodospirillales) | OR400271 |
| 248 | WSP30 | HLS | Vibrio sp. strain PP-XX7<br>Vibrio sp. strain PP-XX4-1<br>Vibrio metschnikovii strain Y2 | OQ712005.1<br>OQ712004.1<br>MK999904.1 | Vibrio cincinnatiensis strain ATCC 35912<br>Vibrio sp. LJC006<br>Vibrio viridaestus strain LJC006 | NR_026122.1<br>MG745796.1<br>NR_179904.1 | *Vibrio metschnikovii* (Vibrionales) | OR400272 |
| 250 | WSP30 | HLS | Alteromonas sp. TO01<br>Alteromonas stellipolaris strain PQQ-42<br>Alteromonas stellipolaris strain R10SW13 | AB049730.1<br>CP015345.1<br>CP014322.1 | Alteromonas stellipolaris strain LMG 21861T<br>Alteromonas stellipolaris strain LMG 21861<br>Alteromonas naphthalenivorans strain SN2 | AJ295715.2<br>CP013926.1<br>CP002339.1 | *Alteromonas stellipolaris* (Alteromonadales) | OR400273 |
| 303 | TSB3+10 | HLS | Gallaecimonas sp. strain K01<br>Uncultured bacterium clone Lc2z_ML_310<br>Uncultured bacterium clone Lc2z_ML_075 | KY819126.1<br>FJ355333.1<br>FJ355332.1 | Gallaecimonas pentaromativorans strain CEE_131<br>Gallaecimonas pentaromativorans strain CEE_131<br>Gallaecimonas pentaromativorans strain CEE_131 | NR_116928.1<br>NR_116926.1<br>NR_116925.1 | *Gallaecimonas pentaromativorans* (Gammaproteobacteria incertae sedis (Gallaecimonas)) | OR400274 |
| 305 | TSB3+10 | RS | Flavobacterium frigidarium strain CBS 21<br>Flavobacterium frigidarium strain CBS 13<br>Flavobacterium frigidarium strain CBS 4 | MT539759.1<br>MT539752.1<br>MT539744.1 | Flavobacterium ovatum strain W201E<br>Flavobacterium muglaense strain F-60<br>Flavobacterium frigidarium strain NCIMB 13737 | NR_159890.1<br>MW555593.1<br>NR_117853.1 | *Flavobacterium* sp. (Flavobacteriales) | OR400275 |
| 306 | TSB3+10 | HLS | Exiguobacterium sp. strain NO6<br>Exiguobacterium artemiae strain NO5<br>Exiguobacterium sp. strain LK8P6F | MT377851.1<br>MT377850.1<br>MN491312.1 | Exiguobacterium artemiae strain 9AN<br>Exiguobacterium sibiricum 255-15<br>Exiguobacterium undae strain DSM 14481 | NR_114970.2<br>CP001022.1<br>NR_043477.1 | *Exiguobacterium* sp. (Caryophanales) | OR400276 |
| 323 | TSB3+10 | HLS | Flavobacterium sp. strain SW-2<br>Flavobacterium sp. SH55<br>Flavobacterium ponti strain GSW-R14 | MW245855.1<br>JQ778313.1<br>NR_104505.1 | Flavobacterium ponti strain GSW-R14<br>Flavobacterium gelidilacus strain R-8899<br>Flavobacterium maris strain KMM 9535 | NR_104505.1<br>NR_025538.1<br>NR_145939.1 | *Flavobacterium ponti* (Flavobacteriales) | OR400277 |
| 325 | TSB3+10 | HLS | Gallaecimonas sp. strain K02<br>Gallaecimonas sp. strain K01<br>Gallaecimonas pentaromativorans strain EP27-14-7 | KY819127.1<br>KY819126.1<br>KY457426.1 | Gallaecimonas pentaromativorans strain CEE_131<br>Gallaecimonas pentaromativorans strain CEE_131<br>Gallaecimonas pentaromativorans strain CEE_131 | NR_116927.1<br>NR_116928.1<br>NR_116925.1 | *Gallaecimonas pentaromativorans* (Gammaproteobacteria incertae sedis (Gallaecimonas)) | OR400278 |

| Isolate no. | Medium | Source | Closest related species (BLAST all) | Acc. no. next related (all) | Closest related species (BLAST type) | Acc. no. next related (type) | Lowest taxonomic classification (order) | Acc. no. |
|---|---|---|---|---|---|---|---|---|
| 334 | TSB3+10 | RI | Breoghania corrubedonensis strain UBF-P1<br>Salaquimonas pukyongi strain RR3-28<br>Salaquimonas pukyongi strain RR3-28 | NR_104495.1<br>NR_173673.1<br>CP019044.1 | Breoghania corrubedonensis strain JC706<br>Nesiotobacter exalbescens strain 0298<br>Uncultured alpha proteobacterium clone B23 | OU501485.1<br>KP236413.1<br>GU259680.1 | *Breoghania corrubedonensis*<br>(Hyphomicrobiales) | OR400280 |
| 431 | PDA | DLI | Mycolicibacterium sediminis JCM 17899<br>Mycolicibacterium sediminis strain PDD-69b-31<br>Mycolicibacterium sediminis strain YIM M13028 | AP022588.1<br>OM827359.1<br>NR_109733.1 | Mycolicibacterium sediminis strain YIM M13028<br>Mycolicibacterium arabiense JCM 18538<br>Mycolicibacterium arabiense strain YIM 121001 | NR_109733.1<br>AP022593.1<br>NR_109734.2 | *Mycolicibacterium sediminis*<br>(Mycobacteriales) | OR400281 |
| 436 | PDA | DLI | Paenibacillus terrae strain Pt05<br>Paenibacillus sp. strain 23TSA30-6<br>Paenibacillus sp. 28ISP30-2 | MN961668.1<br>MK511843.1<br>MK511842.1 | Paenibacillus maysiensis strain 1-49<br>Paenibacillus terrae strain AM141<br>Paenibacillus brasilensis strain PB1 72 | NR_165764.1<br>NR_025170.1<br>NR_025106.1 | *Paenibacillus* sp.<br>(Caryophanales) | OR400282 |
| 609 | TSB+BSW | HLS | Agarivorans sp. strain AR19-05<br>Agarivorans sp. strain S7-5-32<br>Agarivorans gilvus strain IMCC34138 | MW826714.1<br>MK737665.1<br>MG456768.1 | Agarivorans litoreus strain GJSW-6<br>Agarivorans albus JCM 21469<br>Agarivorans albus strain NBRC 102603 | NR_134691.1<br>AP023032.1<br>NR_114160.1 | *Agarivorans* sp.<br>(Alteromonadales) | OR400283 |
| 611 | TSB+BSW | HLS | Micrococcus luteus strain PGHP6<br>Micrococcus luteus strain AJ54<br>Micrococcus sp. strain BSE45 | MT539734.1<br>MT533935.1<br>MT397263.1 | Micrococcus yunnanensis strain YIM 65004<br>Micrococcus luteus strain NCTC 2665<br>Micrococcus aloeverae strain AE-6 | NR_116578.1<br>MN075406.1<br>NR_134088.1 | *Micrococcus* sp.<br>(Micrococcales) | OR400285 |
| 614 | TSB+BSW | DLS | Aquimarina muelleri strain V1SW 51<br>Aquimarina muelleri strain KCTC 12285<br>Aquimarina muelleri strain KMM 6028 | OL744421.1<br>MT759915.1<br>AY608408.1 | Aquimarina muelleri strain KCTC 12285<br>Aquimarina muelleri strain KMM 6020<br>Aquimarina muelleri strain KCTC 12285 | MT759915.1<br>NR_025823.1<br>MT758055.1 | *Aquimarina muelleri*<br>(Flavobacteriales) | OR400286 |
| 616 | TSB+BSW | DLS | Sulfitobacter sp. strain P149-L05b<br>Sulfitobacter pontiacus strain YT163<br>Sulfitobacter pontiacus strain YT143 | MN043891.1<br>MH725557.1<br>MH725556.1 | Sulfitobacter pontiacus strain ChLG-10<br>Sulfitobacter pontiacus strain ChLG-10<br>Sulfitobacter aestuarii strain hydD52 | MF417412.1<br>NR_026418.1<br>MG210570.1 | *Sulfitobacter pontiacus*<br>(Rhodobacterales) | OR400287 |
| 625 | TSB+BSW | HLS | Flavobacterium sp. strain SWA EN P2.6<br>Bacterium AK30<br>Flavobacterium sp. strain SWA CA P2.5 | OL672351.1<br>LC465495.1<br>KY476579.1 | Flavobacterium jumunjinense strain HME7102<br>Flavobacterium jumunjinense strain HME7102<br>Flavobacterium sediminilitoris strain YSM-43 | CP091285.1<br>NR_109367.1<br>NR_159337.1 | *Flavobacterium jumunjinense*<br>(Flavobacteriales) | OR400288 |
| 629 | TSB+BSW | HLS | Cobetia sp. 5-11-6-3<br>Cobetia pacifica strain GPM2<br>Cobetia sp. L2A1 | LC549335.1<br>CP047970.1<br>CP047025.1 | Cobetia marina strain JCM 21022<br>Cobetia marina strain NBRC 102605<br>Cobetia pacifica strain KMM 3879 | CP017114.1<br>NR_114162.1<br>NR_113402.1 | *Cobetia* sp.<br>(Oceanospirillales) | OR400289 |
| 630 | TSB+BSW | RS | Salinimonas sediminis<br>Alteromonas sp.<br>Alteromonas sp. | CP031769.1<br>FM992724.1<br>FM992719.1 | Salinimonas sediminis<br>Alteromonas indica<br>Salinimonas sediminis | CP031769.1<br>MG852173.1<br>MG745360.3 | Alteromonadaceae sp.<br>(Alteromonadales) | OR400290 |
| 632 | TSB+BSW | DLS | Uncultured bacterium clone SS-22G12<br>Alteromonas stellipolaris strain PQQ-44<br>Alteromonas stellipolaris strain PQQ-42 | KX177758.1<br>CP015346.1<br>CP015345.1 | Alteromonas stellipolaris strain LMG 21861<br>Alteromonas stellipolaris type strain LMG 21861T<br>Alteromonas stellipolaris strain LMG 21861 | CP013926.1<br>AJ295715.2<br>NR_025433.1 | *Alteromonas stellipolaris*<br>(Alteromonadales) | OR400291 |
| 639 | TSB+BSW | HLS | Paracoccus sp. HIJAc-3c<br>Paracoccus sp. KJF6-16<br>Paracoccus marcusii strain kbs21 | JX515659.1<br>JQ800107.1<br>AF194399.1 | Paracoccus sediminilitoris strain DSL-16<br>Paracoccus nototheniae strain 41R45<br>Paracoccus seriniphilus strain NBRC 100798 | MH491014.1<br>MH065728.1<br>NR_113939.1 | *Paracoccus* sp.<br>(Rhodobacterales) | OR400292 |
| 640 | TSB+BSW | HLS | Neptunomonas sp. BZm-1<br>Neptunomonas phycophila strain 3SM2.1<br>Neptunomonas phycophila strain 3SH2.3 | LC006855.1<br>LT717324.1<br>LT717324.1 | Neptunomonas phycophila strain SYM1<br>Neptunomonas naphthovorans strain NBRC 101991<br>Neptunomonas antarctica strain S3-22 | NR_134775.1<br>NR_114018.1<br>NR_116775.1 | *Neptunomonas phycophila*<br>(Oceanospirillales) | OR400293 |
| 641 | TSB+BSW | RI | Bacterium AK32<br>Labrenzia sp. strain s93<br>Labrenzia sp. R-66638 | LC465497.1<br>MF796605.1<br>KT185115.1 | Roseibium marinum strain mano18<br>Roseibium album strain 5OM6<br>Roseibium hamelinense JCM 10544 | NR_043040.1<br>NR_042378.1<br>LC506143.1 | *Roseibium marinum*<br>(Hyphomicrobiales) | OR400294 |

| Isolate no. | Medium | Source | Closest related species (BLAST all) | Acc. no. next related (all) | Closest related species (BLAST type) | Acc. no. next related (type) | Lowest taxonomic classification (order) | Acc. no. |
|---|---|---|---|---|---|---|---|---|
| 704 | WSP+BSW | HLI | Bacillus sp. strain HY2-13<br>Bacillus sp. strain HY2-6<br>Bacillus thuringiensis strain IMH/T-1 | OR083872.1<br>OR083868.1<br>OR077585.1 | Bacillus thuringiensis strain ATCC 10792<br>Bacillus thuringiensis strain IAM 12077<br>Bacillus fungorum | MN396730.1<br>MK377087.1<br>MG601116.1 | *Bacillus* sp.<br>(Caryophanales) | OR400295 |
| 706 | WSP+BSW | DLI | Uncultured Oceanimonas sp. clone C6A23<br>Oceanisphaera sediminis strain SK-3<br>Oceanimonas sp. strain YTUF5 | KP016678.1<br>OL839954.1<br>KY425614.1 | Oceanisphaera marina strain YM319<br>Oceanisphaera aquimarina strain S33<br>Oceanisphaera sediminis strain TW92 | NR_157015.1<br>NR_169367.1<br>NR_109063.1 | *Oceanisphaera sediminis*<br>(Aeromonadales) | OR400296 |
| 716 | WSP+BSW | HLS | Pantoea agglomerans strain KABNA4<br>Pantoea agglomerans strain KABNA3<br>Pantoea agglomerans strain KABNA2 | MT605813.1<br>MT605812.1<br>MT605811.1 | Pantoea vagans strain LMG 24199<br>Pantoea agglomerans strain DSM 3493(T)<br>Pantoea pleuroti strain JZB 2120015 | CP038853.1<br>MF289172.1<br>KJ654341.1 | *Pantoea* sp.<br>(Enterobacterales) | OR400299 |
| 725 | WSP+BSW | RS | Bacillus mycoides strain 2861<br>Bacillus sp. (in: Bacteria) strain AVC226<br>Bacillus sp. (in: Bacteria) strain AVC212 | MT586023.1<br>MW477022.1<br>MW477009.1 | Bacillus mycoides strain ATCC 6462<br>Bacillus mycoides strain DSM 2048<br>Bacillus hominis strain BML-BC059 | CP009692.1<br>CP093291.1<br>MW674729.1 | *Bacillus* sp.<br>(Caryophanales) | OR400300 |
| 729 | WSP+BSW | HLS | Microbacterium sp. R-4-H-33<br>Microbacterium sp. AB302d<br>Microbacterium sp. OS-6 | KM362873.1<br>FR821182.1<br>AJ296094.1 | Microbacterium diaminobutyricum strain RZ63<br>Microbacterium terricola strain KV-448<br>Microbacterium terregens strain JCM 1342 | NR_152648.1<br>NR_041329.1<br>MT760352.1 | *Microbacterium* sp.<br>(Micrococcales) | OR400301 |
| 733 | WSP+BSW | S | Rhodococcus sp. strain YF-9<br>Rhodococcus qingshengii strain cqsV23<br>Rhodococcus qingshengii strain cqsV7 | MT631993.1<br>MN826591.1<br>MN826576.1 | Rhodococcus erythropolis strain: JCM 3201<br>Rhodococcus erythropolis strain DSM 43066<br>Nocardia coeliaca strain DSM 44595 | AB429553.1<br>KJ476725.1<br>NR_104776.1 | *Rhodococcus* sp.<br>(Corynebacteriales) | OR400302 |
| 738 | WSP+BSW | S | Streptomyces sp.<br>Streptomyces argenteolus<br>Streptomyces sp. | MT540269.1<br>MN382155.1<br>MK053664.1 | Streptomyces silvae<br>Streptomyces flavovirens<br>Streptomyces flavogriseus | MW479423.1<br>MT760096.1<br>NR_028988.1 | *Streptomyces* sp.<br>(Streptomycetales) | OR400303 |
| 741 | WSP+BSW | HLS | Psychrobacter sp. strain PL 19<br>Psychrobacter cryohalolentis strain 4_KNS_9_Sed_A2<br>Psychrobacter cryohalolentis strain 4_KNS_9_Sed_Z3 | MT594461.1<br>MT309525.1<br>MT309524.1 | Psychrobacter cryohalolentis K5<br>Psychrobacter arcticus strain 273-4<br>Psychrobacter okhotskensis strain MD17 | NR_075055.1<br>NR_075054.1<br>MW228834.1 | *Psychrobacter* sp.<br>(Pseudomonadales) | OR400304 |
| 742 | WSP+BSW | HLI | Bacillus sp. (in: Bacteria) strain H402<br>Bacillus simplex strain EH12<br>Bacillus thuringiensis strain EGI94 | MH669310.1<br>MN750767.1<br>MN704417.1 | Peribacillus simplex strain NBRC 15720 = DSM 1321<br>Peribacillus frigoritolerans strain DSM 8801<br>Peribacillus frigoritolerans strain DSM 8801 (T) | CP017704.1<br>NR_115064.1<br>MK424281.1 | *Bacillus* sp.<br>(Caryophanales) | OR400305 |
| 744 | WSP+BSW | S | Streptomyces prasinosporus strain Ya22<br>Streptomyces scopiformis strain WLY164<br>Streptomyces rubrogriseus strain NM6 | MN315547.1<br>MT322212.1<br>MT071572.1 | Streptomyces scopiformis strain NBRC 100244<br>Streptomyces scopiformis strain A25<br>Streptomyces ambofaciens ATCC 23877 | NR_112586.1<br>NR_028764.1<br>CP012382.1 | *Streptomyces* sp.<br>(Streptomycetales) | OR400306 |
| 747 | WSP+BSW | DLI | Loktanella atrilutea isolate HaHa_7_89<br>Loktanella sp. D15_40<br>Loktanella sp. strain CP8 | LR722705.1<br>KM268056.1<br>MN326783.1 | Loktanella salsilacus strain NBRC 102486<br>Loktanella salsilacus strain LMG21507<br>Loktanella salsilacus strain R-8904 | NR_114115.1<br>NR_115894.1<br>NR_025539.1 | *Loktanella* sp.<br>(Rhodobacterales) | OR400307 |
| 202b | WSP30 | HLS | Pseudoalteromonas sp. strain C52<br>Pseudoalteromonas sp. strain C40<br>Pseudoalteromonas elyakovii strain A43 | MT645494.1<br>MT645493.1<br>MT598003.1 | Pseudoalteromonas carrageenovora IAM 12662<br>Pseudoalteromonas arctica A 37-1-2<br>Pseudoalteromonas issachenkonii strain KMM 3549 | LT965929.1<br>CP011026.1<br>CP011030.1 | *Pseudoalteromonas* sp. (Alteromonadales) | OR400265 |
| 214a | WSP30 | HLI | Bacillus megaterium strain DY14-63<br>Bacillus sp. (in: Bacteria) strain CL03-35<br>Bacillus sp. Ishi 8/5a | KC494006.1<br>MT472067.1<br>AB972819.1 | Bacillus megaterium NBRC 15308 = ATCC 14581<br>Priestia megaterium strain ATCC 14581<br>Bacillus megaterium strain DSM 32 | CP009920.1<br>CP069288.1<br>KJ476721.1 | *Bacillus megaterium*<br>(Caryophanales) | OR400268 |

| Isolate no. | Medium | Source | Closest related species (BLAST all) | Acc. no. next related (all) | Closest related species (BLAST type) | Acc. no. next related (type) | Lowest taxonomic classification (order) | Acc. no. |
|---|---|---|---|---|---|---|---|---|
| 332b | TSB3+10 | DLI | Oceanobacillus sp. strain L38<br>Oceanobacillus sp. strain ZI6<br>Oceanobacillus sp. strain 1823 | MT516462.1<br>MT433859.1<br>MT255006.1 | Oceanobacillus picturae strain R-5321<br>Oceanobacillus kapialis strain PCU 300<br>Oceanobacillus manasiensis strain YD3-56 | NR_028952.1<br>MT758147.1<br>NR_116624.1 | *Oceanobacillus* sp.<br>(Caryophanales) | OR400279 |
| 3LBT | TSB3+10 | HLI | Endophytic bacterium strain B12<br>Bacillus pumilus strain JK-B-11<br>Bacillus pumilus strain JK-B-09 | OM938278.1<br>OM909099.1<br>OM909097.1 | Peribacillus acanthi strain L28<br>Bacillus pumilus strain NCTC10337<br>Bacillus pumilus strain NBRC-12092 | MG719555.1<br>LT906438.1<br>KX261622.1 | *Bacillus pumilus*<br>(Caryophanales) | OR400253 |
| 610a | TSB+BSW | HLS | Flavobacterium sp. strain SWA CA P1.7<br>Flavobacterium sp. strain N1861<br>Flavobacterium jumunjinense strain Hal044 | OL662948.1<br>CP109996.1<br>MT406425.1 | Flavobacterium jumunjinense strain HME7102<br>Flavobacterium jumunjinense strain HME7102<br>Flavobacterium jumunjinense strain CECT 7955 | CP091285.1<br>NR_109367.1<br>MT760282.1 | *Flavobacterium jumunjinense*<br>(Flavobacteriales) | OR400284 |
| 6RBSU | TSB+BSW | RS | Microbacterium sp. strain B8637.14<br>Endophytic bacterium strain B1<br>Microbacterium foliorum strain IITA-TZ090 | OM948661.1<br>OM938267.1<br>OM909339.1 | Microbacterium algeriense strain G1<br>Microbacterium phyllosphaerae strain DSM 13468 (T)<br>Microbacterium hydrocarbonoxydans strain NBRC 103074 (T) | MK480726.1<br>MK424292.1<br>MK424288.1 | *Microbacterium* sp.<br>(Micrococcales) | OR400254 |
| 711b | WSP+BSW | RS | Flavobacterium sp. strain 2157<br>Flavobacterium sp. strain 1639<br>Flavobacterium frigidarium strain CBS 23 | MT585995.1<br>MT585977.1<br>MT539760.1 | Flavobacterium muglaense strain F-60<br>Flavobacterium frigidarium strain NCIMB 13737<br>Flavobacterium frigidarium strain A2i | MW555593.1<br>NR_117853.1<br>NR_025020.1 | *Flavobacterium* sp.<br>(Flavobacteriales) | OR400297 |
| 713b | WSP+BSW | HLS | Proteus cibi<br>Proteus cibi strain FJ2001126-3<br>Proteus vulgaris strain DSM 13387T | MG913248.1<br>NR_179961.1<br>HE978268.1 | Proteus vulgaris strain ZN3<br>Proteus cibi<br>Proteus sp. strain SCAU6 | CP047344.1<br>MG913248.1<br>KP125982.1 | *Proteus* sp.<br>(Enterobacterales) | OR400298 |
| DAI04 | MA | DLS | Uncultured Pseudoalteromonas sp. clone SSU_type_12<br>Pseudoalteromonas tetraodonis strain GFC<br>Pseudoalteromonas sp. strain H12 | MH894289.1<br>CP011041.1<br>KX982224.1 | Pseudoalteromonas tetraodonis strain GFC<br>Pseudoalteromonas tetraodonis GFC<br>Pseudoalteromonas issachenkonii strain KMM 3549 | CP011041.1<br>NR_114187.1<br>CP011030.1 | *Pseudoalteromonas tetraodonis*<br>(Alteromonadales) | OR400308 |
| RBI15 | MA | RS | Uncultured bacterium clone EvDD2-24<br>Uncultured bacterium clone EvDD2-39<br>Uncultured gamma proteobacterium clone LSmat.Fer.04 | MH556117.1<br>MH556130.1<br>HE576775.1 | Pseudoalteromonas caenipelagi strain JBTF-M23<br>Pseudoalteromonas amylolytica strain JW1<br>Pseudoalteromonas byunsanensis strain FR1199 | MT348162.1<br>NR_149821.1<br>DQ011289.2 | *Pseudoalteromonas* sp. (Alteromonadales) | OR400309 |
| x001 | MA | HLS | Cobetia sp. 5-11-6-3<br>Cobetia pacifica strain GPM2<br>Cobetia marina strain W1B | LC549335.1<br>CP047970.1<br>MN326584.1 | Cobetia marina strain JCM 21022<br>Cobetia marina strain NBRC 102605<br>Cobetia pacifica strain KMM 3879 | CP017114.1<br>NR_114162.1<br>NR_113402.1 | *Cobetia* sp.<br>(Oceanospirillales) | OR400310 |
| x005 | MA | RS | Bacterium AK37<br>Dokdonia sp. strain BMW36<br>Dokdonia sp. strain 9a | LC465515.1<br>MH773427.1<br>MT484159.1 | Dokdonia eikasta strain NBRC 100814<br>Dokdonia diaphoros strain MSKK-32<br>Dokdonia eikasta strain PMA-26 | NR_113944.1<br>NR_041274.1<br>NR_041273.1 | *Dokdonia* sp.<br>(Flavobacteriales) | OR400311 |
| x007 | MA | HLS | Bacillus sp. (in: Bacteria) MH10<br>Alkalihalobacillus hwajinpoensis strain HSB097<br>Bacillus sp. (in: Bacteria) strain RKSG232 | LC373524.1<br>MH559620.1<br>MH491170.1 | Alkalihalobacillus hwajinpoensis strain SW-72<br>Alkalihalobacillus hemicentroti strain JSM 076093<br>Alkalihalobacillus hwajinpoensis strain SW-72 | NR_025264.1<br>NR_109010.1<br>MW227498.1 | *Alkalihalobacillus hwajinpoensis*<br>(Caryophanales) | OR400312 |
| x012 | MA | RI | Shewanella basaltis strain 105<br>Shewanella sp. R-2-H-2-1<br>Shewanella sp. ER-Te-42B-Light | MK967079.1<br>KM362877.1<br>MZ905343.1 | Shewanella basaltis strain J83<br>Shewanella inventionis strain KX27<br>Shewanella xiamenensis strain JCM 16212 | NR_044418.1<br>NR_153722.1<br>MT760370.1 | *Shewanella* sp.<br>(Alteromonadales) | OR400313 |
| x014 | MA | DLI | Aquimarina sp. strain SS2-1<br>Aquimarina sp. AD1<br>Aquimarina amphilecti strain 92V | OM763662.1<br>CP031966.1<br>NR_126246.1 | Aquimarina amphilecti strain 92V<br>Aquimarina latercula strain NBRC 15938<br>Aquimarina seongsanensis strain CBA3208 | NR_126246.1<br>NR_113824.1<br>NR_152627.1 | *Aquimarina* sp.<br>(Flavobacteriales) | OR400314 |

| Isolate no. | Medium | Source | Closest related species (BLAST all) | Acc. no. next related (all) | Closest related species (BLAST type) | Acc. no. next related (type) | Lowest taxonomic classification (order) | Acc. no. |
|---|---|---|---|---|---|---|---|---|
| x015 | MA | HLI | Bacillus amyloliquefaciens<br>Bacillus sp. AM1(2019)<br>Bacillus amyloliquefaciens | MK743994.1<br>CP047644.1<br>MN759438.1 | Bacillus amyloliquefaciens<br>Bacillus amyloliquefaciens<br>Bacillus velezensis | NR_117946.1<br>FN597644.1<br>KY694464.1 | *Bacillus* sp.<br>(Caryophanales) | OR400315 |
| x018 | MA | RS | Octadecabacter sp. CAU 1310<br>Uncultured bacterium clone Woods-Hole_a5909<br>Octadecabacter sp. ZS6-8 | KU671053.1<br>KF799439.1<br>FJ196060.1 | Octadecabacter arcticus strain 238<br>Octadecabacter antarcticus strain 307<br>Octadecabacter ponticola strain HDSW-34 | NR_102905.1<br>NR_102908.1<br>NR_152087.1 | *Octadecabacter* sp.<br>(Rhodobacterales) | OR400316 |
| x019 | MA | DLI | Streptomyces sp. strain BSP1<br>Streptomyces sp. 604F<br>Streptomyces violascens strain EGI125 | MT176505.1<br>CP026490.1<br>MN704434.1 | Streptomyces argenteolus strain CGMCC 4.1693<br>Streptomyces sampsonii strain ATCC 25495<br>Streptomyces sampsonii strain NRRL B12325 | EU048540.1<br>NR_025870.2<br>NR_116508.1 | *Streptomyces* sp.<br>(Streptomycetales) | OR400317 |
| x025 | MA | DLS | Streptomyces sp. Esraa<br>Streptomyces sp. strain W54<br>Streptomyces griseorubens strain INA 5148 | KJ777675.3<br>KY402256.1<br>MG200163.1 | Streptomyces griseorubens strain NBRC 12780<br>Streptomyces lateritius strain CSSP722<br>Streptomyces sp. MS 3/20 | NR_041066.1<br>NR_115438.1<br>KR827683.1 | *Streptomyces griseorubens*<br>(Streptomycetales) | OR400318 |
| x027 | MA | HLS | Zobellia amurskyensis strain KMM 3526<br>Zobellia sp. strain SWA EN P1.8<br>Zobellia sp. TBL_82 | NR_024826.1<br>OL672360.1<br>JX854306.1 | Zobellia amurskyensis strain KMM 3526<br>Zobellia laminariae strain KMM 3676<br>Zobellia galactanivorans strain DsiJ | NR_024826.1<br>NR_024827.1<br>NR_074684.1 | *Zobellia amurskyensis*<br>(Flavobacteriales) | OR400319 |
| x028 | MA | HLS | Pseudomonas sp. strain SS NBRI 52<br>Pseudomonas mendocina strain SS NBRI 50<br>Pseudomonas mendocina strain SS NBRI 48 | MT629881.1<br>MT629879.1<br>MT629877.1 | Pseudomonas khazarica strain TBZ2<br>Pseudomonas yangonensis strain MY50<br>Pseudomonas mendocina strain NCTC10897 | NR_169334.1<br>MK907288.1<br>LR134290.1 | *Pseudomonas* sp.<br>(Pseudomonadales) | OR400320 |
| x029 | MA | RI | Rhodococcus jostii strain NW16<br>Rhodococcus koreensis strain N112<br>Rhodococcus koreensis strain N67 | JF915360.1<br>HM244985.1<br>HM244979.1 | Rhodococcus jostii strain DSM 44719 (T)<br>Rhodococcus opacus strain DSM 43205, isolate Clone 5<br>Rhodococcus jostii strain IFO 16295 | MK424304.1<br>LN827922.1<br>NR_118421.1 | *Rhodococcus* sp.<br>(Corynebacteriales) | OR400321 |
| x030 | MA | HLS | Glaciecola sp. strain SC05<br>Uncultured bacterium clone OTU191<br>Glaciecola siphonariae strain J27 | KX922678.1<br>GU451522.1<br>JX976297.1 | Paraglaciecola aquimarina strain GGW-M5<br>Paraglaciecola chathamensis S18K6<br>Paraglaciecola agarilytica strain NO2 | NR_118469.1<br>NR_041397.1<br>NR_043956.1 | *Glaciecola siphonariae*<br>(Alteromonadales) | OR400322 |
| x033 | MA | RS | Olleya marilimosa strain Y23<br>Olleya marilimosa strain SJOD-MR-31<br>Uncultured marine bacterium clone BS_OffshoreExp_T48h_A_H8 | MN746117.1<br>MK955343.1<br>KR054353.1 | Olleya marilimosa CAM030<br>Olleya marilimosa CAM030<br>Olleya algicola strain 3Alg 18 | NR_104945.2<br>NR_116100.1<br>NR_157624.1 | *Olleya marilimosa*<br>(Flavobacteriales) | OR400323 |
| x034 | MA | HLS | Uncultured bacterium clone ZJ155<br>Algoriphagus jejuensis strain CNU040<br>Uncultured bacterium clone: SC-132 | HQ625149.1<br>NR_108184.1<br>AB255108.1 | Algoriphagus jejuensis strain CNU040<br>Algoriphagus terrigena strain DS-44<br>Algoriphagus taeanensis strain HMC4223 | NR_108184.1<br>NR_043616.1<br>NR_125540.1 | *Algoriphagus jejuensis*<br>(Cytophagales) | OR400324 |
| x037 | MA | DLI | Cellulophaga pacifica strain NBRC 101531<br>Cellulophaga pacifica strain KMM 3669<br>Cellulophaga pacifica strain KMM 3664 | NR_114003.1<br>AB100841.1<br>NR_024809.1 | Cellulophaga pacifica strain NBRC 101531<br>Cellulophaga pacifica strain KMM 3664<br>Cellulophaga algicola strain DSM 14237 | NR_114003.1<br>NR_024809.1<br>NR_074452.1 | *Cellulophaga pacifica*<br>(Flavobacteriales) | OR400325 |
| x039 | MA | RI | Hoeflea alexandrii, isolate ZB_4_142<br>Mesorhizobium sp. strain L-B47<br>Hoeflea sp. strain L-B21 | LR722749.1<br>MH650981.1<br>MH650978.1 | Hoeflea alexandrii strain AM1V30<br>Hoeflea alexandrii strain CECT 5682<br>Hoeflea halophila strain JG120-1 | NR_042321.1<br>MT760263.1<br>NR_108835.1 | *Hoeflea alexandrii*<br>(Hyphomicrobiales) | OR400326 |
| xRASU | MA | RS | Rheinheimera sp. 235<br>Rheinheimera sp. 400<br>Rheinheimera sp. 347 | JQ012975.1<br>JQ012974.1<br>JQ012973.1 | Rheinheimera baltica strain OSBAC1<br>Rheinheimera japonica strain KP17<br>Rheinheimera gaetbuli strain H26 | NR_025541.1<br>NR_136858.1<br>NR_149824.1 | *Rheinheimera baltica*<br>(Chromatiales) | OR400327 |

| Isolate no. | Medium | Source | Closest related species (BLAST all) | Acc. no. next related (all) | Closest related species (BLAST type) | Acc. no. next related (type) | Lowest taxonomic classification (order) | Acc. no. |
|---|---|---|---|---|---|---|---|---|
| xSBU1 | MA | S | Pseudoalteromonas ulvae<br>Pseudoalteromonas sp.<br>Pseudoalteromonas sp. | KF472191.1<br>FR821214.1<br>AM990863.1 | Pseudoalteromonas ulvae<br>Pseudoalteromonas caenipelagi<br>Pseudoalteromonas rubra | NR_025032.1<br>MT348162.1<br>LC275063.1 | *Pseudoalteromonas ulvae* (Alteromonadales) | OR400328 |

**Supplementary Table S3:** Taxonomical identification of fungi isolated from *Z. marina* and reference samples. Strains are given with their isolation medium and source as well as Genbank accession number (Acc. no.). Closest three related strains are given according to a full BLAST (Altschul et al., 1990) search (BLAST all) and a BLAST search limited to sequences from type material (BLAST type). Code for isolation source: HLS: Healthy leaf surface, HLI: Healthy leaf inner tissue, DLS: Decaying leaf surface, DLI: Decaying leaf inner tissue, RS: Root surface, RI: Root inner tissue, W: Seawater reference, S: Sediment reference.

| Isolate no. | Medium | Source | Closest related species (BLAST all) | Acc. no. next related (all) | Closest related species (BLAST type) | Acc. no. next related (type) | Lowest taxonomic classification (order) | Acc. no. |
|---|---|---|---|---|---|---|---|---|
| 105 | MMN | DLS | Trichoderma harzianum clone HC-1<br>Trichoderma harzianum strain ZG-2-2-1<br>Trichoderma sp. 3 DoF17 | MK552405.1<br>KT192387.1<br>JQ388262.1 | Trichoderma lixii CBS 110080, TYPE material<br>Trichoderma asiaticum strain YMF1.00352<br>Trichoderma neotropicale strain CBS 130633 | NR_131264.1<br>MH113930.1<br>MH865818.1 | *Trichoderma harzianum* (Hypocreales) | OR400216 |
| 110 | MMN | W | Cladosporium cladosporioides isolate R13<br>Cladosporium sp. isolate CLAD300<br>Cladosporium sp. isolate CLAD284 | MK513830.1<br>MK111611.1<br>MK111606.1 | Cladosporium magnoliigena voucher C463<br>Cladosporium subuliforme strain CBS 126500<br>Cladosporium verrucocladosporioides strain CBS 126363 | MK347813.1<br>MH864124.1<br>MH863939.1 | *Cladosporium* sp. (Cladosporiales) | OR400217 |
| 147 | MMN | S | Arthrinium sp. strain CS06<br>Arthrinium arundinis isolate: TS08-4-1<br>Arthrinium arundinis strain MFG 70050 | KX015984.1<br>AB470870.1<br>OK563249.1 | Apiospora stipae CBS 146804 TYPE<br>Apiospora stipae strain CPC:38101<br>Apiospora chromolaenae MFLUCC 17-1505 | NR_173005.1<br>MW883403.1<br>NR_168859.1 | *Arthrinium arundinis* (Xylariales) | OR400218 |
| 210 | WSP30 | DLS | Pestalotiopsis chamaeropis isolate LB11-2<br>Pestalotiopsis chamaeropis isolate LB11-3<br>Pestalotiopsis chamaeropis isolate LB5-2 | MH425901.1<br>MH425902.1<br>MH425899.1 | Pestalotiopsis scoparia strain CBS 176.25<br>Pestalotiopsis chamaeropis strain CBS 186.71<br>Pestalotiopsis scoparia CBS 176.25 TYPE | MH854838.1<br>KM199326.1<br>NR_145238.1 | *Pestalotiopsis* sp. (Amphisphaeriales) | OR400219 |
| 233 | WSP30 | DLS | Cladosporium sp. strain 17BPLE002<br>Cladosporium sp. MV-2018B isolate MLT-27<br>Cladosporium halotolerans isolate fung15 | MT645909.1<br>MT636978.1<br>MT635287 | Cladosporium halotolerans isolate CBS119416<br>Cladosporium halotolerans ITS region; TYPE material<br>Cladosporium endophyticum MFLUCC 17-0599 TYPE | KJ596569.1<br>NR_119605.1<br>NR_158360.1 | *Cladosporium halotolerans* (Cladosporiales) | OR400220 |
| 402 | PDA | DLS | Neoascochyta exitialis strain ZMXL10<br>Neoascochyta exitialis strain ZMXL1<br>Neoascochyta graminicola isolate WAC13710 | MT446179.1<br>MT446179.1<br>MK190675.1 | Neoascochyta europaea CBS 820.84; TYPE<br>Neoascochyta dactylidis voucher MFLU:19-2859<br>Neoascochyta dactylidis MFLUCC 13-0495; TYPE | NR_136131.1<br>MT185527.1<br>NR_170041.1 | *Neoascochyta* sp. (Pleosporales) | OR400221 |
| 403 | PDA | RS | Penicillium glabrum isolate SER15_I.1<br>Penicillium glabrum strain Y56<br>Penicillium glabrum isolate P_7 | MN995823.1<br>MK793228.1<br>MK761053.1 | Penicillium glabrum strain CBS 125543<br>Penicillium glabrum CBS 125543<br>Penicillium bussumense CBS 138160 | MH863551.1<br>NR_163530.1<br>NR_137895.1 | *Penicillium glabrum* (Eurotiales) | OR400222 |
| 407 | PDA | S | Trichoderma viride strain NEFU26<br>Trichoderma viride isolate 1056F1<br>Trichoderma viride isolate T01 | KF944470.1<br>KU202215.1<br>FJ481123.1 | Hypocrea koningii strain ATCC 64262<br>Trichoderma hispanicum CBS 130540; TYPE<br>Trichoderma caribbaeum var. caribbaeum CBS; TYPE | AJ301990.1<br>NR_138451.1<br>NR_166015.1 | *Trichoderma viride* (Hypocreales) | OR400223 |
| 417 | PDA | RS | Cystofilobasidium bisporidii CBS 6346, TYPE<br>Cystofilobasidium bisporidii strain CBS 6347 | NR_154785.1<br>AF444299.1<br>KY103153.1 | Cystofilobasidium bisporidii CBS 6346; TYPE<br>Cystofilobasidium infirmominiatum CBS 323; TYPE<br>Cystofilobasidium ferigula culture CBS:7202 | NR_154785.1<br>NR_073232.1<br>KY103159.1 | *Cystofilobasidium bisporidii* (Cystofilobasidiales) | OR400224 |

| Isolate no. | Medium | Source | Closest related species (BLAST all) | Acc. no. next related (all) | Closest related species (BLAST type) | Acc. no. next related (type) | Lowest taxonomic classification (order) | Acc. no. |
|---|---|---|---|---|---|---|---|---|
| | | | Cystofilobasidium bisporidii culture CBS:6347 | | | | | |
| 420 | PDA | DLS | Starmerella vitis strain UWO.PS_00.426.2<br>Starmerella sp. CBMA19.25<br>Starmerella sp. isolate CHF-3E | MN317382.1<br>KC992848.1<br>MN861515.1 | Starmerella lactis-condensi CBS 52; TYPE<br>Starmerella riodocensis CBS 10087; TYPE<br>Starmerella neotropicalis CBS 12811; TYPE | NR_155822.1<br>NR_137870.1<br>NR_160316.1 | *Starmerella vitis*<br>(Saccharomycetales) | OR400225 |
| 434 | PDA | HLS | Parathyridaria robiniae MFLUCC 14-1119, TYPE material<br>Roussoellaceae sp. culture MUT<ITA>:2452<br>Parathyridaria robiniae voucher MFLUCC 14-1119 | NR_168161.1<br>MG813183.1<br>KY511142.1 | Parathyridaria robiniae MFLUCC 14-1119;TYPE<br>Parathyridaria clematidis isolate MFLUCC 17-2185 T<br>Parathyridaria serratifoliae isolate MFLUCC 17-2210 | NR_168161.1<br>MT310642.1<br>MT310646.1 | *Parathyridaria* sp.<br>(Pleosporales) | OR400227 |
| 444 | PDA | W | Cystofilobasidium bisporidii culture CBS:6346<br>Cystofilobasidium bisporidii culture CBS:6347<br>Cystofilobasidium bisporidii culture CBS:6348 | KY103154.1<br>KY103153.1<br>KY103151.1 | Cystofilobasidium bisporidii CBS 6346; TYPE<br>Cystofilobasidium infirmominiatum CBS 323; TYPE<br>Cystofilobasidium ferigula culture CBS:7202 | NR_154785.1<br>NR_073232.1<br>KY103159.1 | *Cystofilobasidium bisporidii*<br>(Cystofilobasidiales) | OR400228 |
| 719 | WSP+BSW | DLS | Fusarium asiaticum isolate MSBL-4<br>Fusarium asiaticum strain TK2<br>Fusarium asiaticum voucher BJ2-10 | MT322117.1<br>MK791240.1<br>KX527878.1 | Fusarium boothii NRRL 29011; TYPE material<br>Fusarium aethiopicum strain NRRL 46738<br>Fusarium boothii strain NRRL29011 | NR_121203.1<br>FJ240310.1<br>DQ459846.1 | *Fusarium* sp.<br>(Hypocreales) | OR400230 |
| 720 | WSP+BSW | DLI | Acrostalagmus luteoalbus strain CH-6<br>Acrostalagmus sp. isolate CLE7<br>Acrostalagmus luteoalbus strain TK43 | MT367202.1<br>MN543911.1<br>MH836621.1 | Sodiomyces alkalinus strain F11<br>Sodiomyces alcalophilus strain CBS 114.92<br>Sodiomyces alcalophilus JCM 7366 | OM238133.1<br>MH862344.1<br>NR_077123.1 | *Acrostalagmus luteoalbus*<br>(Glomerellales) | OR400231 |
| 737 | WSP+BSW | S | Plectosphaerella cucumerina strain PN-G-2018.6.15-#2<br>Plectosphaerella cucumerina clone 2014_1630<br>Plectosphaerella cucumerina clone 2014_1622 | MK409997.1<br>MN523210.1<br>MN523202.1 | Plectosphaerella plurivora strain Plect 365<br>Plectosphaerella niemeijerarum CBS 143233 ; TYPE<br>Plectosphaerella plurivora CBS:131742; TYPE | HQ238975.1<br>NR_156677.1<br>LR026829.1 | *Plectosphaerella* sp.<br>(Glomerellales) | OR400232 |
| 739 | WSP+BSW | S | Talaromyces barcinensis strain CBS 649.95<br>Talaromyces helicus var. helicus strain CBS 335.48<br>Talaromyces sp. voucher UNASAM-FUN-0112 | MH862547.1<br>MH856373.1<br>MT762733.1 | Talaromyces barcinensis strain CBS 649.95<br>Talaromyces helicus var. helicus strain CBS 335.48<br>Talaromyces sp. 8 AJC-2016 strain CBS 140611 | MH862547.1<br>MH856373.1<br>KU866647.1 | *Talaromyces* sp.<br>(Eurotiales) | OR400233 |
| 805 | ZMB | S | Mucor circinelloides strain CMRC 558<br>Mucor circinelloides strain CMRC 508<br>Mucor circinelloides strain CMRC 507 | MT603942.1<br>MT603901.1<br>MT603900.1 | Mucor circinelloides f. circinelloides strain CBS 195.68<br>Mucor circinelloides f. circinelloides strain CBS 195.68<br>Mucor circinelloides f. circinelloides strain CBS 195.68 | AY243943.1<br>NR_126116.1<br>HQ154604.1 | *Mucor circinelloides*<br>(Mucorales) | OR400234 |
| 809 | ZMB | S | Didymella pinodella isolate fung39<br>Penicillium olsonii strain DUCC5743<br>Penicillium coffeae strain DTO 273-A7 | MT635311.1<br>MT582783.1<br>MT309664.1 | Penicillium olsonii CBS 232.60; TYPE<br>Penicillium astrolabium NRRL 35611; TYPE<br>Penicillium salamii ITEM 15291; TYPE | NR_163546.1<br>NR_137678.1<br>NR_156542.1 | *Penicillium olsonii*<br>(Eurotiales) | OR400235 |

| Isolate no. | Medium | Source | Closest related species (BLAST all) | Acc. no. next related (all) | Closest related species (BLAST type) | Acc. no. next related (type) | Lowest taxonomic classification (order) | Acc. no. |
|---|---|---|---|---|---|---|---|---|
| 810 | ZMB | DLS | Uncultured Cladosporium clone 64<br>Uncultured Cladosporium clone 13<br>Aff. Cladosporium sp. strain B64 | MG976310.1<br>MG976260.1<br>MF615037.1 | Cladosporium iridis strain CBS 138.40<br>Cladosporium ossifragi strain CBS 842.91<br>Cladosporium sp. 5 SDM-2014 | EU167591.1<br>MH862342.1<br>LN834417.1 | *Cladosporium* sp.<br>(Cladosporiales) | OR400236 |
| 813 | ZMB | W | Aureobasidium pullulans isolate PE_11<br>Aureobasidium pullulans strain CSK3<br>Aureobasidium pullulans strain KUC3002 | MT035961.1<br>MK460802.1<br>KY294714.1 | Aureobasidium lini strain CBS 125.21<br>Aureobasidium melanogenum CBS 105.22; TYPE<br>Aureobasidium pullulans strain CBS 584.75 | MH854694.1<br>NR_159598.1<br>KT693733.1 | *Aureobasidium pullulans* (Dothideales) | OR400237 |
| 815 | ZMB | W | Sarocladium strictum isolate Zb-32<br>Sarocladium sp. isolate MBD_3441<br>Sarocladium strictum isolate VGSS15-1 | OR346314.1<br>MK595587.1<br>MF663649.1 | Sarocladium strictum isolate CBS 346.70<br>Sarocladium strictum genogroup I strain CBS 346.70T<br>Sarocladium bactrocephalum strain CBS:749.69 | NR_111145.1<br>AY138845.1<br>MH859409.1 | *Sarocladium strictum*<br>(Hypocreales) | OR400238 |
| 818 | ZMB | DLS | Paradendryphiella arenariae strain CBS 181.58<br>Dendryphiella arenaria strain CBS 181.58<br>Paradendryphiella sp. IOC-1364 | MH857747.1<br>DQ411539.1<br>AB975287.1 | Paradendryphiella arenariae strain CBS 181.58<br>Dendryphiella arenaria strain CBS 181.58<br>Stemphylium triglochinicola strain CBS 718.68 | MH857747.1<br>DQ411539.1<br>MH859210.1 | *Paradendryphiella arenariae*<br>(Pleosporales) | OR400239 |
| 901 | ZMF | S | Trichoderma sp. 3 DoF17<br>Trichoderma sp. SQR582<br>Trichoderma harzianum voucher TriH_JSB22 | JQ388262.1<br>GQ497169.1<br>KC569353.1 | Trichoderma lixii CBS 110080; TYPE<br>Trichoderma asiaticum strain YMF1.00352; HOLOTYPE<br>Trichoderma neotropicale strain CBS 130633 | NR_131264.1<br>MH113930.1<br>MH865818.1 | *Trichoderma* sp.<br>(Hypocreales) | OR400240 |
| 906 | ZMF | HLS | Penicillium expansum strain DUCC5734<br>Penicillium crustosum strain DUCC5730<br>Penicillium crustosum strain DTO 403-D6 | MT582774.1<br>MT582770.1<br>MT316358.1 | Penicillium crustosum strain CBS 115503<br>Penicillium crustosum FRR 1669<br>Penicillium fuscoglaucum isolate 2010F32 | MH862985.1<br>NR_077153.1<br>MT558936.1 | *Penicillium* sp.<br>(Eurotiales) | OR400243 |
| 907 | ZMF | DLS | Cladosporium marinum strain SFC20230103-M38<br>Cladosporium perangustum strain SFC20230103-M25<br>Cladosporium rectoides strain SFC20230103-M60 | OQ186134.1<br>OQ186121.1<br>OQ165259.1 | Cladosporium vicinum culture CPC:22316<br>Cladosporium brigadeirensis strain COAD 2257<br>Cladosporium chusqueae strain COAD 2258 | MF473311.1<br>MZ318435.1<br>MZ318430.1 | *Cladosporium* sp.<br>(Cladosporiales) | OR400244 |
| 910 | ZMF | DLS | Asteromyces cruciatus<br>Asteromyces cruciatus<br>Asteromyces cruciatus | MK432716.1<br>NR_159604.1<br>KM272370.1 | Asteromyces cruciatus<br>Stemphylium gracilariae<br>Paradendryphiella arenariae | NR_159604.1<br>MH862230.1<br>MH857747.1 | *Asteromyces cruciatus*<br>(Helotiales) | OR400245 |
| 912 | ZMF | DLS | Cladosporium allicinum isolate 9<br>Cladosporium herbarum isolate 15-027<br>Cladosporium herbarum isolate KoRLI047279 | MT573471.1<br>MK919499.1<br>MN341231.1 | Cladosporium variabile strain CBS 121636<br>Cladosporium variabile strain CBS 121635<br>Cladosporium tenellum culture CBS:121634 | MH863132.1<br>MH863131.1<br>MH863130.1 | *Cladosporium* sp.<br>(Cladosporiales) | OR400246 |
| 425a | PDA | HLS | Parathyridaria sp. voucher G.M. 2015-10-04.3<br>Thyridariaceae sp. culture MUT<ITA>:2452<br>Pleosporales sp. MUT 4893 internal transcribed spacer 1 | MW296932.1<br>MG813183.1<br>KM355998.1 | Parathyridaria robiniae MFLUCC 14-1119; TYPE<br>Parathyridaria robiniae voucher MFLUCC 14-1119<br>Parathyridaria clematidis isolate MFLUCC 17-2185 | NR_168161.1<br>KY511142.1<br>MT310642.1 | *Parathyridaria* sp.<br>(Pleosporales) | OR400226 |

| Isolate no. | Medium | Source | Closest related species (BLAST all) | Acc. no. next related (all) | Closest related species (BLAST type) | Acc. no. next related (type) | Lowest taxonomic classification (order) | Acc. no. |
|---|---|---|---|---|---|---|---|---|
| 617b | TSB+BSW | S | Trichoderma harzianum isolate MT2<br>Trichoderma harzianum isolate MT1<br>Trichoderma lixii strain F-2 | MT577837.1<br>MT577649.1<br>MT434003.1 | Trichoderma harzianum isolate MT2<br>Trichoderma harzianum isolate MT1<br>Trichoderma lixii strain F-2 | MT577837.1<br>MT577649.1<br>MT434003.1 | *Trichoderma* sp.<br>(Hypocreales) | OR400229 |
| 902b | ZMF | S | Trichoderma harzianum isolate MT2<br>Trichoderma lixii strain F-2<br>Trichoderma effusum voucher research collection Farrer lab 265 | MT577837.1<br>MT434003.1<br>MN644616.1 | Trichoderma lixii CBS 110080<br>Trichoderma asiaticum<br>Trichoderma neotropicale strain CBS 130633 | NR_131264.1<br>MH113930.1<br>MH865818.1 | *Trichoderma* sp.<br>(Hypocreales) | OR400241 |
| 903a | ZMF | RS | Alternaria infectoria<br>Lewia sp. isolate UASWS2033<br>Alternaria infectoria isolate 113 | MT635276.1<br>MN833932.1<br>MN534845.1 | Alternaria sp. 5 II 2018<br>Alternaria conjuncta strain CBS 196.86<br>Alternaria dactylidicola MFLUCC 15-0466 | LR133913.1<br>MH861940.1<br>NR_137964.1 | *Alternaria* sp.<br>(Pleosporales) | OR400242 |

**Supplementary Table S4:** Bioactivity of bacterial GYM medium extracts against aquatic panel. $IC_{50}$ values are in µg/ml. Code for isolation source: HLS: Healthy leaf surface, HLI: Healthy leaf inner tissue, DLS: Decaying leaf surface, DLI: Decaying leaf inner tissue, RS: Root surface, RI: Root inner tissue, W: Seawater reference, S: Sediment reference. La: *Leifsonia aquatica*, Lg: *Lactococcus garvieae*, Ab: *Algicola bacteriolytica*, Pe: *Pseudoalteromonas elyacovii*, Sha: *Shewanella algae*, Vae: *Vibrio aestuarianus*, Val: *Vibrio alginolyticus*, Va: *Vibrio anguillarum*, Vch: *Vibrio cholerae*, Vco: *Vibrio coralliilyticus*, Vf: *Vibrio fischeri*, Vh: *Vibrio harveyi*, Vi: *Vibrio ichthyoenteri*, Vp: *Vibrio parahaemolyticus*, Vsp: *Vibrio splendidus*, Vv: *Vibrio vulnificus*. Positive control: chloramphenicol, except for *Lactococcus garvieae* (ampicillin).

| Strain | Identification | Origin | La | Lg | Ab | Pe | Sha | Vae | Val | Va | Vch | Vco | Vf | Vh | Vi | Vp | Vsp | Vv |
|---|---|---|---|---|---|---|---|---|---|---|---|---|---|---|---|---|---|---|
| 111 | *Pedobacter silvitoris* | HLS | - | - | - | - | - | - | - | - | - | - | - | - | - | - | - | - |
| 113 | *Brevundimonas sp.* | HLS | - | - | - | - | - | - | - | - | - | - | - | - | - | - | - | - |
| 117 | *Kocuria* sp. | HLS | 94.4 | - | - | - | - | 75.3 | - | 75.9 | 28.2 | - | 60.4 | - | 48.1 | - | - | 77.6 |
| 122 | *Microbacterium* sp. | HLS | - | - | - | - | - | - | - | - | - | - | - | - | - | - | - | 79.6 |
| 131 | *Streptomyces* sp. | HLS | 1.8 | 6.9 | 1.8 | - | - | - | - | - | - | - | - | - | - | - | - | - |
| 150 | *Stenotrophomonas* sp. | HLS | 6.6 | 8.6 | - | - | - | - | - | - | - | - | - | - | - | - | - | - |
| 204 | *Falsirhodobacter* sp. | HLS | 15.5 | - | - | - | - | - | - | - | - | - | - | - | - | - | - | - |
| 248 | *Vibrio metschnikovii* | HLS | 6.5 | 15.0 | - | - | - | - | - | - | - | - | - | - | - | 27.6 | - | 56.1 |
| 250 | *Alteromonas stellipolaris* | HLS | 6.2 | 53.0 | - | - | - | - | - | - | - | - | - | - | - | - | - | - |
| 303 | *Gallaecimonas pentaromativorans* | HLS | 6.3 | 92.2 | - | - | - | - | - | - | - | - | - | - | - | - | - | - |
| 306 | *Exiguobacterium* sp. | HLS | - | - | - | - | - | - | - | - | - | - | - | - | - | - | - | - |
| 323 | *Flavobacterium ponti* | HLS | 17.7 | - | - | - | - | - | - | - | - | - | - | - | - | - | - | - |
| 325 | *Gallaecimonas pentaromativorans* | HLS | 6.8 | 78.2 | - | - | - | - | - | - | - | - | - | - | - | - | - | - |
| 609 | *Agarivorans* sp. | HLS | 90.2 | - | - | - | - | - | - | - | - | - | - | - | - | - | 60.3 | - |
| 611 | *Micrococcus* sp. | HLS | - | 97.8 | - | - | - | - | - | - | - | - | - | - | 63.0 | - | - | - |
| 625 | *Flavobacterium jumunjinense* | HLS | 6.2 | 25.8 | - | - | - | - | - | - | - | - | - | - | - | - | - | - |
| 629 | *Cobetia* sp. | HLS | 7.0 | 84.6 | - | - | - | - | - | - | - | - | - | - | - | - | - | - |
| 639 | *Paracoccus* sp. | HLS | - | - | - | - | - | - | - | - | - | - | - | - | - | - | 85.1 | - |
| 640 | *Neptunomoas phycophila* | HLS | 15.0 | 27.6 | - | - | - | - | - | - | - | - | - | - | - | - | - | - |
| 716 | *Pantoea* sp. | HLS | 6.5 | 93.4 | - | - | - | - | - | - | - | - | - | - | - | - | - | - |
| 729 | *Microbacterium* sp. | HLS | - | - | - | - | - | 80.4 | - | - | - | 72.8 | - | - | - | - | 73.7 | - |
| 741 | *Psychrobacter* sp. | HLS | 7.0 | - | - | - | - | - | - | - | - | - | - | - | - | - | - | - |
| 202b | *Pseudoalteromonas* sp. | HLS | 6.2 | 25.4 | - | - | - | - | - | - | - | - | - | - | - | - | - | - |
| 610a | *Flavobacterium jumunjinense* | HLS | 17.2 | - | - | - | - | - | - | - | - | - | - | - | - | - | - | - |
| 713b | *Proteus* sp. | HLS | 7.5 | - | - | - | - | - | - | - | - | - | - | - | - | - | - | - |
| x001 | *Cobetia* sp. | HLS | 5.9 | 21.0 | 16.7 | - | - | - | - | - | - | - | - | - | 14.0 | - | - | - |
| x007 | *Alkalihalobacillus hwajinpoensis* | HLS | 5.5 | 19.6 | - | - | - | - | - | - | - | - | - | - | - | - | - | - |
| x027 | *Zobellia amurskyensis* | HLS | - | - | - | - | - | - | - | - | - | - | - | - | - | - | - | - |
| x028 | *Pseudomonas* sp. | HLS | 6.6 | - | - | - | - | - | - | - | - | - | - | - | - | - | - | - |
| x030 | *Glaciecola siphonariae* | HLS | - | - | - | - | - | - | - | - | - | - | - | - | - | - | 92.8 | - |
| x034 | *Algoriphagus jejuensis* | HLS | - | - | - | - | - | - | - | - | - | - | - | - | - | - | - | - |
| 205 | Alteromonadaceae sp. | DLS | 6.2 | 70.7 | - | - | - | - | - | - | - | - | - | - | 29.5 | - | - | - |
| 229 | *Novosphingobium decolorationis* | DLS | 40.6 | - | - | - | - | - | - | - | - | - | - | - | 19.1 | - | - | - |
| 614 | *Aquimarina muelleri* | DLS | 11.7 | - | - | - | - | - | - | - | - | - | - | - | 46.6 | - | - | - |
| 616 | *Sulfitobacter pontiacus* | DLS | 68.0 | - | - | - | - | - | - | - | - | - | - | - | - | - | - | - |
| 632 | *Alteromonas stellipolaris* | DLS | 8.3 | 75.9 | - | - | - | - | - | - | - | - | - | - | 65.4 | - | - | - |
| DAI04 | *Pseudoalteromonas tetraodonis* | DLS | 9.1 | 21.1 | - | - | - | - | - | - | - | - | - | - | - | - | - | - |
| x025 | *Streptomyces griseorubens* | DLS | 11.0 | 76.1 | 9.4 | - | - | - | - | - | 52.4 | 97.3 | 68.7 | - | 39.5 | 28.4 | - | 18.8 |
| 115 | *Novosphingobium* sp. | RS | - | - | - | - | - | - | - | - | - | - | - | - | - | - | - | - |

| Strain | Identification | Origin | La | Lg | Ab | Pe | Sha | Vae | Val | Va | Vch | Vco | Vf | Vh | Vi | Vp | Vsp | Vv |
|---|---|---|---|---|---|---|---|---|---|---|---|---|---|---|---|---|---|---|
| 137 | *Rhizobium* sp. | RS | 7.0 | - | - | - | - | - | - | - | - | - | - | - | - | - | - | - |
| 305 | *Flavobacterium* sp. | RS | - | - | - | - | - | - | - | - | - | - | - | - | - | - | - | - |
| 630 | Alteromonadaceae sp. | RS | 6.2 | 34.7 | - | - | - | - | - | - | - | - | - | - | 53.1 | - | - | - |
| 725 | *Bacillus* sp. | RS | 6.2 | 21.1 | - | - | - | - | - | - | - | - | - | - | - | - | - | - |
| 6RBSU | *Microbacterium* sp. | RS | - | - | - | - | - | - | - | - | - | - | - | - | - | - | - | - |
| 711b | *Flavobacterium* sp. | RS | 74.8 | - | - | - | - | - | - | - | - | - | - | - | 63.9 | - | - | - |
| RBI15 | *Pseudoalteromonas* sp. | RS | 4.4 | 12.2 | - | - | - | - | - | - | - | - | 46.8 | - | - | - | - | - |
| x005 | *Dokdonia* sp. | RS | - | - | - | - | - | - | - | - | - | - | - | - | - | - | - | - |
| x018 | *Octadecabacter* sp. | RS | - | - | - | - | - | - | - | - | - | - | - | - | - | - | - | - |
| x033 | *Olleya marilimosa* | RS | 12.2 | - | - | - | - | - | - | - | - | - | - | - | - | - | - | - |
| xRASU | *Rheinheimera baltica* | RS | 7.6 | 21.7 | - | - | - | - | - | - | - | - | - | - | - | - | - | - |
| 704 | *Bacillus* sp. | HLI | 6.9 | 34.0 | - | - | - | - | - | - | - | - | - | - | - | - | - | - |
| 742 | *Bacillus* sp. | HLI | 6.3 | 9.0 | - | - | - | - | - | - | - | - | - | - | - | - | - | - |
| 214a | *Bacillus megaterium* | HLI | - | - | - | - | - | - | - | - | - | - | - | - | - | - | - | - |
| 3LBT | *Bacillus pumilus* | HLI | 31.8 | - | - | - | - | - | - | - | - | - | - | - | - | - | - | - |
| x015 | *Bacillus* sp. | HLI | 7.5 | - | - | 76.7 | - | - | - | - | - | - | - | - | 2.1 | - | - | - |
| 144 | *Pseudoalteromonas* sp. | DLI | - | - | - | - | - | - | - | - | - | - | - | - | - | - | - | - |
| 237 | *Thalassospira lucentensis* | DLI | 7.1 | - | - | - | - | - | - | - | - | - | - | - | - | - | - | - |
| 431 | *Mycolicibacterium sediminis* | DLI | - | - | - | - | - | - | - | - | - | - | - | - | - | - | - | - |
| 436 | *Paenibacillus sp.* | DLI | 7.2 | 7.4 | - | - | - | - | - | - | - | - | - | - | - | - | - | - |
| 706 | *Oceanisphaera sediminis* | DLI | 6.5 | - | - | - | - | - | - | - | - | 94.0 | - | - | - | - | 45.7 | - |
| 747 | *Loktanella* sp. | DLI | - | - | - | - | - | - | - | - | - | - | - | - | - | - | - | - |
| 332b | *Oceanobacillus* sp. | DLI | 8.6 | 24.0 | - | - | - | - | - | - | - | - | - | - | - | - | - | - |
| x014 | *Aquimarina* sp. | DLI | 11.6 | - | - | - | - | - | - | - | - | - | - | - | - | - | - | - |
| x019 | *Streptomyces* sp. | DLI | 3.1 | 20.9 | - | 11.4 | 6.6 | 7.9 | - | 15.5 | 14.0 | 7.4 | 1.9 | 51.8 | 2.1 | 2.2 | - | 1.5 |
| x037 | *Cellulophaga pacifica* | DLI | - | - | - | - | - | - | - | - | - | - | - | - | 62.7 | - | - | - |
| 334 | *Breoghania corrubedonensis* | RI | 9.4 | - | - | - | - | - | - | - | - | - | - | - | - | - | - | - |
| 641 | *Roseibium marinum* | RI | 46.5 | - | - | - | - | 83.0 | - | - | - | - | - | - | 29.0 | - | 19.3 | - |
| x012 | *Shewanella* sp. | RI | 7.1 | 24.7 | - | - | - | - | - | - | - | - | - | - | - | - | - | - |
| x029 | *Rhodococcus sp.* | RI | - | - | - | - | - | - | - | - | - | - | - | - | - | - | - | - |
| x039 | *Hoeflea alexandrii* | RI | 7.0 | - | - | - | - | - | - | - | - | - | - | - | - | - | - | - |
| 218 | *Curtobacterium herbarum* | W | - | - | - | - | 23.0 | - | - | - | - | - | - | - | 65.3 | - | - | - |
| 146 | *Micromonospora* sp. | S | - | - | - | - | - | - | - | - | - | - | - | - | - | - | - | - |
| 733 | *Rhodococcus sp.* | S | - | - | - | - | - | - | - | - | - | - | - | - | - | - | - | - |
| 738 | *Streptomyces* sp. | S | 6.6 | 21.4 | - | - | - | - | - | - | - | - | - | - | 4.6 | - | - | - |
| 744 | *Streptomyces* sp. | S | 11.2 | 76.2 | - | - | - | - | - | - | - | - | - | - | - | - | - | - |
| xSBU1 | *Pseudoalteromonas ulvae* | S | 5.7 | 21.5 | - | - | - | - | - | - | - | - | - | - | - | - | - | - |
| Pos | Positive control | - | 0.4 | 0.5 | 2.9 | 2.3 | 4.1 | 1.2 | 0.8 | 0.4 | 0.5 | 0.7 | 0.6 | 0.7 | 0.4 | 0.4 | 0.5 | 0.2 |

**Supplementary Table S5:** Bioactivity of bacterial GYM medium extracts against fecal, human and plant pathogenic panel. $IC_{50}$ values are in µg/ml.HLS: Healthy leaf surface, HLI: Healthy leaf inner tissue, DLS: Decaying leaf surface, DLI: Decaying leaf inner tissue, RS: Root surface, RI: Root inner tissue, W: Seawater reference, S: Sediment reference. Fecal pathogens (positive control): Ecas*: Enterococcus casseliflavus* (ampicillin), Ef: *Enterococcus faecalis* (ampicillin), Efm: *Enterococcus faecium* (ampicillin), Eh: *Enterococcus hirae* (ampicillin), Ec: *Escherichia coli* (chloramphenicol). Human pathogens (positive control) MRSA: Methicillin resistant *Staphylococcus aureus* (chloramphenicol). Phytopathogens (positive control) Ea: *Erwinia amylovora* (chloramphenicol), Pi: *Phytophthora infestans* (cycloheximid), Mg: *Magnaphorte grisea* (nystatin), Pss: *Pseudomonas syringae* (chloramphenicol), Rs: *Ralstonia solanacearum* (tetracycline), Xc: *Xanthomonas campestris* (chloramphenicol).

| Strain | Identification | Source | Ecas | Ef | Efm | Eh | Ec | MRSA | Ea | Pi | Mg | Pss | Rs | Xc |
|---|---|---|---|---|---|---|---|---|---|---|---|---|---|---|
| 111 | *Pedobacter silvitoris* | HLS | - | - | - | - | - | - | - | - | - | - | - | - |
| 113 | *Brevundimonas sp.* | HLS | - | - | - | - | - | 62.0 | - | - | - | - | - | - |
| 117 | *Kocuria* sp. | HLS | - | - | - | - | - | 55.6 | - | 50.1 | 59.9 | - | - | - |
| 122 | *Microbacterium* sp. | HLS | 53.1 | - | - | - | - | 41.6 | - | 31.1 | - | - | - | - |
| 131 | *Streptomyces* sp. | HLS | 1.6 | 2.0 | 2.3 | 6.4 | - | 2.3 | - | 6.7 | 1.2 | - | - | - |
| 150 | *Stenotrophomonas* sp. | HLS | 4.6 | 8.1 | 6.8 | 12.2 | - | 7.2 | - | - | - | - | - | - |
| 204 | *Falsirhodobacter* sp. | HLS | - | - | - | - | - | 8.9 | - | - | - | - | - | - |
| 248 | *Vibrio metschnikovii* | HLS | 5.7 | 7.2 | 8.6 | 17.5 | - | 4.9 | - | - | 46.3 | - | - | - |
| 250 | *Alteromonas stellipolaris* | HLS | 9.3 | 20.1 | 22.4 | - | - | 8.1 | - | - | - | - | - | - |
| 303 | *Gallaecimonas pentaromativorans* | HLS | 11.8 | 23.2 | 22.6 | - | - | 9.3 | - | - | - | - | - | - |
| 306 | *Exiguobacterium* sp. | HLS | - | - | - | - | - | 58.9 | - | 66.9 | - | - | - | - |
| 323 | *Flavobacterium ponti* | HLS | 8.5 | - | - | - | - | 21.0 | - | - | - | - | - | - |
| 325 | *Gallaecimonas pentaromativorans* | HLS | 12.5 | 23.1 | 23.6 | - | - | 13.9 | - | - | - | - | - | - |
| 609 | *Agarivorans* sp. | HLS | - | - | - | - | - | 15.2 | - | - | - | - | - | - |
| 611 | *Micrococcus* sp. | HLS | 14.9 | 82.7 | 75.0 | - | - | 37.4 | - | - | 53.4 | - | - | - |
| 625 | *Flavobacterium jumunjinense* | HLS | 6.9 | 21.7 | 18.4 | 35.4 | - | 16.8 | - | - | - | - | - | - |
| 629 | *Cobetia* sp. | HLS | 11.8 | 36.9 | 28.6 | - | - | 18.8 | - | - | - | - | - | - |
| 639 | *Paracoccus* sp. | HLS | - | - | - | - | - | 19.5 | - | - | - | - | - | - |
| 640 | *Neptunomoas phycophilaNeptunomoas* | HLS | 5.6 | 20.6 | 15.5 | 22.5 | - | 21.7 | - | - | - | - | - | - |
| 716 | *Pantoea* sp. | HLS | 6.3 | 23.8 | 13.0 | 23.3 | - | 0.9 | - | - | - | - | - | - |
| 729 | *Microbacterium* sp. | HLS | - | - | - | - | - | - | - | 53.7 | - | - | - | - |
| 741 | *Psychrobacter* sp. | HLS | 12.2 | 55.0 | - | - | - | 14.5 | - | - | - | - | - | - |
| 202b | *Pseudoalteromonas* sp. | HLS | 11.7 | 18.8 | 22.0 | 70.8 | - | 19.4 | - | - | - | - | - | - |
| 610a | *Flavobacterium jumunjinense* | HLS | 21.9 | - | 86.8 | - | - | 48.8 | - | 5.7 | - | - | - | - |
| 713b | *Proteus* sp. | HLS | 5.8 | 50.1 | 23.6 | - | - | 10.7 | - | - | - | - | - | - |
| x001 | *Cobetia* sp. | HLS | 2.1 | 6.1 | 6.8 | 6.6 | - | 8.2 | - | - | - | - | - | - |
| x007 | *Alkalihalobacillus hwajinpoensis* | HLS | 6.1 | 18.4 | 14.3 | 22.2 | - | 12.6 | - | - | - | - | - | 6.2 |
| x027 | *Zobellia amurskyensis* | HLS | - | - | - | - | - | - | - | - | - | - | - | - |
| x028 | *Pseudomonas* sp. | HLS | 5.7 | - | - | - | - | 19.7 | - | - | - | - | - | - |
| x030 | *Glaciecola siphonariae* | HLS | - | - | - | - | - | - | - | - | - | - | - | - |
| x034 | *Algoriphagus jejuensis* | HLS | - | - | - | - | - | 57.6 | - | - | - | - | - | - |
| 205 | Alteromonadaceae sp. | DLS | 11.3 | 20.3 | 82.9 | - | - | 8.0 | - | - | - | - | - | - |
| 229 | *Novosphingobium decolorationis* | DLS | - | - | - | - | - | 9.1 | - | 54.2 | - | - | - | - |
| 614 | *Aquimarina muelleri* | DLS | 10.9 | 69.6 | 68.4 | - | - | 26.9 | - | - | - | - | - | - |
| 616 | *Sulfitobacter pontiacus* | DLS | - | - | - | - | - | 6.8 | - | - | - | - | - | - |
| 632 | *Alteromonas stellipolaris* | DLS | 10.9 | 22.0 | 38.5 | - | - | 16.1 | - | - | - | - | - | - |
| DAI04 | *Pseudoalteromonas tetraodonis* | DLS | 7.1 | 18.5 | 20.9 | 20.9 | - | 7.5 | - | - | - | - | - | - |
| x025 | *Streptomyces griseorubens* | DLS | 20.8 | 65.7 | 7.0 | 72.9 | - | 6.6 | 74.3 | 45.0 | - | - | - | - |

| **Strain** | **Identification** | **Source** | **Ecas** | **Ef** | **Efm** | **Eh** | **Ec** | **MRSA** | **Ea** | **Pi** | **Mg** | **Pss** | **Rs** | **Xc** |
|---|---|---|---|---|---|---|---|---|---|---|---|---|---|---|
| 115 | *Novosphingobium* sp. | RS | - | - | - | - | - | - | - | - | - | - | - | - |
| 137 | *Rhizobium* sp. | RS | - | - | - | - | - | 7.3 | - | - | - | - | - | - |
| 305 | *Flavobacterium* sp. | RS | - | - | - | - | - | - | - | - | - | - | - | - |
| 630 | Alteromonadaceae sp. | RS | 7.3 | 16.9 | 24.5 | 78.0 | - | 8.0 | - | - | - | - | - | - |
| 725 | *Bacillus* sp. | RS | 6.6 | 17.9 | 18.1 | 21.8 | - | 15.2 | - | - | - | - | - | - |
| 6RBSU | *Microbacterium* sp. | RS | - | - | - | - | - | - | - | - | 93.9 | - | - | - |
| 711b | *Flavobacterium* sp. | RS | - | - | - | - | - | 42.1 | - | - | - | - | - | - |
| RBI15 | *Pseudoalteromonas* sp. | RS | 4.4 | 7.1 | 6.3 | 17.5 | - | 6.7 | - | 85.8 | - | - | - | - |
| x005 | *Dokdonia* sp. | RS | - | - | - | - | - | 32.8 | - | - | - | - | - | - |
| x018 | *Octadecabacter* sp. | RS | - | - | - | - | - | 8.8 | - | - | - | - | - | - |
| x033 | *Olleya marilimosa* | RS | 14.5 | 82.7 | 95.0 | - | - | 44.9 | - | - | - | - | - | - |
| xRASU | *Rheinheimera baltica* | RS | 8.6 | 17.7 | 21.0 | 25.4 | - | 12.2 | - | - | - | - | - | - |
| 704 | *Bacillus* sp. | HLI | 10.0 | 23.2 | 21.3 | 31.3 | - | 23.3 | - | - | 46.5 | - | - | - |
| 742 | *Bacillus* sp. | HLI | 2.7 | 12.7 | 6.9 | 15.5 | - | 7.5 | - | 28.2 | 28.7 | - | - | 6.2 |
| 214a | *Bacillus megaterium* | HLI | - | - | - | - | - | - | - | - | - | - | - | - |
| 3LBT | *Bacillus pumilus* | HLI | - | - | - | - | - | - | - | - | - | - | - | - |
| x015 | *Bacillus* sp. | HLI | 35.7 | - | 6.9 | 19.3 | - | 10.2 | - | - | - | - | - | 2.1 |
| 144 | *Pseudoalteromonas* sp. | DLI | - | - | - | - | - | - | - | - | - | - | - | - |
| 237 | *Thalassospira lucentensis* | DLI | 6.4 | 30.5 | 72.5 | - | - | 7.7 | - | - | - | - | - | - |
| 431 | *Mycolicibacterium sediminis* | DLI | - | - | - | - | - | - | - | - | - | - | - | - |
| 436 | *Paenibacillus sp.* | DLI | 3.8 | 6.7 | 7.1 | 7.2 | - | 5.3 | - | 35.1 | 23.4 | - | - | 5.4 |
| 706 | *Oceanisphaera sediminis* | DLI | - | - | - | - | - | 78.9 | - | - | - | - | - | - |
| 747 | *Loktanella* sp. | DLI | - | - | - | - | - | - | - | - | - | - | - | - |
| 332b | *Oceanobacillus* sp. | DLI | 8.7 | 21.6 | 24.8 | 25.1 | - | 19.3 | - | - | - | - | - | 7.4 |
| x014 | *Aquimarina* sp. | DLI | 6.3 | 55.1 | 36.3 | - | - | 20.5 | - | - | - | - | - | - |
| x019 | *Streptomyces* sp. | DLI | 8.5 | 19.5 | 18.0 | 19.4 | - | 20.1 | 18.5 | 6.5 | **0.6** | - | - | - |
| x037 | *Cellulophaga pacifica* | DLI | - | - | - | - | - | 46.0 | - | - | - | - | - | - |
| 334 | *Breoghania corrubedonensis* | RI | - | - | - | - | - | 4.9 | - | - | - | - | - | - |
| 641 | *Roseibium marinum* | RI | - | - | - | - | - | 10.3 | - | - | - | - | - | - |
| x012 | *Shewanella* sp. | RI | 7.7 | 17.6 | 19.5 | 21.1 | - | 9.7 | - | - | - | - | - | - |
| x029 | *Rhodococcus* | RI | - | - | - | - | - | - | - | - | - | - | - | - |
| x039 | *Hoeflea alexandrii* | RI | 4.4 | - | - | - | - | 10.5 | - | - | - | - | - | - |
| 218 | *Curtobacterium herbarum* | W | - | - | - | - | - | - | - | - | - | - | - | - |
| 733 | *Rhodococcus* sp. | S | - | - | - | - | - | - | - | - | - | - | - | - |
| 738 | *Streptomyces* sp. | S | 7.4 | 17.1 | 15.5 | 21.2 | - | 19.5 | - | - | - | - | - | - |
| 744 | *Streptomyces* sp. | S | 17.3 | 54.2 | 23.9 | 74.9 | - | 25.7 | - | 1.3 | 2.8 | - | - | - |
| xSBU1 | *Pseudoalteromonas ulvae* | S | 7.1 | 17.3 | 17.7 | 20.6 | - | 8.8 | - | - | - | - | - | - |
| Pos | Positive control | - | 2.4 | 0.5 | 0.2 | 0.8 | 6.4 | 1.5 | 1.1 | 0.01 | 0.3 | 1.6 | 0.8 | 2.4 |

**Supplementary Table S6:** Bioactivity of bacterial MA medium extracts against aquatic panel. $IC_{50}$ values are in µg/ml. HLS: Healthy leaf surface, HLI: Healthy leaf inner tissue, DLS: Decaying leaf surface, DLI: Decaying leaf inner tissue, RS: Root surface, RI: Root inner tissue, W: Seawater reference, S: Sediment reference. La: *Leifsonia aquatica*, Lg: *Lactococcus garvieae*, Ab: *Algicola bacteriolytica*, Pe: *Pseudoalteromonas elyacovii*, Sha: *Shewanella algae*, Vae: *Vibrio aestuarianus*, Val: *Vibrio alginolyticus*, Va: *Vibrio anguillarum*, Vch: *Vibrio cholerae*, Vco: *Vibrio coralliilyticus*, Vf: *Vibrio fischeri*, Vh: *Vibrio harveyi*, Vi: *Vibrio ichthyoenteri*, Vp: *Vibrio parahaemolyticus*, Vsp: *Vibrio splendidus*, Vv: *Vibrio vulnificus*. Positive control: chloramphenicol, except for *Lactococcus garvieae* (ampicillin).

| Strain | Identification | Origin | La | Lg | Ab | Pe | Sha | Vae | Val | Va | Vch | Vco | Vf | Vh | Vi | Vp | Vsp | Vv |
|---|---|---|---|---|---|---|---|---|---|---|---|---|---|---|---|---|---|---|
| 111 | *Pedobacter silvitoris* | HLS | - | - | - | - | - | - | - | - | - | - | - | - | - | - | - | - |
| 113 | *Brevundimonas vesicularis* | HLS | - | - | - | - | - | - | - | - | - | - | - | - | - | - | - | - |
| 117 | *Kocuria* sp. | HLS | - | - | - | - | - | - | - | - | - | - | - | - | - | - | - | - |
| 122 | *Microbacterium* sp. | HLS | 93.1 | - | - | - | 56.6 | - | - | - | - | - | 86.6 | - | 71.2 | - | - | - |
| 131 | *Streptomyces* sp. | HLS | **0.6** | **1.1** | **0.8** | - | - | - | - | - | - | - | - | - | - | - | - | - |
| 150 | *Stenotrophomonas* sp. | HLS | 5.9 | 8.4 | - | - | - | - | - | - | - | - | - | - | - | - | - | - |
| 204 | *Falsirhodobacter* sp. | HLS | - | - | - | - | - | - | - | - | - | - | - | - | - | - | - | - |
| 248 | *Vibrio metschnikovii* | HLS | 17.5 | - | - | - | - | - | - | - | - | - | - | - | - | - | - | - |
| 250 | *Alteromonas stellipolaris* | HLS | 21.5 | - | - | - | - | - | - | - | - | - | - | - | - | - | - | - |
| 303 | *Gallaecimonas pentaromativorans* | HLS | 20.3 | - | - | - | - | - | - | - | - | - | - | - | - | - | - | - |
| 306 | *Exiguobacterium* sp. | HLS | - | - | - | - | - | - | - | - | - | - | - | - | - | - | - | - |
| 323 | *Flavobacterium ponti* | HLS | - | - | - | - | - | - | - | - | - | - | - | - | - | - | - | - |
| 325 | *Gallaecimonas pentaromativorans* | HLS | 11.9 | - | - | - | - | - | - | - | - | - | - | - | - | - | - | - |
| 609 | *Agarivorans* sp. | HLS | 6.2 | 21.3 | - | - | - | - | - | - | - | - | - | - | - | - | - | - |
| 611 | *Micrococcus* sp. | HLS | 26.4 | - | - | - | - | - | - | - | - | - | - | - | - | - | - | - |
| 625 | *Flavobacterium jumunjinense* | HLS | 25.6 | - | - | - | - | - | - | - | - | - | - | - | - | - | - | - |
| 629 | *Cobetia* sp. | HLS | 14.1 | - | - | - | - | - | - | - | - | - | - | - | - | - | - | - |
| 639 | *Paracoccus* sp. | HLS | - | - | - | - | - | - | - | - | - | - | - | - | - | - | - | - |
| 640 | *Neptunomoas phycophila* | HLS | 23.3 | - | - | - | - | - | - | - | - | - | - | - | - | - | - | - |
| 716 | *Pantoea* sp. | HLS | 8.7 | 58.6 | - | - | - | - | - | - | - | - | - | - | - | - | - | - |
| 729 | *Microbacterium* sp. | HLS | 20.0 | - | - | - | - | - | - | - | - | - | - | - | - | - | - | - |
| 741 | *Psychrobacter* sp. | HLS | 10.0 | - | - | - | - | - | - | - | - | - | - | - | - | - | - | - |
| 202b | *Pseudoalteromonas* sp. | HLS | 6.3 | 78.9 | - | - | - | - | - | - | - | - | - | - | - | - | - | - |
| 610a | *Flavobacterium jumunjinense* | HLS | 85.2 | - | - | - | - | - | - | - | - | - | - | - | - | - | - | - |
| 713b | *Proteus* sp. | HLS | 7.7 | 27.1 | 16.9 | - | - | - | - | - | - | - | - | - | - | - | - | - |
| x001 | *Cobetia* sp. | HLS | 6.8 | 18.4 | 48.1 | - | - | - | - | - | - | - | - | - | 17.6 | - | - | - |
| x007 | *Alkalihalobacillus hwajinpoensis* | HLS | 7.0 | - | - | - | - | - | - | - | - | - | - | - | - | - | - | - |
| x027 | *Zobellia amurskyensis* | HLS | - | - | - | - | - | - | - | - | - | - | - | - | - | - | - | - |
| x028 | *Pseudomonas* sp. | HLS | - | - | - | - | - | - | - | - | - | - | - | - | - | - | - | - |
| x030 | *Glaciecola* sp. | HLS | 15.7 | - | - | - | - | - | - | - | - | - | - | - | - | - | - | - |
| x034 | *Algoriphagus jejuensis* | HLS | 26.2 | - | - | - | - | - | - | - | - | - | - | - | - | - | - | - |
| 205 | Alteromonadaceae sp. | DLS | 21.5 | - | - | - | - | - | - | - | - | - | - | - | - | - | - | - |
| 229 | *Novosphingobium decolorationis* | DLS | 45.6 | - | - | - | - | - | - | - | - | - | - | - | - | - | - | - |
| 614 | *Aquimarina muelleri* | DLS | 86.9 | - | - | - | - | - | - | - | - | - | - | - | 81.7 | - | - | - |
| 616 | *Sulfitobacter pontiacus* | DLS | - | - | - | - | - | - | - | - | - | - | - | - | - | - | - | - |
| 632 | *Alteromonas stellipolaris* | DLS | 17.4 | - | - | - | - | - | - | - | - | - | - | - | 85.2 | - | - | - |
| DAI04 | *Pseudoalteromonas tetraodonis* | DLS | 6.2 | 54.9 | - | - | - | - | - | - | - | - | - | - | - | - | - | - |
| x025 | *Streptomyces griseorubens* | DLS | 19.5 | - | - | - | - | - | - | - | - | - | - | - | - | - | - | - |
| 115 | *Novosphingobium* sp. | RS | - | - | - | - | - | - | - | - | - | - | - | - | - | - | - | - |

| Strain | Identification | Origin | La | Lg | Ab | Pe | Sha | Vae | Val | Va | Vch | Vco | Vf | Vh | Vi | Vp | Vsp | Vv |
|---|---|---|---|---|---|---|---|---|---|---|---|---|---|---|---|---|---|---|
| 137 | Rhizobium sp. | RS | 10.8 | - | - | - | - | - | - | - | - | - | - | - | - | - | - | - |
| 305 | *Flavobacterium* sp. | RS | - | - | - | - | - | - | - | - | - | - | - | - | - | - | - | - |
| 630 | Alteromonadaceae sp. | RS | 20.8 | - | - | - | - | - | - | - | - | - | - | - | - | - | - | - |
| 725 | *Bacillus* sp. | RS | 8.1 | - | - | - | - | - | - | - | - | - | - | - | - | - | - | - |
| 6RBSU | *Microbacterium* sp. | RS | - | - | - | - | - | - | - | - | - | - | - | - | - | - | - | - |
| 711b | *Flavobacterium* sp. | RS | 19.7 | - | - | - | - | - | - | - | - | - | - | - | - | - | - | - |
| RBI15 | *Pseudoalteromonas* sp. | RS | 16.6 | - | - | - | - | - | - | - | - | - | - | - | - | - | - | - |
| x005 | *Dokdonia* sp. | RS | 21.4 | - | - | - | - | - | - | - | - | - | - | - | - | - | - | - |
| x018 | *Octadecabacter* sp. | RS | - | - | - | - | - | - | - | - | - | - | - | - | - | - | - | - |
| x033 | *Olleya marilimosa* | RS | 68.6 | - | - | - | - | - | - | - | - | - | - | - | - | - | - | - |
| xRASU | *Rheinheimera baltica* | RS | 22.0 | - | - | - | - | - | - | - | - | - | - | - | - | - | - | - |
| 704 | *Bacillus* sp. | HLI | 10.3 | - | - | - | - | - | - | - | - | - | - | - | - | - | - | - |
| 742 | *Bacillus* sp. | HLI | 5.9 | 34.2 | - | - | - | - | - | - | - | - | - | - | - | - | - | - |
| 214a | *Bacillus megaterium* | HLI | 73.2 | - | - | - | - | - | - | - | - | - | 76.1 | - | - | - | - | - |
| 3LBT | *Bacillus pumilus* | HLI | - | - | - | - | - | - | - | - | - | - | - | - | - | - | - | - |
| x015 | *Bacillus* sp. | HLI | 12.5 | - | - | - | - | - | - | - | - | - | - | - | - | - | - | - |
| 144 | *Pseudoalteromonas* sp. | DLI | - | - | - | - | - | - | - | - | - | - | - | - | - | - | - | - |
| 237 | *Thalassospira lucentensis* | DLI | 9.0 | - | - | - | - | - | - | - | - | - | - | - | - | - | - | - |
| 431 | *Mycolicibacterium sediminis* | DLI | - | - | - | - | - | - | - | - | - | - | - | - | - | - | - | - |
| 436 | *Paenibacillus sp.* | DLI | 71.8 | - | - | - | - | - | - | - | - | - | - | - | - | - | - | - |
| 706 | *Oceanisphaera sediminis* | DLI | - | - | - | - | - | - | - | - | - | - | - | - | - | - | - | - |
| 747 | *Loktanella* sp. | DLI | - | - | - | - | - | - | - | - | - | - | - | - | - | - | - | - |
| 332b | *Oceanobacillus* sp. | DLI | 7.6 | - | - | - | - | - | - | - | - | - | - | - | - | - | - | - |
| x014 | *Aquimarina* sp. | DLI | 8.1 | 28.5 | - | - | - | - | - | - | - | - | - | - | - | - | - | - |
| x019 | *Streptomyces* sp. | DLI | 3.3 | 20.9 | - | - | 60.8 | - | - | - | - | - | 45.0 | - | 36.3 | - | - | - |
| x037 | *Cellulophaga pacifica* | DLI | 41.7 | - | - | - | - | - | - | - | - | - | - | - | 57.9 | - | - | - |
| 334 | *Breoghania corrubedonensis* | RI | 30.2 | - | - | - | - | - | - | - | - | - | - | - | - | - | - | - |
| 641 | *Roseibium marinum* | RI | 73.1 | - | - | - | - | - | - | - | - | - | - | - | - | - | - | - |
| x012 | *Shewanella* sp. | RI | 6.9 | 22.0 | - | - | - | - | - | - | - | - | - | - | - | - | - | - |
| x029 | Rhodococcus sp. | RI | - | - | - | - | - | - | - | - | - | - | - | - | - | - | - | - |
| x039 | *Hoeflea alexandrii* | RI | 6.6 | - | - | - | - | - | - | - | - | - | - | - | - | - | - | - |
| 218 | *Curtobacterium herbarum* | W | - | - | - | - | - | - | - | - | - | - | - | - | - | - | - | - |
| 146 | *Micromonospora* sp. | S | - | - | - | - | - | - | - | - | - | - | - | - | - | - | - | - |
| 733 | *Rhodococcus* sp. | S | - | - | - | - | - | - | - | - | - | - | - | - | - | - | - | - |
| 738 | *Streptomyces* sp. | S | 8.1 | 21.7 | - | - | - | - | - | - | 64.3 | - | - | - | 20.9 | - | - | - |
| 744 | *Streptomyces* sp. | S | - | - | - | - | - | - | - | - | - | - | - | - | - | - | - | - |
| xSBU1 | *Pseudoalteromonas ulvae* | S | 15.5 | - | - | - | - | - | - | - | - | - | - | - | - | - | - | - |
| Pos | Positive control | - | 0.4 | 0.5 | 2.9 | 2.3 | 4.1 | 1.2 | 0.8 | 0.4 | 0.5 | 0.7 | 0.6 | 0.7 | 0.4 | 0.4 | 0.5 | 0.2 |

**Supplementary Table S7:** Bioactivity of bacterial MA medium extracts against fecal, human and plant pathogenic panel. $IC_{50}$ values are in µg/ml. HLS: Healthy leaf surface, HLI: Healthy leaf inner tissue, DLS: Decaying leaf surface, DLI: Decaying leaf inner tissue, RS: Root surface, RI: Root inner tissue, W: Seawater reference, S: Sediment reference. Fecal pathogens (positive control) Ecas*:* *Enterococcus casseliflavus* (ampicillin), Ef: *Enterococcus faecalis* (ampicillin), Efm: *Enterococcus faecium* (ampicillin), Eh: *Enterococcus hirae* (ampicillin), Ec: *Escherichia coli* (chloramphenicol). Human pathogens (positive control) MRSA: Methicillin resistant *Staphylococcus aureus* (chloramphenicol). Phytopathogens (positive control) Ea: *Erwinia amylovora* (chloramphenicol), Pi: *Phytophthora infestans* (cycloheximid), Mg: *Magnaphorte grisea* (nystatin), Pss: *Pseudomonas syringae* (chloramphenicol), Rs: *Ralstonia solanacearum* (tetracycline), Xc: *Xanthomonas campestris* (chloramphenicol).

| Strain | Identification | Source | Ecas | Ef | Efm | Eh | Ec | MRSA | Ea | Pi | Mg | Pss | Rs | Xc |
|---|---|---|---|---|---|---|---|---|---|---|---|---|---|---|
| 111 | *Pedobacter silvitoris* | HLS | - | - | - | - | - | - | - | - | - | - | - | - |
| 113 | *Brevundimonas vesicularis* | HLS | - | - | - | - | - | - | - | - | - | - | - | - |
| 117 | *Kocuria* sp. | HLS | - | - | - | - | - | - | - | 32.8 | - | - | - | - |
| 122 | *Microbacterium* sp. | HLS | - | - | - | - | - | - | - | 23.9 | - | - | - | - |
| 131 | *Streptomyces* sp. | HLS | 0.3 | 0.2 | 0.6 | 0.7 | - | 0.6 | - | - | 0.5 | - | - | - |
| 150 | *Stenotrophomonas* sp. | HLS | 4.9 | 6.5 | 6.6 | 9.2 | - | 15.1 | - | - | - | - | - | - |
| 204 | *Falsirhodobacter* sp. | HLS | - | - | - | - | - | 54.5 | - | - | - | - | - | - |
| 248 | *Vibrio metschnikovii* | HLS | 5.8 | - | 9.8 | - | - | 22.8 | - | 0.9 | - | - | - | - |
| 250 | *Alteromonas stellipolaris* | HLS | - | - | - | - | - | 22.4 | - | - | - | - | - | - |
| 303 | *Gallaecimonas pentaromativorans* | HLS | 26.7 | - | 66.4 | - | - | 50.8 | - | - | - | - | - | - |
| 306 | *Exiguobacterium* sp. | HLS | - | - | - | - | - | - | - | - | - | - | - | - |
| 323 | *Flavobacterium ponti* | HLS | - | - | - | - | - | - | - | - | - | - | - | - |
| 325 | *Gallaecimonas pentaromativorans* | HLS | 31.5 | 52.5 | 53.8 | - | - | 25.4 | - | - | - | - | - | - |
| 609 | *Agarivorans* sp. | HLS | 12.8 | 17.0 | 16.7 | 20.2 | - | 17.6 | - | - | 59.9 | - | - | - |
| 611 | *Micrococcus* sp. | HLS | - | - | - | - | - | 73.7 | - | - | - | - | - | - |
| 625 | *Flavobacterium jumunjinense* | HLS | 20.6 | 59.2 | 47.5 | - | - | 26.3 | - | - | - | - | - | - |
| 629 | *Cobetia* sp. | HLS | 20.0 | 25.9 | 28.9 | - | - | 22.4 | - | - | - | - | - | - |
| 639 | *Paracoccus* sp. | HLS | - | - | - | - | - | 83.0 | - | - | - | - | - | - |
| 640 | *Neptunomonas phycophila* | HLS | 27.5 | 81.8 | 97.6 | - | - | 77.0 | - | - | - | - | - | - |
| 716 | *Pantoea* sp. | HLS | 12.0 | 17.9 | 19.1 | 22.9 | - | 18.7 | - | - | - | - | - | - |
| 729 | *Microbacterium* sp. | HLS | - | - | - | - | - | - | - | 80.6 | - | - | - | - |
| 741 | *Psychrobacter* sp. | HLS | 17.0 | 60.7 | - | - | - | 11.7 | - | 64.3 | - | - | - | - |
| 202b | *Pseudoalteromonas* sp. | HLS | 18.1 | 20.5 | 37.2 | 69.3 | - | 20.9 | - | - | - | - | - | - |
| 610a | *Flavobacterium jumunjinense* | HLS | - | - | - | - | - | 65.8 | - | 9.6 | - | - | - | - |
| 713b | *Proteus* sp. | HLS | 14.6 | 21.2 | 48.7 | 39.4 | - | 18.9 | - | - | - | - | - | - |
| x001 | *Cobetia* sp. | HLS | 2.5 | 3.1 | 6.1 | 6.2 | - | 6.9 | - | - | - | - | - | - |
| x007 | *Alkalihalobacillus hwajinpoensis* | HLS | 25.8 | 67.0 | 24.3 | - | - | 68.9 | - | 57.6 | - | - | - | - |
| x027 | *Zobellia amurskyensis* | HLS | - | - | - | - | - | - | - | - | - | - | - | - |
| x028 | *Pseudomonas* sp. | HLS | - | - | - | - | - | 90.6 | - | - | - | - | - | - |
| x030 | *Glaciecola siphonariae* | HLS | 18.2 | 22.7 | 23.5 | - | - | 20.1 | - | - | - | - | - | - |
| x034 | *Algoriphagus jejuensis* | HLS | - | - | - | - | - | 55.5 | - | - | - | - | - | - |
| 205 | Alteromonadaceae sp. | DLS | 26.0 | 92.7 | - | - | - | 38.1 | - | - | - | - | - | - |
| 229 | *Novosphingobium decolorationis* | DLS | - | - | - | - | - | 10.2 | - | - | - | - | - | - |
| 614 | *Aquimarina muelleri* | DLS | 29.8 | - | - | - | - | 80.3 | - | - | - | - | - | - |
| 616 | *Sulfitobacter pontiacus* | DLS | - | - | - | - | - | 20.6 | - | - | - | - | - | - |
| 632 | *Alteromonas stellipolaris* | DLS | 25.4 | - | - | - | - | 13.3 | - | - | - | - | - | - |
| DAI04 | *Pseudoalteromonas tetraodonis* | DLS | 13.4 | 28.0 | 65.9 | - | - | 21.9 | | | | | | |
| x025 | *Streptomyces griseorubens* | DLS | 21.5 | 56.9 | 19.5 | - | - | 21.9 | - | - | - | - | - | - |

| Strain | Identification | Source | Ecas | Ef | Efm | Eh | Ec | MRSA | Ea | Pi | Mg | Pss | Rs | Xc |
|---|---|---|---|---|---|---|---|---|---|---|---|---|---|---|
| 115 | *Novosphingobium* sp. | RS | - | - | - | - | - | - | - | - | - | - | - | - |
| 137 | *Rhizobium* sp. | RS | - | - | - | - | - | 15.9 | - | - | - | - | - | - |
| 305 | *Flavobacterium* sp. | RS | - | - | - | - | - | - | - | - | - | - | - | - |
| 630 | Alteromonadaceae sp. | RS | 33.6 | 67.2 | 51.7 | - | - | 28.3 | - | - | - | - | - | - |
| 725 | *Bacillus* sp. | RS | 17.8 | 66.0 | 68.9 | - | - | 49.4 | - | - | - | - | - | - |
| 6RBSU | *Microbacterium* sp. | RS | - | - | - | - | - | - | - | 53.2 | - | - | - | - |
| 711b | *Flavobacterium* sp. | RS | 21.4 | 36.0 | 69.6 | - | - | 27.3 | - | - | - | - | - | - |
| RBI15 | *Pseudoalteromonas* sp. | RS | 19.8 | 24.6 | 27.9 | - | - | 22.7 | - | - | - | - | - | - |
| x005 | *Dokdonia* sp. | RS | 4.7 | - | 25.1 | - | - | 23.1 | - | - | - | - | - | - |
| x018 | *Octadecabacter* sp. | RS | - | - | - | - | - | 71.4 | - | - | - | - | - | - |
| x033 | *Olleya marilimosa* | RS | - | - | - | - | - | - | - | - | - | - | - | - |
| xRASU | *Rheinheimera baltica* | RS | 23.4 | 40.6 | 66.6 | - | - | 26.1 | - | - | - | - | - | - |
| 704 | *Bacillus* sp. | HLI | 24.4 | 66.1 | 25.5 | - | - | 58.8 | - | - | - | - | - | - |
| 742 | *Bacillus* sp. | HLI | 16.5 | 25.4 | 21.9 | 53.4 | - | 41.2 | - | 67.5 | 84.2 | - | - | - |
| 214a | *Bacillus megaterium* | HLI | 95.4 | - | - | - | - | - | - | - | - | - | - | - |
| 3LBT | *Bacillus pumilus* | HLI | - | - | - | - | - | - | - | - | - | - | - | - |
| x015 | *Bacillus* sp. | HLI | 10.8 | 25.1 | 15.3 | 23.1 | - | 2.2 | - | - | - | - | - | - |
| 144 | *Pseudoalteromonas* sp. | DLI | - | - | - | - | - | - | - | - | - | - | - | - |
| 237 | *Thalassospira lucentensis* | DLI | 21.8 | 54.0 | - | - | - | 10.3 | - | - | - | - | - | - |
| 431 | *Mycolicibacterium sediminis* | DLI | - | - | - | - | - | - | - | - | - | - | - | - |
| 436 | *Paenibacillus sp.* | DLI | 5.3 | - | 18.4 | - | - | 14.5 | - | - | - | - | - | - |
| 706 | *Oceanisphaera sediminis* | DLI | - | - | - | - | - | 98.6 | - | - | - | - | - | - |
| 747 | *Loktanella* sp. | DLI | - | - | - | - | - | - | - | - | - | - | - | - |
| 332b | *Oceanobacillus* sp. | DLI | - | - | - | - | - | 89.3 | - | 60.6 | - | - | - | - |
| x014 | *Aquimarina* sp. | DLI | 14.6 | 21.7 | 18.0 | 32.6 | - | 22.0 | - | - | - | - | - | - |
| x019 | *Streptomyces* sp. | DLI | 12.3 | 22.3 | 17.6 | 53.9 | - | 19.7 | - | 4.0 | 0.2 | - | - | - |
| x037 | *Cellulophaga pacifica* | DLI | 20.7 | - | 51.9 | - | - | 72.2 | - | - | - | - | - | - |
| 334 | *Breoghania corrubedonensis* | RI | - | - | - | - | - | 8.3 | - | - | - | - | - | - |
| 641 | *Roseibium marinu* | RI | - | - | - | - | - | - | - | - | - | - | - | - |
| x012 | *Shewanella* sp. | RI | 11.0 | 15.1 | 20.0 | 21.5 | - | 16.7 | - | - | - | - | - | - |
| x029 | *Rhodococcus* sp. | RI | - | - | - | - | - | - | - | - | - | - | - | - |
| x039 | *Hoeflea alexandrii* | RI | - | - | - | - | - | 10.1 | - | - | - | - | - | - |
| 218 | *Curtobacterium herbarum* | W | - | - | - | - | - | - | - | - | - | - | - | - |
| 146 | *Micromonospora* sp. | S | - | - | - | - | - | 86.3 | - | - | - | - | - | - |
| 733 | *Rhodococcus* sp. | S | - | - | - | - | - | - | - | - | - | - | - | - |
| 738 | *Streptomyces* sp. | S | 8.9 | 17.4 | 19.3 | 20.6 | - | 18.2 | - | - | - | - | - | - |
| 744 | *Streptomyces* sp. | S | - | - | - | - | - | - | - | 2.2 | 2.1 | - | - | - |
| xSBU1 | *Pseudoalteromonas ulvae* | S | 23.4 | 24.1 | 56.4 | - | - | 23.1 | - | - | - | - | - | - |
| Pos | Positive control | - | 2.4 | 0.5 | 0.2 | 0.8 | 6.4 | 1.5 | 1.1 | 0.01 | 0.3 | 1.6 | 0.8 | 2.4 |

**Supplementary Table S8:** Bioactivity of fungal M34 medium extracts against aquatic panel. $IC_{50}$ values are in µg/ml. HLS: Healthy leaf surface, HLI: Healthy leaf inner tissue, DLS: Decaying leaf surface, DLI: Decaying leaf inner tissue, RS: Root surface, RI: Root inner tissue, W: Seawater reference, S: Sediment reference. La: *Leifsonia aquatica*, Lg: *Lactococcus garvieae*, Ab: *Algicola bacteriolytica*, Pe: *Pseudoalteromonas elyacovii*, Sha: *Shewanella algae*, Vae: *Vibrio aestuarianus*, Val: *Vibrio alginolyticus*, Va: *Vibrio anguillarum*, Vch: *Vibrio cholerae*, Vco: *Vibrio coralliilyticus*, Vf: *Vibrio fischeri*, Vh: *Vibrio harveyi*, Vi: *Vibrio ichthyoenteri*, Vp: *Vibrio parahaemolyticus*, Vsp: *Vibrio splendidus*, Vv: *Vibrio vulnificus*. Positive control: chloramphenicol, except for *Lactococcus garvieae* (ampicillin).

| **Strain** | **Identification** | **Source** | **La** | **Lg** | **Ab** | **Pe** | **Sha** | **Vae** | **Val** | **Va** | **Vch** | **Vco** | **Vf** | **Vh** | **Vi** | **Vp** | **Vsp** | **Vv** |
|---|---|---|---|---|---|---|---|---|---|---|---|---|---|---|---|---|---|---|
| 425a | *Parathyridaria* sp. | HLS | 42.9 | 52.8 | - | - | - | - | - | - | - | - | - | - | - | - | - | - |
| 434 | *Parathyridaria* sp. | HLS | 41.4 | - | - | - | - | - | - | - | - | - | - | - | - | - | - | - |
| 906 | *Penicillium* sp. | HLS | - | - | - | - | - | - | - | - | - | - | - | - | - | - | - | - |
| 105 | *Trichoderma harzianum* | DLS | 79.0 | - | 29.9 | - | - | - | - | - | - | | - | - | - | - | - | - |
| 210 | *Pestalotiopsis* sp. | DLS | - | - | - | - | - | - | - | - | - | - | - | - | - | - | - | - |
| 233 | *Cladosporium halotolerans* | DLS | 42.2 | - | - | 90.4 | - | - | - | - | 19.7 | - | - | - | 61.0 | - | - | - |
| 402 | *Neoascochyta* sp. | DLS | - | - | - | - | - | - | - | - | - | - | - | - | - | - | - | - |
| 420 | *Starmerella vitis* | DLS | 54.8 | - | - | - | - | - | - | - | - | - | - | - | - | - | - | - |
| 719 | *Fusarium* sp. | DLS | 15.6 | 11.0 | - | - | - | - | - | - | - | - | - | - | - | - | - | 26.9 |
| 810 | *Cladosporium* sp. 10 | DLS | - | - | - | - | - | - | - | - | - | - | - | - | - | - | - | - |
| 818 | *Paradendryphiella arenariae* | DLS | - | - | - | - | - | - | - | - | - | - | - | - | - | - | - | - |
| 907 | *Cladosporium* sp. | DLS | 10.9 | - | - | - | - | - | - | - | - | - | - | - | 89.2 | - | 56.9 | - |
| 910 | *Asteromyces cruciatus* | DLS | 40.1 | - | - | - | - | - | - | - | - | - | - | - | - | - | - | - |
| 912 | *Cladosporium* sp. | DLS | | - | - | - | - | - | - | - | - | - | - | - | - | - | - | - |
| 403 | *Penicillium glabrum* | RS | 72.0 | - | - | - | - | - | - | - | - | - | - | - | - | - | - | - |
| 417 | *Cystofilobasidium bisporidii* | RS | 8.6 | 89.2 | - | - | - | - | - | - | - | - | - | - | - | - | - | - |
| 903a | *Alternaria* sp. | RS | - | - | - | - | - | - | - | - | - | - | - | - | - | - | - | - |
| 720 | *Acrostalagmus luteoalbus* | DLI | 2.1 | 9.9 | 9.2 | 3.8 | 15.2 | 8.7 | 31.8 | 4.1 | 5.0 | 34.3 | 4.8 | 22.2 | 17.9 | 4.6 | - | 3.5 |
| 147 | *Arthrinium arundinis* | S | - | - | - | - | - | - | - | - | - | - | - | - | - | - | - | - |
| 407 | *Trichoderma viride* | S | - | - | - | - | - | - | - | - | - | - | - | - | - | - | - | - |
| 617b | *Trichoderma* sp. | S | - | - | 54.6 | - | - | - | - | - | - | - | - | - | - | - | - | - |
| 737 | *Plectosphaerella* sp. | S | - | - | - | - | - | - | - | - | - | - | - | - | - | - | - | - |
| 739 | *Talaromyces* sp. | S | 50.1 | - | - | - | - | - | - | - | - | - | - | - | - | - | - | 73.3 |
| 805 | *Mucor* sp. | S | - | - | - | - | - | - | - | - | - | - | - | - | - | - | - | - |
| 809 | *Penicillium olsonii* | S | 9.9 | - | - | - | - | - | - | - | - | - | - | - | - | 21.8 | - | 15.8 |
| 901 | *Trichoderma* sp. | S | 51.3 | - | 35.1 | - | - | - | - | - | - | - | - | - | - | - | - | - |
| 902b | *Trichoderma* sp. | S | 48.5 | - | 54.4 | - | - | - | - | - | - | - | - | - | - | - | - | - |
| 110 | *Cladosporium* sp. | W | 14.5 | - | - | - | - | - | - | - | - | - | - | - | 97.2 | - | - | - |
| 444 | *Cystofilobasidium bisporidii* | W | 16.0 | - | - | - | - | - | - | - | - | - | - | - | - | - | - | - |
| 813 | *Aureobasidium pullulans* | W | 12.4 | - | 30.8 | - | - | - | - | - | - | - | - | - | 5.2 | - | - | - |
| 815 | *Sarocladium strictum* | W | 34.3 | - | 13.0 | - | - | - | - | - | - | - | - | - | - | - | - | - |
| Pos | Positive control | - | 0.4 | 0.5 | 2.9 | 2.3 | 4.1 | 1.2 | 0.8 | 0.4 | 0.5 | 0.7 | 0.6 | 0.7 | 0.4 | 0.4 | 0.5 | 0.2 |

**Supplementary Table S9:** Bioactivity of fungal M34 medium extracts against fecal, human and plant pathogens. $IC_{50}$ values are in µg/ml. HLS: Healthy leaf surface, HLI: Healthy leaf inner tissue, DLS: Decaying leaf surface, DLI: Decaying leaf inner tissue, RS: Root surface, RI: Root inner tissue, W: Seawater reference, S: Sediment reference. Fecal pathogens (positive control) Ecas: *Enterococcus casseliflavus* (ampicillin), Ef: *Enterococcus faecalis* (ampicillin), Efm: *Enterococcus faecium* (ampicillin), Eh: *Enterococcus hirae* (ampicillin), Ec: *Escherichia coli* (chloramphenicol). Human pathogens (positive control) MRSA: Methicillin resistant *Staphylococcus aureus* (chloramphenicol). Phytopathogens (positive control) Ea: *Erwinia amylovora* (chloramphenicol), Pi: *Phytophthora infestans* (cycloheximid), Mg: *Magnaphorte grisea* (nystatin), Pss: *Pseudomonas syringae* (chloramphenicol), Rs: *Ralstonia solanacearum* (tetracycline), Xc: *Xanthomonas campestris* (chloramphenicol).

| Strain | Identification | Source | Ecas | Ef | Efm | Eh | Ec | MRSA | Ea | Pi | Mg | Pss | Rs | Xc |
|---|---|---|---|---|---|---|---|---|---|---|---|---|---|---|
| 425a | *Parathyridaria* sp. | HLS | 16.3 | 12.5 | 13.6 | 31.5 | - | 4.1 | - | - | 4.5 | - | - | - |
| 434 | *Parathyridaria* sp. | HLS | 18.6 | 17.3 | 18.4 | 55.9 | - | 6.5 | - | - | 2.9 | - | - | - |
| 906 | *Penicillium* sp. | HLS | - | - | - | - | - | - | - | - | - | - | - | - |
| 105 | *Trichoderma harzianum* | DLS | - | - | - | - | - | 20.3 | - | - | - | - | - | - |
| 210 | *Pestalotiopsis* sp. | DLS | - | - | - | - | - | - | - | - | - | - | - | - |
| 233 | *Cladosporium halotolerans* | DLS | - | - | - | - | - | 46.4 | - | 2.2 | 38.4 | - | - | - |
| 402 | *Neoascochyta* sp. | DLS | - | - | - | - | - | - | - | - | - | - | - | - |
| 420 | *Starmerella vitis* | DLS | - | - | - | - | - | - | - | - | - | - | - | - |
| 719 | *Fusarium* sp. | DLS | 18.1 | 19.1 | 8.9 | 55.0 | - | 5.8 | - | 1.2 | 9.4 | - | - | - |
| 810 | *Cladosporium* sp. | DLS | - | - | - | - | - | - | - | - | - | - | - | - |
| 818 | *Paradendryphiella arenariae* | DLS | - | - | - | - | - | 21.9 | - | 55.2 | - | - | - | - |
| 907 | *Cladosporium* sp. | DLS | 94.6 | 39.1 | 36.7 | - | - | 17.6 | - | - | - | - | - | - |
| 910 | *Asteromyces cruciatus* | DLS | - | - | - | - | - | 39.4 | - | - | - | - | - | - |
| 912 | *Cladosporium* sp. | DLS | - | - | - | - | - | 58.9 | - | - | - | - | - | - |
| 403 | *Penicillium glabrum* | RS | - | - | - | - | - | 54.2 | - | - | - | - | - | - |
| 417 | *Cystofilobasidium bisporidii* | RS | 23.1 | 37.6 | 26.9 | - | - | 20.4 | - | - | - | - | - | - |
| 903a | *Alternaria* sp. | RS | - | - | - | - | - | 27.7 | - | 24.4 | - | - | - | - |
| 720 | *Acrostalagmus luteoalbus* | DLI | 10.8 | 0.8 | 3.9 | 3.5 | - | 0.2 | 36.4 | 0.5 | - | 11.5 | 8.7 | 3.2 |
| 147 | *Arthrinium arundinis* | S | - | - | - | - | - | 72.5 | - | 69.2 | - | - | - | - |
| 407 | *Trichoderma viride* | S | - | - | - | - | - | 7.0 | - | - | - | - | - | - |
| 617b | *Trichoderma* sp. | S | - | - | - | - | - | 35.7 | - | - | - | - | - | - |
| 737 | *Plectosphaerella* sp. | S | - | - | - | - | - | - | - | - | - | - | - | - |
| 739 | *Talaromyces* sp. | S | - | - | - | - | - | 41.2 | - | - | - | - | - | - |
| 805 | *Mucor* sp. | S | - | - | - | - | - | - | - | - | - | - | - | - |
| 809 | *Penicillium olsonii* | S | 15.4 | - | 14.2 | - | - | 17.1 | - | 28.4 | 13.3 | - | - | - |
| 901 | *Trichoderma* sp. | S | - | - | - | - | - | 27.5 | - | - | - | - | - | - |
| 902b | *Trichoderma* sp. | S | - | - | - | - | - | - | - | - | - | - | - | - |
| 110 | *Cladosporium* sp. | W | 37.4 | 69.6 | - | - | - | 22.5 | - | - | - | - | - | - |
| 444 | *Cystofilobasidium bisporidii* | W | - | 33.3 | 26.5 | - | - | 21.5 | - | - | - | - | - | - |
| 813 | *Aureobasidium pullulans* | W | 2.8 | 43.1 | 28.5 | - | - | 24.4 | - | 89.3 | - | - | - | 8.3 |
| 815 | *Sarocladium strictum* | W | 65.4 | 92.8 | - | - | - | 8.3 | - | - | 74.6 | - | - | - |
| Pos | Positive control | - | 2.4 | 0.5 | 0.2 | 0.8 | 6.4 | 1.5 | 1.1 | 0.01 | 0.3 | 1.6 | 0.8 | 2.4 |

**Supplementary Table S10:** Bioactivity of fungal PDA medium extracts against aquatic panel. $IC_{50}$ values are in µg/ml HLS: Healthy leaf surface, HLI: Healthy leaf inner tissue, DLS: Decaying leaf surface, DLI: Decaying leaf inner tissue, RS: Root surface, RI: Root inner tissue, W: Seawater reference, S: Sediment reference. La: *Leifsonia aquatica*, Lg: *Lactococcus garvieae*, Ab: *Algicola bacteriolytica*, Pe: *Pseudoalteromonas elyacovii*, Sha: *Shewanella algae*, Vae: *Vibrio aestuarianus*, Val: *Vibrio alginolyticus*, Va: *Vibrio anguillarum*, Vch: *Vibrio cholerae*, Vco: *Vibrio coralliilyticus*, Vf: *Vibrio fischeri*, Vh: *Vibrio harveyi*, Vi: *Vibrio ichthyoenteri*, Vp: *Vibrio parahaemolyticus*, Vsp: *Vibrio splendidus*, Vv: *Vibrio vulnificus*. Positive control: chloramphenicol, except for *Lactococcus garvieae* (ampicillin).

| Strain | Identification | Source | La | Lg | Ab | Pe | Sha | Vae | Val | Va | Vch | Vco | Vf | Vh | Vi | Vp | Vsp | Vv |
|---|---|---|---|---|---|---|---|---|---|---|---|---|---|---|---|---|---|---|
| 425a | *Parathyridaria* sp. | HLS | 100.0 | 38.9 | - | - | - | - | - | - | - | - | - | - | - | 32.0 | - | 16.3 |
| 434 | *Parathyridaria* sp. | HLS | 20.7 | 37.2 | - | - | - | - | - | - | - | - | - | - | - | 29.2 | - | 16.0 |
| 906 | *Penicillium* sp. | HLS | - | - | - | - | - | - | - | - | - | - | - | - | - | - | - | - |
| 105 | *Trichoderma harzianum* | DLS | 64.9 | - | 19.3 | - | - | - | - | - | - | - | - | - | - | - | - | - |
| 210 | *Pestalotiopsis* sp. | DLS | - | - | - | - | - | - | - | - | - | - | - | - | - | - | - | - |
| 233 | *Cladosporium halotolerans* | DLS | 28.4 | - | - | - | - | - | - | 75.9 | 15.2 | - | - | - | 67.2 | - | - | - |
| 402 | *Neoascochyta* sp. | DLS | - | - | - | - | - | - | - | - | - | - | - | - | - | - | - | - |
| 420 | *Starmerella* sp. | DLS | 54.8 | - | - | - | - | - | - | - | - | - | - | - | - | - | - | - |
| 719 | *Fusarium* sp. | DLS | 12.2 | 10.8 | - | - | - | - | - | - | - | - | - | - | - | 73.9 | - | 32.0 |
| 810 | *Cladosporium* sp. | DLS | - | - | - | - | - | - | - | - | - | - | - | - | - | - | - | - |
| 818 | *Paradendryphiella arenariae* | DLS | - | - | - | - | - | - | - | - | - | - | - | - | - | - | - | - |
| 907 | *Cladosporium* sp. | DLS | 16.8 | - | - | - | - | - | - | - | - | - | - | - | - | - | - | - |
| 910 | *Asteromyces cruciatus* | DLS | 29.5 | - | - | - | - | - | - | - | - | - | - | - | - | - | - | - |
| 912 | *Cladosporium* sp. | DLS | | - | - | - | - | - | - | - | - | - | - | - | - | - | - | - |
| 403 | *Penicillium glabrum* | RS | - | - | - | - | - | - | - | - | - | - | - | - | - | - | - | - |
| 417 | *Cystofilobasidium bisporidii* | RS | 46.5 | - | - | - | - | - | - | - | - | - | - | - | - | - | - | - |
| 903a | *Alternaria* sp. | RS | - | - | - | - | - | - | - | - | - | - | - | - | - | - | - | - |
| 720 | *Acrostalagmus luteoalbus* | DLI | 2.2 | 6.3 | 15.9 | 3.6 | 14.3 | 5.3 | 17.2 | 7.1 | 2.6 | 26.1 | 3.5 | 8.7 | 13.3 | 4.2 | | 3.7 |
| 147 | *Arthrinium arundinis* | S | - | - | - | - | - | - | - | 44.6 | - | - | 76.8 | - | 72.0 | - | 33.9 | 57.9 |
| 407 | *Trichoderma viride* | S | - | - | 70.8 | - | - | - | - | - | - | - | - | - | - | - | - | - |
| 617b | *Trichoderma* sp. | S | 21.3 | - | 14.7 | - | - | - | - | - | - | - | - | - | - | - | - | - |
| 737 | *Plectosphaerella* sp. | S | - | - | - | - | - | - | - | - | - | - | - | - | - | - | - | - |
| 739 | *Talaromyces* sp. | S | - | - | - | - | - | - | - | - | - | - | 61.0 | - | - | - | - | 35.5 |
| 805 | *Mucor* sp. | S | - | - | - | - | - | - | - | - | - | - | - | - | - | - | - | - |
| 809 | *Penicillium olsonii* | S | 2.4 | 3.2 | - | - | - | - | - | - | - | - | - | - | - | 36.2 | - | 8.5 |
| 901 | *Trichoderma* sp. | S | 55.6 | - | 16.3 | - | - | - | - | - | - | - | - | - | - | - | - | - |
| 902b | *Trichoderma* sp. | S | 44.7 | - | 12.4 | - | - | - | - | - | - | - | - | - | - | - | - | - |
| 110 | *Cladosporium* sp. | W | 17.3 | - | - | - | - | - | - | - | - | - | - | - | - | - | - | - |
| 444 | *Cystofilobasidium bisporidii* | W | 35.3 | - | - | - | - | - | - | - | - | - | - | - | - | - | - | - |
| 813 | *Aureobasidium pullulans* | W | 27.7 | - | 19.4 | - | - | - | - | - | - | - | - | - | - | - | - | - |
| 815 | *Sarocladium strictum* | W | - | - | - | - | - | - | - | - | - | - | - | - | - | - | - | - |
| Pos | Positive control | - | 0.4 | 0.5 | 2.9 | 2.3 | 4.1 | 1.2 | 0.8 | 0.4 | 0.5 | 0.7 | 0.6 | 0.7 | 0.4 | 0.4 | 0.5 | 0.2 |

**Supplementary Table S11:** Bioactivity of fungal PDA medium extracts against fecal, human and plant pathogens. $IC_{50}$ values are in µg/ml HLS: Healthy leaf surface, HLI: Healthy leaf inner tissue, DLS: Decaying leaf surface, DLI: Decaying leaf inner tissue, RS: Root surface, RI: Root inner tissue, W: Seawater reference, S: Sediment reference. Fecal pathogens (positive control) Ecas: *Enterococcus casseliflavus* (ampicillin), Ef: *Enterococcus faecalis* (ampicillin), Efm: *Enterococcus faecium* (ampicillin), Eh: *Enterococcus hirae* (ampicillin), Ec: *Escherichia coli* (chloramphenicol). Human pathogens (positive control) MRSA: Methicillin resistant *Staphylococcus aureus* (chloramphenicol). Phytopathogens (positive control) Ea: *Erwinia amylovora* (chloramphenicol), Pi: *Phytophthora infestans* (cycloheximid), Mg: *Magnaphorte grisea* (nystatin), Pss: *Pseudomonas syringae* (chloramphenicol), Rs: *Ralstonia solanacearum* (tetracycline), Xc: *Xanthomonas campestris* (chloramphenicol).

| Strain | Identification | Source | Ecas | Ef | Efm | Eh | Ec | MRSA | Ea | Pi | Mg | Pss | Rs | Xc |
|---|---|---|---|---|---|---|---|---|---|---|---|---|---|---|
| 425a | *Parathyridaria* sp. | HLS | 17.9 | 7.8 | 6.8 | 17.0 | - | 2.1 | - | 5.1 | 26.4 | - | - | - |
| 434 | *Parathyridaria* sp. | HLS | 9.1 | 6.8 | 9.2 | 27.7 | - | 2.5 | - | 54.0 | 8.3 | - | - | - |
| 906 | *Penicillium* sp. | HL-S | - | - | - | - | - | - | - | 87.8 | - | - | - | - |
| 105 | *Trichoderma harzianum* | DLS | - | - | - | - | - | 86.2 | - | 68.8 | - | - | - | - |
| 210 | *Pestalotiopsis* sp. | DLS | - | - | - | - | - | - | - | - | - | - | - | - |
| 233 | *Cladosporium halotolerans* | DLS | - | - | - | - | - | 50.7 | - | 0.7 | 16.6 | - | - | - |
| 402 | *Neoascochyta* sp. | DLS | - | - | - | - | - | - | - | - | - | - | - | - |
| 420 | *Starmerella* sp. | DLS | 61.5 | - | - | - | - | 80.4 | - | - | - | - | - | - |
| 719 | *Fusarium* sp. | DLS | 3.7 | 18.4 | 5.4 | 46.8 | - | 3.9 | - | 2.6 | 14.1 | - | - | - |
| 810 | *Cladosporium* sp. | DLS | - | - | - | - | - | - | - | - | - | - | - | - |
| 818 | *Paradendryphiella arenariae* | DLS | - | - | - | - | - | 40.5 | - | 10.9 | - | - | - | - |
| 907 | *Cladosporium* sp. | DLS | 37.4 | 47.6 | - | - | - | 32.7 | - | - | - | - | - | - |
| 910 | *Asteromyces cruciatus* | DLS | 61.4 | 79.0 | - | - | - | 25.6 | - | - | - | - | - | - |
| 912 | *Cladosporium* sp. | DLS | - | - | - | - | - | 55.7 | - | - | - | - | - | - |
| 403 | *Penicillium glabrum* | RS | - | - | - | - | - | - | - | - | - | - | - | - |
| 417 | *Cystofilobasidium bisporidii* | RS | - | 98.3 | 79.5 | - | - | 51.6 | - | - | - | - | - | - |
| 903a | *Alternaria* sp. | RS | - | - | - | - | - | 78.5 | - | 18.3 | - | - | - | - |
| 720 | *Acrostalagmus luteoalbus* | DLI | 17.6 | 1.0 | 6.0 | 4.1 | - | 0.5 | 44.4 | 0.7 | - | 15.0 | 9.1 | 4.1 |
| 147 | *Arthrinium arundinis* | S | - | - | - | - | - | 66.1 | - | 19.7 | 65.3 | - | - | - |
| 407 | *Trichoderma viride* | S | - | - | - | - | - | - | - | - | - | - | - | - |
| 617b | *Trichoderma* sp. | S | 93.9 | - | 70.8 | - | - | 31.1 | - | - | 75.9 | - | - | - |
| 737 | *Plectosphaerella* sp. | S | - | - | - | - | - | - | - | - | - | - | - | - |
| 739 | *Talaromyces* sp. | S | - | - | - | - | - | 67.6 | - | - | - | - | - | - |
| 805 | *Mucor* sp. | S | - | - | - | - | - | - | - | - | - | - | - | - |
| 809 | *Penicillium olsonii* | S | 1.9 | 1.7 | 0.9 | 1.8 | - | 1.0 | - | 0.7 | 0.2 | - | - | - |
| 901 | *Trichoderma* sp. | S | - | - | - | - | - | - | - | - | - | - | - | - |
| 902b | *Trichoderma* sp. | S | - | - | - | - | - | 47.1 | - | - | - | - | - | - |
| 110 | *Cladosporium* sp. | W | 39.8 | 56.6 | - | - | - | 25.6 | - | - | - | - | - | - |
| 444 | *Cystofilobasidium bisporidii* | W | 65.0 | 83.7 | - | - | - | 77.6 | - | - | - | - | - | - |
| 813 | *Aureobasidium pullulans* | W | 19.2 | 20.5 | 11.7 | 62.4 | - | 19.8 | - | - | - | - | - | 7.1 |
| 815 | *Sarocladium strictum* | W | - | - | - | - | - | 62.4 | - | - | - | - | - | - |
| Pos | Positive control | - | 2.4 | 0.5 | 0.2 | 0.8 | 6.4 | 1.5 | 1.1 | 0.01 | 0.3 | 1.6 | 0.8 | 2.4 |

**Supplementary Table S12.** The most active ($IC_{50} \leq 10$ µg/ml) bacterial extracts selected for metabolome analysis. HLS: Healthy leaf surface, HLI: Healthy leaf inner tissue, DLS: Decaying leaf surface, DLI: Decaying leaf inner tissue, RS: Root surface, RI: Root inner tissue, W: Seawater reference, S: Sediment reference. Aquatic pathogens (positive control) La: *Leifsonia aquatica* (chloramphenicol), Lg: *Lactococcus garvieae* (Ampicillin), Ab: *Algicola bacteriolytica* (chloramphenicol), Sha: *Shewanella algae* (chloramphenicol), Vae: *Vibrio aestuarianus* (chloramphenicol), Vco: *Vibrio coralliilyticus* (chloramphenicol), Vf: *Vibrio fischeri* (chloramphenicol), Vi: *Vibrio ichthyoenteri* (chloramphenicol), Vp: *Vibrio parahaemolyticus* (chloramphenicol), Vv: *Vibrio vulnificus* (chloramphenicol). Fecal pathogens (positive control) Ecas*:* *Enterococcus casseliflavus* (ampicillin), Ef: *Enterococcus faecalis* (ampicillin), Efm: *Enterococcus faecium* (ampicillin), Eh: *Enterococcus hirae* (ampicillin). Human pathogen (positive control) MRSA: Methicillin resistant *Staphylococcus aureus* (chloramphenicol). Phytopathogens (positive control) Pi: *Phytophthora infestans* (cycloheximid), Mg: *Magnaphorte grisea* (nystatin), Xc: *Xanthomonas campestris* (chloramphenicol).

| | Strain | Identification | Source | Media | Aquatic test panel | | | | | | | | | | Fecal test panel | | | | Human test panel | Plant pathogenic test panel | | |
|---|---|---|---|---|---|---|---|---|---|---|---|---|---|---|---|---|---|---|---|---|---|---|
| | | | | | La | Lg | Ab | Sha | Vae | Vco | Vf | Vi | Vp | Vv | Ecas | Ef | Efm | Eh | MRSA | Pi | Mg | Xc |
| | **Pos** | Positive control | | | 0.4 | 0.5 | 2.9 | 4.1 | 1.2 | 0.7 | 0.6 | 0.4 | 0.4 | 0.2 | 2.4 | 0.5 | 0.2 | 0.8 | 1.5 | 0.01 | 0.3 | 2.4 |
| 1 | **131** | ***Streptomyces* sp.** | **HLS** | MA | **0.6** | **1.1** | **0.8** | | | | | | | | **0.3** | **0.2** | **0.6** | **0.7** | **0.6** | | **0.5** | |
| 2 | | | | GYM | **1.8** | 6.9 | **1.8** | | | | | | | | **1.6** | **2.0** | **2.3** | **6.4** | **2.3** | 6.7 | 1.2 | |
| 3 | 150 | *Stenotrophomonas* sp. | HLS | MA | 5.9 | 8.4 | | | | | | | | | 4.9 | 6.5 | 6.6 | 9.2 | | | | |
| 4 | | | | GYM | 6.6 | 8.6 | | | | | | | | | 4.6 | 8.1 | 6.8 | | 7.2 | | | |
| 5 | 204 | *Falsirhodobacter* sp. | HLS | GYM | | | | | | | | | | | | | | | 8.9 | | | |
| 6 | 248 | *Vibrio metschnikovii* | HLS | MA | | | | | | | | | | | 5.8 | | 9.8 | | | 0.9 | | |
| 7 | | | | GYM | 6.5 | | | | | | | | | | 5.7 | 7.2 | 8.6 | | 4.9 | | | |
| 8 | 250 | *Alteromonas stellipolaris* | HLS | GYM | 6.2 | | | | | | | | | | 9.3 | | | | 8.1 | | | |
| 9 | 303 | *Gallaecimonas pentaromativorans* | HLS | GYM | 6.3 | | | | | | | | | | | | | | 9.3 | | | |
| 10 | 323 | *Flavobacterium ponti* | HLS | GYM | | | | | | | | | | | 8.5 | | | | | | | |
| 11 | 325 | *Gallaecimonas pentaromativorans* | HLS | GYM | 6.8 | | | | | | | | | | | | | | | | | |
| 12 | 609 | *Agarivorans* sp. | HLS | MA | 6.2 | | | | | | | | | | | | | | | | | |
| 13 | 625 | *Flavobacterium jumunjinense* | HLS | GYM | 6.2 | | | | | | | | | | 6.9 | | | | | | | |
| 14 | 629 | *Cobetia* sp. | HLS | GYM | 7.0 | | | | | | | | | | | | | | | | | |
| 15 | 640 | *Neptunomoas phycophila* | HLS | GYM | | | | | | | | | | | 5.6 | | | | | | | |
| 16 | 716 | *Pantoea* sp. | HLS | MA | 8.7 | | | | | | | | | | | | | | | | | |
| 17 | | | | GYM | 6.5 | | | | | | | | | | 6.3 | | | | **0.9** | | | |
| 18 | 741 | *Psychrobacter* sp. | HLS | MA | 10.0 | | | | | | | | | | | | | | | | | |
| 19 | | | | GYM | 7.0 | | | | | | | | | | | | | | | | | |
| 20 | 202b | *Pseudoalteromonas* sp. | HLS | MA | 6.3 | | | | | | | | | | | | | | | | | |
| 21 | | | | GYM | 6.2 | | | | | | | | | | | | | | | | | |
| 22 | 610a | *Flavobacterium jumunjinense* | HLS | MA | | | | | | | | | | | | | | | | 9.6 | | |
| 23 | | | | GYM | | | | | | | | | | | | | | | | 5.7 | | |
| 24 | 713b | *Proteus* sp. | HLS | MA | 7.7 | | | | | | | | | | | | | | | | | |
| 25 | | | | GYM | 7.5 | | | | | | | | | | 5.8 | | | | | | | |
| 26 | x001 | *Cobetia* sp. | HLS | MA | 6.8 | | | | | | | | | | 2.5 | 3.1 | 6.1 | 6.2 | 6.9 | | | |
| 27 | | | | GYM | 5.9 | | | | | | | | | | **2.1** | 6.1 | 6.8 | 6.6 | 8.2 | | | |
| 28 | x007 | *Alkalihalobacillus hwajinpoensis* | HLS | MA | 7.0 | | | | | | | | | | | | | | | | | |
| 29 | | | | GYM | 5.5 | | | | | | | | | | 6.1 | | | | | | | 6.2 |
| 30 | x028 | *Pseudomonas* sp. | HLS | GYM | 6.6 | | | | | | | | | | 5.7 | | | | | | | |
| 31 | 205 | Alteromonadaceae sp. | DLS | GYM | 6.2 | | | | | | | | | | | | | | 8.0 | | | |
| 32 | 229 | *Novosphingobium decolorationis* | DLS | GYM | | | | | | | | | | | | | | | 9.1 | | | |
| 33 | 616 | *Sulfitobacter pontiacus* | DLS | GYM | | | | | | | | | | | | | | | 6.8 | | | |
| 34 | 632 | *Alteromonas stellipolaris* | DLS | GYM | 8.3 | | | | | | | | | | | | | | | | | |
| 35 | DAI04 | *Pseudoalteromonas tetraodonis* | DLS | MA | 6.2 | | | | | | | | | | | | | | | | | |
| 36 | | | | GYM | 9.1 | | | | | | | | | | 7.1 | | | | 7.5 | | | |
| 37 | x025 | *Streptomyces griseorubens* | DLS | GYM | | | 9.4 | | | | | | | | | | 7.0 | | 6.6 | | | |

| | Strain | Identification | Source | Media | Aquatic test panel | | | | | | | | | | Fecal test panel | | | | Human test panel | Plant pathogenic test panel | | |
|---|---|---|---|---|---|---|---|---|---|---|---|---|---|---|---|---|---|---|---|---|---|---|
| | | | | | **La** | **Lg** | **Ab** | **Sha** | **Vae** | **Vco** | **Vf** | **Vi** | **Vp** | **Vv** | **Ecas** | **Ef** | **Efm** | **Eh** | **MRSA** | **Pi** | **Mg** | **Xc** |
| 38 | 137 | *Rhizobium* sp. | RS | GYM | 7.0 | | | | | | | | | | | | | | 7.3 | | | |
| 39 | 630 | Alteromonadaceae sp. | RS | GYM | 6.2 | | | | | | | | | | 7.3 | | | | 8.0 | | | |
| 40 | 725 | *Bacillus* sp. | RS | MA | 8.1 | | | | | | | | | | | | | | | | | |
| 41 | | | | GYM | 6.2 | | | | | | | | | | 6.6 | | | | | | | |
| 42 | RBI15 | *Pseudoalteromonas* sp. | RS | GYM | 4.4 | | | | | | | | | | 4.4 | 7.1 | 6.3 | | 6.7 | | | |
| 43 | x018 | *Octadecabacter* sp. | RS | GYM | | | | | | | | | | | | | | | 8.8 | | | |
| 44 | xRASU | *Rheinheimera baltica* | RS | GYM | 7.6 | | | | | | | | | | 8.6 | | | | | | | |
| 45 | x005 | *Dokdonia* sp. | RS | MA | | | | | | | | | | | 4.7 | | | | | | | |
| 46 | 704 | *Bacillus* sp. | HLI | GYM | 6.9 | | | | | | | | | | 10.0 | | | | | | | |
| 47 | 742 | *Bacillus sp.* | HLI | MA | 5.9 | | | | | | | | | | | | | | | | | |
| 48 | | | | GYM | 6.3 | 9.0 | | | | | | | | | 2.7 | | 6.9 | | 7.5 | | | 6.2 |
| 49 | x015 | *Bacillus* sp. | HLI | MA | | | | | | | | | | | | | | | 2.2 | | | |
| 50 | | | | GYM | 7.5 | | | | | | | 2.1 | | | | | 6.9 | | | | | 2.1 |
| 51 | 237 | *Thalassospira lucentensis* | DLI | MA | 9.0 | | | | | | | | | | | | | | | | | |
| 52 | | | | GYM | 7.1 | | | | | | | | | | 6.4 | | | | 7.7 | | | |
| 53 | 436 | *Paenibacillus* sp. | DLI | MA | | | | | | | | | | | 5.3 | | | | | | | |
| 54 | | | | GYM | 7.2 | 7.4 | | | | | | | | | 3.8 | 6.7 | 7.1 | 7.2 | 5.3 | | | 5.4 |
| 55 | 706 | *Oceanisphaera sediminis* | DLI | GYM | 6.5 | | | | | | | | | | | | | | | | | |
| 56 | 332b | *Oceanobacillus* sp. | DLI | MA | 7.6 | | | | | | | | | | | | | | | | | |
| 57 | | | | GYM | 8.6 | | | | | | | | | | 8.7 | | | | | | | 7.4 |
| 58 | x014 | *Aquimarina* sp. | DLI | MA | 8.1 | | | | | | | | | | | | | | | | | |
| 59 | | | | GYM | | | | | | | | | | | 6.3 | | | | | | | |
| 60 | x019 | *Streptomyces* sp. | DLI | MA | 3.3 | | | | | | | | | | | | | | | 4.0 | 0.2 | |
| 61 | | | | GYM | 3.1 | | | 6.6 | 7.9 | 7.4 | 1.9 | 2.1 | 2.2 | 1.5 | 8.5 | | | | | 6.5 | 0.6 | |
| 62 | 334 | *Breoghania corrubendonensis* | RI | MA | | | | | | | | | | | | | | | 8.3 | | | |
| 63 | | | | GYM | 9.4 | | | | | | | | | | | | | | 4.9 | | | |
| 64 | x012 | *Shewanella* sp. | RI | MA | 6.9 | | | | | | | | | | | | | | | | | |
| 65 | | | | GYM | 7.1 | | | | | | | | | | 7.7 | | | | 9.7 | | | |
| 66 | x039 | *Hoeflea alexandrii* | RI | GYM | 7.0 | | | | | | | | | | 4.4 | | | | | | | |
| 67 | 738 | *Streptomyces* sp. | S | MA | 8.1 | | | | | | | | | | 8.9 | | | | | | | |
| 68 | | | | GYM | 6.6 | | | | | | | 4.6 | | | 7.4 | | | | | | | |
| 69 | 744 | *Streptomyces scopiformis* | S | MA | | | | | | | | | | | | | | | | 2.2 | 2.1 | |
| 70 | | | | GYM | | | | | | | | | | | | | | | | 1.3 | 2.8 | |
| 71 | xSBU1 | *Pseudoalteromonas ulvae* | S | GYM | 5.7 | | | | | | | | | | 7.1 | | | | 8.8 | | | |

**Supplementary Table S13.** The most active ($IC_{50} \leq 10$ µg/ml) fungal extracts selected for metabolome analysis. HLS: Healthy leaf surface, DLS: Decaying leaf surface, DLI: Decaying leaf inner tissue, RS: Root surface, W: Seawater reference, S: Sediment reference. Aquatic pathogens (positive control) La: *Leifsonia aquatica* (chloramphenicol), Lg: *Lactococcus garvieae* (Ampicillin) Ab: *Algicola bacteriolytica* (chloramphenicol), Sha: *Shewanella algae* (chloramphenicol), Pe: *Pseudoalteromonas elyacovii* (chloramphenicol), Vae: *Vibrio aestuarianus* (chloramphenicol), , Va: *Vibrio anguillarum* (chloramphenicol), Vch: *Vibrio cholerae* (chloramphenicol), Vf: *Vibrio fischeri* (chloramphenicol), Vh: *Vibrio harveyi* (chloramphenicol), Vi: *Vibrio ichthyoenteri* (chloramphenicol), Vp: *Vibrio parahaemolyticus* (chloramphenicol), Vv: *Vibrio vulnificus* (chloramphenicol). Fecal pathogens (positive control) Ecas*:* *Enterococcus casseliflavus* (ampicillin), Ef: *Enterococcus faecalis* (ampicillin), Efm: *Enterococcus faecium* (ampicillin), Eh: *Enterococcus hirae* (ampicillin). Human pathogen (positive control) MRSA: Methicillin resistant *Staphylococcus aureus* (chloramphenicol). Phytopathogens (positive control) Pi: *Phytophthora infestans* (cycloheximid), Mg: *Magnaphorte grisea* (nystatin), Rs: *Ralstonia solanacearum* (tetracycline), Xc: *Xanthomonas campestris* (chloramphenicol).

| | Strain | Identification | Source | Media | Aquatic test panel | | | | | | | | | | | | | Fecal test panel | | | | Human test panel | Plant pathogenic test panel | | | |
|---|---|---|---|---|---|---|---|---|---|---|---|---|---|---|---|---|---|---|---|---|---|---|---|---|---|---|
| | | | | | La | Lg | Ab | Sha | Pe | Vae | Va | Vch | Vf | Vh | Vi | Vp | Vv | Ecas | Ef | Efm | Eh | MRSA | Pi | Mg | Rs | Xc |
| | **Pos** | Positive control | | | 0.4 | 0.5 | 2.9 | 4.1 | 2.3 | 1.2 | 0.4 | 0.5 | 0.6 | 0.7 | 0.4 | 0.4 | 0.2 | 2.4 | 0.5 | 0.2 | 0.8 | 1.5 | 0.01 | 0.3 | 0.8 | 2.4 |
| 1 | 425a | *Parathyridaria* sp. | HLS | M34 | | | | | | | | | | | | | | | | | | 4.1 | | 4.5 | | |
| 2 | | | | PDA | | | | | | | | | | | | | | | 7.8 | 6.8 | | 2.1 | 5.1 | | | |
| 3 | 434 | *Parathyridaria* sp. | HLS | M34 | | | | | | | | | | | | | | | | | | 6.5 | | 2.9 | | |
| 4 | | | | PDA | | | | | | | | | | | | | | 9.1 | 6.8 | 9.2 | | 2.5 | | 8.3 | | |
| 5 | 233 | *Cladosporium halotolerans* | DLS | M34 | | | | | | | | | | | | | | | | | | | 2.2 | | | |
| 6 | | | | PDA | | | | | | | | | | | | | | | | | | | 0.7 | | | |
| 7 | 719 | *Fusarium* sp. | DLS | M34 | | | | | | | | | | | | | | | | 8.9 | | 5.8 | 1.2 | 9.4 | | |
| 8 | | | | PDA | | | | | | | | | | | | | | 3.7 | | 5.4 | | 3.9 | 2.6 | | | |
| 9 | 417 | *Cystofilobasidium bisporidii* | RS | M34 | 8.6 | | | | | | | | | | | | | | | | | | | | | |
| 10 | 720 | *Acrostalagmus luteoalbus* | DLI | M34 | 2.1 | 9.9 | 9.2 | | 3.8 | 8.7 | 4.1 | 5.0 | 4.8 | | | 4.6 | 3.5 | | 0.8 | 3.9 | 3.5 | 0.2 | 0.5 | | 8.7 | 3.2 |
| 11 | | | | PDA | 2.2 | 6.3 | | | 3.6 | 5.3 | 7.1 | 2.6 | 3.5 | 8.7 | | 4.2 | 3.7 | | 1.0 | 6.0 | 4.1 | 0.5 | 0.7 | | 9.1 | 4.1 |
| 12 | 407 | *Trichoderma viride* | S | M34 | | | | | | | | | | | | | | | | | | 7.0 | | | | |
| 13 | 809 | *Penicillium olsonii* | S | M34 | 9.9 | | | | | | | | | | | | | | | | | | | | | |
| 14 | | | | PDA | 2.4 | 3.2 | | | | | | | | | | | 8.5 | 1.9 | 1.7 | 0.9 | 1.8 | 1.0 | 0.7 | 0.2 | | |
| 15 | 813 | *Aureobasidium pullulans* | W | M34 | | | | | | | | | | | 5.2 | | | 2.8 | | | | | | | | 8.3 |
| 16 | | | | PDA | | | | | | | | | | | | | | | | | | | | | | 7.1 |
| 17 | 815 | *Sarocladium strictum* | W | M34 | | | | | | | | | | | | | | | | | | 8.3 | | | | |

**Supplementary Table S14.** Putative annotation of metabolites produced by the extracts of the most active bacteria isolated from eelgrass surfaces: i.e., healthy leaves (HLS), decaying leaves (DLS), roots (RS), plus sediment (S) and seawater (W) references, detected in two culture media.

| **Experiment. *m/z*** | **$t_R$ Min** | **Putative annotation** | **Molecular formula** | **Strain number, source bacterium, origin** | **Culture Media** | **Adduct** | **Chemical family** | **Reported bioactivity** | **Biological source** | **References** |
|---|---|---|---|---|---|---|---|---|---|---|
| 197.1290 | 0.6 | Cyclo(L-Pro-LVal) | $C_{10}H_{16}N_2O_2$ | 131 *Streptomyces* sp. HLS<br>137 *Rhizobium* sp. RS<br>150 *Stenotrophomonas* sp. HLS<br>202b *Pseudoalteromonas* sp. HLS<br>204 *Falsirhodobacter* sp. HLS<br>205 Alteromonadaceae sp. DLS<br>229 *Novosphingobium decolorationis* DLS<br>248 *Vibrio metschnikovii* HLS<br>250 *Alteromonas stellipolaris* HLS<br>303 *Gallaecimonas pentaromativorans* HLS<br>325 *Gallaecimonas pentaromativorans* HLS<br>609 *Agarivorans* sp.HLS<br>610a *Flavobacterium jumunjinense* HLS<br>625 *Flavobacterium jumunjinense* HLS<br>629 *Cobetia* sp. HLS<br>630 Alteromonadaceae sp. RS<br>632 *Alteromonas stellipolaris* DLS<br>640 *Neptunomoas phycophila* HLS<br>713b *Proteus* sp.HLS<br>716 *Pantoea* sp.HLS<br>725 *Bacillus* sp.RS<br>738 *Streptomyces* sp. S<br>741 *Psychrobacter* sp. HLS<br>744 *Streptomyces* sp. S<br>DAI04 *Pseudoalteromonas tetraodonis* DLS<br>RBI15 *Pseudoalteromonas* sp. RS<br>x001 *Cobetia* sp. HLS<br>x007 *Alkalihalobacillus hwajinpoensis* HLS<br>x018 *Octadecabacter* sp. RS<br>x025 *Streptomyces griseorubens* DLS<br>x028 *Pseudomonas* sp. HLS | GYM, MA | $[M+H]^+$ | Diketopiperazine | Bioactivity against marine pathogenic bacteria and MRSA. | Various bacterial and fungal species | Borthwick, 2012;<br>Qi et al., 2009;<br>Alshaibani et al., 2017 |
| 211.1451 | 0.74 | Cyclo(L-Pro-L-Leu) | $C_{11}H_{18}N_2O_2$ | 131 *Streptomyces* sp. HLS<br>137 *Rhizobium* sp. RS<br>150 *Stenotrophomonas* sp. HLS<br>202b *Pseudoalteromonas* sp. HLS<br>204 *Falsirhodobacter* sp. HLS<br>205 Alteromonadaceae sp. DLS<br>229 *Novosphingobium decolorationis* DLS<br>248 *Vibrio metschnikovii* HLS<br>250 *Alteromonas stellipolaris* HLS<br>303 *Gallaecimonas pentaromativorans* HLS<br>323 *Flavobacterium ponti* HLS<br>325 *Gallaecimonas pentaromativorans* HLS<br>609 *Agarivorans* sp.HLS<br>610a *Flavobacterium jumunjinense* HLS<br>616 *Sulfitobacter pontiacus* DLS<br>625 *Flavobacterium jumunjinense* HLS<br>629 *Cobetia* sp. HLS<br>630 Alteromonadaceae sp. RS<br>632 *Alteromonas stellipolaris* DLS<br>640 *Neptunomoas phycophila* HLS | GYM, MA | $[M+H]^+$ | Diketopiperazine | Potent antibiotic against VRE strains, incl. *E. faecium* and *E. faecalis,* antiphytopatho genic vs *M. oryzae* | Various bacterial and fungal species including *Streptomyces* spp. | Rhee, 2002;<br>Rhee, 2003 |

| Experiment. *m/z* | $t_R$ Min | Putative annotation | Molecular formula | Strain number, source bacterium, origin | Culture Media | Adduct | Chemical family | Reported bioactivity | Biological source | References |
|---|---|---|---|---|---|---|---|---|---|---|
| | | | | 713b *Proteus* sp.HLS<br>716 *Pantoea* sp.HLS<br>725 *Bacillus* sp.RS<br>738 *Streptomyces* sp. S<br>741 *Psychrobacter* sp. HLS<br>744 *Streptomyces* sp. S<br>DAI04 *Pseudoalteromonas tetraodonis* DLS<br>RBI15 *Pseudoalteromonas* sp. RS<br>x001 *Cobetia* sp. HLS<br>x005 *Dokdonia* sp. RS<br>x007 *Alkalihalobacillus hwajinpoensis* HLS<br>x018 *Octadecabacter* sp. RS<br>x025 *Streptomyces griseorubens* DLS<br>x028 *Pseudomonas* sp. HLS<br>xRASU *Rheinheimera baltica* RS<br>xSBU1 *Pseudoalteromonas ulvae* S | | | | | | |
| 247.1448 | 0.9 | Cyclo(L-Val-L-Phe) | $C_{14}H_{18}N_2O_2$ | 616 *Sulfitobacter pontiacus* DLS<br>640 *Neptunomoas phycophila* HLS<br>725 *Bacillus* sp.RS<br>741 *Psychrobacter* sp. HLS<br>RBI15 *Pseudoalteromonas* sp. RS<br>x001 *Cobetia* sp. HLS<br>x028 *Pseudomonas* sp. HLS | GYM, MA | $[M+H]^+$ | Diketopiperazine | Quorum sensing regulator | *Pseudoalteromonas* sp. isolated from the marine sponge *Hymeniacidon perleve* | Guo et al.,2011 |
| 284.1401 | 0.82 | Cyclo(L-Trp-L-Pro) | $C_{16}H_{17}N_3O_2$ | 131 *Streptomyces* sp. HLS<br>137 *Rhizobium* sp. RS<br>150 *Stenotrophomonas* sp. HLS<br>202b *Pseudoalteromonas* sp. HLS<br>204 *Falsirhodobacter* sp. HLS<br>205 Alteromonadaceae sp. DLS<br>248 *Vibrio metschnikovii* HLS<br>609 *Agarivorans* sp.HLS<br>610a *Flavobacterium jumunjinense* HLS<br>616 *Sulfitobacter pontiacus* DLS<br>625 *Flavobacterium jumunjinense* HLS<br>629 *Cobetia* sp. HLS<br>630 Alteromonadaceae sp. RS<br>632 *Alteromonas stellipolaris* DLS<br>640 *Neptunomoas phycophila* HLS<br>713b *Proteus* sp.HLS<br>716 *Pantoea* sp.HLS<br>725 *Bacillus* sp.RS<br>738 *Streptomyces* sp. S<br>741 *Psychrobacter* sp. HLS<br>DAI04 *Pseudoalteromonas tetraodonis* DLS<br>RBI15 *Pseudoalteromonas* sp. RS<br>x001 *Cobetia* sp. HLS<br>x005 *Dokdonia* sp. RS<br>x007 *Alkalihalobacillus hwajinpoensis* HLS<br>x028 *Pseudomonas* sp. HLS<br>xSBU1 *Pseudoalteromonas ulvae* S | GYM, MA | $[M+H]^+$ | Diketopiperazine | Bioactivity against Gram-positive bacteria incl. *S. aureus*, antifungal against human pathogenic fungi | Various bacterial and fungal species including *Streptomyces* spp. | Borthwick, 2012; Ben Ameur Mehdi et al., 2009; Nishanth Kumar et al.,2014 |
| 347.0919 | 1.95 | Wailupemycin G | $C_{21}H_{14}O_5$ | 738 *Streptomyces* sp. S | GYM, MA | $[M+H]^+$ | Polyketide | Bacteriostatic against Gram-positive & -negative bacteria | Marine-derived *Streptomyces* sp. | Xiang et al., 2002; Sitachitta et al.,1996 |

| Experiment. *m/z* | $t_R$ Min | Putative annotation | Molecular formula | Strain number, source bacterium, origin | Culture Media | Adduct | Chemical family | Reported bioactivity | Biological source | References |
|---|---|---|---|---|---|---|---|---|---|---|
| 365.1020 | 1.4 | Wailupemycin F | $C_{21}H_{16}O_6$ | 738 *Streptomyces* sp. S | GYM | $[M+H]^+$ | Polyketide | Bacteriostatic against Gram-positive & -negative bacteria | Marine-derived *Streptomyces* sp. | Xiang et al., 2002; Sitachitta et al.,1996 |
| 387.2379 | 2.25 | Nonactyl nonactate | $C_{20}H_{34}O_7$ | 131 *Streptomyces* sp. HLS | GYM | $[M+H]^+$ | Polyketide | Not reported | Soil-derived *Streptomyces globisporus* | Řezanka et al., 2004 |
| 397.1759 | 4.11 | Streptophenazine E | $C_{22}H_{24}N_2O_5$ | 131 *Streptomyces* sp. HLS | MA | $[M+H]^+$ | Phenazine alkaloid | Bioactive against Gram-positive bacteria | *Streptomyces* sp. isolated from the marine sponge *Halichondria panicea* | Mitova et al.,2008; Kunz et al., 2014 |
| 400.1867 | 0.6 | Bioxalomycin Beta 2 | $C_{21}H_{25}N_3O_5$ | x025 *Streptomyces griseorubens* DLS | GYM | $[M+H]^+$ | Tetrahydroisoquinoline alkaloid | Broad-spectrum antibiotic vs. Gram-positive and -negative bacteria, incl. MRSA, *E. faecium;* cytotoxic | Marine sediment derived *Streptomyces* sp. | Bernan et al.,1994; Zaccardi et al., 1994; Scott et al., 2002 |
| 401.2538 | 2.8 | Bonactin | $C_{21}H_{36}O_7$ | 131 *Streptomyces* sp. HLS | GYM, MA | $[M+H]^+$ | Polyketide | Antimicrobial vs Gram-positive (incl. *S.aureus*) and Gram-negative bacteria, antifungal antiphytopathogenic vs *Magnaporthe oryzae* | Marine sediment-derived *Streptomyces* sp. | Schumacher et al., 2003; Rabby et al., 2022 |
| 409.2199 | 2.26 | Nonactyl nonactate | $C_{20}H_{34}O_7$ | 131 *Streptomyces* sp. HLS | GYM, MA | $[M+Na]^+$ | Polyketide | Not reported | Soil-derived *Streptomyces globisporus* | Řezanka et al.,2004 |
| 411.1918 | 4.5 | Streptophenazine C | $C_{23}H_{26}N_2O_5$ | 131 *Streptomyces* sp. HLS | MA | $[M+H]^+$ | Phenazine alkaloid | Bioactivite against Gram-positive bacteria | *Streptomyces* sp. Isolated from the marine sponge *Halichondria panicea* | Mitova et al., 2008; Kunz et al., 2014 |
| 415.2694 | 3.31 | Homononactyl Homononactate | $C_{22}H_{38}O_7$ | 131 *Streptomyces* sp. HLS | GYM, MA | $[M+H]^+$ | Polyketide | Weak antibacterial activity | *Bacillus pumilus* isolated from a plant sample (*Breynia fruticosa*) | Han et al., 2014: Huang et al., 2015 |
| 418.1969 | 0.54 | Naphthyridinomycin | $C_{21}H_{27}N_3O_6$ | x025 *Streptomyces griseorubens* DLS | GYM | $[M+H]^+$ | Tetrahydroisoquinoline alkaloid | Broad-spectrum antibiotic vs Gram-positive and -negative bacteria, incl. *S. aureus* and *E. faecium,* cytotoxic | Soil-derived *Streptomyces lusitanus* | Kluepfel et al., 1975; Zaccardi et al., 1994; Scott et al., 2002 |
| 423.1921 | 5.81 | Oxo-streptophenazine A | $C_{24}H_{26}N_2O_5$ | 131 *Streptomyces* sp. HLS | MA | $[M+H]^+$ | Phenazine alkaloid | Weak or no antimicrobial activity | Marine-derived *Streptomyces* sp. | Bauman et al., 2019 |

| Experiment. *m/z* | $t_R$ Min | Putative annotation | Molecular formula | Strain number, source bacterium, origin | Culture Media | Adduct | Chemical family | Reported bioactivity | Biological source | References |
|---|---|---|---|---|---|---|---|---|---|---|
| 423.2363 | 2.8 | Bonactin | $C_{21}H_{36}O_7$ | 131 *Streptomyces* sp. HLS | GYM, MA | $[M+Na]^+$ | Polyketide | Antibiotic vs Gram-positive (incl *S. aureus*) and -negative bacteria, antifungal, anti-phytopathogenic vs *M. oryzae* | Marine sediment-derived *Streptomyces* sp. | Schumacher et al., 2003; Rabby et al., 2022 |
| 425.2075 | 4.98 | Streptophenazine B | $C_{24}H_{28}N_2O_5$ | 131 *Streptomyces* sp. HLS | GYM, MA | $[M+H]^+$ | Phenazine alkaloid | Bioactivity against Gram-positive bacteria, incl. MRSA | *Streptomyces* sp. isolated from sponge *Halichondria panicea* | Mitova et al., 2008; Kunz et al., 2014; Liang et al.,2017 |
| 429.1186 | 0.86 | 8-deoxyenterocin | $C_{22}H_{20}O_9$ | 738 *Streptomyces* sp. S | GYM, MA | $[M+H]^+$ | Polyketide | Not reported | Terrestrial-derived *Streptomyces* sp. | Sitachitta et al.,1996 |
| 437.2078 | 6.41 | Oxo-streptophenazine G | $C_{25}H_{28}N_2O_5$ | 131 *Streptomyces* sp. HLS | MA | $[M+H]^+$ | Phenazine alkaloid | Weak or absent antimicrobial activity | marine-derived *Streptomyces* sp. | Bauman et al., 2019 |
| 437.2522 | 3.36 | Homononactyl Homononactate | $C_{22}H_{38}O_7$ | 131 *Streptomyces* sp. HLS | GYM, MA | $[M+Na]^+$ | Polyketide | Weak antibacterial activity | *Bacillus pumilus* isolated from a plant sample *(Breynia fruticosa)* | Han et al.,2014: Huang et al., 2015 |
| 439.2227 | 5.66 | Streptophenazine G | $C_{25}H_{30}N_2O_5$ | 131 *Streptomyces* sp. HLS | GYM, MA | $[M+H]^+$ | Phenazine alkaloid | Active against Gram-positive bacteria | *Streptomyces* sp. isolated from sponge *Halichondria panicea* | Mitova et al., 2008; Kunz et al., 2014 |
| 441.2026 | 2.11 | Streptophenazine H | $C_{24}H_{28}N_2O_6$ | 131 *Streptomyces* sp. HLS | MA | $[M+H]^+$ | Phenazine alkaloid | Active against Gram-positive bacteria | *Streptomyces* sp. isolated from sponge *Halichondria panicea* | Mitova et al., 2008; Kunz et al., 2014 |
| 445.1130 | 0.7 | Enterocin | $C_{22}H_{20}O_{10}$ | 738 *Streptomyces* sp. S | GYM, MA | $[M+H]^+$ | Polyketide | Active against Gram-positive and Gram-negative bacteria, food pathogens | *Streptomyces* sp. | Miyairi et al., 1976 |
| 451.2669 | 3.89 | Ethyl Homononactyl Nonactate | $C_{23}H_{40}O_7$ | 131 *Streptomyces* sp. HLS | MA | $[M+Na]^+$ | Polyketide | Not reported | *Bacillus pumilus* isolated from a plant sample *(Breynia fruticosa)* | Han et al., 2014 |
| 455.2179 | 2.58 | Streptophenazine I | $C_{25}H_{30}N_2O_6$ | 131 *Streptomyces* sp. HLS | MA | $[M+H]^+$ | Phenazine alkaloid | Inhibitory activity against the enzyme phosphodiesterase (PDE 4B) | *Streptomyces* sp. isolated from the sponge *Halichondria panicea* | Kunz et al., 2014 |
| 467.0941 | 0.7 | Enterocin | $C_{22}H_{20}O_{10}$ | 738 *Streptomyces* sp. S | GYM, MA | $[M+Na]^+$ | Polyketide | Active against food pathogens | *Streptomyces* sp. | Miyairi et al., 1976 |
| 627.3869 | 7.54 | Bafilomycin D | $C_{35}H_{56}O_8$ | 744 *Streptomyces* sp. S | GYM | $[M+Na]^+$ | Bafilomycin | Active vs Gram-positive bacteria, fungi, yeast, anti-neoplastic, immunosuppressive, | Soil-derived *Streptomyces* sp. | Kretschmer et al., 1985; Zhang et al., 2011; Xu et al.,2013 |

| Experiment. *m/z* | $t_R$ Min | Putative annotation | Molecular formula | Strain number, source bacterium, origin | Culture Media | Adduct | Chemical family | Reported bioactivity | Biological source | References |
|---|---|---|---|---|---|---|---|---|---|---|
| | | | | | | | | antiphytopathogenic | | |
| 641.4033 | 7.19 | 17,18-Dehydro-19,21-di-O-methyl-24-demethyl-bafilomycin A1 | $C_{36}H_{58}O_8$ | 744 *Streptomyces* sp. S | GYM | $[M+Na]^+$ | Bafilomycin | Active vs Gram-positive bacteria, fungi, yeast, anti-neoplastic, immunosuppressive, antiphytopathogenic | *Streptomyces* sp. isolated from the callus of *Maytenus hookeri* | Li et al., 2010; Zhang et al., 2011; |
| 645.3982 | 6.35 | Bafilomycin A1 | $C_{35}H_{58}O_9$ | 744 *Streptomyces* sp. S | GYM, MA | $[M+Na]^+$ | Bafilomycin | Active vs Gram-positive bacteria, fungi, yeast, anti-neoplastic, immunosuppressive, antiphytopathogenic | Soil-derived *Streptomyces griseus* | Werner et al.,1984; Zhang et al., 2011; Xu et al., 2013 |
| 659.4135 | 7.88 | Bafilomycin A2 | $C_{36}H_{60}O_9$ | 744 *Streptomyces* sp. S | GYM, MA | $[M+Na]^+$ | Bafilomycin | Active vs Gram-positive bacteria, fungi, yeast, anti-neoplastic, immunsuppressive | Soil-derived *Streptomyces griseus* | Werner et al., 1984; Zhang et al., 2011 |
| 751.4638 | 8.72 | Monactin | $C_{41}H_{66}O_{12}$ | 131 *Streptomyces* sp. HLS | GYM, MA | $[M+H]^+$ | Macrotetrolide | Antibacterial, antifungal, insecticidial, antiproliferative, inhibits zoosporogenesis & motility of peronosporomycete zoospores | Soil-derived *Streptomyces* sp. | Beck et al., 1962; Zizka, 1998; Shishlyannikova et al., 2017; Islam et al., 2016 |
| 757.4503 | 6.49 | Dermostatin B | $C_{41}H_{66}O_{11}$ | 744 *Streptomyces* sp. S | MA | $[M+Na]^+$ | Polyene macrolide | Antifungal | Soil-derived *Streptomyces virdigreseus* | Thirumalachar et al., 1962 |
| 759.4294 | 8.39 | Nonactin | $C_{40}H_{64}O_{12}$ | 131 *Streptomyces* sp. HLS | GYM | $[M+Na]^+$ | Macrotetrolide | Antibacterial, antifungal, antiproliferative, inhibitseffects on zoosporogenesis and motility of peronosporomycete zoospores | Soil-derived *Streptomyces* sp. | Corbaz et al., 1955; Kusche et al., 2009; Zizka, 1998; Shishlyannikova et al., 2017; Islam et al., 2016 |
| 773.4458 | 8.72 | Monactin | $C_{41}H_{66}O_{12}$ | 131 *Streptomyces* sp. HLS | GYM, MA | $[M+Na]^+$ | Macrotetrolide | Antibacterial, antifungal, | Soil-derived *Streptomyces* sp. | Beck et al., 1962; Zizka 1998; |

| Experiment. *m/z* | $t_R$ Min | Putative annotation | Molecular formula | Strain number, source bacterium, origin | Culture Media | Adduct | Chemical family | Reported bioactivity | Biological source | References |
|---|---|---|---|---|---|---|---|---|---|---|
| | | | | | | | | insecticidial antiproliferative, inhibits zoosporogenesis & motility of peronosporomycete zoospores | | Islam et al., 2016; Shishlyannikova et al., 2017 |
| 773.4457 | 7.22 | Pn00053 | $C_{41}H_{66}O_{12}$ | 744 *Streptomyces* sp. S | GYM, MA | $[M+Na]^+$ | Polyene macrolide | Broad spectrum antifungal activity | Soil-derived *Streptomyces* sp. | Vartak et al., 2014 |
| 783.4906 | 6.71 | Seco-dinactin | $C_{42}H_{70}O_{13}$ | 131 *Streptomyces* sp. HLS | MA | $[M+H]^+$ | Polyketide | Not reported | Soil-derived *Streptomyces griseus* | Xie et al., 2014 |
| 787.4627 | 9.01 | Dinactin | $C_{42}H_{68}O_{12}$ | 131 *Streptomyces* sp. HLS | GYM, MA | $[M+Na]^+$ | Macrotetrolide | Antibacterial, antifungal, insecticidial antiproliferative, inhibits zoosporogenesis & motility of peronosporomycete zoospores, antifungal vs phytopathogens. | Soil-derived *Streptomyces* sp. | Beck et al.,1962; Zizka 1998; Islam et al.,2016; Shishlyannikova et al., 2017; Liu et al., 2019; Zhang et al.,2020 |
| 801.4775 | 9.33 | Trinactin | $C_{43}H_{70}O_{12}$ | 131 *Streptomyces* sp. HLS | GYM, MA | $[M+Na]^+$ | Macrotetrolide | Antibacterial, antifungal, insecticidial antiproliferative, inhibits zoosporogenesis & motility of peronosporomycete zoospores | Soil-derived *Streptomyces* sp. | Beck et al., 1962; Zizka, 1998; Islam et al., 2016; Shishlyannikova et al., 2017 |
| 815.4927 | 9.63 | Tetranactin | $C_{44}H_{72}O_{12}$ | 131 *Streptomyces* sp. HLS | GYM, MA | $[M+Na]^+$ | Macrotetrolide | Antibacterial, antifungal, insecticidial antiproliferative, antifungal vs phytopathogens | Soil-derived *Streptomyces* sp. | Smith, 1975; Zizka 1998; Shishlyannikova et al., 2017; Liu et al., 2019 |
| 829.5083 | 10.02 | Macrotetrolide C | $C_{45}H_{74}O_{12}$ | 131 *Streptomyces* sp. HLS | GYM, MA | $[M+Na]^+$ | Macrotetrolide | Not reported | Soil-derived *Streptomyces* sp. | Smith, 1975; Zizka 1998 |

**Supplementary Table S15.** Putative annotation of metabolites produced by the extracts of the most active bacteria isolated from the inner tissues of eelgrass, i.e., healthy leaves (HLI), decaying leaves (DLI) and roots (RI) detected in two culture media.

| Experiment. m/z | $t_R$ Min | Putative annotation | Molecular formula | Strain number, Source bacterium, origin | Culture Media | Adduct | Chemical family | Reported bioactivity | Biological source | References |
|---|---|---|---|---|---|---|---|---|---|---|
| 197.1289 | 0.6 | Cyclo(L-Pro-L-Val) | $C_{10}H_{16}N_2O_2$ | 237 *Thalassospira lucentensis* DLI<br>332b *Oceanobacillus* sp. DLI<br>334 *Breoghania corrubendonensis* RI<br>436 *Paenibacillus* sp. DLI<br>706 *Oceanisphaera sediminis* DLI<br>x012 *Shewanella* sp. RI<br>x015 *Bacillus* sp. HLI<br>x019 *Streptomyces* sp. DLI | MA, GYM | $[M+H]^+$ | Diketopiperazine | Bioactivity against marine pathogenic bacteria and MRSA. | Various bacterial and fungal species | Borthwick, 2012;<br>Qi et al., 2009;<br>Alshaibani et al., 2017 |
| 211.1445 | 0.72 | Cyclo(L-Pro-L-Leu) | $C_{11}H_{18}N_2O_2$ | 237 *Thalassospira lucentensis* DLI<br>334 *Breoghania corrubendonensis* RI<br>436 *Paenibacillus* sp. DLI<br>704 *Bacillus* sp. HLI<br>742 *Bacillus* sp. HLI<br>x012 *Shewanella* sp. RI<br>x014 *Aquimarina* sp. DLI<br>x015 *Bacillus* sp. HLI<br>x019 *Streptomyces* sp. DLI<br>x039 *Hoeflea alexandrii* RI | MA, GYM | $[M+H]^+$ | Diketopiperazine | Potent activity against twelve VRE strains, including *E. faecium* and *E. faecalis*, antitumor, anti-phytopathogenic vs *M. oryzae* | Various bacterial and fungal species including *Streptomy ces* spp. | Rhee, 2002;<br>Rhee, 2003 |
| 247.1449 | 0.89 | Cyclo-(L-Val-L-Phe) | $C_{14}H_{18}N_2O_2$ | 436 *Paenibacillus* sp. DLI<br>706 *Oceanisphaera sediminis* DLI | MA, GYM | $[M+H]^+$ | Diketopiperazine | Quorum sensing regulator | *Pseudoalteromonas sp. i*solated from the marine sponge *Hymeniacidon perleve* | Guo et al.,2011 |
| 284.1397 | 0.70 | Cyclo(L-Trp-L-Pro) | $C_{16}H_{17}N_3O_2$ | 237 *Thalassospira lucentensis* DLI<br>704 *Bacillus* sp. HLI<br>x014 *Aquimarina* sp. DLI<br>x015 *Bacillus* sp. HLI | MA, GYM | $[M+H]^+$ | Diketopiperazine | Bioactivity against Gram-positive bacteria incl. *S. aureus*, antifungal against human pathogenic fungi | Various bacterial and fungal species including *Streptomyces* spp. | Borthwick, 2012;<br>Ben Ameur Mehdi et al., 2009;<br>Nishanth Kumar et al.,2014 |
| 429.1182 | 0.86 | 8-Deoxyenterocin | $C_{22}H_{20}O_9$ | x019 *Streptomyces* sp. DLI | MA | $[M+H]^+$ | Polyketide | Not reported | Terrestrial-derived *Streptomyces* sp. | Sitachitta et al.,1996 |
| 445.1130 | 0.71 | Enterocin | $C_{22}H_{20}O_{10}$ | x019 *Streptomyces* sp. DLI | GYM | $[M+H]^+$ | Polyketide | Broad-spectrum bactericidal, bacteriostatic activity | Terrestrial and marine *Streptomyces* sp. | Miyairi et al., 1976 |
| 449.8101 | 2.61 | Surugamide E | $C_{47}H_{79}N_9O_8$ | x019 *Streptomyces* sp. DLI | MA, GYM | $[M+2H]^{++}$ | Cyclic peptide | Anticancer activity | Deep-sea sediment-derived *Streptomyces* sp. | Takada et al., 2013;<br>Almeida et al.,2019 |
| 451.1002 | 0.86 | 8-Desoxyenterocin | $C_{22}H_{20}O_9$ | x019 *Streptomyces* sp. DLI | MA | $[M+Na]^+$ | Polyketide | Not reported | Terrestrial-derived *Streptomyces* sp. | Sitachitta et al.,1996 |
| 456.8173 | 2.61 | Surugamide A | $C_{48}H_{81}N_9O_8$ | x019 *Streptomyces* sp. DLI | MA, GYM | $[M+2H]^{++}$ | Cyclic peptide | Anticancer activity | Deep-sea sediment -derived *Streptomyces* sp. | Takada et al., 2013;<br>Almeida et al., 2019 |
| 479.2906 | 3.85 | Ikarugamycin | $C_{29}H_{38}N_2O_4$ | x019 *Streptomyces* sp. DLI | GYM | $[M+H]^+$ | Macrocyclic tetramic acid | Moderate activities against Gram-positive bacteria and fungi | Soil-derived *Streptomyces* sp. | Jomon et al., 1972 |
| 495.2857 | 4.44 | Ikarugamycin epoxide | $C_{29}H_{38}N_2O_5$ | x019 *Streptomyces* sp. DLI | GYM | $[M+H]^+$ | Macrocyclic tetramic acid | Moderate activities against Gram-positive bacteria and fungi | Soil-derived *Streptomyces* sp. | Bertasso et al., 2003 |
| 507.2703 | 6.54 | Deformylated antimycin A2a | $C_{26}H_{38}N_2O_8$ | x019 *Streptomyces* sp. DLI | MA | $[M+H]^+$ | Antimycin | Antifungal agent | Marine sediment-derived *Streptomyces* sp. | Zhang et al., 2017;<br>Belakhov et al., 2018;<br>Paul et al., 2022 |

| Experiment. m/z | $t_R$ Min | Putative annotation | Molecular formula | Strain number, Source bacterium, origin | Culture Media | Adduct | Chemical family | Reported bioactivity | Biological source | References |
|---|---|---|---|---|---|---|---|---|---|---|
| 511.2804 | 3.94 | Maltophilin | $C_{29}H_{38}N_2O_6$ | x019 *Streptomyces* sp. DLI | MA, GYM | $[M+H]^+$ | Macrocyclic tetramic acid | Broad spectrum fungicide; anti phytopathogenic vs *Plasmopara viticola* | *Stenotrophomonas maltophilia* from the rhizosphere of oilseed rape | Jakobi et al.,1996 |
| 513.2968 | 3.36 | Dihydromaltophilin | $C_{29}H_{40}N_2O_6$ | x019 *Streptomyces* sp. DLI | MA, GYM | $[M+H]^+$ | Macrocyclic tetramic acid | Not reported | Soil-derived *Streptomyces* sp. | Graupner et al., 1997 |
| 521.2855 | 6.88 | Deformylated antimycin A1a | $C_{27}H_{40}N_2O_8$ | x019 *Streptomyces* sp. DLI | MA | $[M+H]^+$ | Antimycin | Antifungal agent | Marine sediment-derived *Streptomyces* sp. | Zhang et al., 2017; Belakhov et al., 2018; Paul et al., 2022 |
| 535.2653 | 6.19 | Antimycin A2 | $C_{27}H_{38}N_2O_9$ | x019 *Streptomyces* sp. DLI | MA | $[M+H]^+$ | Antimycin | Antifungal agent | Soil-derived *Streptomyces* sp. | Barrow et al.,1997; Belakhov et al., 2018; Paul et al., 2022 |
| 549.2814 | 6.64 | Antimycin A1 | $C_{28}H_{40}N_2O_9$ | x019 *Streptomyces* sp. DLI | MA | $[M+H]^+$ | Antimycin | Antifungal agent | Soil-derived *Streptomyces* sp. | Dunshee et al., 1949 |
| 563.2969 | 7.04 | Antimycin A15 | $C_{29}H_{42}N_2O_9$ | x019 *Streptomyces* sp. DLI | MA | $[M+H]^+$ | Antimycin | Antifungal agent | Soil-derived *Streptomyces* sp. | Hosotani et al.,2005; Belakhov et al., 2018; Paul et al., 2022 |
| 577.3124 | 7.42 | Antimycin A16 | $C_{30}H_{44}N_2O_9$ | x019 *Streptomyces* sp. DLI | MA | $[M+H]^+$ | Antimycin | Antifungal agent | Soil-derived *Streptomyces* sp. | Hosotani et al.,2005; Belakhov et al., 2018; Paul et al., 2022 |
| 804.5026 | 3.81 | Koranimine | $C_{44}H_{65}N_7O_7$ | 742 *Bacillus* sp. HLI | MA, GYM | $[M+H]^+$ | Cyclic peptide | Potent nematocidal activity | Terrestrial-derived *Bacillus* sp. | Evans et al., 2011; Montecillo et al., 2022 |
| 822.5129 | 3.42 | Koranimine B | $C_{44}H_{67}N_7O_8$ | 742 *Bacillus* sp. HLI | MA | $[M+H]^+$ | Cyclic peptide | Not reported | terrestrial-derived *Bacillus* sp. | Malins et al., 2017 |
| 898.6139 | 2.61 | Surugamide E | $C_{47}H_{79}N_9O_8$ | x019 *Streptomyces* sp. DLI | MA, GYM | $[M+H]^+$ | Cyclic peptide | Anticancer activity | Deep-sea sediment -derived *Streptomyces* sp. | Takada et al., 2013; Almeida et al., 2019 |
| 912.6287 | 2.61 | Surugamide A | $C_{48}H_{81}N_9O_8$ | x019 *Streptomyces* sp. DLI | MA, GYM | $[M+H]^+$ | Cyclic peptide | Anticancer activity | Deep-sea sediment -derived *Streptomyces* sp. | Takada et al., 2013; Almeida et al., 2019 |
| 980.6286 | 7.08 | Surfactin $C_{-11}$ | $C_{49}H_{85}N_7O_{13}$ | x015 *Bacillus* sp. HLI | MA, GYM | $[M+H]^+$ | Lipopeptide | Antimicrobial vs Gram-positive (*Staphylococcus*, *Streptococcus*, *Enterococcus*), Gram-negative (*Flexibacter*, *Vibrio*, *Escherichia*, *Salmonella*) bacteria; anti-phytopathogenic vs *Xanthomonas campestris* | Soil-derived *Bacillus subtilis* | Fei et al.,2020; Bartal et al., 2023; Kim et al., 2009 |
| 994.6445 | 7.39 | [Val7]-Surfactin | $C_{50}H_{87}N_7O_{13}$ | x015 *Bacillus* sp. HLI | MA, GYM | $[M+H]^+$ | Lipopeptide | Antimicrobial vs Gram-positive (*Staphylococcus*, *Streptococcus*, and *Enterococcus*), Gram-negative (*Flexibacter*, *Vibrio*, *Escherichia*, and *Salmonella*) bacteria; antiphytopathogenic vs *Xanthomonas campestris* | Terrestrial-derived *Bacillus subtilis* | Peypoux et al.,1991; Kim et al.,2009; Bartal et al., 2023 |
| 1008.6600 | 7.84 | Surfactin A | $C_{51}H_{89}N_7O_{13}$ | x015 *Bacillus* sp. HLI | MA, GYM | $[M+H]^+$ | Lipopeptide | Antimicrobial vs Gram-positive | Several strains of terrestrial-derived | Arima et al, 1968; Bartal et al., 2023; |

| Experiment. m/z | $t_R$ Min | Putative annotation | Molecular formula | Strain number, Source bacterium, origin | Culture Media | Adduct | Chemical family | Reported bioactivity | Biological source | References |
|---|---|---|---|---|---|---|---|---|---|---|
| | | | | | | | | (*Staphylococcus*, *Streptococcus*, and *Enterococcus*), Gram-negative (*Flexibacter*, *Vibrio*, *Escherichia*, and *Salmonella*) bacteria; antiphytopathogenic vs *Xanthomonas campestris* | *Bacillus subtilis* | Kim et al., 2009 |
| 1022.6779 | 7.81 | Surfactin B | $C_{52}H_{91}N_7O_{13}$ | x015 *Bacillus* sp. HLI | MA, GYM | $[M+H]^+$ | Lipopeptide | Antimicrobial vs Gram-positive (*Staphylococcus*, *Streptococcus*, and *Enterococcus*), Gram-negative (*Flexibacter*, *Vibrio*, *Escherichia*, and *Salmonella*) bacteria; antiphytopathogenic vs *Xanthomonas campestris* | Several strains of terrestrial-derived *Bacillus subtilis* | Oka et al., 1993; Bartal et al., 2023, Kim et al.,2009 |
| 1025.6843 | 7.81 | Surfactin A | $C_{51}H_{89}N_7O_{13}$ | x015 *Bacillus* sp. HLI | MA, GYM | $[M+NH_4]^+$ | Lipopeptide | Antimicrobial vs Gram-positive (*Staphylococcus*, *Streptococcus*, and *Enterococcus*), Gram-negative (*Flexibacter*, *Vibrio*, *Escherichia*, and *Salmonella*) bacteria; antiphytopathogenic vs *Xanthomonas campestris* | Several strains of terrestrial-derived *Bacillus subtilis* | Arima et al.,1968; Bartal et al., 2023; Kim et al.,2009 |
| 1030.6418 | 7.85 | Surfactin A | $C_{51}H_{89}N_7O_{13}$ | x015 *Bacillus* sp. HLI | MA, GYM | $[M+Na]^+$ | Lipopeptide | Antimicrobial vs Gram-positive (*Staphylococcus*, *Streptococcus*, and *Enterococcus*), Gram-negative (*Flexibacter*, *Vibrio*, *Escherichia*, and *Salmonella*) bacteria; antiphytopathogenic vs *Xanthomonas campestris* | Several strains of terrestrial-derived *Bacillus subtilis* | Oka et al., 1993; Bartal et al., 2023, Kim et al., 2009 |
| 1036.6927 | 8.09 | Surfactin C | $C_{53}H_{93}N_7O_{13}$ | x015 *Bacillus* sp. HLI | MA, GYM | $[M+H]^+$ | Lipopeptide | Antimicrobial vs Gram-positive (*Staphylococcus*, *Streptococcus*, and *Enterococcus*), Gram-negative (*Flexibacter*, *Vibrio*, *Escherichia*, and *Salmonella*) bacteria; antiphytopathogenic | Several strains of terrestrial-derived *Bacillus subtilis* | Oka et al., 1993; Bartal et al., 2023; Kim et al., 2009 |

| Experiment. m/z | $t_R$ Min | Putative annotation | Molecular formula | Strain number, Source bacterium, origin | Culture Media | Adduct | Chemical family | Reported bioactivity | Biological source | References |
|---|---|---|---|---|---|---|---|---|---|---|
| | | | | | | | | vs *Xanthomonas campestris* | | |
| 1039.7003 | 8.08 | Surfactin B | $C_{52}H_{91}N_7O_{13}$ | x015 *Bacillus* sp. HLI | MA, GYM | $[M+NH_4]^+$ | Lipopeptide | Antimicrobial vs Gram-positive (*Staphylococcus*, *Streptococcus*, and *Enterococcus*), Gram-negative (*Flexibacter*, *Vibrio*, *Escherichia*, and *Salmonella*) bacteria; antiphytopathogenic vs *Xanthomonas campestris* | Several strains of terrestrial-derived *Bacillus subtilis* | Oka et al., 1993; Bartal et al., 2023; Kim et al.,2009 |
| 1040.6858 | 6.5 | Gageostatin A | $C_{52}H_{93}N_7O_{14}$ | x015 *Bacillus* sp. HLI | MA, GYM | $[M+H]^+$ | Lipopeptide | Moderate antimicrobial properties against Gram-positive (incl. *S.aureus*) and Gram-negative bacteria, and good antifungal activities | Marine sediment-derived *Bacillus* sp. | Tareq et al., 2014 |
| 1044.6581 | 7.81 | Surfactin B | $C_{52}H_{91}N_7O_{13}$ | x015 *Bacillus* sp. HLI | MA, GYM | $[M+Na]^+$ | Lipopeptide | Antimicrobial vs Gram-positive (*Staphylococcus, Streptococcus, and Enterococcus*), Gram-negative (*Flexibacter, Vibrio, Escherichia, and Salmonella*) bacteria; antiphytopathogenic vs *Xanthomonas campestris* | Several strains of terrestrial-derived *Bacillus subtilis* | Oka et al., 1993; Bartal et al., 2023; Kim et al., 2009 |
| 1045.5575 | 2.78 | C15-bacillomycin D | $C_{49}H_{76}N_{10}O_{15}$ | x015 *Bacillus* sp. HLI | MA | $[M+H]^+$ | Lipopeptide | Strong activity against *Staphylococcus* spp., incl MRSA | Several strains of terrestrial-derived *Bacillus subtilis* | Peypoux et al., 1984; Nam et al., 2021 |
| 1050.7074 | 8.29 | Surfactin D | $C_{54}H_{95}N_7O_{13}$ | x015 *Bacillus* sp. HLI | MA, GYM | $[M+H]^+$ | Lipopeptide | Antimicrobial vs Gram-positive (*Staphylococcus*, *Streptococcus*, and *Enterococcus*), Gram-negative (*Flexibacter*, *Vibrio*, *Escherichia*, *and Salmonella*) bacteria; antiphytopathogenic vs *Xanthomonas campestris* | Several strains of terrestrial-derived *Bacillus subtilis* | Oka et al., 1993; Bartal et al., 2023; Kim et al., 2009 |
| 1054.7015 | 6.8 | Gageostatin B | $C_{53}H_{95}N_7O_{14}$ | x015 *Bacillus* sp. HLI | MA, GYM | $[M+H]^+$ | Lipopeptide | Moderate antimicrobial properties against Gram-positive (incl. *S.aureus*) and Gram-negative bacteria, and | Marine sediment-derived *Bacillus* sp. | Tareq et al.,2014 |

| Experiment. m/z | $t_R$ Min | Putative annotation | Molecular formula | Strain number, Source bacterium, origin | Culture Media | Adduct | Chemical family | Reported bioactivity | Biological source | References |
|---|---|---|---|---|---|---|---|---|---|---|
| | | | | | | | | good antifungal activities | | |
| 1058.6731 | 8.09 | Surfactin C | $C_{53}H_{93}N_7O_{13}$ | x015 *Bacillus* sp. HLI | MA, GYM | $[M+Na]^+$ | Lipopeptide | Antimicrobial vs Gram-positive (*Staphylococcus*, *Streptococcus*, and *Enterococcus*), Gram-negative (*Flexibacter*, *Vibrio*, *Escherichia*, *and Salmonella*) bacteria; antiphytopathogenic vs *Xanthomonas campestris* | Several strains of terrestrial-derived *Bacillus subtilis* | Oka et al., 1993; Bartal et al., 2023; Kim et al., 2009 |

**Supplementary Table S16.** Putative annotation of metabolites produced by the extracts of the most active fungi isolated from eelgrass surfaces, i.e., healthy leaves (HLS), decaying leaves (DLS), roots (RS), and sediment (S) and seawater (W) references, detected in two culture media.

| **Experiment. *m/z*** | **$t_R$ Min** | **Putative annotation** | **Molecular formula** | **Strain number, source fungus, origin** | **Culture Media** | **Adduct** | **Chemical family** | **Reported bioactivity** | **Biological source** | **References** |
|---|---|---|---|---|---|---|---|---|---|---|
| 197.1290 | 0.59 | Cyclo(L-Pro-L-Val) | $C_{10}H_{16}N_2O_2$ | 233 *Cladosporium halotolerans* DLS<br>407 *Trichoderma viride* S<br>417 *Cystofilobasidium bisporidii* RS<br>425a, *Parathyridaria* sp. HLS<br>719 *Fusarium* sp. DLS<br>809 *Penicillium olsonii* S<br>813 *Aureobasidium pullulans* W<br>815 *Sarocladium strictum* W | M34, PDA | $[M+H]^+$ | Diketopiperazine | Bioactivity against marine pathogenic bacteria and MRSA. | Various bacterial and fungal species | Borthwick, 2012;<br>Qi et al., 2009;<br>Alshaibani et al., 2017 |
| 211.1445 | 0.65 | Cyclo(L-Pro-L-Leu) | $C_{11}H_{18}N_2O_2$ | 233 *Cladosporium halotolerans* DLS<br>407 *Trichoderma viride* S<br>417 *Cystofilobasidium bisporidii* RS<br>425a, *Parathyridaria* sp. HLS<br>813 *Aureobasidium pullulans* W | M34, PDA | $[M+H]^+$ | Diketopiperazine | Potent activity against twelve VRE strains, including *E. faecium* and *E. faecalis*, antitumor, anti-phytopathogenic vs *M. oryzae* | Various bacterial and fungal species including *Strept omyces* spp. | Rhee, 2002;<br>Rhee, 2003 |
| 247.1452 | 0.90 | Cyclo-(L-Val-L-Phe) | $C_{14}H_{18}N_2O_2$ | 233 *Cladosporium halotolerans* DLS<br>417 *Cystofilobasidium bisporidii* RS<br>809 *Penicillium olsonii* S | M34 | $[M+H]^+$ | Diketopiperazine | Quorum sensing regulator | *Pseudoalteromo nas sp.* isolated from the marine sponge *Hymeniacidon perleve* | Guo et al.,2011 |
| 261.1245 | 0.57 | Cyclo(L-Tyr-L-Pro) | $C_{14}H_{16}N_2O_3$ | 233 *Cladosporium halotolerans* DLS<br>407 *Trichoderma viride* S<br>813 *Aureobasidium pullulans* W<br>417 *Cystofilobasidium bisporidii* RS<br>719 *Fusarium* sp. DLS<br>809 *Penicillium olsonii* S<br>815 *Sarocladium strictum* W | M34 | $[M+H]^+$ | Diketopiperazine | Antimicrobial vs Gram-positive (incl. *S. aureus*) and Gram-negative bacteria, antifungal against human and plant pathogen | Several microganisms | Kumar et al., 2013;<br>Wattana-Amorn et al., 2016 |
| 301.1467 | 3.13 | Zearalenone | $C_{18}H_{22}O_5$ | 719 *Fusarium* sp. DLS | M34 | $[M-H_2O+H]^+$ | Polyketide | Mycotoxic, uterotropic | Moulded corn-derived *Fusarium graminearum* | Urry et al., 1966;<br>Utermark et al., 2007;<br>Saetang et al., 2016 |
| 303.1596 | 2.05 | Zearalenol | $C_{18}H_{24}O_5$ | 719 *Fusarium* sp. DLS | M34 | $[M-H_2O+H]^+$ | Polyketide | Mycotoxic, uterotropic | *Fusarium roseum* from corn or grain sorghum | Stipanovic et al., 1975 |
| 317.1386 | 2.27 | 5β-Hydroxyzearalenone | $C_{18}H_{22}O_6$ | 719 *Fusarium* sp. DLS | M34 | $[M-H_2O+H]^+$ | Polyketide | Mycotoxic, uterotropic | Seagrass-derived *Fusarium* sp. | Saetang et al., 2016 |
| 319.1569 | 3.13 | Zearalenone | $C_{18}H_{22}O_5$ | 719 *Fusarium* sp. DLS | M34, PDA | $[M+H]^+$ | Polyketide | Mycotoxic, uterotropic | Terrestrial *Fusarium graminearum* | Urry et al., 1966;<br>Utermark et al., 2007;<br>Saetang et al., 2016 |
| 335.1493 | 2.27 | 5β-Hydroxyzearalenone | $C_{18}H_{22}O_6$ | 719 *Fusarium* sp. DLS | M34 | $[M+H]^+$ | Polyketide | Mycotoxic, uterotropic | Seagrass - derived *Fusarium* sp. | Saetang et al.,2016 |

| Experiment. *m/z* | $t_R$ Min | Putative annotation | Molecular formula | Strain number, source fungus, origin | Culture Media | Adduct | Chemical family | Reported bioactivity | Biological source | References |
|---|---|---|---|---|---|---|---|---|---|---|
| 447.2380 | 3.3 | Atranone B | $C_{25}H_{34}O_7$ | 425a, *Parathyridaria* sp. HLS<br>434 *Parathyridaria* sp. HLS | M34, PDA | $[M+H]^+$ | Diterpenoid | Enhancer in neurite outgrowth | Terrestrial *Stachybotrys atra* | Hinkley et al.,1999 |
| 469.2202 | 3.3 | Atranone B | $C_{25}H_{34}O_7$ | 425a *Parathyridaria* sp.HLS | M34, PDA | $[M+Na]^+$ | Diterpenoid | Enhancer in neurite outgrowth | Terrestrial *Stachybotrys atra* | Hinkley et al.,1999 |
| 479.2642 | 3.2 | Atranone O | $C_{26}H_{38}O_8$ | 425a *Parathyridaria* sp.HLS<br>434 *Parathyridaria* sp. HLS | PDA | $[M+H]^+$ | Diterpenoid | Not reported | Marine crinoid-derived *Stachybotrys chartarum* | Li et al., 2017 |
| 501.2463 | 3.22 | Atranone O | $C_{26}H_{38}O_8$ | 425a *Parathyridaria* sp. HLS<br>434 *Parathyridaria* sp. HLS | M34, PDA | $[M+Na]^+$ | Diterpenoid | Not reported | Marine crinoid-derived *Stachybotrys chartarum* | Li et al., 2017 |
| 507.2283 | 4.65 | Asperphenamate | $C_{32}H_{30}N_2O_4$ | 809 *Penicillium olsonii* S | M34, PDA | $[M+H]^+$ | Peptide | Anticancer | Terrestrial *Aspergillus flavipes* | Clark et al.,1977; Frisvad et al., 2004; Frisvad et al., 2013 |
| 523.2237 | 3.38 | Asperphenamate analogue phenylalanine to tyrosine substitution | $C_{32}H_{30}N_2O_5$ | 809 *Penicillium olsonii* S | M34, PDA | $[M+H]^+$ | Peptide | Not reported | Marine-derived *Penicillium bialowiezense* | Kildgaard et al., 2014 |
| 529.0774 | 3.68 | Verrulactone B | $C_{28}H_{18}O_{12}$ | 809 *Penicillium olsonii* S | PDA | $[M-H_2O+H]^+$ | Benzocoumarin | Potent antibacterial activity against MRSA | Soil-derived *Penicillium verruculosum* | Kim et al., 2012 |
| 529.2108 | 4.65 | Asperphenamate | $C_{32}H_{30}N_2O_4$ | 809 *Penicillium olsonii* S | M34, PDA | $[M+Na]^+$ | Peptide | Anticancer | Terrestrial *Aspergillus flavipes* | Clark et al., 1977; Frisvad et al., 2004; Frisvad et al., 2013 |
| 543.1289 | 4.26 | Dimeric rubrofusarin | $C_{30}H_{22}O_{10}$ | 809 *Penicillium olsonii* S<br>719 *Fusarium* sp. DLS | M34, PDA | $[M+H]^+$ | Napthoquinone | Antimicrobial vs Gram-positive & -negative bacteria and *Candida* | *Fusarium graminearum* isolated from wheat kernels | Frandsen et al., 2006; Westphal et al., 2018 |
| 545.2057 | 3.38 | Asperphenamate analogue phenylalanine to tyrosine substitution | $C_{32}H_{30}N_2O_5$ | 809 *Penicillium olsonii* S | PDA | $[M+Na]^+$ | Peptide | Not reported | Marine-derived *Penicillium bialowiezense* | Kildgaard et al., 2014 |
| 557.0719 | 3.04 | Morakotin D | $C_{29}H_{16}O_{12}$ | 719 *Fusarium* sp. DLS | M34 | $[M+H]^+$ | Polyketide | Bioactivity against both *B. cereus* and *S. aureus* | *Cordyceps morakotii* isolated from an ant | Wang et al.,2019 |
| 557.1086 | 4.43 | Fuscofusarin | $C_{30}H_{20}O_{11}$ | 719 *Fusarium* sp. DLS | M34, PDA | $[M+H]^+$ | Napthoquinone | Antimicrobial vs Gram-positive & -negative bacteria and *Candida* | *Fusarium culmorum* from blighted wheat seedlings | Takeda et al., 1968 |
| 559.1180 | 4.33 | Dimeric rubrofusarin-9-hydroxyrubrofusarin | $C_{30}H_{22}O_{11}$ | 719 *Fusarium* sp. DLS | M34, PDA | $[M+H]^+$ | Napthoquinone | Antimicrobial vs Gram-positive & -negative bacteria and *Candida* | *Fusarium* graminearum from wheat kernels | Frandsen et al., 2006; Westphal et al., 2018 |
| 559.3850 | 5.15 | Exophilin A1 | $C_{30}H_{56}O_{10}$ | 813 *Aureobasidium pullulans* W | M34, PDA | $[M-H_2O+H]^+$ | Polyol lipid | Antimicrobial vs Gram-positive bacteria (i.e. *Enterococci* and *S. aureus*) | Marine sponge-derived *Exophiala pisciphila* | Doshida et al., 1996; Price et al., 2013; Bischoff et al., 2015 |
| 571.0877 | 3.45 | Aurofusarin | $C_{30}H_{18}O_{12}$ | 719 *Fusarium* sp. DLS | M34, PDA | $[M+H]^+$ | Napthoquinone | Bioactivity against Gram-positive (i.e *Lactobacilli*) | *Fusarium culmorum* from blighted wheat seedlings | Ashley et al., 1937; |

| Experiment. *m/z* | $t_R$ Min | Putative annotation | Molecular formula | Strain number, source fungus, origin | Culture Media | Adduct | Chemical family | Reported bioactivity | Biological source | References |
|---|---|---|---|---|---|---|---|---|---|---|
| 571.0879 | 3.72 | 3,4,3',4'-Bisdehydroxanthomegnin | $C_{30}H_{18}O_{12}$ | 809 *Penicillium olsonii* S<br>719 *Fusarium* sp. DLS | M34, PDA | $[M+H]^+$ | Napthoquinone | Antimicrobial vs Gram-positive and Gram-negative bacteria | Monascospore isolated *Nannizzia cajetalli* | Sedmera et al., 1981; Zeeck et al., 1979 |
| 573.1031 | 3.48 | Dimeric 9-hydroxyrubrofusarin-fuscofusarin | $C_{30}H_{20}O_{12}$ | 809 *Penicillium olsonii* S<br>719 *Fusarium* sp. DLS | M34, PDA | $[M+H]^+$ | Napthoquinone | Antimicrobial vs Gram-positive and Gram-negative bacteria and *Candida* | *Fusarium graminearum* from wheat kernels | Frandsen et al., 2006; Westphal et al., 2018 |
| 575.1187 | 3.79 | Xanthomegnin | $C_{30}H_{22}O_{12}$ | 719 *Fusarium* sp. DLS | M34, PDA | $[M+H]^+$ | Napthoquinone | Bioactivite vs Gram-positive and Gram-negative bacteria | Monascospore isolated *Nannizzia cajetalli* | Sedmera et al., 1981; Zeeck et al., 1979 |
| 589.0986 | 3.48 | Xanthoepocin | $C_{30}H_{22}O_{14}$ | 809 *Penicillium olsonii* S | M34, PDA | $[M-H_2O+H]^+$ | Anthraquinone | Bioactive vs pathogenic yeasts, MRSA, vancomycin-resistant *Enterococcus faecium* weak cytotoxic | *Penicillium simplicissimum* | Igarashi et al., 2000; Vrabl et al., 2022 |
| 599.3805 | 5.15 | Exophilin A1 | $C_{30}H_{56}O_{10}$ | 813 *Aureobasidium pullulans* W | M34, PDA | $[M+Na]^+$ | Polyol lipid | Bioactive against Gram-positive bacteria (*Enterococci* and *S. aureus*) | Marine sponge-derived *Exophiala pisciphila* | Doshida et al., 1996; Price et al., 2013; Bischoff et al., 2015 |
| 607.1093 | 3.68 | Xanthoepocin | $C_{30}H_{22}O_{14}$ | 809 *Penicillium olsonii* S | PDA | $[M+H]^+$ | Anthraquinone | Broad spectrum activity vs pathogenic yeasts, MRSA, vancomycin-resistant *Enterococcus faecium*, cytotoxic | *Penicillium simplicissimum* | Igarashi et al., 2000; Vrabl et al., 2022 |
| 633.4208 | 4.77 | Glycerolliamocin A1 | $C_{33}H_{62}O_{12}$ | 813 *Aureobasidium pullulans* W | M34 | $[M-H_2O+H]^+$ | Polyol lipid | Antibacterial specifically vs *Streptococcus* sp. | Diverse terrestrial *Aureobasidium pullulans* | Manitchotpisit et al., 2011; Price et al., 2013; Leathers et al., 2015; Bischoff et al., 2015 |
| 659.4385 | 5.84 | Fusaristatin A | $C_{36}H_{58}N_4O_7$ | 719 *Fusarium* sp. DLS | M34, PDA | $[M+H]^+$ | Cyclic lipopeptide | Cytotoxic, antiphytopathogenic vs *Glomerella acutata* | *Fusarium* sp. isolated from the stem of *Maackia chinensis* | Shiono et al., 2007; Li et al., 2016 |
| 673.4146 | 4.76 | Glycerolliamocin A1 | $C_{33}H_{62}O_{12}$ | 813 *Aureobasidium pullulans* W | M34, PDA | $[M+Na]^+$ | Polyol lipid | Antibacterial specifically vs *Streptococcus* sp. | Diverse terrestrial *Aureobasidium pullulans* | Manitchotpisit et al., 2011; Price et al., 2013; Leathers et al., 2015; Bischoff et al., 2015 |
| 674.4374 | 6.98 | Fusaristatin B | $C_{37}H_{59}N_3O_8$ | 719 *Fusarium* sp. DLS | M34 | $[M+H]^+$ | Cyclic lipopeptide | Cytotoxic | *Fusarium* sp. isolated from the stem of *Maackia chinensis* | Shiono et al., 2007 |

| Experiment. *m/z* | $t_R$ Min | Putative annotation | Molecular formula | Strain number, source fungus, origin | Culture Media | Adduct | Chemical family | Reported bioactivity | Biological source | References |
|---|---|---|---|---|---|---|---|---|---|---|
| 675.4319 | 4.29 | Arabinitol liamocin A1 | $C_{35}H_{66}O_{14}$ | 813 *Aureobasidium pullulans* W | M34, PDA | $[M-2H_2O+H]^+$ | Polyol lipid | Antibacterial specifically vs *Streptococcus* sp. | Diverse terrestrial-*Aureobasidium pullulans* | Manitchotpisit et al., 2011; Price et al., 2013; Leathers et al., 2015; Bischoff et al., 2015 |
| 693.4431 | 4.29 | Arabinitol liamocin A1 | $C_{35}H_{66}O_{14}$ | 813 *Aureobasidium pullulans* W | M34, PDA | $[M-H_2O+H]^+$ | Polyol lipid | Antibacterial specifically vs *Streptococcus* sp. | Diverse terrestrial-*Aureobasidium pullulans* | Manitchotpisit et al., 2011; Price et al., 2013; Leathers et al., 2015; Bischoff et al., 2015 |
| 703.4242 | 4.48 | Threitol liamocin A1 | $C_{34}H_{64}O_{13}$ | 813 *Aureobasidium pullulans* W | M34 | $[M+Na]^+$ | Polyol lipid | Antibacterial specifically vs *Streptococcus* sp. | Diverse terrestrial-*Aureobasidium pullulans* | Manitchotpisit et al., 2011; Price et al., 2013; Leathers 2015; Bischoff et al., 2015 |
| 705.4430 | 4.2 | Liamocin A1 | $C_{36}H_{68}O_{15}$ | 813 *Aureobasidium pullulans* W | M34, PDA | $[M-2H_2O+H]^+$ | Polyol lipid | Antibacterial specifically vs *Streptococcus* sp. | Diverse terrestrial-*Aureobasidium pullulans* | Manitchotpisit et al., 2011; Price et al., 2013; Leathers et al., 2015; Bischoff et al., 2015 |
| 723.4531 | 4.22 | Liamocin A1 | $C_{36}H_{68}O_{15}$ | 809 *Penicillium olsonii* S, 813 *Aureobasidium pullulans* W | M34, PDA | $[M-H_2O+H]^+$ | Polyol lipid | Antibacterial specifically vs *Streptococcus* sp. | Diverse terrestrial-*Aureobasidium pullulans* | Manitchotpisit et al., 2011; Price et al., 2013; Leathers et al., 2015; Bischoff et al., 2015 |
| 727.4991 | 6.51 | Exophilin B1 | $C_{40}H_{74}O_{13}$ | 813 *Aureobasidium pullulans* W | M34, PDA | $[M-2H_2O+H]^+$ | Polyol lipid | Antimicrobial vs Gram-positive bacteria (*Enterococci, S. aureus*) | Diverse terrestrial-*Aureobasidium pullulans* | Price et al., 2013; Bischoff et al., 2015; Price et al., 2017 |
| 733.4375 | 4.29 | Arabinitol liamocin A1 | $C_{35}H_{66}O_{14}$ | 813 *Aureobasidium pullulans* W | M34, PDA | $[M+Na]^+$ | Polyol lipid | Antibacterial specifically vs *Streptococcus* sp. | Diverse terrestrial-*Aureobasidium pullulans* | Manitchotpisit et al., 2011; Price et al., 2013; Leathers et al., 2015; Bischoff et al., 2015 |
| 735.4522 | 4.87 | Arabinitol liamocin A2 | $C_{37}H_{68}O_{15}$ | 813 *Aureobasidium pullulans* W | M34, PDA | $[M-H_2O+H]^+$ | Polyol lipid | Antibacterial specifically vs *Streptococcus* sp. | Diverse terrestrial-*Aureobasidium pullulans* | Manitchotpisit et al., 2011; Price et al., 2013; Leathers et al., 2015; Bischoff et al., 2015 |
| 745.4344 | 4.85 | Threitol liamocin A2 | $C_{36}H_{66}O_{14}$ | 813 *Aureobasidium pullulans* W | M34 | $[M+Na]^+$ | Polyol lipid | Antibacterial specifically vs *Streptococcus* sp. | Diverse terrestrial-*Aureobasidium pullulans* | Manitchotpisit et al., 2011; Price et al., 2013; Leathers et al., 2015; Bischoff et al., 2015 |
| 745.5101 | 6.51 | Exophilin B1 | $C_{40}H_{74}O_{13}$ | 813 *Aureobasidium pullulans* W | M34, PDA | $[M-H_2O+H]^+$ | Polyol lipid | Antibacterial specifically vs *Streptococcus* sp. | Diverse terrestrial-*Aureobasidium pullulans* | Price et al., 2013; Bischoff et al., 2015; Price et al., 2017 |
| 747.4531 | 4.81 | Liamocin A2 | $C_{38}H_{70}O_{16}$ | 813 *Aureobasidium pullulans* W | M34, PDA | $[M-2H_2O+H]^+$ | Polyol lipid | Antibacterial specifically vs *Streptococcus* sp. | Diverse terrestrial-*Aureobasidium pullulans* | Manitchotpisit et al., 2011; Price et al., 2013; Leathers et al., 2015; Bischoff et al., 2015 |

| Experiment. *m/z* | $t_R$ Min | Putative annotation | Molecular formula | Strain number, source fungus, origin | Culture Media | Adduct | Chemical family | Reported bioactivity | Biological source | References |
|---|---|---|---|---|---|---|---|---|---|---|
| 763.4459 | 4.19 | Liamocin A1 | $C_{36}H_{68}O_{15}$ | 809 *Penicillium olsonii* S, 813 *Aureobasidium pullulans* W | M34, PDA | $[M+Na]^+$ | Polyol lipid | Antibacterial specifically vs *Streptococcus* sp. | Diverse terrestrial-*Aureobasidium pullulans* | Manitchotpisit et al., 2011; Price et al., 2013; Leathers et al., 2015; Bischoff et al., 2015 |
| 765.4638 | 4.81 | Liamocin A2 | $C_{38}H_{70}O_{16}$ | 813 *Aureobasidium pullulans* W | M34, PDA | $[M-H_2O+H]^+$ | Polyol lipid | Antibacterial specifically vs *Streptococcus* sp. | Diverse terrestrial-*Aureobasidium pullulans* | Manitchotpisit et al., 2011; Price et al., 2013; Leathers et al., 2015; Bischoff et al., 2015 |
| 775.4442 | 4.82 | Arabinitol liamocin A2 | $C_{37}H_{68}O_{15}$ | 813 *Aureobasidium pullulans* W | M34, PDA | $[M+Na]^+$ | Polyol lipid | Antibacterial specifically vs *Streptococcus* sp. | Diverse terrestrial-*Aureobasidium pullulans* | Manitchotpisit et al., 2011; Price et al., 2013; Leathers et al., 2015; Bischoff et al., 2015 |
| 785.5050 | 6.51 | Exophilin B1 | $C_{40}H_{74}O_{13}$ | 813 *Aureobasidium pullulans* W | M34, PDA | $[M+Na]^+$ | Polyol lipid | Antibacterial specifically vs *Streptococcus* sp. | Diverse terrestrial-*Aureobasidium pullulans* | Price et al., 2013; Bischoff et al., 2015; Price et al., 2017 |
| 787.5227 | 7.18 | Halymecin A | $C_{42}H_{76}O_{14}$ | 813 *Aureobasidium pullulans* W | M34, PDA | $[M-H_2O+H]^+$ | Polyol lipid | Antimicroalgal, low activity vs Gram-positive and Gram-negative bacteria | Marine algae-derived *Fusarium* sp. | Chen et al.,1996; Le Dang et al., 2014 |
| 805.4571 | 4.81 | Liamocin A2 | $C_{38}H_{70}O_{16}$ | 809 *Penicillium olsonii* S 813 *Aureobasidium pullulans* W | M34, PDA | $[M+Na]^+$ | Polyol lipid | Antibacterial specifically vs *Streptococcus* sp. | Diverse terrestrial-*Aureobasidium pullulans* | Manitchotpisit et al., 2011; Price et al., 2013; Leathers et al., 2015; Bischoff et al., 2015 |
| 817.4919 | 4.36 | Halymecin D | $C_{40}H_{74}O_{15}$ | 813 *Aureobasidium pullulans* W | M34 | $[M+Na]^+$ | Polyol lipid | Not reported | Marine algae-derived *Acremonium* sp. | Chen et al.,1996 |
| 827.5162 | 7.19 | Halymecin A | $C_{42}H_{76}O_{14}$ | 813 *Aureobasidium pullulans* W | M34, PDA | $[M+Na]^+$ | Polyol lipid | Antimicroalgal, low activity vs Gram-positive and Gram-negative bacteria | Marine algae-derived *Fusarium* sp. | Chen et al., 1996; Le Dang et al., 2014 |
| 859.5401 | 6.26 | Glycerol liamocin B1 | $C_{43}H_{80}O_{15}$ | 813 *Aureobasidium pullulans* W | M34, PDA | $[M+Na]^+$ | Polyol lipid | Antibacterial specifically vs *Streptococcus* sp. | Diverse terrestrial-*Aureobasidium pullulans* | Manitchotpisit et al., 2011; Price et al., 2013; Leathers et al., 2015; Bischoff et al., 2015 |
| 884.5278 | 5.19 | Trichorzin HA-2 | $C_{78}H_{138}N_{20}O_{20}$ | 407 *Trichoderma viride* S | M34 | $[M+2H]^{++}$ | Peptaibol | Potent antibiotic vs Gram-positive bacteria incl. *S.aureus*, anti-phytopathogenic vs fungus *Sclerotium cepivorum.* | Soil-derived *Trichoderma harzianum* | Goulard et al., 1995 |
| 889.5466 | 6.18 | Threitol liamocin B1 | $C_{44}H_{82}O_{16}$ | 813 *Aureobasidium pullulans* W | M34, PDA | $[M+Na]^+$ | Polyol lipid | Antibacterial specificly vs *Streptococcus* sp. | Diverse terrestrial-*Aureobasidium pullulans* | Manitchotpisit et al., 2011; Price et al., 2013; Leathers et al., 2015; Bischoff et al., 2015 |

| Experiment. *m/z* | $t_R$ Min | Putative annotation | Molecular formula | Strain number, source fungus, origin | Culture Media | Adduct | Chemical family | Reported bioactivity | Biological source | References |
|---|---|---|---|---|---|---|---|---|---|---|
| 891.5394 | 5.27 | Trichorzin PA VI | $C_{84}H_{140}N_{20}O_{22}$ | 407 *Trichoderma viride* S | M34 | $[M+2H]^{++}$ | Peptaibol | Potent activity against Gram-positive bacteria incl. *S.aureus* | Soil-derived *Trichoderma harzianum* | Duval et al.,1997 |
| 898.5440 | 5.47 | Trichorzin PA VIII | $C_{85}H_{142}N_{20}O_{22}$ | 407 *Trichoderma viride* S | M34 | $[M+2H]^{++}$ | Peptaibol | Potent activity against Gram-positive bacteria incl. *S.aureus* | Soil-derived *Trichoderma harzianum* | Duval et al., 1997 |
| 913.0174 | 5.4 | Zervamicin II-4 | $C_{89}H_{137}N_{19}O_{22}$ | 407 *Trichoderma viride* S | M34 | $[M+2H]^{++}$ | Peptaibol | Potent activity against Gram-positive bacteria incl. *S.aureus* | Marine fungus *Emericellopsis salmosynnemat a* | Rinehart et al., 1981 |
| 913.5140 | 6.03 | Zervamicin IB | $C_{89}H_{136}N_{18}O_{23}$ | 407 *Trichoderma viride* S | M34 | $[M+2H]^{++}$ | Peptaibol | Potent activity vs Gram-positive bacteria incl. *S.aureus* | Marine fungus *Emericellopsis salmosynnemat a* | Argoudelis et al., 1974 |
| 919.5644 | 5.81 | Arabinitol liamocin B1 | $C_{45}H_{84}O_{17}$ | 813 *Aureobasidium pullulans* W | M34, PDA | $[M+Na]^{+}$ | Polyol lipid | Antibacterial specificly vs *Streptococcus* sp. | Diverse terrestrial- *Aureobasidium pullulans* | Manitchotpisit et al., 2011; Price et al., 2013; Leathers et al., 2015; Bischoff et al., 2015 |
| 920.0230 | 6.14 | Zervamicin IIB | $C_{90}H_{139}N_{19}O_{22}$ | 407 *Trichoderma viride* S | M34 | $[M+2H]^{++}$ | Peptaibol | Potent activity against Gram-positive bacteria incl. *S.aureus* | Marine fungus *Emericellopsis salmosynnemat a* | Argoudelis et al., 1974 |
| 949.5727 | 5.53 | Liamocin B1 | $C_{46}H_{86}O_{18}$ | 813 *Aureobasidium pullulans* W | M34, PDA | $[M+Na]^{+}$ | Polyol lipid | Antibacterial specifically vs *Streptococcus* sp. | Diverse terrestrial- *Aureobasidium pullulans* | Manitchotpisit et al., 2011; Price et al., 2013; Leathers et al., 2015; Bischoff et al., 2015 |
| 1009.5922 | 4.23 | Halymecin F | $C_{50}H_{88}O_{20}$ | 813 *Aureobasidium pullulans* W | M34, PDA | $[M+H]^{+}$ | Polyol lipid | Antiphytopathogenic vs bacteria *A. konjaci, A. tumefaciens, B. glumae* | *Simplicillium lamellicola* isolated from the mycelia of *Botrytis cinerea* | Le Dang et al., 2014 |

**Supplementary Table S17**. Putative annotation of metabolites produced by two media extracts of *Acrostalagmus luteoalbus* 720 isolated from the inner tissues of the decaying eelgrass leaves (DLI).

| **Experimental *m/z*** | **$t_R$ Min** | **Putative annotation** | **Molecular formula** | **Culture Media** | **Adduct** | **Chemical family** | **Reported bioactivity** | **Biological source** | **References** |
|---|---|---|---|---|---|---|---|---|---|
| 197.1290 | 0.60 | Cyclo(L-Pro-L-Val) | $C_{10}H_{16}N_2O_2$ | PDA, M34 | $[M+H]^+$ | Diketopiperazine | Bioactivity against marine pathogenic bacteria and MRSA. | Various bacterial and fungal species | Borthwick, 2012; Qi et al., 2009; Alshaibani et al., 2017 |
| 211.1448 | 0.75 | Cyclo(L-Pro-L-Leu) | $C_{11}H_{18}N_2O_2$ | PDA, M34 | $[M+H]^+$ | Diketopiperazine | Antimicrobial against VRE strains, incl. *E. faecium* and *E. faecalis*, anti-phytopathogenic vs *M. oryzae* | Various bacterial and fungal species including *Streptomyces* spp. | Rhee, 2002; Rhee, 2003 |
| 245.1293 | 0.8 | Cyclo(L-Phe-L-Pro) | $C_{14}H_{16}N_2O_2$ | PDA,M34 | $[M+H]^+$ | Diketopiperazine | Antibacterial vs VRE strains incl *E. faecium* and *E. faecalis* and *S.aureus*, antifungal against human and plant pathogens | Isolated from *Pseudomonas fluorescens* and *P. alcaligenes* | Rhee et al., 2001; Ström et al.,2002 |
| 261.1239 | 0.74 | Cyclo(L-Tyr-L-Pro) | $C_{14}H_{16}N_2O3$ | M34 | $[M+H]^+$ | Diketopiperazine | Antimicrobial vs Gram-positive (incl. *S. aureus*) and Gram-negative bacteria, antifungal against human and plant pathogens | Several microganisms | Kumar et al., 2013; Wattana-Amorn et al., 2016 |
| 681.1089 | 3.37 | 11'-deoxyverticillin A | $C_{30}H_{28}N_6O_5S_4$ | PDA, M34 | $[M+H]^+$ | Epipolythiodioxopiperazine | Cytotoxic | Epiphytic *Penicillium* sp. isolated from green alga *Avruinvillea longicuulis* | Son et al., 1999 |
| 697.1032 | 3.15 | Chaetocin | $C_{30}H_{28}N_6O_6S_4$ | PDA, M34 | $[M+H]^+$ | Epipolythiodioxopiperazine | Cytotoxic, antibacterial vs MRSA, *Enterococcus faecalis* | *Chaetomium minutum* | Hauser et al.,1970 |
| 713.0995 | 3.36 | Verticillin B | $C_{30}H_{28}N_6O_7S_4$ | PDA, M34 | $[M+H]^+$ | Epipolythiodioxopiperazine | Cytotoxic | *Verticillium* sp. isolated from a basidiocarp of *Coltricia cinnamomea* | Minato et al., 1973 |
| 745.0707 | 3.46 | Verticillin C | $C_{30}H_{28}N_6O_7S_5$ | PDA, M34 | [M+H]+ | Epipolythiodioxopiperazine | Cytotoxic | *Verticillium* sp. isolated from a basidiocarp of *Coltricia cinna-momea* | Minato et al.,1973 |
| 761.0667 | 2.63 | Chetracin B | $C_{30}H_{28}N_6O_8S_5$ | PDA, M34 | $[M+H]^+$ | Epipolythiodioxopiperazine | Cytotoxic | Soil collected under lichens-derived *Oidiodendron truncatum* | Li et al., 2012 |
| 793.0389 | 2.73 | Chetracin C | $C_{30}H_{28}N_6O_8S_6$ | PDA, M34 | $[M+H]^+$ | Epipolythiodioxopiperazine | Cytotoxic | Soil collected under lichens-derived *Oidiodendron truncatum* | Li et al., 2012 |

**Supplementary Fig. S1.** FBMN showing the relative abundance in MA and GYM media of ions detected in the most active bacterial isolates from the surface of *Zostera marina*, i.e., healthy leaves (HLS), decaying leaves (DLS), roots (RS), plus the sediment (S) and seawater (W) references. Primary metabolite clusters are coloured in grey. Annotated molecular families (MF) are highlighted in frames.

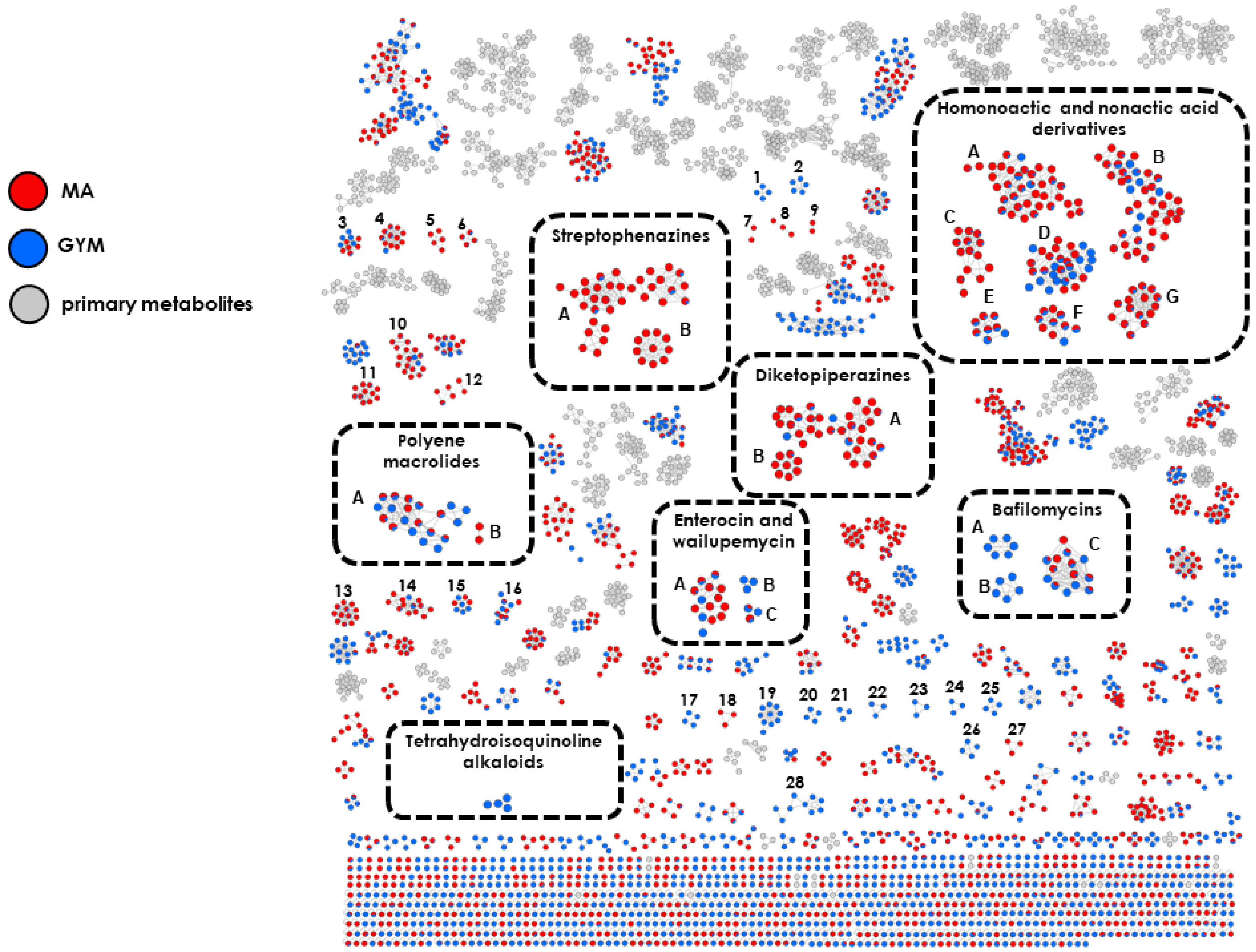

**Supplementary Fig. S2.** FBMN showing the relative abundance in MA and GYM media of ions detected in the most active bacterial isolates from the inner tissues of *Zostera marina*, i.e., healthy leaves (HLI), decaying leaves (DLI), roots (RI). Primary metabolite clusters are coloured in grey. Annotated molecular families (MF) are highlighted in frames.

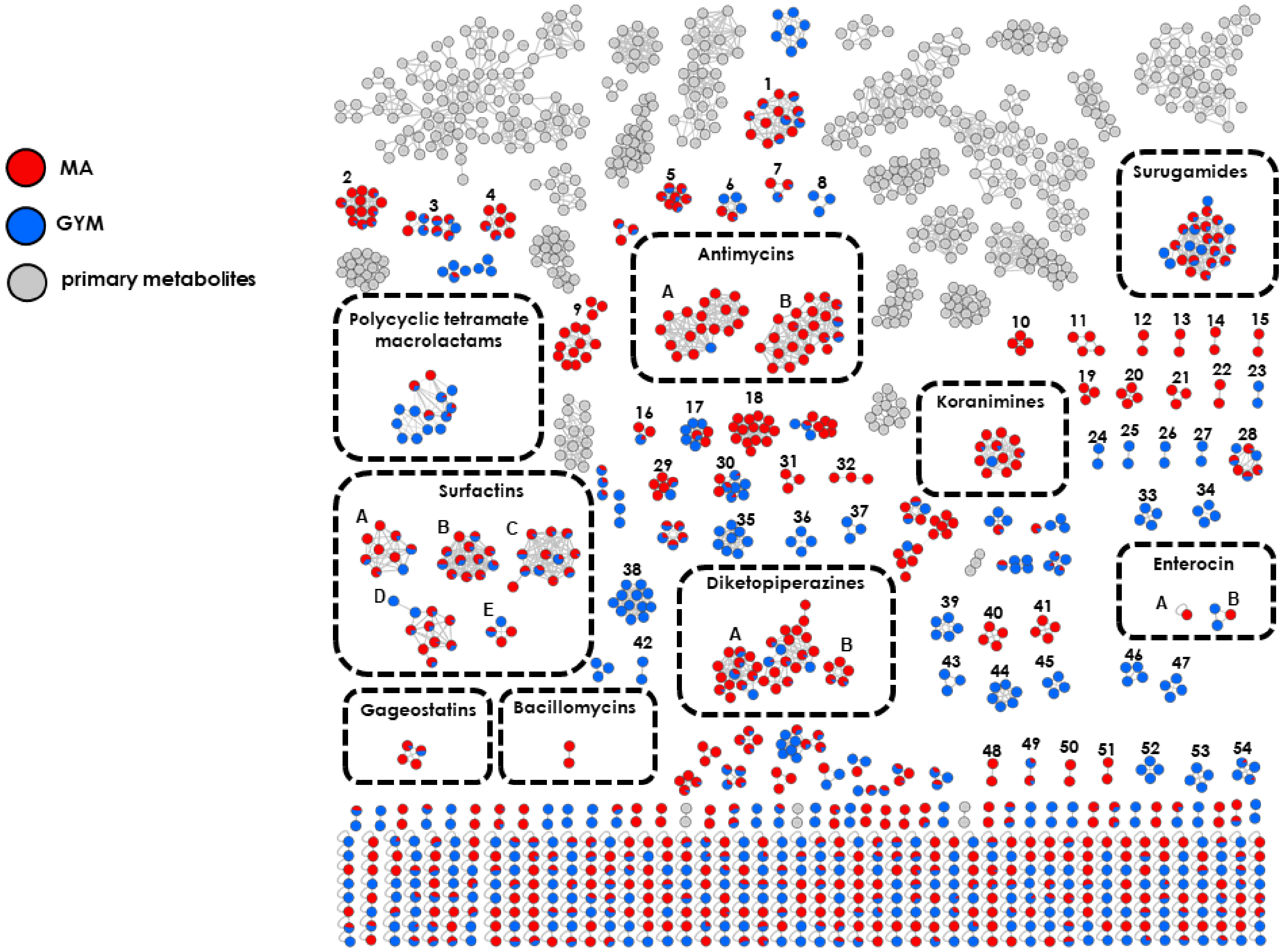

**Supplementary Fig. S3.** FBMN showing the relative abundance of ions detected in M34 and PDA media extracts of the most active fungi isolated from all eelgras surfaces, i.e., healthy leaves (HLS), decaying leaves (DLS), roots (RS), plus and the sediment (S) and seawater (W) references. Annotated molecular families (MF) are highlighted in frames. Primary metabolites clusters are shown in grey. The Venn diagrams show the number of exclusive and shared nodes detected in two culture media.

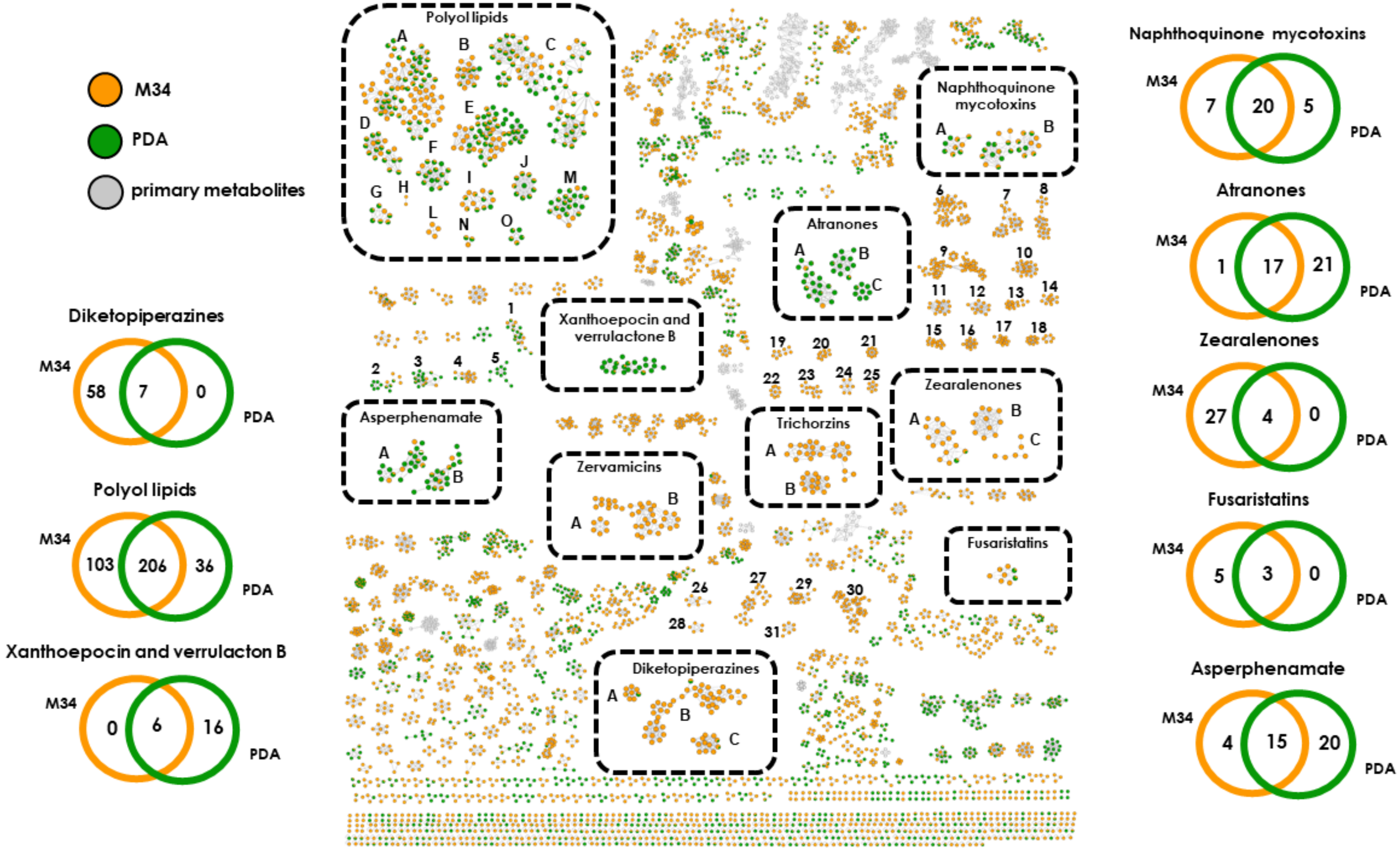

**Supplementary Fig. S4.** Comparative metabolomics of five *Streptomyces* species cultured in two culture media (MA and GYM).

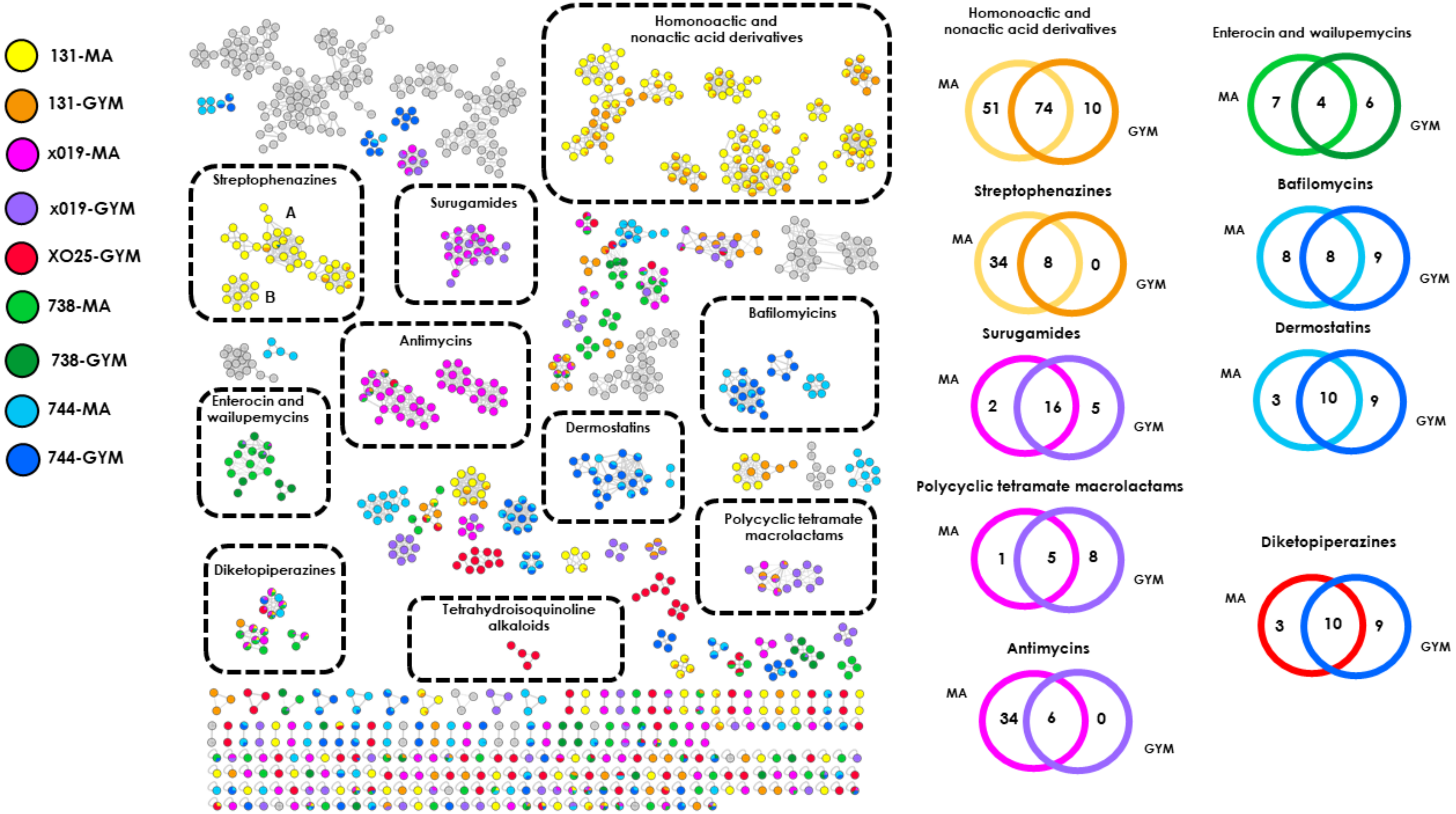

**Supplementary Fig. S5**: Chemical structures of annotated compounds

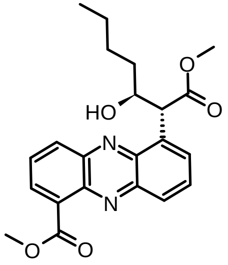

Streptophenazine E

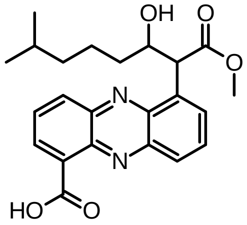

Streptophenazine C

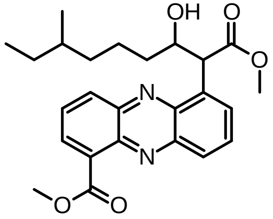

Streptophenazine B

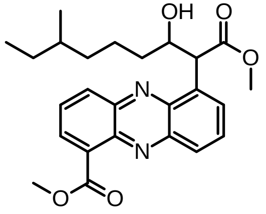

Streptophenazine G

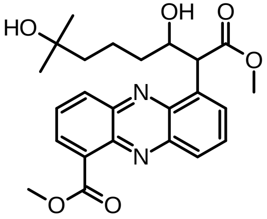

Streptophenazine H

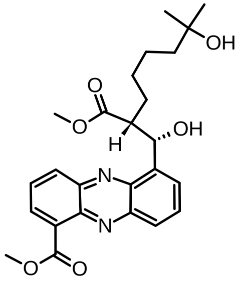

Streptophenazine I

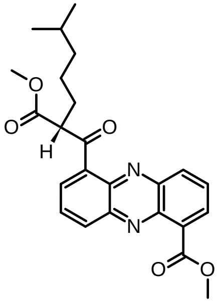

Oxo-streptophenazine A

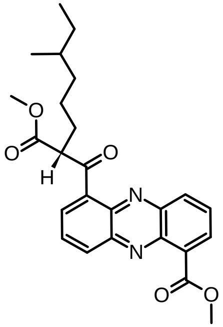

Oxo-streptophenazine G

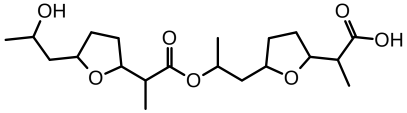

Nonactyl nonactate

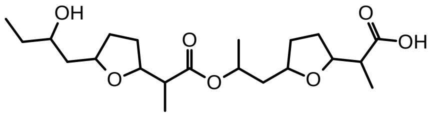

Bonactin

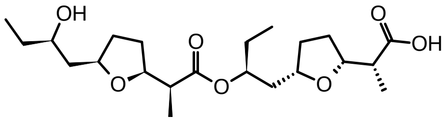

Homononactyl Homononactate

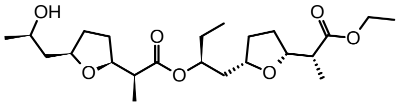

Ethyl Homononactyl Nonactate

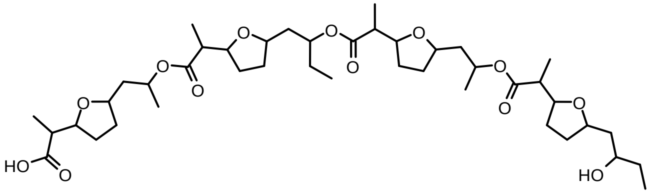

Seco-dinactin

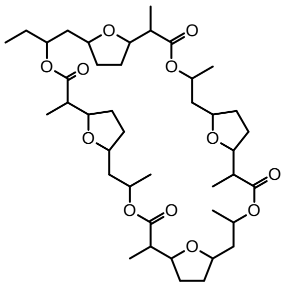

Monactin

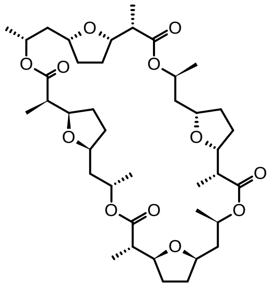

Nonactin

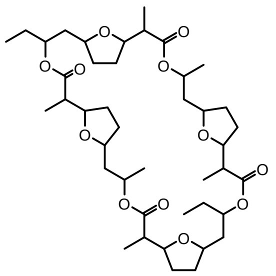

Dinactin

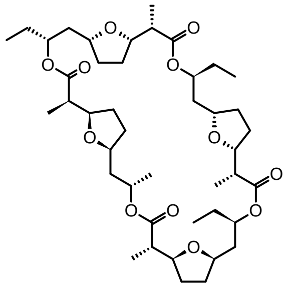

Trinactin

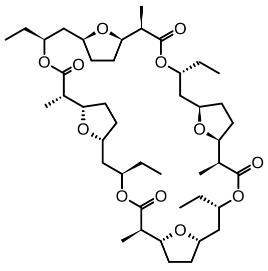

Tetranactin

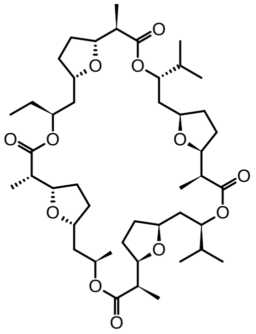

Macrotetrolide C

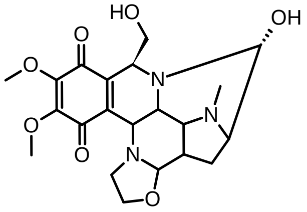

Naphthyridinomycin

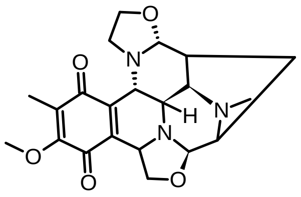

Bioxalomycin Beta 2

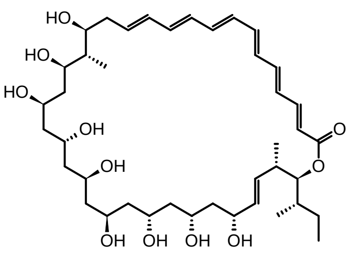
Dermostatin B

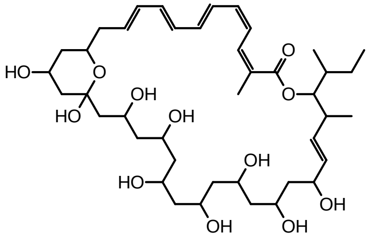
Pn00053

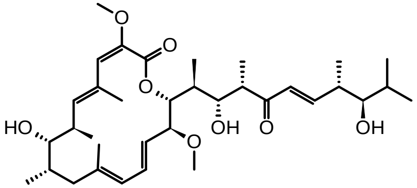
Bafilomycin D

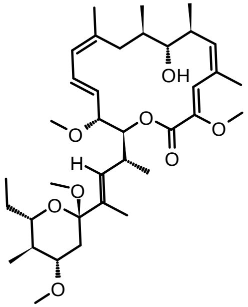
17,18-Dehydro-19,21-di-O-methyl-24-demethyl-bafilomycin A1

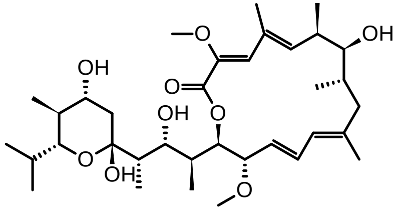
Bafilomycin A1

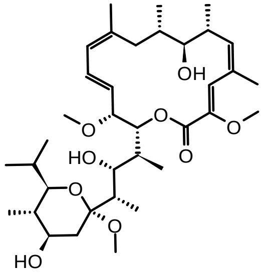
Bafilomycin A2

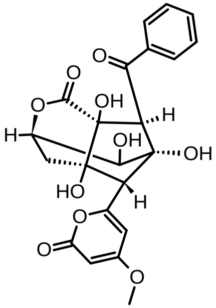
Enterocin

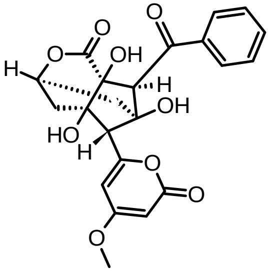
8-Deoxyenterocin

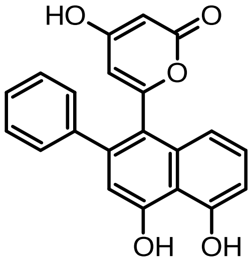
Wailupemycin G

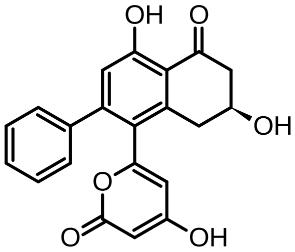
Wailupemycin F

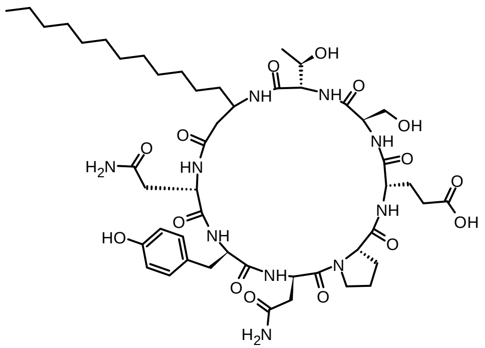

C15-bacillomycin D

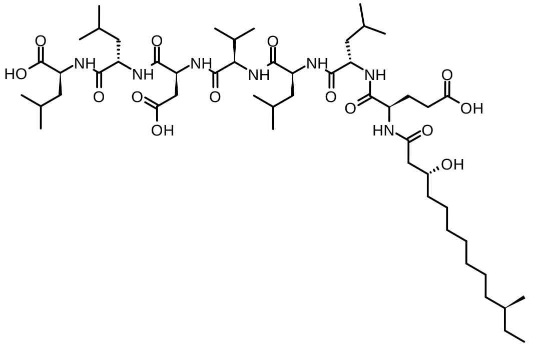

Gageostatin A

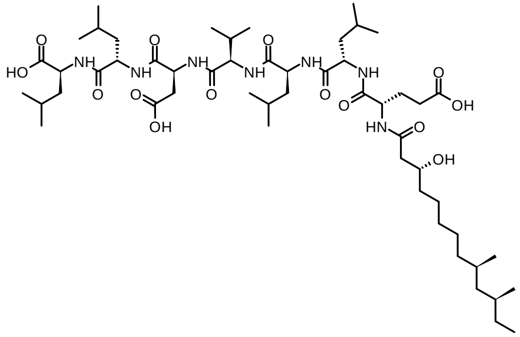

Gageostatin B

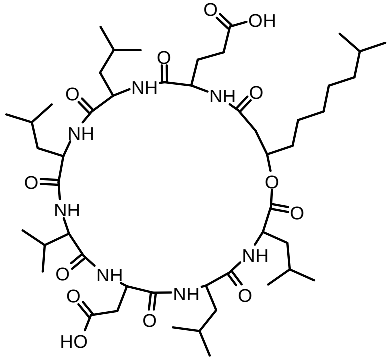

Surfactin C-11

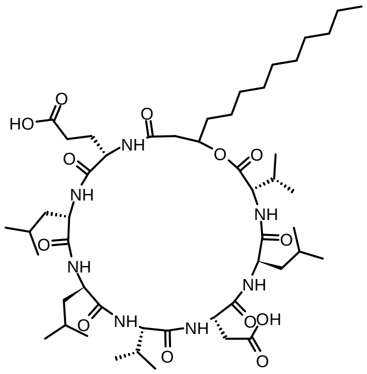

[Val7]-Surfactin

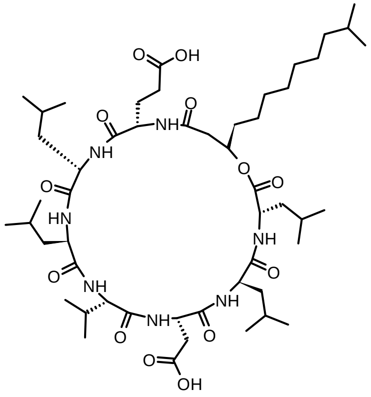

Surfactin A

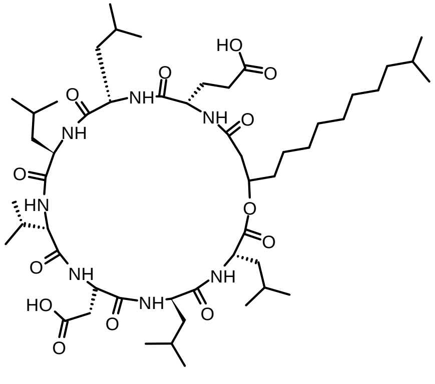

Surfactin B

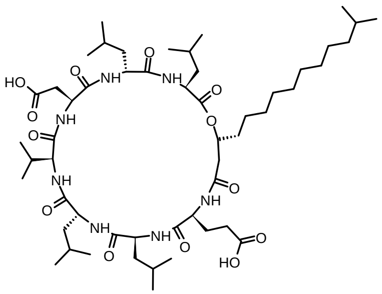

Surfactin C

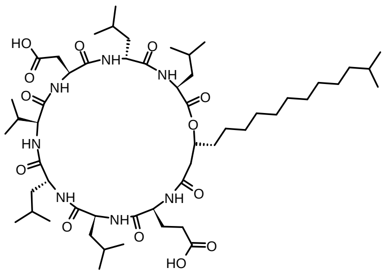

Surfactin D

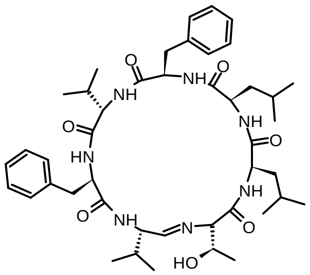

Koranimine

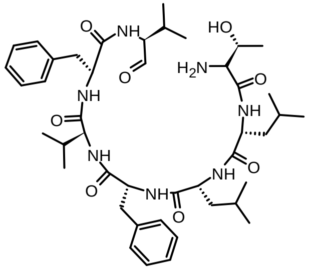

Koranimine B

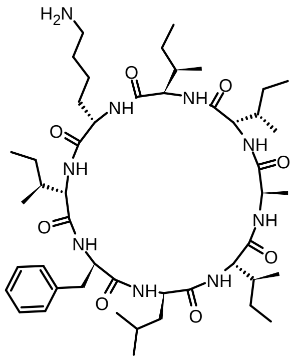

Surugamide A

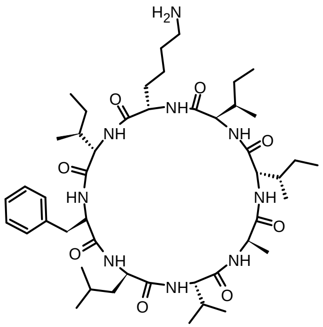

Surugamide E

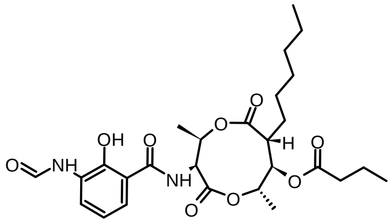
Antimycin A2

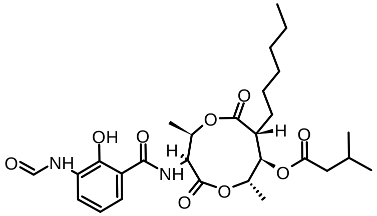
Antimycin A1

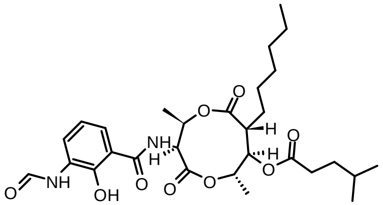
Antimycin A15

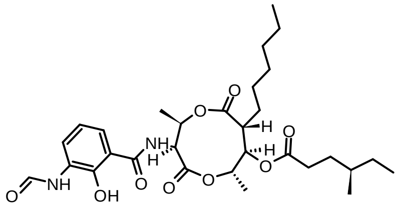
Antimycin A16

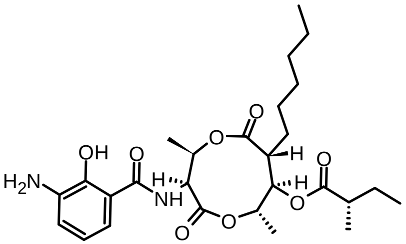
Deformylated antimycin A1a

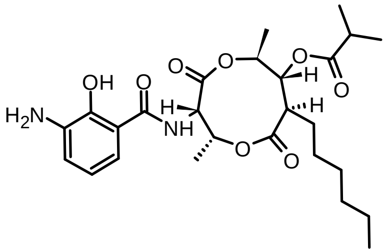
Deformylated antimycin A2a

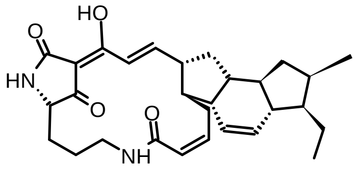
Ikarugamycin

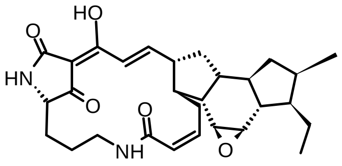
Ikarugamycin epoxide

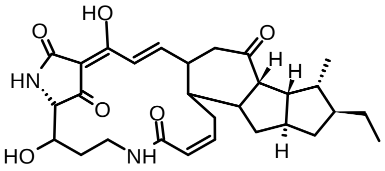
Maltophilin

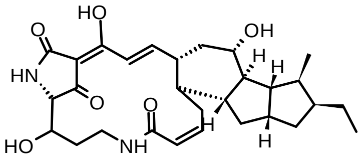
Dihydromaltophilin

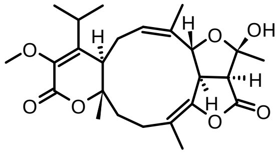
Atranone B

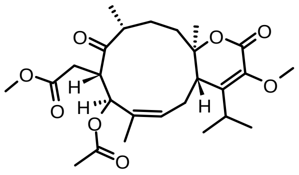
Atranone O

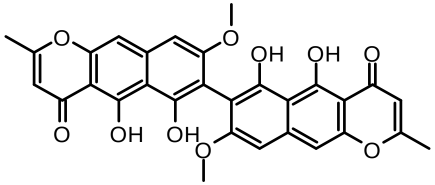

Dimeric rubrofusarin

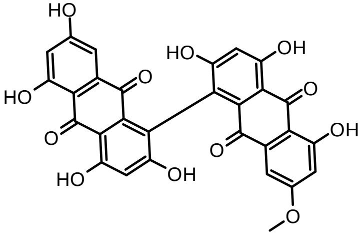

Morakotin D

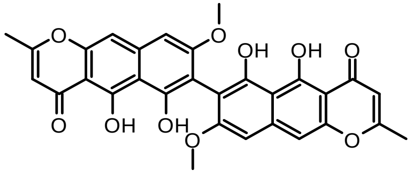

Fuscofusarin

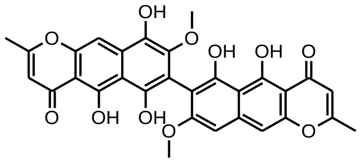

Dimeric rubrofusarin-9-hydroxyrubrofusarin

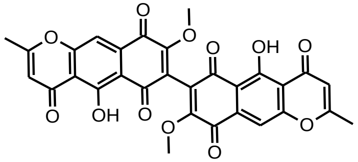

Aurofusarin

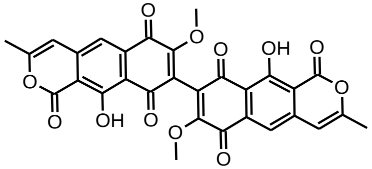

3,4,3',4'-Bisdehydroxanthomegnin

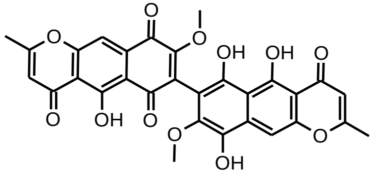

Dimeric 9-hydroxyrubrofusarin-fuscofusarin

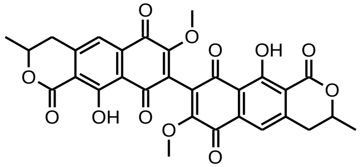

Xanthomegnin

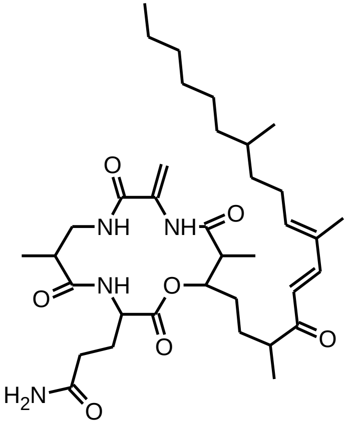

Fusaristatin A

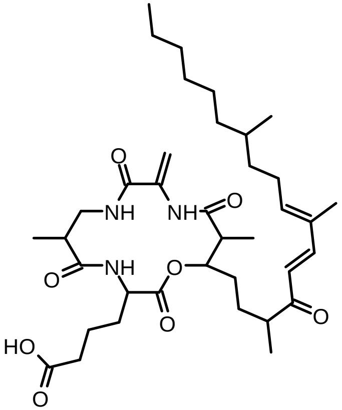

Fusaristatin B

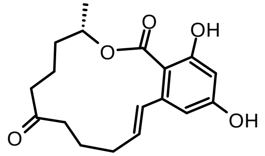

Zearalenone

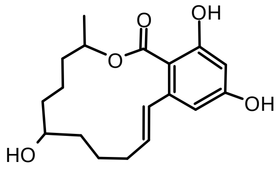

Zearalenol

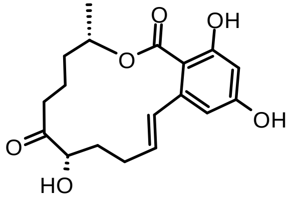

5′β-Hydroxyzearalenone

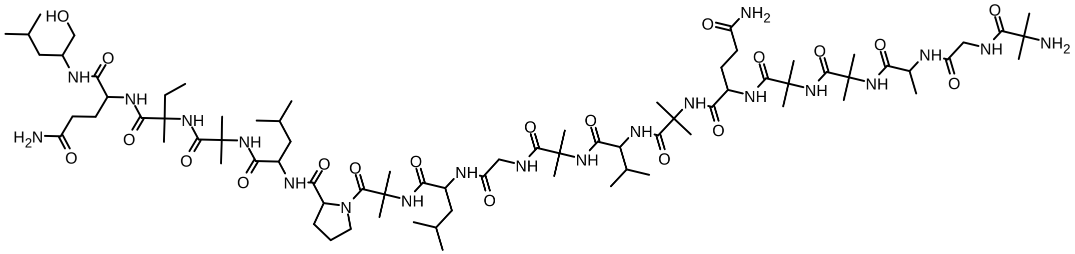

Trichorzin HA II

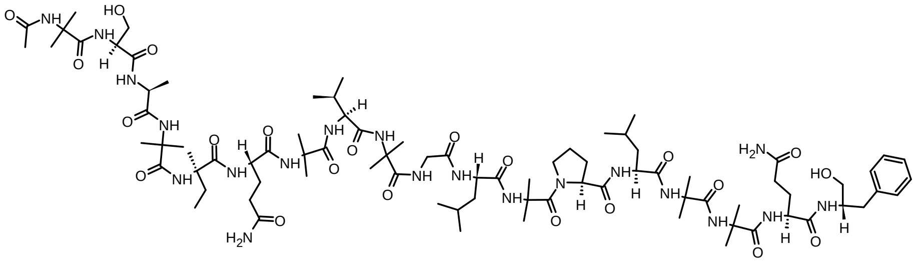

Trichorzin PA VI

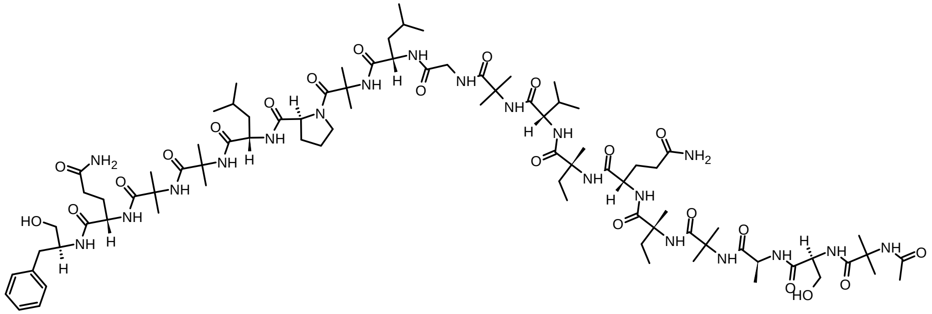

Trichorzin PA VIII

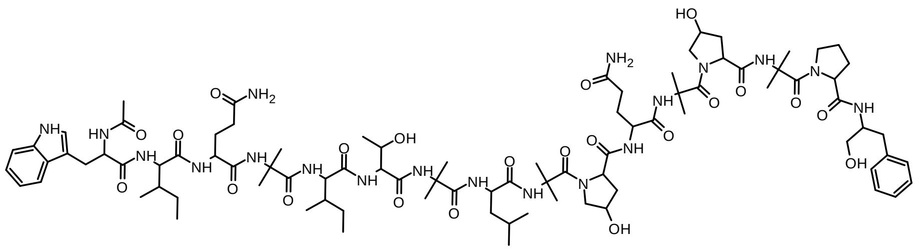

Zervamicin II-4

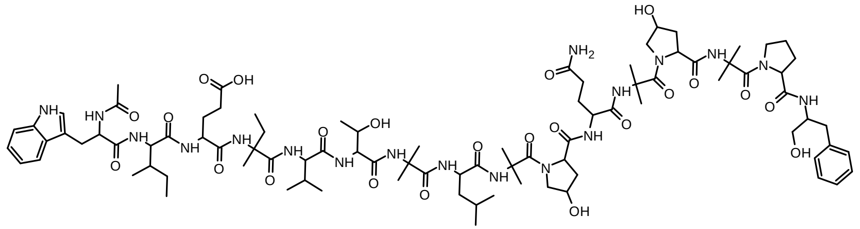

Zervamicin IB

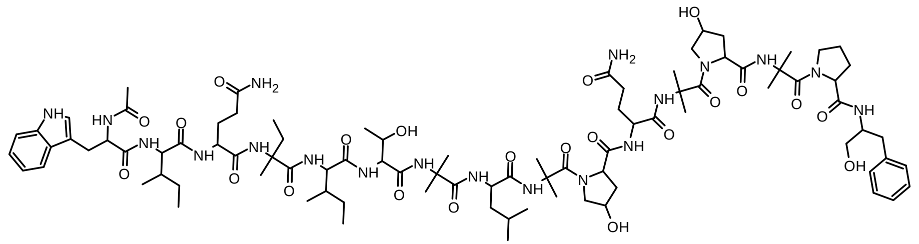

Zervamicin IIB

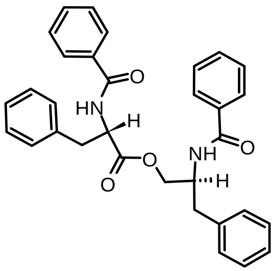

Asperphenamate

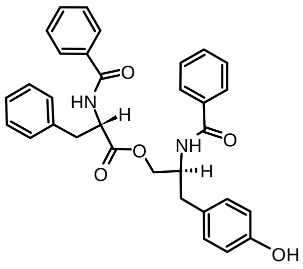

Asperphenamate analogue

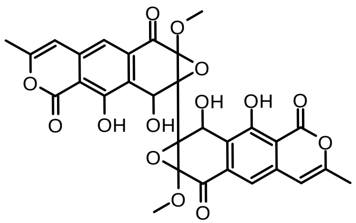

Xanthoepocin

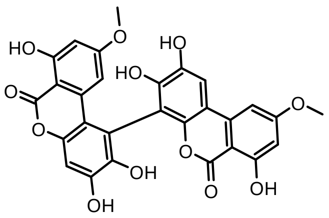

Verrulactone B

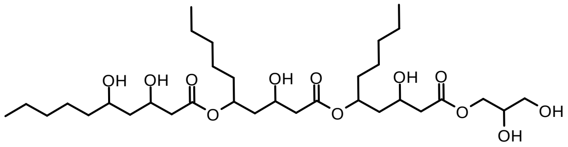

Glycerolliamocin A1

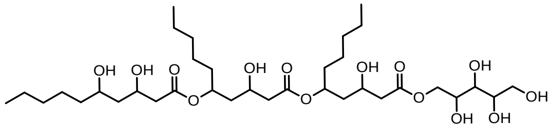

Arabinitol liamocin A1

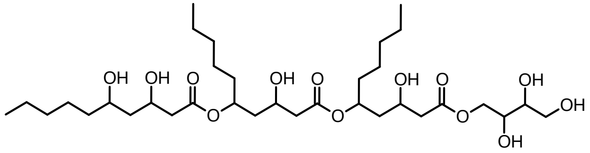

Threitol liamocin A1

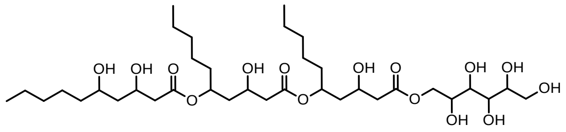

Liamocin A1

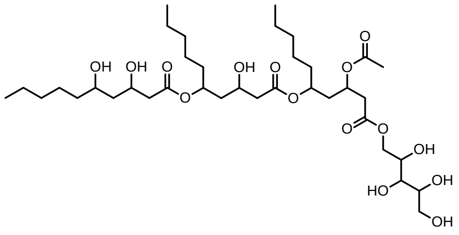

Arabinitol liamocin A2

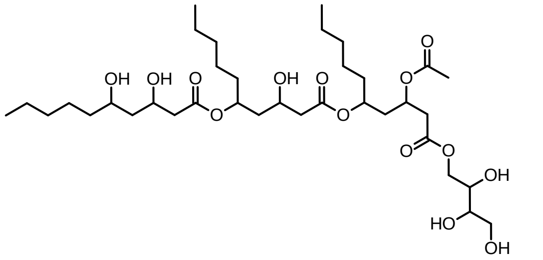

Threitol liamocin A2

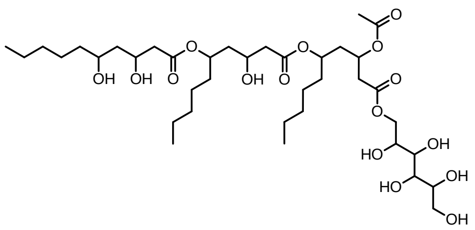

Liamocin A2

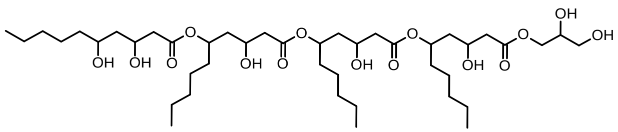

Glycerol liamocin B1

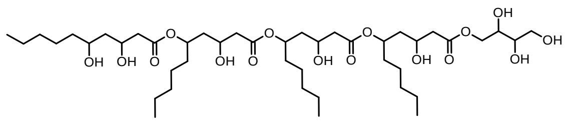

Threitol liamocin B1

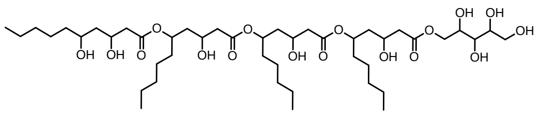

Arabinitol liamocin B1

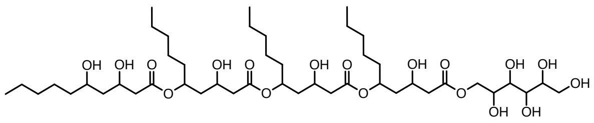

Liamocin B1

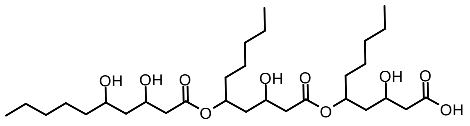
Exophilin A1

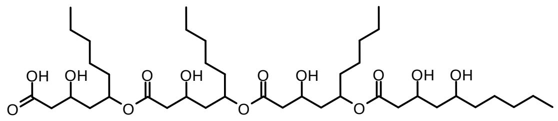
Exophilin B1

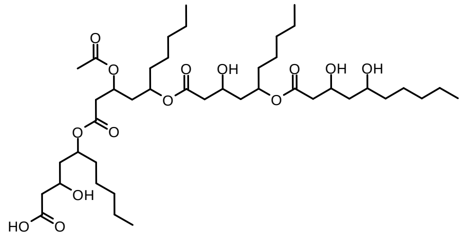
Halymecin A

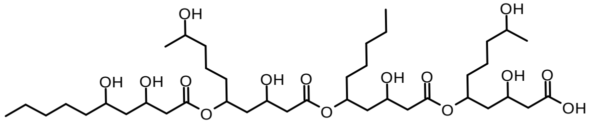
Halymecin D

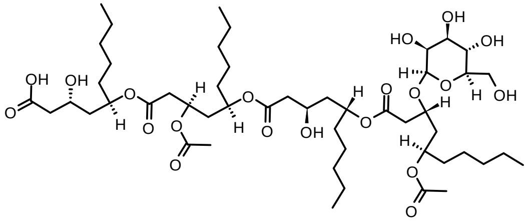
Halymecin F

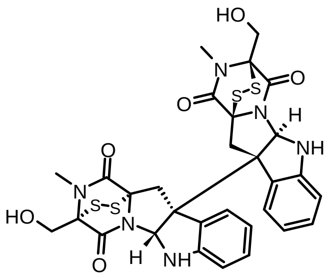
Chaetocin

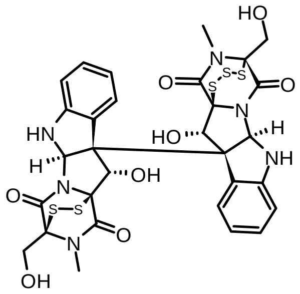
Chetracin B

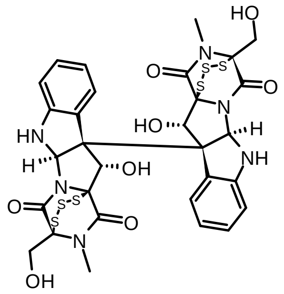
Chetracin C

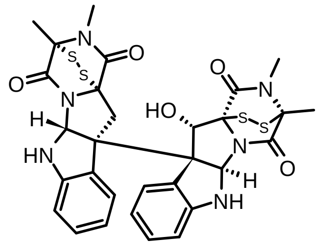
11'-deoxyverticillin A

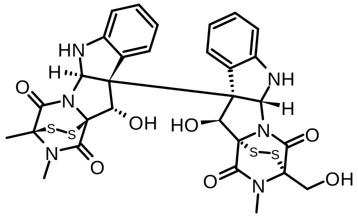
Verticillin B

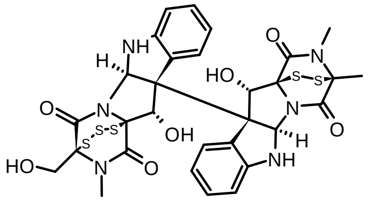
Verticillin C

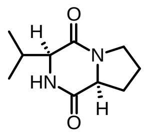

Cyclo(L-Pro-L-Val)

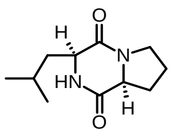

Cyclo(L-Pro-L-Leu)

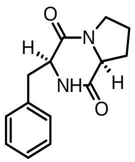

Cyclo(L-Phe-L-Pro)

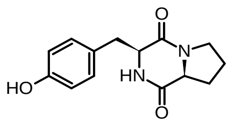

Cyclo(L-Tyr-L-Pro)

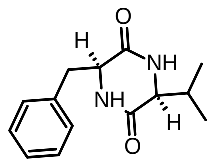

Cyclo(L-Val-L-Phe)
